# Nanoscale Imaging and Microanalysis of Ice Age Bone Offers New Perspective on “Subfossils” and Fossilization

**DOI:** 10.1101/2023.12.05.570041

**Authors:** Landon A. Anderson

**Affiliations:** Department of Biology, North Carolina State University, Raleigh, North Carolina, United States of America

**Author notes:** Corresponding author: Landon A. Anderson.

**Keywords:** Fossil, subfossil, biomolecular histology, collagen, bone, electron microscopy, blood vessels, diagenesis

## Abstract

The 3-D structure and organization of type-1 collagen protein and vasculature for a set of ancient permafrost bones is extensively documented at the nanoscale (up to 150,000× magnification) for the first time. The chemical mapping technique ToF-SIMS is additionally used to directly localize chemical signal to these structures; C:N and isotope measurements are also reported for the bulk organic bone matrix. These analyses test the hypothesis that biomolecular histology of collagen and vasculature from the permafrost bones supports their taphonomic classification as “subfossils” rather than “fossils”. Results indicate the original collagenous scaffolding and vasculature are still present, the former of which is well-preserved, thus supporting the hypothesis. This study is the first to taphonomically classify a set of pre-Holocene bones as “subfossils” based on the preserved state of their biomolecular histology. These methods can be readily expanded to specimens of warmer thermal settings and earlier geologic strata. Doing so has potential to establish/formalize at what point a bone has been truly “fossilized”; that is, when it has transitioned from “subfossil” status to being a true “fossil” bone. This will elucidate the fossilization process for ancient vertebrates and lead to a deeper understanding of what it means to be a “fossil”.

## Introduction

A sizeable body of evidence suggests that biological “soft” tissues can preserve during fossilization, even into the early Neogene, Paleogene, and Mesozoic. Organisms trapped within fossil resins, for example, are renowned for their life-like preservation [1], and ancient remains of pigmented skin [2, 3], blubber [3], cuticular and chitinous coverings [4, 5], and even internal organs [3, 6] have all been reported. Even specimens preserving only biomineralized remains, such as bones and/or teeth, are reported to preserve osseous soft tissues including collagenous fibers, cells, and vascular tissue [7–10].

While pre-Pleistocene specimens can preserve soft tissues and cells, relatively little is known regarding how the chemical processes responsible for this phenomenon progress [11, 12]. Presumably, the soft tissues of most “subfossil” specimens (defined here as not being fully “fossilized”; see Comment 1 of Supplemental Discussion) are eventually degraded during fossilization, preserving only the mineral portions [13–16]. Soft tissues that avoid degradation undergo chemical transformation towards a more recalcitrant state [12, 17, 18]. For example, soft tissues of specimens as recent as the Pliocene generally exhibit extensive *in-situ* polymerization and/or carbonization [19–22]. Additionally, degree of ancient DNA and protein sequence recovery exhibits substantial drop-offs for strata dated prior to ∼0.13-0.24 Ma [23] and ∼0.8-1.0 Ma [24–26], respectively. Combined, this suggests diagenetic reactions progress significantly within soft tissue specimens as the late/mid-Pleistocene transitions into the early Pleistocene/Pliocene. By the Pliocene, the biomolecules of most soft tissues would either be chemically transformed into organic, diagenetic macromolecules, or degraded and potentially replaced with recrystallized mineral [12, 15, 16, 19–22]. However, empirical observations of the progression of these diagenetic reactions in soft tissue specimens spanning the Pleistocene into the Pliocene have generally, to this point, not been reported [11, 12, 27]. In fact, the point at which a “subfossil” bone is considered truly “fossilized” and thus a fossil, is poorly defined in the primary literature [28–33]. This is significant, as understanding the changes biological tissues undergo during fossilization is linked to predicting a specimen’s potential for harboring sequence-able DNA and proteins [11, 12, 27].

Analysis of bone biomolecular histology has recently been proposed for addressing the above stated gaps in understanding regarding the fossilization process. Biomolecular histology refers to histological structure that is the direct manifestation of constituent biomolecules [27]. Cellular membranes, which are composed primarily of phospholipids along with various associated proteins and sterols [34], are one example of biomolecular histology. Another would be the banded fibrils of the collagenous scaffolding of bone; these banded fibrils are the direct, structural manifestation of type-1 collagen sequences [35, 36]. Biogenic minerals, including the bioapatite of bone, are strictly excluded from this definition. A biomolecular histological approach would thus report how the morphology and chemistry of such structures compare for specimens spanning the geologic record. Bone specifically is suggested as a model tissue because its non-mineral portion (defined herein as organic bone matrix (OBM), includes the type-1 collagenous scaffold and associated cellular portions such as vasculature, osteocytes, nerve cells, etc.) consists overwhelmingly (>90%) of a single biomolecule, type-1 collagen [35, 36]. This simplifies comparisons of ancient OBM against modern-day collagen controls and standards. Furthermore, fossil bone specimens are often more accessible for molecular analyses relative to rarer, non-biomineralized remains, and soft tissues have been readily reported to preserve within ancient bone specimens from a variety of sedimentary contexts [7, 9, 10, 20, 37–39].

A pilot study is presented here to demonstrate morphological and chemical approaches to studying the biomolecular histology of ancient bone. This will provide an initial dataset for comparing bone biomolecular histology across the late/mid-Pleistocene into the Pliocene/Miocene. Permafrost bones are specifically chosen for this initial investigation because they are well studied regarding gross biochemical preservation. Genomic studies of permafrost bone extracts [40–43], as well as some proteomic [40, 44, 45] and isotopic investigations [46–48], have reported excellent biochemical preservation. DNA fragments and a diversity of proteins are generally reported [23, 40, 44]. Sequence coverage for type-1 collagen [40, 44, 45] is often comparable to that of extant bone [49, 50]. This may suggest the original biological structure of many permafrost bones is largely intact and not fully “fossilized”. Yet, the term “fossil” is still often used to describe such specimens [40, 51–54] (see Comment 1 of Supplemental Discussion) and the preserved state of their biomolecular histology remains ambiguous within the primary literature. Indeed, direct investigation of underlying biomolecular histology is poorly documented for most Pleistocene specimens, excluding the exceptional cases of frozen Pleistocene “mummies” [55–58]. A few studies have reported images for OBM of latest Pleistocene bones using transmission electron microscopy (TEM) showing 2-dimensional sections of collagen fibril banding [59–62]. A recent study even reported nanoscale, 3-D imaging of microstructures within the bones of Silurian and Devonian fish; however, the reported images were of empty osteocyte lacunae lacking any original biomolecular tissue [63].

This study is the first to present extensive nanoscale (up to 150,000× magnification) imaging data characterizing the 3-dimensional structure and organization of type-1 collagen protein fibers and vasculature for *any* ancient bone (two preliminary images of ancient collagen at 50,000× mag. were presented in Anderson 2022 [27]). Time-of-flight secondary ionization mass spectrometry (ToF-SIMS) is also used to directly investigate the preserved chemical state of these structures. This is significant because such chemical data is localized to the specimen’s biomolecular histology itself [64], as shown in the presented microscope images. Past studies have instead generally used homogenized extracts which preclude localizing chemical signal to biomolecular histology [27]. Finally, carbon:nitrogen (C:N) ratios are reported for the various specimens herein to allow for direct comparison against previous studies of permafrost bone [46–48]. The atomic C:N ratio, as well as %C and %N by mass, have historically been commonly used proxies for collagen preservation and are practical for comparing specimens across studies [65]. These analyses will test the hypothesis that the biomolecular histological preservation of collagen fibers and vascular tissue for permafrost bone is consistent with prior genomic, proteomic, and isotopic data; that is, these permafrost specimens are not fully “fossilized” and should be considered “subfossils”. This study is the first to formally distinguish between “fossils” and “subfossils” based on a given specimen’s biomolecular histology. Further, the above methods can be readily applied in future studies to ancient bone specimens from temperate and subtropical thermal settings, as well as earlier geologic timepoints. Doing so will elucidate the diagenetic changes OBM undergoes during fossilization and facilitate recovery of ancient biomolecules [12, 27].

## Results and Discussion

### Permafrost Specimens and Elemental/Isotopic Analysis

The Pleistocene permafrost specimens comprise six bones recovered from the Little Blanche Creek and Irish Gulch localities of the Yukon Canadian territory (Figure 1 (B1-G1)). The bones were each recovered as isolated, disarticulate remains which is typical for many Pleistocene specimens from eastern Beringia [52, 66]. Notably, the *B. priscus* tibia from Irish Gulch preserved remnant soft tissue (potentially ligaments and tendons, or even remnant musculature) still attached to the bone’s external surface. The Beringian permafrost specimens were overall more brittle than the extant controls (Figure 1(A1) and Figure S1(A and F)), and external bone portions were generally discolored and somewhat friable. Care was taken to sample only the internal cortical bone layers which displayed minimal discoloration and brittleness. Post-demineralization, extant control OBM was stiff and rubber-like, and sterile razor blades were necessary to cut smaller sections. In comparison, the demineralized permafrost specimen OBM readily frayed into smaller fibers and was soft and pliable.

**Figure 1.**
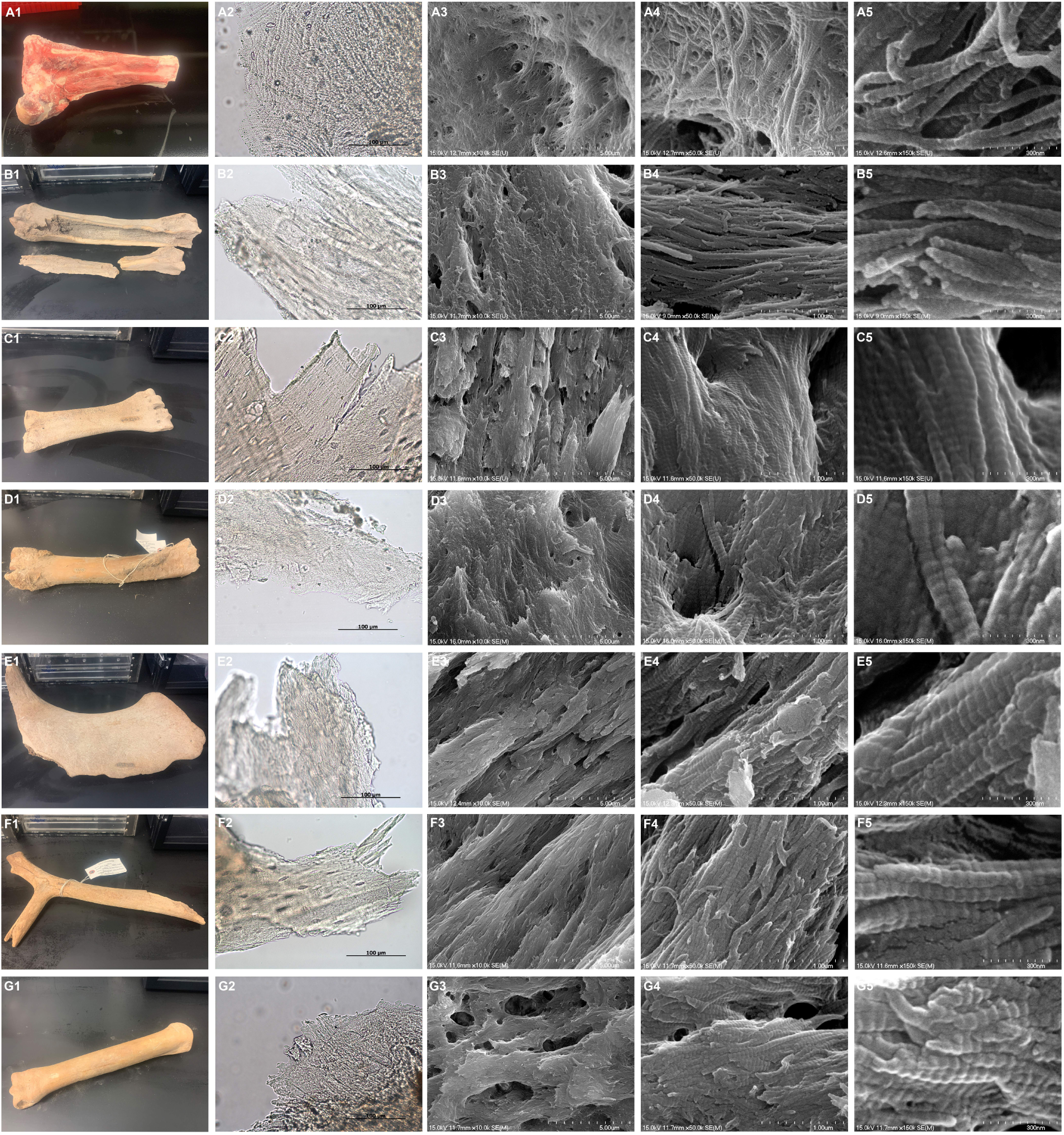
Images of permafrost and extant organic bone matrix (OBM). First shown are macroscopic images of the mineralized bone specimens (A1-G1). Then, transmitted light microscope imaging depicts the microscale, fibrous structure of demineralized OBM sections taken from the bone specimens (A2-G2). Lastly, scanning electron microscope (SEM) images of increasing magnification reveal how the fibrous structure shown by the transmitted light images is formed of nanoscopic type-1 collagen fibrils (A3-G5). **(A1-A5)** Extant B. taurus long bone. **(B1-B5)** YG 610.2365 (B. priscus radius). **(C1-C5)** YG 610.2363 (B. priscus metatarsal). **(D1-D5)** YG 126.115 (B. priscus tibia). Desiccated soft tissue can be observed on the exterior surface of this specimen. **(E1-E5)** YG 610.2397 (M. primigenius innominate). **(F1-F5)** YG 610.2305 (R. tarandus antler). **(G1-G5)** YG 610.2364 (E. lambei metatarsal).

Stable isotope and C:N measurements obtained from demineralized bone samples are shown in Table 1. Both the permafrost and extant specimen C:N ratios were within the range 2.9-3.6, which has historically been accepted as indicating well-preserved collagen. Values for %C and %N (by mass) ranged from 41.0-46.0% and 15.2-17.1%, respectively, which is likewise consistent with the collagenous scaffolding having undergone minimal diagenesis [46–48, 65]. Measurements of δ^13^C and δ^15^N are also reported; while the obtained values are consistent with prior values obtained from permafrost megafaunal bones, they are more so related to organismal diet/trophic-level and thus are not further discussed [46–48].

**Table 1.** Stable isotope and C:N measurements.

| **Specimen Number** | **Sample** | **C:N (atomic)** | **%C (by mass)** | **%N (by mass)** | **$\delta^{13}\text{C}$ (‰) VPDB** | **$\delta^{15}\text{N}$ (‰) AIR** |
| --- | --- | --- | --- | --- | --- | --- |
| **n/a** | *B. taurus* long-bone | 3.1 | 41.0 | 15.3 | -20.39 | 4.96 |
| **n/a** | *A. mississippiensis* long-bone | 3.0 | 44.6 | 17.1 | -17.45 | 8.75 |
| **n/a** | *S. camelus* long-bone | 3.4 | 44.1 | 15.3 | -20.31 | 4.93 |
| **YG 126.115** | *B. priscus* tibia | 3.3 | 46.0 | 16.2 | -19.76 | 6.46 |
| **YG 610.2305** | *R. tarandus* antler | 3.1 | 44.4 | 16.7 | -17.19 | 2.00 |
| **YG 610.2363** | *B. priscus* metatarsal | 3.1 | 45.2 | 16.8 | -19.84 | 5.53 |
| **YG 610.2364** | *E. lambei* metatarsal | 3.1 | 44.2 | 16.4 | -19.77 | 5.03 |
| **YG 610.2365** | *B. priscus* radius | 3.1 | 41.8 | 15.6 | -19.09 | 4.22 |
| **YG 610.2397** | *M. primigenius* innominate | 3.2 | 41.0 | 15.2 | -20.38 | 7.21 |

### Type-1 Collagen Protein Fibers

Demineralized OBM sections exhibited an overall fibrous structure when visualized with transmitted light microscopy (Figure 1(A2-G2); Figure S1(B and G)). Higher magnification images (Figure 1(A3-G5); Figure S1(C-E and H-J)) confirmed this fibrous structure to be the manifestation of collagen fibers, themselves consisting of type-1 collagen fibril bundles. For extant bone collagen, these individual fibrils consist of parallel-packed chains of peptide helices (each helix consists of two α1 peptides and a single α2 peptide, intertwined). The peptide helices are bound one-to-another near their termini via covalent crosslinks; these chains are packed parallel to each other down the length of the fibril and with small longitudinal offsets that result in alternating regions of tightly/loosely packed peptides (see Figures 1 and 4 of Minary-Jolandan and Yu 2009 [67]; Figure 3 of Orgel et al. [68]). These alternating regions of density manifest as the ∼67 nm banding pattern characteristic of extant type-1 collagen fibrils [35, 36, 67–69]. Such a pattern is clearly present throughout the high magnification images of both the permafrost and extant OBM. The presence of this banding in the permafrost specimens indicates the collagen fibrils are, to an extent, well preserved. Chemical degradation has not occurred to substantially disrupt the packed structure of the peptide helices within an individual fibril [69]. The arrangement of the fibrils themselves with respect to one another does, however, potentially indicate some degree of chemical degradation. The fibrils of the Pleistocene specimens potentially display a somewhat “looser” association with one another relative to those of the extant controls, which are tightly intertwined. This suggests that the intermolecular forces holding the bundled fibrils together have been disrupted somewhat, perhaps due to chemical degradation of the constituent amino acids [40, 44]. This would also explain why, post-acid demineralization, the OBM of the permafrost samples tended to easily fray in comparison to the stiffer, rubber-like matrix of the extant controls.

**Figure 2.**
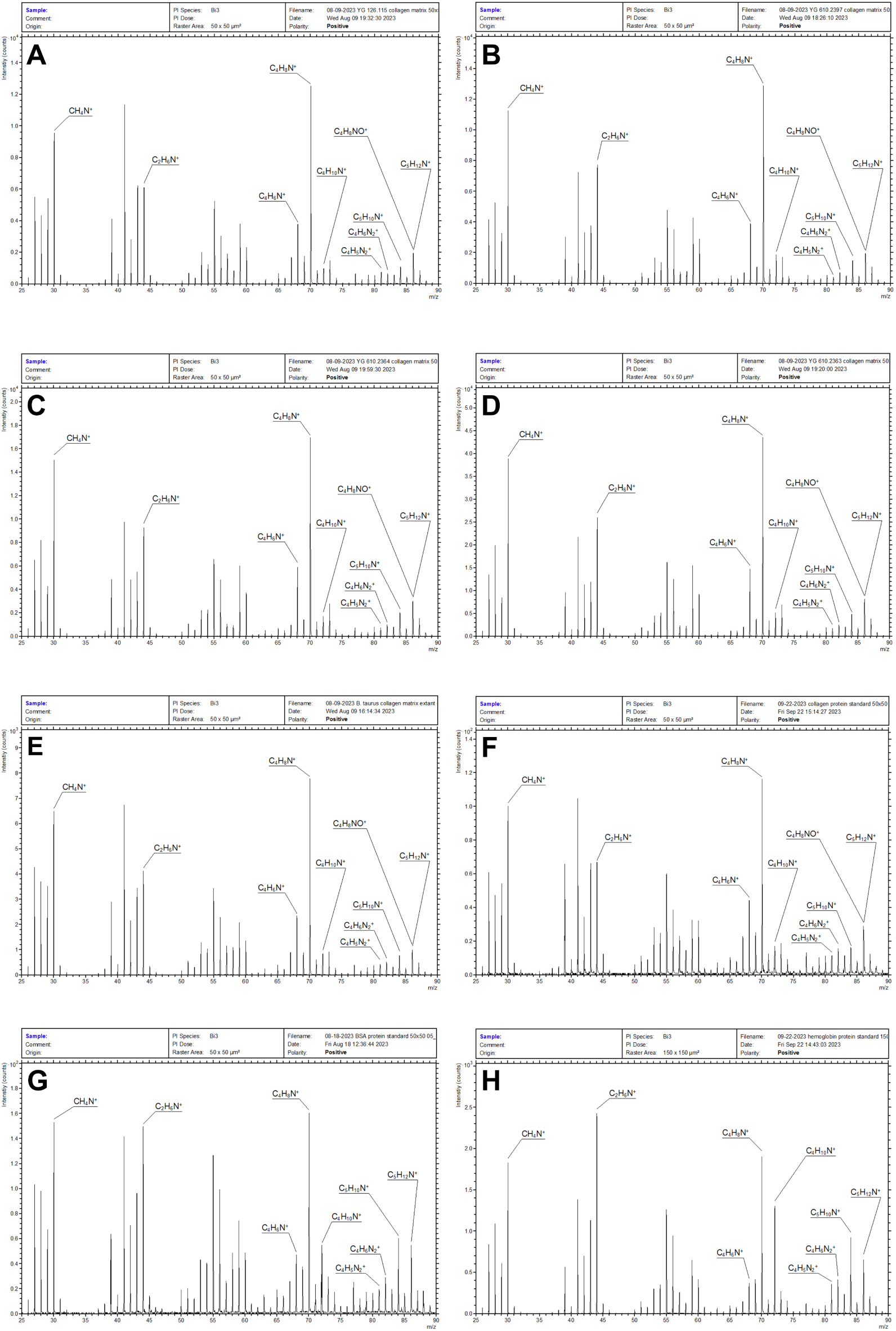
ToF-SIMS spectra of permafrost and extant OBM, and of purified protein standards. Secondary ion peaks for CH_4_N^+^, C_2_H_6_N^+^, C_4_H_6_N^+^, C_4_H_8_N^+^, C_4_H_10_N^+^, C_4_H_5_N_2_^+^, C_4_H_6_N_2_^+^, C_5_H_10_N^+^, and C_5_H_12_N^+^ are labeled for all spectra; the range of each spectrum is limited to 25-90 m/z to aid readability. The ion CH_4_N^+^ is highly abundant across all the protein-related spectra [71–73]. Both C_4_H_6_N^+^ and C_4_H_8_N^+^ (proline [71–73]) present as prominent peaks for the collagenous spectra; this peak is less intense in the hemoglobin standard. At a nominal m/z of 86, substantial peaks for both C_4_H_8_NO^+^ and C_5_H_12_N^+^ are present for all collagenous spectra (see Figure 3). In contrast, the secondary ion C_4_H_8_NO^+^ (hydroxyproline [77]) is not present in the BSA and hemoglobin spectra, thus only C_5_H_12_N^+^ is labeled. The BSA and hemoglobin spectra also exhibit elevated intensities for C_4_H_10_N^+^ (valine [71–73]) and C_5_H_10_N^+^ (leucine/lysine [71–73]) relative to the collagenous spectra. **(A)** YG 126.115 OBM spectrum. **(B)** YG 610.2397 OBM spectrum. **(C)** YG 610.2364 OBM spectrum. **(D)** YG 610.2363 OBM spectrum. **(E)** Extant B. taurus OBM spectrum. **(F)** Purified type-1 collagen protein standard (bovine) spectrum. **(G)** Purified BSA protein standard spectrum. **(H)** Purified hemoglobin (porcine) protein.

**Figure 3.**
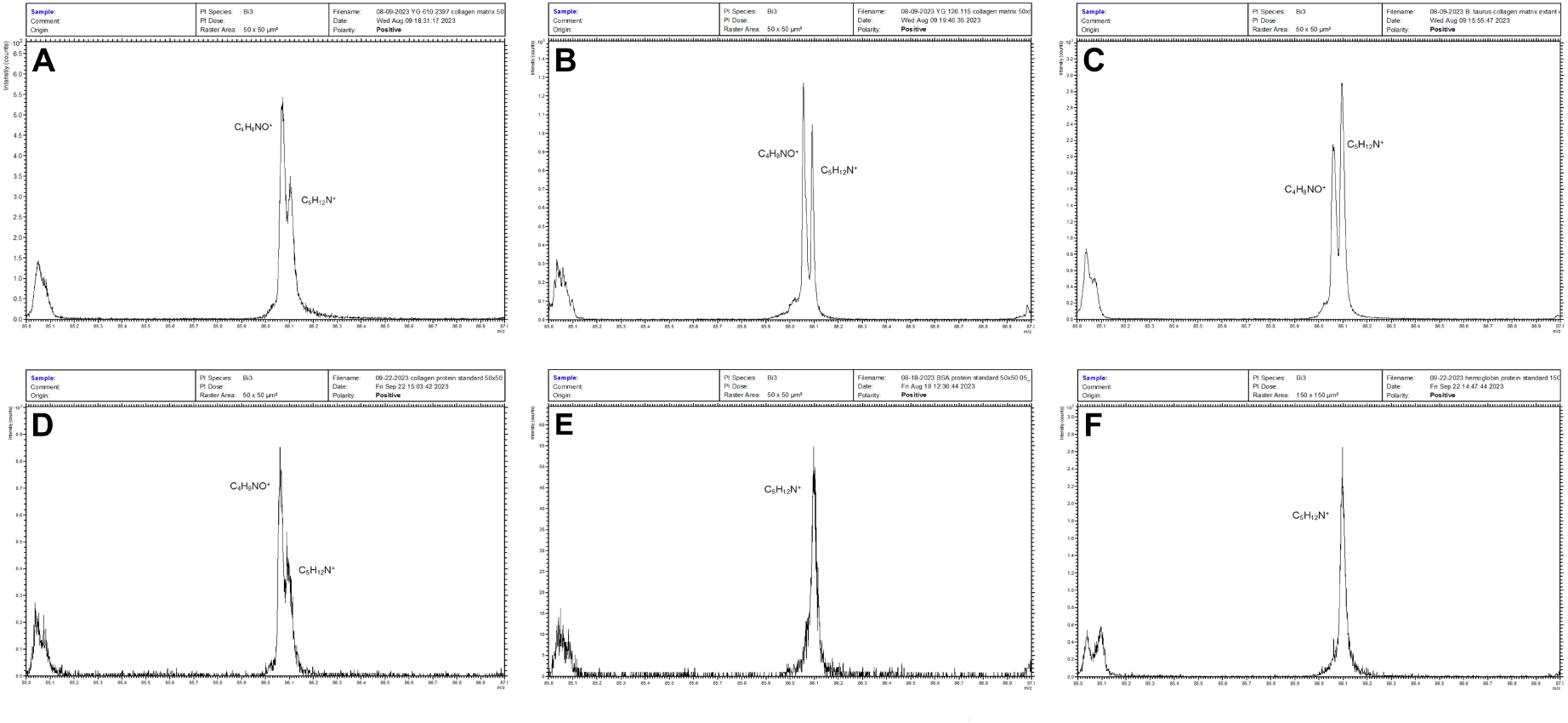
Hydroxyproline-related secondary ion peak for collagenous specimens and protein standards. Enlarged view of secondary ion peaks with a nominal m/z of 86. In studies of proteinaceous materials, the ion C_4_H_8_NO^+^ is primarily formed by fragmentation of hydroxyproline [77]. A peak corresponding to C_4_H_8_NO^+^ is present for the OBM spectra (A-C) and the type-1 collagen protein standard (D). This peak is absent in both the BSA and hemoglobin standard spectra (E-F). **(A)** YG 610.2397. **(B)** YG 126.115. **(C)** Extant B. taurus. **(D)** Purified type-1 collagen protein standard (bovine). **(E)** Purified BSA protein standard. **(F)** Purified hemoglobin (porcine) protein standard.

**Figure 4.**
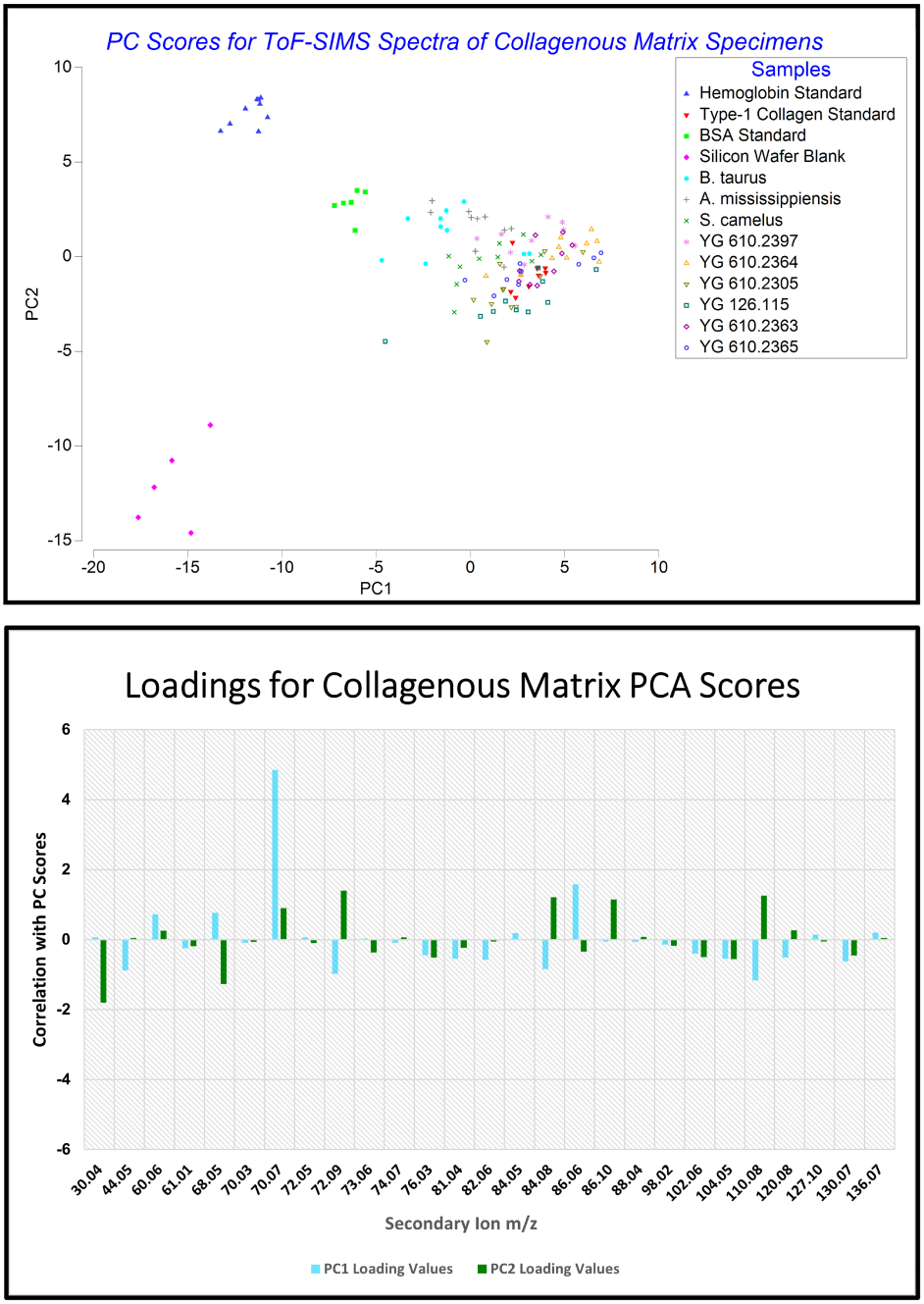
Score and loadings plots for OBM and protein standard ToF-SIMS spectra. Determining the relative abundance of a given secondary ion (and thus amino acid) for a given spectra using PCA requires comparison of score plot values with the loadings plot values. For example, the PC1 loading value for an m/z of 70.07 (C_4_H_8_N^+^) is 4.85. This indicates a strong positive correlation for C_4_H_8_N^+^ peak intensity with higher PC1 scores. Thus, sample spectra with PC1 scores that are more positive exhibit a greater relative C_4_H_8_N^+^ peak intensity. The collagenous spectra exhibited strong positive correlations for PC1 with the ions C_4_H_8_N^+^ (70.07, proline) and C_4_H_8_NO^+^ (86.06, hydroxyproline), and some notable negative correlations with C_4_H_10_N^+^ (72.09, valine), C_4_H_5_N_2_^+^ (81.04, histidine), C_4_H_6_N_2_^+^ (82.06, histidine), C_5_H_10_N^+^ (84.08, leucine and lysine), and C_5_H_8_N_3_^+^ (110.08, histidine).

Chemical characterization of type-1 collagen fibrils as shown in Figure 1 was accomplished with ToF-SIMS. ToF-SIMS is a chemical mapping technique that rasters a beam of primary ions in a square, grid-like manner across a sample surface. Under static conditions (used in this study), the ToF-SIMS primary ion beam ablates only the uppermost ∼1-2 nm of a given sample’s surface. A small portion of these ablated atoms/molecules ionize, and it is these secondary ions the TOF analyzer/detector sorts into and records as mass spectra. A unique spectrum is recorded for each raster point in the square, grid-like analysis area. The sum of these spectra for a given analysis area, or a selected subregion within it, forms a single mass spectrum representative of the given region’s overall secondary ion yield [64]; this summed spectrum effectively functions as a chemical “fingerprint” that can be compared against other analysis areas/samples [70].

ToF-SIMS spectra collected for both permafrost and extant collagen fibers displayed high intensities for secondary ions characteristic of proteins [71–73] (Figures 2-3; Figures S4-S16). The distribution and intensities of their secondary ions closely mirrored the type-1 collagen standard spectra. Relative to the hemoglobin and BSA standards, yields for C_4_H_6_N^+^ (68.05), C_4_H_8_N^+^ (70.07), and C_4_H_8_NO^+^ (86.06) secondary fragment ions were particularly high. In SIMS protein analyses, C_4_H_6_N^+^ and C_4_H_8_N^+^ are generally produced by the amino acid proline [71–73] (C_4_H_8_N^+^ also corresponds to arginine [73], which is somewhat common in type-1 collagen [74–76]), and C_4_H_8_NO^+^ by hydroxyproline [77] (see Figure 3 for the hydroxyproline-related ion peak); both amino acids are enriched within the type-1 collagen sequence. Additionally, glycine makes up approximately one-third of type-1 collagen [74–77]. The CH_4_N^+^ (30.04) ion formed by SIMS fragmentation of glycine is present in high abundances for the permafrost and extant OBM. However, this ion is ubiquitous to most amino acids and generally shows limited variation between spectra of different proteins, as was observed in this study [71–73]. These three amino acids are the most abundant within type-1 collagen and are integral to forming the tight turns of its helical peptides [67, 68, 74, 78]. The substantial presence of these ions, as well as overall spectral agreement with the purified type-1 collagen standard, evidences the permafrost collagen scaffolding is relatively well preserved. The BSA and hemoglobin spectra also exhibit elevated intensities for C_4_H_10_N^+^ and C_5_H_10_N^+^ relative to the collagenous spectra. These ions are typically produced by valine and lysine/leucine [71–73], respectively, which are less prevalent in the type-1 collagen sequence relative to BSA and hemoglobin [74–76].

Peak areas for 27 amino acid-related ions [71–73, 77] (Table S1) were calculated for each protein spectrum, placed in a data matrix, and analyzed with principal component analysis (PCA) (Figure 4). PC1 scores for the collagenous sample spectra were generally more positive (56.5% explained variation), and PC2 scores (23.2% explained variation) were generally closer to zero. Loading values for C_4_H_8_N^+^ and C_4_H_8_NO^+^ show a substantial positive correlation with scores for PC1, supporting a relatively high abundance of proline and hydroxyproline within the collagenous sample spectra. Additionally, several secondary ions, especially those corresponding to valine (C_4_H_10_N^+^, 72.09) and lysine/leucine (C_5_H_10_N^+^, 84.08), correlate negatively with PC1 scores. This was also the case for C_4_H_5_N_2_^+^ (81.04), C_4_H_6_N_2_^+^ (82.06), and C_5_H_8_N_3_^+^ (110.08), all of which correspond to histidine [71–73]. This agrees with the type-1 collagen sequence having a lower abundance of these amino acids relative to BSA and hemoglobin [74–76]. PC2 scores further separated the hemoglobin and collagenous spectra. PC2 loading values for CH_4_N^+^ and C_4_H_6_N^+^ were negatively correlated with the hemoglobin spectral scores, while C_4_H_10_N^+^, C_5_H_10_N^+^, C_5_H_12_N^+^, and C H N ^+^ demonstrated positive correlations; this likewise agrees with the known amino acid compositions for hemoglobin vs type-1 collagen [74–76].

Together, the C:N measurements, microscope images, and ToF-SIMS spectral data and PCA support that the permafrost collagen fibers share substantial morphological and chemical similarity with extant analogs. The type-1 collagen scaffolding of these permafrost bones has undergone limited diagenesis relative to the extensive *in-situ* polymerization/carbonization observed for soft tissue specimens of earlier geologic strata [18, 19, 21, 22, 79]. This agrees with prior studies on bulk permafrost bone extracts that have reported C:N ratios of ∼2.9-3.6, implying relatively minimal alteration of the collagen [46–48]. Likewise, paleoproteomic sequencing studies of permafrost bone generally report sequence coverage for both peptide chains of type-1 collagen (α1 and α2) >50-60% [40, 44, 45], roughly comparable to what is reported for extant bone [49, 50]. The lack of observed degradation within the type-1 collagen scaffolding supports these permafrost bones should be considered “subfossils”. While extensive alteration of the permafrost collagen was not detected herein, this is unlikely to be the case with specimens from warmer thermal settings and/or earlier geologic strata. Future studies on such specimens can document OBM structural alteration via electron microscopy, as shown by the preliminary data in Figure 1 of Anderson 2022 [27]. For ToF-SIMS, indicators of chemical diagenesis would likely include alteration of the intensity and distribution of protein-related ion peaks, as well as an increased presence of secondary ions related to aliphatic and aromatic hydrocarbons and heterocycles [3, 12, 80].

### Vascular Tissue

“Blood vessels” were isolated/obtained for all specimens by enzymatic digestion of the OBM type-1 collagen scaffolding (Figure 5(A1-G1); Figure S2(A and E)). The vessels (interpreted here as representing basal endothelium [81]) were generally clear and flexible and readily sheared in response to excessive turbulence or mechanical force. Some loose, exogenous sedimentary matter was present in portions of the vessel lumens for the permafrost specimens, and fungal hyphae were found to have colonized the vessel lumens of YG 610.2363 and YG 610.2365. Electron microscope imaging revealed a substantial divergence in structure between the permafrost vessels and the extant controls (Figure 5(A2-G4); Figure S2(B-D and F-H)). The vessel “endothelium” of the permafrost specimens consisted of a single, thin “membrane”. In contrast, the extant vessel endothelium exhibited more substantial structure, possibly portions of the cytoskeleton as well as cytosolic membranes [34, 82]. This disparity suggests that non-collagenous, cellular portions have undergone extensive diagenesis within the permafrost-frozen bones. One exception was YG 126.115 which possessed vessel endothelium somewhat closer in structure to the extant specimens, rather than a single, thin “membrane”. Osteocytes (bone cells) were also observed for YG 126.115 as well as for each extant specimen; of the Little Blanche Creek specimens, only YG 610.2364 was observed to harbor osteocytes (Figure S3). Additionally, YG 126.115 possesses soft tissue preserved on the bone’s external surface. It is the only specimen recovered from the Irish Gulch locality (the others being from Little Blanche Creek), suggesting a potential disparity in diagenetic processes between the two sites.

**Figure 5.**
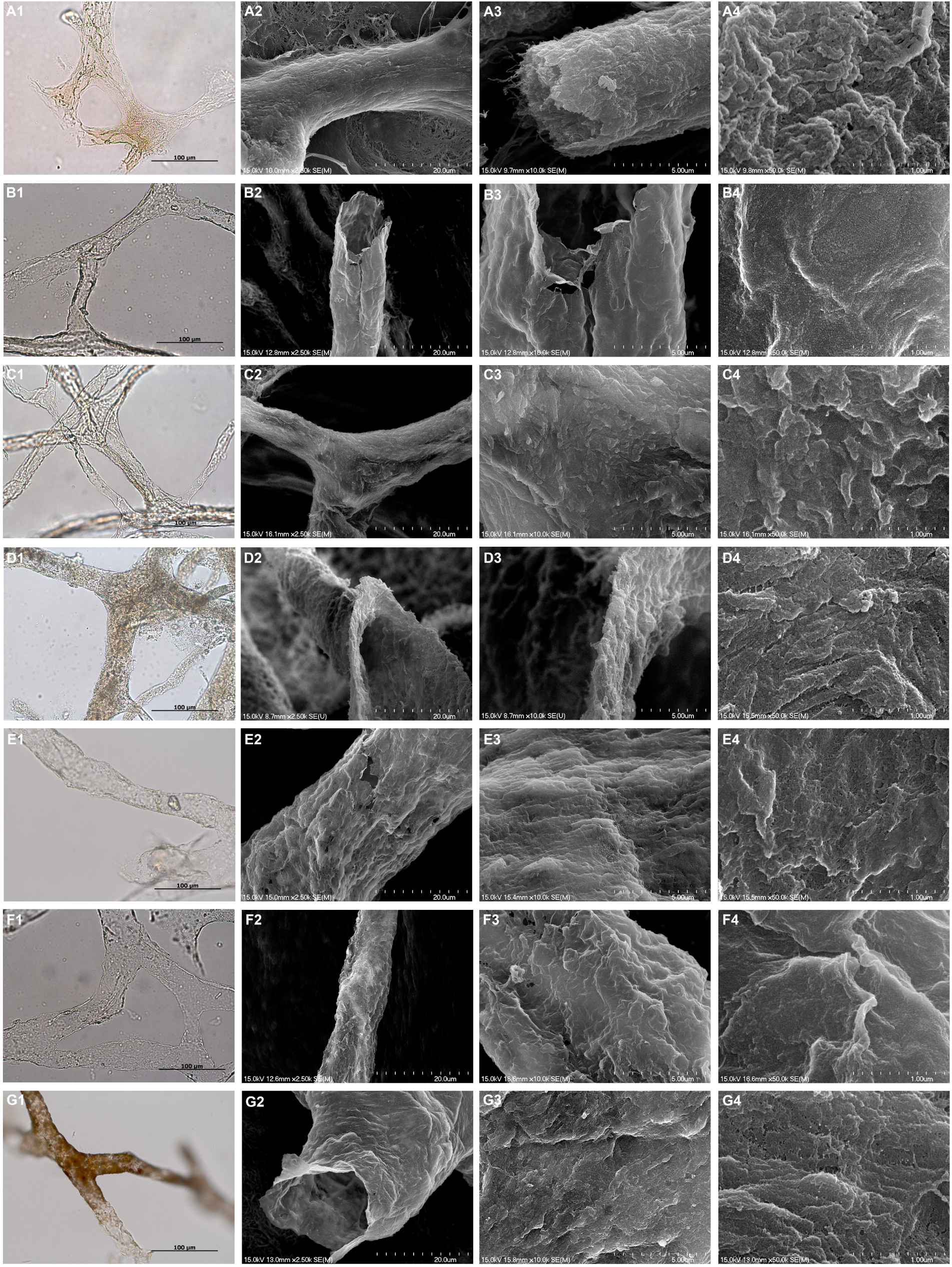
Images of permafrost and extant vascular tissue. (A1-A4) Extant B. taurus. The capillary lumen of (A3) reveals the fibrous and membranous structure of the endothelium. **(B1-B4)** YG 610.2365 (B. priscus radius). The vessel endothelium (B3) presents as a single, thin “membrane” lacking significant structure. This contrasts with the proteinaceous fibers and membranous portions of the extant B. taurus capillary (A3). **(C1-C4)** YG 610.2363 (B. priscus metatarsal). Images of the vessel lumen are not shown but were comparable to the other Little Blanche Creek specimens. **(D1-D4)** YG 126.115 (B. priscus tibia). The endothelial membrane (D3) is thin but presents with more structure relative to the Little Blanche Creek specimens. **(E1-E4)** YG 610.2397 (M. primigenius innominate). An example of loose sediment in some of the vessel lumens is shown in E1. Additionally, the small tear of the vessel membrane in (E2) shows similar morphology with images (B2-B3). **(F1-F4)** YG 610.2305 (R. tarandus antler). **(G1-G4)** YG 610.2364 (E. lambei metatarsal). The vessel “membrane” morphology in (G2) is comparable to vascular tissue for the other Little Blanche Creek specimens.

Negative ion ToF-SIMS spectra (Figure 6; Figures S17-S25) were collected for vascular tissue as shown in Figure 5. Protein-related peaks were not studied because the enzymatic digestion used to isolate the vessels introduces a potential source of non-specific peptide signal. In vessels from the extant controls, molecular ions (non-fragmentary secondary ions) corresponding to a variety fatty acids were present; this was especially the case for palmitic (C_16_H_33_O_2_^-^) and oleic (C_18_H_33_O_2_^-^) acids [83]. These two fatty acids tend to be the most abundant within extant mammalian tissue including bone [84, 85]. Molecular ions for stearic acid, as well as potentially palmitoleic, myristic, margaric, and linoleic acids were also consistently observed; these latter ions, however, may also be attributable to fragment ions of palmitic, oleic, and stearic acids [83]. For the Little Blanche Creek permafrost specimens, no secondary ions could be confidently attributed to fatty acids. Secondary ions corresponding to stearic, and particularly palmitic, acids were observed sporadically but generally of limited intensity. The preferential preservation of these fatty acids would be expected in ancient specimens; their lack of hydrocarbon alkene (-CH=CH-) bonds increases chemical stability [12, 86]. Previous lipidomic studies have likewise reported the presence of palmitic and stearic acids in ancient remains [45, 87, 88]. The limited mass resolution of the ToF mass analyzer, however, hinders identification of these “fatty acid” molecular ion peaks for most of the permafrost specimens due to their sporadic nature [89] (see Comment 2 of the Supplemental Discussion). This was compounded by many of these “peaks” for the ancient specimens expressing low signal relative to instrumental background noise. An exception was YG 126.115, which consistently exhibited ion peaks with similar distributions and intensity to those of the extant control fatty acid molecular ions. This agrees with the morphological disparities documented in Figure 5 and further suggests the cellular components of YG 126.115 have undergone limited chemical diagenesis relative to the Little Blanche Creek specimens.

**Figure 6.**
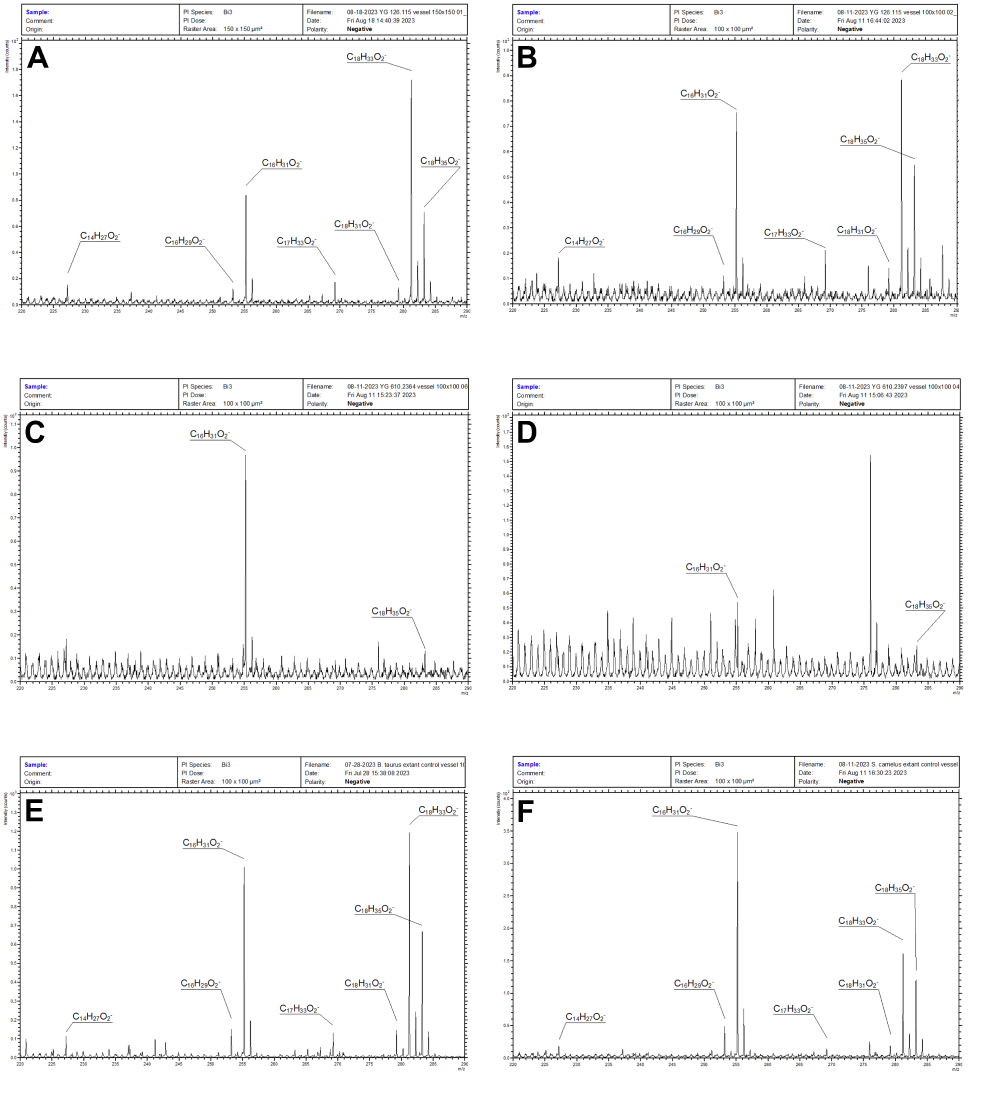
ToF-SIMS spectra of permafrost and extant vascular tissue. Both the YG 126.115 (A-B) and B. taurus/S. camelus extant control (E-F) vascular tissue spectra show similar peak distributions for m/z values corresponding to fatty acids. In contrast, the spectra for YG 610.2364 and YG 610.2397 (C-D) show few prominent peaks matching those of the extant controls in (E-F). Additionally, the secondary ion peaks in (C-D) are labeled for clarity only; the labeling is not necessarily an assertion that these secondary ions result from fatty acids. Making such an assertion for these two figured spectra is difficult due to their disparities with the extant controls, and the limited mass resolution of the ToF analyzer. **(A)** YG 126.115 vascular tissue spectrum. **(B)** YG 126.115 vascular tissue spectrum (separate replicate from (A)). **(C)** YG 610.2364 vascular tissue spectrum. **(D)** YG 610.2397 vascular tissue spectrum. **(E)** Extant B. taurus vascular tissue spectrum. **(F)** Extant S. camelus vascular tissue spectrum.

Figure 7 shows PC1 (70.9% explained variation; PC2 explained 25.5%; see Table S2 for selected peaks) scores for both YG 126.115 and the extant controls were generally negative and clustered together. However, the PC1 scores for these specimens also showed substantial variance and several “outliers” were observed. The loading values for PC1 indicate this was largely due to inconsistency of the palmitic and oleic acid molecular ion signal intensities. Fatty acid composition is known to differ throughout the cellular membranes of eukaryotic organisms [90] and potentially explains this observance. Compounding this, damage to the vessels may have exposed internal cytosolic membranes, exacerbating sample inhomogeneity. Such damage is likely attributable to the method of air-drying the vessels onto silicon wafers; the surface tension of the evaporating water droplet potentially damages the vessel membranes. Regardless, this is one limitation to analyzing structures that are chemically complex, such as cellular membranes, as opposed to the collagen fibers of OBM. In contrast, positive PC1 scores were consistently obtained for the permafrost specimens (minus YG 126.115). The loading value for the peak corresponding to palmitic acid shows a positive correlation with PC1 scores. This suggests that potential fatty acids preserved (if the ions do indeed result from fatty acids, which is itself questionable) within the permafrost vascular tissue are enriched with palmitic acid relative to extant specimens. Previous lipidomic studies of ancient mammalian tissues have also consistently reported this phenomenon [45, 87, 88, 91]. A potential explanation may be found in considering the predominance of palmitic and oleic acids within mammalian tissues during life [84, 85]. Post-mortem, palmitic acid, having a saturated hydrocarbon chain, would preferentially preserve over the unsaturated, more labile oleic acid [12, 86]. Stearic acid, the saturated analog of oleic acid, is also favored to preserve; however, initial concentrations of stearic acid in animal tissues tend to be lower relative to both palmitic and oleic acids [84, 85]. Thus, palmitic acid may be predicted to be the fatty acid species with the highest preservation potential within diagenetically altered mammalian tissues.

**Figure 7.**
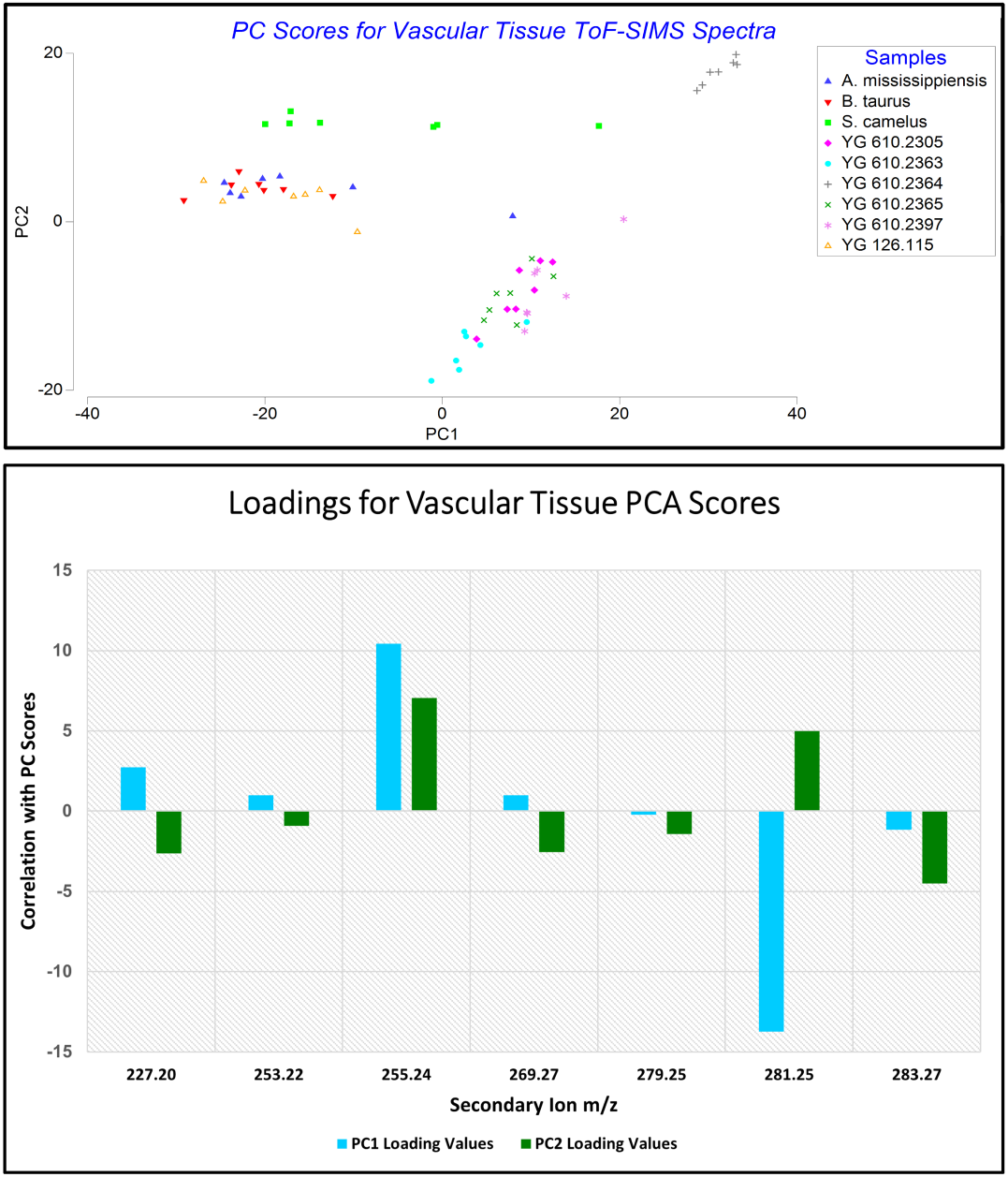
Score and loadings plots for vascular tissue ToF-SIMS spectra. The most predominant molecular ions observed in the extant/YG 126.115 spectra corresponded to palmitic (255.24 m/z) and oleic (281.25 m/z) acids. A majority of the extant/YG 126.115 scores fell within the range –27 to –9, indicating a higher abundance of oleic acid relative to palmitic acid. However, several outliers were observed in the more positive region of the score plot, making it difficult to draw conclusions regarding specific fatty acid prevalence for these specimens. In contrast, the Little Blanche Creek vascular tissue spectra yielded generally higher scores for PC1, indicating a positive correlation with “palmitic acid” molecular ion abundance (whether the observed peak for the Little Blanche Creek spectra represents palmitic acid, however, is questionable).

The data support that vascular tissue of the Little Blanche Creek specimens has undergone substantial diagenesis. Vascular structure is reduced to a single, thin “membrane”, and ion signal corresponding to fatty acids was detected sporadically (if at all) and with limited intensity. These findings suggest the original lipid bilayers have been disrupted, and much of the original cellular machinery is no longer present. This contrasts sharply with, for example, the frozen cells of one *M. primigenius* carcass in which the cellular mechanisms for DNA repair were able to be reactivated post-thawing [56]. As a comparison from this study, the YG 126.115 tibia demonstrated secondary ion distributions (for fatty acid m/z values) mirroring those of the extant control spectra; its vessels also presented with more structure relative to the other permafrost specimens. This suggests that bones preserved in permafrost vary widely regarding degree of cellular preservation. The freezing process largely arrests diagenetic reactions [92, 93]. Thus, these taphonomic disparities are likely attributable to events that occurred prior to freezing, or perhaps during a period of thawing. This is supported by the presence of desiccated soft tissue on the external surface of YG 126.115. For such tissue to preserve, relatively rapid freezing likely must have occurred. The Little Blanche Creek bones may have taken longer to be incorporated into permafrost, or local conditions at the Little Blanche Creek site may have promoted diagenesis prior to freezing [93, 94]. Such disparities in preservational state for permafrost bones have previously been explored [93, 95] but until this study had not been documented at the nanoscale in terms of biomolecular histology.

Regarding chemical diagenesis, molecular ions for fatty acids were generally absent from the Little Blanche Creek vessel spectra, suggesting many of the original biomolecules have undergone chemical alteration. Direct examination of fragmentary organic ions in the lower m/z spectral regions (∼20-100 m/z, spectral region not shown) didn’t reveal any particularly striking differences between the Little Blanche Creek and extant specimens. Tissues that preserve in pre-Pleistocene strata are known to be highly cross-linked and possess a higher carbon content relative to extant tissues [12, 18, 96]; however, the C:N ratios and %C and %N measurements reported herein likely rule out extensive carbonization. This is unsurprising given the colder thermal setting and somewhat recent geologic timepoint. Extensive crosslinking is also likely ruled out as it generally leads to the formation of new molecular structures (such as heterocycles [8, 12]) that would alter the secondary fragment ions observed in the lower m/z region. Limited crosslinking, such as that observed by Poinar et al. 1998 [97], could be present but is difficult to confirm with the current data. Additionally, as static-SIMS only samples the uppermost ∼1-2 nm of specimen surface, a higher degree of biomolecule preservation may be present within the internal portions of the thin vascular “membrane” observed for these specimens. Regardless, the original fatty acids/phospholipids have, to an extent, been altered for the Little Blanche Creek bones, suggesting DNA preservation and protein diversity are also likely reduced relative to YG 126.115.

### Broader Implications for the Study of Ancient Bone

The methods used in this study can be readily applied to fossil and subfossil bones with warmer thermal histories. This includes Pleistocene bones of temperate and subtropical latitudes as well as specimens from pre-Pleistocene strata. The biochemical preservation of such Cenozoic specimens is not well known, particularly in comparison to the wealth of data reported for permafrost remains [11, 12, 27]. Furthermore, the biomolecular histology of such Cenozoic specimens has only been documented in a few isolated studies [27, 60–62, 79], including for nanoscale structures such as collagen fibrils and cellular membranes. Due to these gaps in the primary literature, the fossilization process, as relates to biomolecular tissue, remains poorly understood.

The nebulously defined term “subfossil” epitomizes the above stated gaps in knowledge. A “subfossil” is generally defined as a specimen that is not fully fossilized (see Comment 1 of the Supplemental Discussion). Thus far, criteria for determining whether a bone has been truly “fossilized”, and thus no longer considered a “subfossil”, have not been well established. “Subfossils” are currently classified using varying criteria, including degree of exogenous mineralization, whether a ^14^C date is returned, and geologic age [28–33, 98]. Such criteria are either somewhat subjective, not well defined, and/or not particularly insightful regarding underlying biochemical diagenesis. Rigorous criteria would facilitate consistent application of the term across studies and provide insight regarding biomolecular tissue diagenesis. Broader application of the methods used in this study can help establish such rigor. For example, this pilot study is the first to distinguish a set of pre-Holocene bones (not including remains of exceptional permafrost “mummies”) as “subfossils” rather than “fossils” using a rigorous criterium founded in the specimens’ underlying biomolecular histology. The permafrost bones examined by this study would certainly be classifiable as “subfossils” as their collagenous scaffolding is largely intact, and original vascular tissue is present, albeit having undergone relatively substantial alteration. These permafrost specimens are still biological bone and should not be considered “fossils” [40, 51–54], at least from a taphonomic standpoint.

Particularly, the preserved state of the type-1 collagen scaffolding (or lack thereof) is a potential criterium for establishing when a given bone specimen has been truly “fossilized”, thus ceasing to be a “subfossil”. Under this criterium, “subfossil” bone would correspond to bone preserving a relatively intact collagenous scaffolding. A “fossil” bone would correspond to a specimen in which either the collagen scaffolding has been mostly degraded and replaced with recrystallized mineral [14–17], or in which the constituent biomolecules have undergone extensive chemical transformation (i.e. *in-situ* polymerization and/or carbonization) [12, 18]. The preserved state of collagen scaffolding is an ideal criterium because type-1 collagen constitutes >90% of the OBM and is a literal “scaffold” onto which the bone’s mineral is deposited [35, 36]. Extensive application of the methods used within this pilot study to non-permafrost ancient bone is needed, however, to support the broader use of such a definition.

Few studies have recovered sequence-able proteins from pre-Pleistocene bones [99, 100]. Indeed, the only pre-Pleistocene bone from which peptide sequences have (thus far) been reported with minimal controversy is an isolated camel tibia from the Canadian Yukon territory [101, 102]. This prompts the question of how such an exceptional Pliocene bone harboring sequence-able proteins might be differentially preserved relative to the many Pliocene bones lacking such sequences. A potential answer to this question may be found by evaluating the sequence-preserving specimen’s biomolecular histology, as was done for the permafrost specimens of this present study. How does the OBM of the sequence preserving Pliocene bone compare with OBM of Pliocene bones (if OBM is even present) lacking recoverable sequences? Is the characteristic ∼67 nm banding pattern observable for OBM of the sequence preserving bone, or are “collagen” fibers present at all? Are cellular tissues present with associated membrane lipids, or have most biomolecules undergone *in-situ* polymerization? Is any biomolecular histology present at all, or are endogenous peptides preserved as small, isolated fragments in an otherwise mineral replica of the original bone? Should the bone be classified as a “fossil” or a “subfossil”? The approach used by this pilot study can resolve these questions and yield insight into the taphonomy of ancient tissues and their constituent biomolecules.

Finally, vascular tissue of specimen YG 126.115 did not exhibit the degree of diagenetic alteration observed for the other permafrost specimens. The presented data support YG 126.115 to have a diagenetic history unique from that of the other permafrost specimens and would therefore be preferable for molecular sequencing. This is a preliminary example of how biomolecular histology may be useful as a proxy for screening specimens for ancient sequencing studies; such an approach has previously been discussed in-depth by Anderson 2022 [27]. To truly test this, the permafrost bones of this study should be subjected to sequencing. This would allow cross-comparison of recovered peptide sequences with the data reported herein but is outside the scope of this current study.

## Conclusion

The first extensive nanoscale (up to 150,000× magnification) 3-D imaging data of soft tissues for *any* ancient bone are presented in Figures 1 and 5 of this study. The figured data demonstrate the presence of vascular tissue and well-preserved type-1 collagen fibrils within the Pleistocene permafrost specimens. This supports the taphonomic classification of these specimens as “subfossils”. The presence of vascular tissue and well-preserved collagen scaffolding suggests the examined specimens are incompletely “fossilized”, which is the current, but vague, definition of a “subfossil”. This is the first time that pre-Holocene bones (excluding the exceptional permafrost “mummies”) have been formally classifiable as “subfossils” rather than “fossils” based on underlying biomolecular histology. Broader application of the methods used herein to bone specimens of warmer thermal settings and earlier geologic strata can expand this insight and elucidate the changes that biomolecular tissues undergo during fossilization. This would help reveal the fate of biomolecules during fossilization and inform on what it means for a bone to be truly “fossilized”.

## Methods

### Specimens

Pleistocene bone specimens preserved in permafrost of eastern Beringia were received on loan from the Yukon government via the Yukon Paleontology program. The specimens consisted of 6 isolated, disarticulate bones, as follows: YG 610.2397 (*Mammuthus primigenius* innominate fragment; Little Blanche Creek, Yukon territory), YG 610.2364 (*Equus lambei* metatarsal; Little Blanche Creek, Yukon territory), YG 610.2305 (*Rangifer tarandus* antler; Little Blanche Creek, Yukon territory), YG 610.2365 (*Bison priscus* radius; Little Blanche Creek, Yukon territory), YG 610.2363 (*Bison priscus* metatarsal; Little Blanche Creek, Yukon territory), YG 126.115 (*Bison priscus* tibia; Irish Gulch, Yukon territory). Specimen YG 126.115 exhibited the preservation of desiccated soft tissue on the external surface of the bone, potentially remnant ligaments or tendons.

For the extant control bone specimens, fresh *Bos taurus* long bones (a humerus and a femur) were obtained from a local butcher (Schrock’s Slaughterhouse) immediately post-slaughter of the animals. The animals were not slaughtered for the purpose of this study, and the 2 bones were able to be acquired only because the butcher had no use for them. The bones were immediately placed on ice and stored at –20 degrees Celsius for a period of several months, until they were able to be transported to a lab at North Carolina State University, where they were stored at –80 degrees Celsius. The *Struthio camelus* long-bone shaft section is from the same specimen used in Schweitzer et al. 2007 [10]. The bone section has been stored under laboratory conditions for the past ∼16 years. *Alligator mississippiensis* long bones were obtained fresh from Cordray’s alligator farm and stored at –20 degrees Celsius; as with the *Bos taurus* long bones, these animals were not slaughtered for the purpose of this study. Rather, their bones were obtained as a by-product of the farm’s business practice.

Information for the protein standards purchased for the ToF-SIMS analyses are as follows: type-1 collagen protein (Sigma, CAS: 9007-34-5, C-9879, bovine achilles tendon); bovine serum albumin (BSA) (Fisher, CAS: 9048-46-8, BP 1600-100, heat-shock treated); hemoglobin (Sigma, CAS: 9008-02-0, H4131-1G, porcine).

Pleistocene permafrost bone samples were prepared in a dedicated “ancient” clean lab isolated from the extant controls and protein standards, while wearing nitrile gloves, a laboratory coat, a surgical mask, and a bouffant cap. Permafrost specimens were sampled using a hammer and chisel cleaned with 10% bleach followed by 70% ethanol. Preparatory area surfaces for permafrost samples were also sterilized with 10% bleach followed by 70% ethanol. Glassware and consumables were autoclaved prior to use. Solutions for ancient specimens were vacuum filtered (0.220 microns) prior to use in preparation protocols.

### Microscopy

Bone fragments (∼200-400 mg) were collected from each specimen and incubated in EDTA (0.5 M, pH 8.0) at room temperature for ∼1–5 days, until demineralized OBM was achieved. Care was taken to only sample the interior cortical bone that was the least discolored (in the case of the Pleistocene specimens). For light microscopy of demineralized OBM samples, a Zeiss Axioskop 2 plus microscope was used to image specimens mounted on glass slides, and resultant images were saved as “Tiff” files. For SEM imaging of OBM samples, samples were fixed for 1 h on ice in 2.5% glutaraldehyde with washes in phosphate-buffered saline (PBS) before and after fixation. A graded series of ethanol incubations (1 h at 50%, 1 h 70%, 1 h 95%, 3× 1 h 100% ethanol) was then used to dehydrate the samples. During the series of ethanol incubations, samples were placed within microporous specimen capsules (30 microns, SKU: 70187-20). For critical point drying, sputter coating, and imaging, samples were transported from the clean lab to the CHANL core facility at the University of North Carolina at Chapel Hill. Samples were critical point dried (Tousimis Autosamdri-931) and sputter coated (Cressington 108 Auto) with ∼80 angstroms of palladium-gold metal. Imaging was performed with a Hitachi S-4700 Cold Cathode Field Emission Scanning Electron Microscope with the accelerating voltage set to 15.0 kV. Collected images were processed in Adobe Photoshop 2021, using the Levels tool; a histogram stretch, followed by a gamma adjustment, and then a second histogram stretch were applied to each image. The protocol for light microscopy and SEM imaging of bone vasculature tissue was identical to that used for the OBM, except that a collagenase digestion was performed immediately post demineralization to isolate blood vessels from the OBM. This involved thorough washing of the demineralized OBM (ten times in water purified via a Barnstead E-Pure water purification system) to remove EDTA. The matrix sections were then incubated overnight at 37 degrees Celsius in a solution of ∼1 mg/mL collagenase (collagenase type I, Worthington Biochemical Corporation, 55B7870) dissolved in Dulbecco’s PBS (pH 7.2, with 0.1 g/L calcium chloride and 0.1 g/L magnesium chloride added to the standard PBS recipe) with 1% sodium azide (to inhibit potential microbial growth). Isolated vessels were washed in PBS buffer. At this point, either light microscopy or further processing for SEM (starting with glutaraldehyde fixation) was performed, as described for the OBM sections.

### ToF-SIMS and PCA

Small bone fragments (∼100 mg) were placed in 0.24 M hydrochloric acid to demineralize. After complete demineralization (∼1-2 days), OBM sections were removed and washed with E-Pure water. For vascular tissue analysis, the OBM sections were then incubated overnight at 37 degrees Celsius in 1 mg/mL collagenase (collagenase type I, Worthington Biochemical Corporation, 55B7870) with 1% sodium azide dissolved in Dulbecco’s PBS. Isolated blood vessels were washed twice with E-Pure water. For the type-1 collagen, BSA, and hemoglobin protein standards, chloroform/methanol extraction was necessary to remove residual lipids, despite the “purified” standards having been purchased from companies. A few grams of each standard was incubated for ∼30 minutes with agitation in a 2:1 mixture of chloroform:methanol. The chloroform:methanol solution was then decanted off, and the protein standards washed once with fresh chloroform:methanol solution. The wash solution was decanted off, and the protein standards were left to air dry in a chemical fume hood. Immediately (a few hours before) prior to specimen mounting, silicon wafers (provided courtesy of the Analytical Instrumentation Facility at NC State University) were subjected to an RCA-1 cleaning protocol. The RCA-1 cleaning solution consisted of a heated 5:1:1 mixture of 5 parts E-Pure H_2_O, 1 part 25% NH_4_OH, and 1 part 30% H_2_O_2_. Cleaned wafers were washed twice with E-Pure water and left to dry in a laminar flow hood.

For specimen mounting, OBM and blood vessel samples were suspended in sterile E-Pure water and pipetted directly onto the RCA-1 cleaned silicon wafers and allowed to air dry in a laminar flow hood. Protein standards were mounted directly onto copper conducting tape, provided courtesy of the CHANL core facility at UNC Chapel Hill. The specimens were then immediately transported to the Analytical Instrumentation Facility at North Carolina State University and placed under vacuum within the ToF-SIMS instrument. Sample analysis took place the following day.

A TOF-SIMS V (ION TOF, Inc.) was used to analyze the prepared OBM, blood vessel, and protein standard samples. Analyses were performed with a Bi_3_^+^ liquid metal ion gun at 45° incident to the sample surface under an ultra-high vacuum of ∼7.9×10^-10^ mbar. Regions of 50×50, 100×100, or 150×150 µm^2^ were scanned using the instrument’s high current-bunched (high mass resolution) setting with a beam diameter ∼10 microns, a pulse width of ∼1 ns, and a cycle time of 100 microseconds. Data was recorded in the instrument’s positive mode for OBM sections and protein standards, and in the negative mode for vascular tissue. Images obtained were 128×128 pixels with 1 shot/pixel at an approximate target current of 0.33 pA (this was checked using a Faraday cup prior to each analysis session to ensure the static “limit” was not exceeded). Primary ion dosage was kept below 10^12^ ions/cm^2^ to ensure all analyses were performed under static conditions. Resultant spectra had an approximate mass resolution of 3,000-6,000 m/Δm, dependent on degree of charging and analysis area topography. An electron flood gun was used for charge compensation to mitigate charging-related issues such as degraded mass resolution and attenuated secondary ion signal. When degraded mass resolution (or even peak splitting) due to sample topography were encountered, relatively flat regions of interest within the analysis area were selected to mitigate this issue.

A Poisson correction was applied to the spectra to correct for missed ion counts during instrument dead time. Positive mode spectra were calibrated using the secondary ions C_2_H_3_^+^, C_3_H_5_^+^, C_3_H_7_^+^, C_4_H_7_^+^, and C_5_H_7_^+^. Negative mode spectra were calibrated with the secondary ions C_3_H^-^, C_5_H^-^, C_6_^-^, and C_6_H^-^. Manually selected peak lists of 27 amino acid-related secondary ion fragments (Table S1) and 7 fatty acid-related ions (Table S2) were applied to the positive and negative mode spectra, respectively. The amino acid-related peak list was selected based on Mantus et al. 1993, Wagner et al. 2002, Wagner et al. 2002, and Henss et al. 2013 [71–73, 77]. Areas for secondary ion peaks within the peak lists were calculated and placed in respective data matrices. Resultant data matrices (one each for the positive and negative mode spectra, respectively) were processed using Primer 7 software (version 7.0.23; Primer-E, Quest Research Lmtd., Auckland, New Zealand). Extracted peak areas were standardized to their respective spectrum’s summed intensity for all secondary ions included in the respective peak list. PCA was then performed on each of the 2 standardized data matrices, with resultant PC scores and loadings plotted on separate graphs. Loading values were calculated from the eigenvectors (these contain the loading values scaled to unity) by multiplying each eigenvector value with the square root of the respective PC’s eigenvalue.

### Isotopic and C:N Measurement

Large bone fragments (several grams) were collected from each of the Pleistocene permafrost and extant bone specimens, with care taken to only sample interior cortical bone with minimal discoloration (in the case of the Pleistocene samples). completely demineralized in ∼0.24 M HCl (the individual pieces in the tubes were all easily cut with sterile razor blades prior to freeze-drying) and incubated for 24 hrs in 0.1 M NaOH (with thorough wash steps in E-Pure water in between). The samples were then frozen at –80 degrees Celsius and lyophilized overnight.

Lyophilized OBM was then shipped to the University of Delaware Environmental Isotope Science Laboratory, where the remaining analysis protocol was performed by the laboratory manager on a fee-for-service basis. Samples and reference materials (provided by the UD Environmental Isotope Science Laboratory) were ground and loaded into standard weight pressed tin capsules (8×5 mm; Elemental Microanalysis). The loaded capsules were then elementally and isotopically analyzed using a Thermo Scientific EA IsoLink IRMS System. Sample combustion/elemental analysis was performed with a Flash 2000 EA equipped with a MAS200R autosampler. The Flash EA (elemental analyzer) was configured with a single reactor combustion/reduction tube at 1000 degrees C, helium flow of 180 mL/min., and a GC (gas chromatograph) oven at 40 degrees C. The Flash EA was connected via a ConFlo IV universal continuous flow interface to a Delta V IRMS (isotope ratio mass spectrometer), which allowed collection of stable isotope δ^13^C and δ^15^N values from evolved CO_2_ and N_2_ (from sample combustion in the EA).

The L-glutamic acid reference samples USGS40 and USGS41a were used for instrument calibration, and the glycine reference samples USGS64, USGS65, and USGS66 were analyzed as comparative controls. Two of the extant and three of the Pleistocene samples were analyzed in-duplicate to ensure reproducibility; resultant standard deviations were ∼0.2 ‰. All values measured for δ^13^C are reported relative to Vienna PeeDee belemnite (VPDB) on a scale “normalized such that the of δ^13^C values of NBS 19 calcium carbonate and L-SVEC lithium carbonate are +1.95 ‰ and –46.6 ‰, respectively” (see Coplen et al. 2006 [103]). All measured δ^15^N values are reported relative to atmospheric nitrogen (δ^15^N AIR).

## Competing Interests

The author has no relevant financial or non-financial interests to disclose.

## Data Availability Statement

All underlying/supporting data is either provided in the supplemental material or available from the corresponding author upon reasonable request.

## Supporting information

Supplemental datasheet 1

Supplemental datasheet 2

Unprocessed microscope images

## Notes

### Competing Interest Statement

The authors have declared no competing interest.

