## Supplementary material for "Nanoscale Imaging and Microanalysis of Ice Age Bone Offers New Perspective on “Subfossils” and Fossilization": Supplemental Information.pdf

### Supplemental Discussion

***Comment 1*** *on the definition of "subfossil":*

"Subfossil" has historically been defined one of two ways. One is the definition given in the main text, referring to a specimen that is not yet fully "fossilized" [1]. Currently, this definition is vague as to what is meant by "not yet fully fossilized", but it does at least attempt to consider a specimen's underlying taphonomy. A primary aim of the main text is to more rigorously define at which point bone has been fully "fossilized". An alternative operational definition has also been used in past studies that classifies all organismal remains (in some stage of diagenesis) with assigned dates <~10 Ka as "subfossils" [2]. Such a definition provides nominal insight regarding the taphonomy/preserved state of a given ancient specimen and can be problematic for molecular studies. For example, extant bones have been observed to undergo substantial mineralization towards a "fossil"-like state over a timescale of decades [3]. Likewise, Pleistocene megafaunal carcasses frozen in permafrost have been recovered such that researchers were able to consume remnant musculature in a "stew" [4]. An arbitrary cutoff of ~10 Ka for distinguishing fossils from subfossils introduces potential for ambiguity and confusion regarding these two cases. As a further example, in Orlando et al. 2013, the permafrost frozen, mid-Pleistocene *Equus spp.* metapodial is consistently referred to as a "fossil". However, the study reports extensive DNA and protein sequence preservation, suggesting the specimen itself is not fully "fossilized" [5]. This introduces ambiguity regarding preservational state because the study itself does not define what is meant by the term "fossil" as used in the paper. Indeed, most molecular paleontological and archaeological studies have adhered to this same standard, leading to ambiguity regarding the preserved state of reported specimens.

***Comment 2*** *on the limited mass resolution of the ToF mass analyzer in context of this study:*

A limitation of ToF analyzers is that mass resolution of spectra they produce (m/Δm = ~7,000-10,000, assuming minimal charging and flat sample topography) is insufficient for identifying many individual spectral ions without *a priori* knowledge of sample composition [6]. For the collagenous specimens analyzed by this study, this was mitigated by comparing sample fragmentation patterns (chemical "fingerprints") against those of known standards and extant controls. Furthermore, as organic bone matrix (OBM) is upwards of 90% type-1 collagen protein by composition, it functions somewhat similarly to a homogenous substance which simplifies spectral interpretation [7]. This is not the case for vascular tissue as basal endothelium is cellular and thus inherently heterogenous/complex.

Even still, for the extant controls, because vascular tissue endothelium is a cellular structure, phospholipids would be expected to be present in high abundances as they are primary components of cellular membranes. In SIMS analyses, phospholipids readily ionize as fatty acid molecular ions; past studies have shown that, for extant cells (including those of vascular tissue), signal for a variety of fatty acid molecular ions is readily obtained [8, 9]. Indeed, molecular ion peaks matching "fatty acid" m/z values are observed in the extant vasculature spectra of the present study; these molecular ion peaks are thus attributable to fatty acid molecular ions. Furthermore, the YG 126.115 spectra (as regards the fatty acid molecular ion peaks) closely match the extant controls, which lends confidence to the presence of preserved fatty acids within this ancient specimen. This is not the case for the Little Blanche Creek specimens, despite the presence of sporadic ions (of limited signal intensity) corresponding to fatty acids. Their spectra deviate substantially from the extant controls as regards "fatty acid" molecular ion peak distributions/intensities. Rather, what can be said is that surface signal corresponding to fatty acid m/z values is substantially reduced in the Little Blanche Creek bones relative to extant blood vessel endothelium, thus supporting substantial diagenetic alteration of the original membrane lipid bilayers.

#### Supplemental Tables

***Table S1. Secondary ion peaks selected for PCA of ToF-SIMS organic bone matrix (OBM) and protein standard spectra.*** *Included secondary ion masses were based on past studies reporting ToF-SIMS spectra of amino acid homopolymers [10-12] and bone [13]. Values for secondary ion observed masses are only listed to 2 decimal places because of the ToF analyzer's limited mass resolution. For this table, amino acids are generally listed as corresponding to a given secondary ion fragment if they produce a substantial yield of that given ion. For example,* $C_4H_8N^+$ *can be produced by valine, but it is only observed in minor amounts for this amino acid and thus is not listed [10]. Additionally, a variety of organic compounds aside from proteins/amino acids can produce some of these ions, and individually, no one ion can be definitively attributed to a given amino acid using ToF-SIMS alone. Rather, the combination of the presented microscopy data, C:N ratios, and secondary ion intensities and distributions relative to the type-1 collagen protein standard, allow for the confident identification of type-1 collagen within the permafrost specimens.*

| Secondary Ion | Observed Mass for PCA | Corresponding Amino Acid(s) |
|---|---|---|
| $\mathbf{CH_4N^+}$ | 30.04 | Glycine, lysine, most others |
| $\mathbf{C_2H_6N^+}$ | 44.05 | Alanine, various |

| | | |
|---|---|---|
| | | others |
| **$C_2H_6NO^+$** | 60.06 | Serine |
| **$C_2H_5S^+$** | 61.01 | Methionine |
| **$C_4H_6N^+$** | 68.05 | Proline |
| **$C_3H_4NO^+$** | 70.03 | Asparagine |
| **$C_4H_8N^+$** | 70.07 | Proline, Arginine |
| **$C_3H_6NO^+$** | 72.05 | Aspartic acid |
| **$C_4H_{10}N^+$** | 72.09 | Valine |
| **$C_2H_7N_3^+$** | 73.06 | Arginine |
| **$C_3H_8NO^+$** | 74.07 | Threonine |
| **$C_2H_6NS^+$** | 76.03 | Cysteine |
| **$C_4H_5N_2^+$** | 81.04 | Histidine |
| **$C_4H_6N_2^+$** | 82.06 | Histidine |
| **$C_4H_6NO^+$** | 84.05 | Glutamine, glutamic acid |
| **$C_5H_{10}N^+$** | 84.08 | Lysine, leucine |
| **$C_4H_8NO^+$** | 86.06 | Hydroxyproline (small amounts can be formed by aspartic acid [10]) |
| **$C_5H_{12}N^+$** | 86.10 | Leucine, isoleucine |
| **$C_3H_6NO_2^+$** | 88.04 | Asparagine, aspartic acid |
| **$C_4H_4NO_2^+$** | 98.02 | Asparagine |
| **$C_4H_8NO_2^+$** | 102.06 | Glutamic acid |
| **$C_4H_{10}NS^+$** | 104.05 | Methionine |

| | | |
|---|---|---|
| $C_5H_8N_3^+$ | 110.08 | Histidine |
| $C_8H_{10}N^+$ | 120.08 | Phenylalanine |
| $C_5H_{11}N_4^+$ | 127.10 | Arginine |
| $C_9H_8N^+$ | 130.07 | Tryptophan |
| $C_8H_{10}NO^+$ | 136.07 | Tyrosine |

***Table S2. Secondary ion peaks selected for PCA of ToF-SIMS vascular tissue spectra.*** *Secondary ion peaks corresponding to the (M-H$^+$) ions for myristic, palmitoleic, palmitic, margaric, linoleic, oleic, and stearic acids were initially selected for PCA of the vascular tissue spectra. Some of these secondary ions can also correspond to fragment ions of other fatty acids [14], as shown in the table. Values for secondary ion observed masses are only listed to 2 decimal places because of the ToF analyzer's limited mass resolution. See Comment 2 under the Supplemental Discussion section for a description of how these ions were assigned within the extant and permafrost vascular tissue spectra. Briefly, only the permafrost specimen YG 126.115 shared similar secondary ion distribution and intensities (for the secondary ions listed in Table S2) with the extant control spectra. This supported attribution of these secondary ion peaks to fatty acids. For the other permafrost specimens, some secondary ions potentially corresponding to fatty acids were observed, but the disparity in distribution and intensity (particularly their generally low intensity, often barely above background) with the extant control spectra prevented confident attribution of these peaks to fatty acids.*

| Secondary Ion | Observed Mass for PCA | Corresponding Fatty Acid(s) |
|---|---|---|
| $C_{14}H_{27}O_2^-$ | 227.20 | Myristic, palmitic, stearic, sphingosine |

| | | |
|---|---|---|
| $C_{14}H_{29}O_2^-$ | 253.22 | Palmitoleic, palmitic, stearic |
| $C_{16}H_{31}O_2^-$ | 255.24 | Palmitic, stearic, sphingosine |
| $C_{14}H_{27}O_2^-$ | 269.27 | Margaric, stearic |
| $C_{18}H_{31}O_2^-$ | 279.25 | Stearic, oleic, linoleic, sphingosine |
| $C_{18}H_{33}O_2^-$ | 281.25 | Oleic, stearic |
| $C_{18}H_{35}O_2^-$ | 283.27 | Stearic |

#### Supplemental Figures

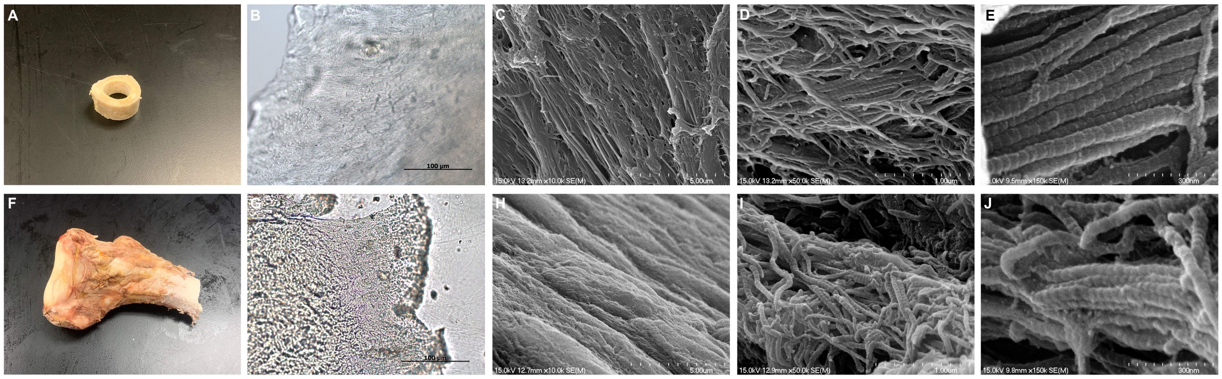


***Figure S1. Light and electron microscope images of extant S. camelus (A-E) and A. mississippiensis (F-J) OBM.***

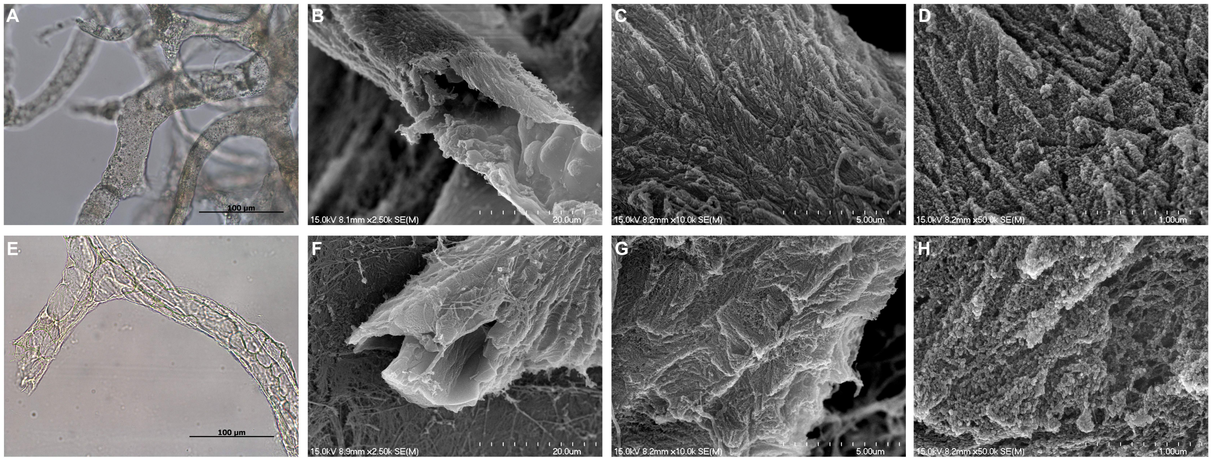

***Figure S2. Light and electron microscope images of extant S. camelus (A-D) and A. mississippiensis (E-H) vascular tissue.*** *The endothelial membranes tend to be thicker and more structured in these extant vessels relative to the Little Blanche Creek specimens.*

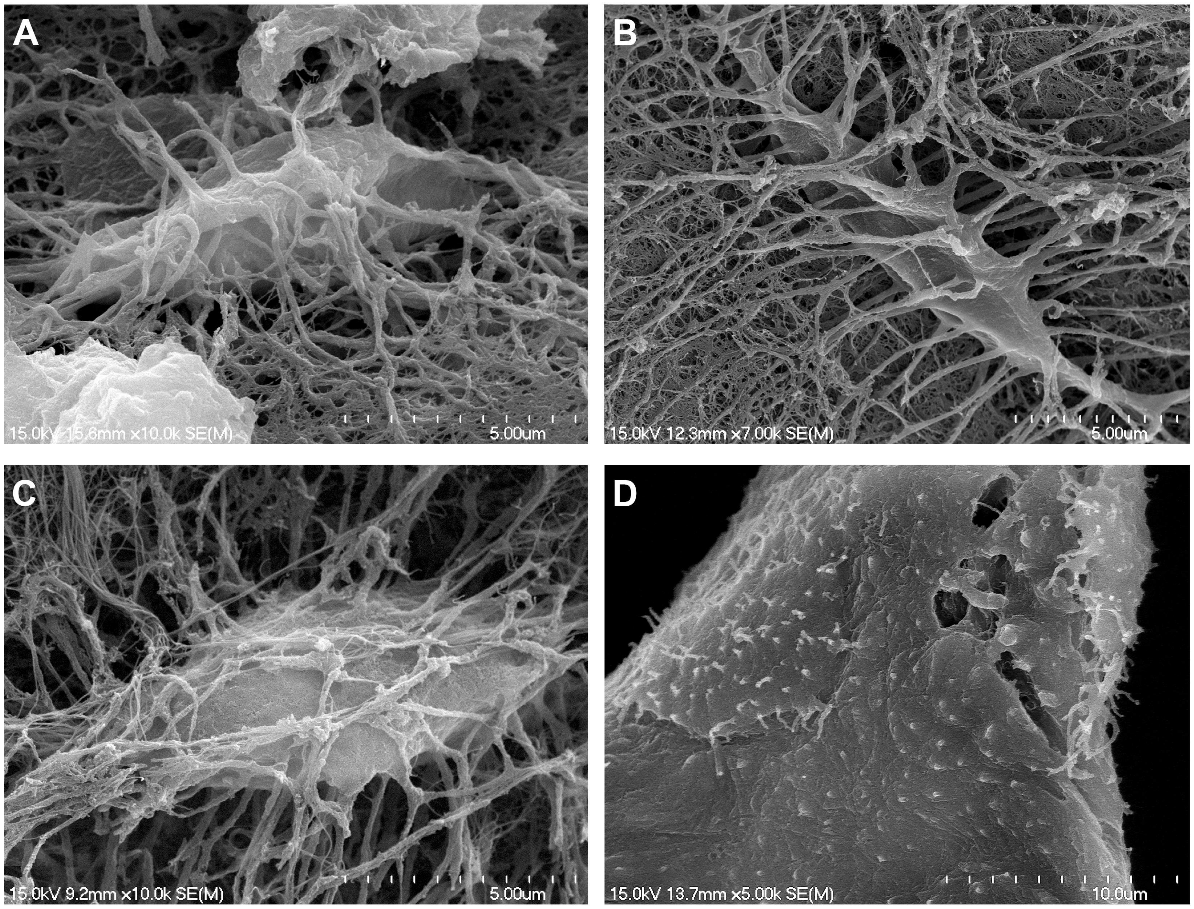


***Figure S3. Electron microscope images of Pleistocene permafrost and extant osteocytes.*** *Osteocytes for the extant A. mississippiensis specimen are not shown but were observed in high abundance. No osteocytes were observed for the Little Blanche Creek specimens except YG 610.2364, as shown in (B).* ***(A)*** *YG 126.115 (B. priscus tibia).* ***(B)*** *YG 610.2364 (E. lambei metatarsal).* ***(C)*** *Extant B. taurus.* ***(D)*** *Extant S. camelus.*

** Note that absolute secondary ion peak intensities cannot be directly compared between the various ToF-SIMS spectra sets shown below as they have not been standardized to account for differences in overall secondary ion yield between analysis areas. However, the relative intensities of the secondary ion peaks can be compared across the spectra. For the PCA shown in the main text, the data was standardized prior to running the statistical analysis.

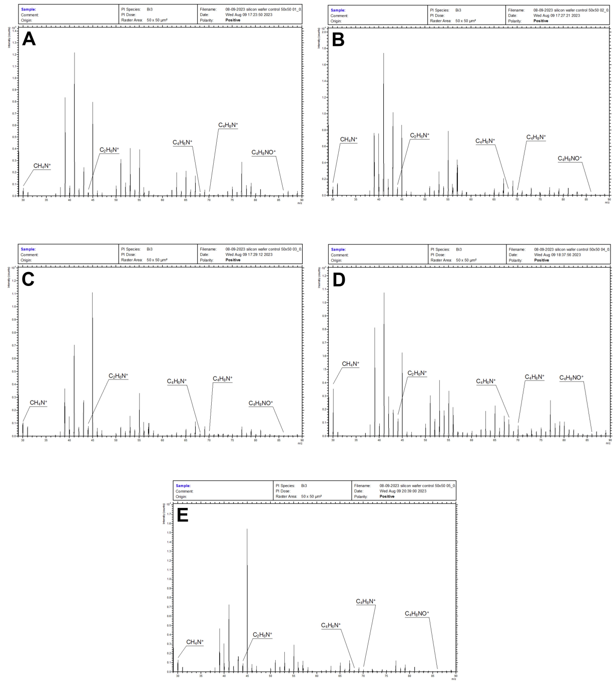


***Figure S4. ToF-SIMS spectra collected of the blank silicon wafer surface (A-E)*** *The spectra shown above were taken of the blank silicon wafer surface. OBM samples were mounted onto silicon wafers for ToF-SIMS analyses. The range of m/z values is shown from 29-90 so as to exclude the* $Si^{+}$ *secondary ion peak. Otherwise, its high intensity relative to the other observed secondary ions would affect the y-axis scaling of the shown spectra.*

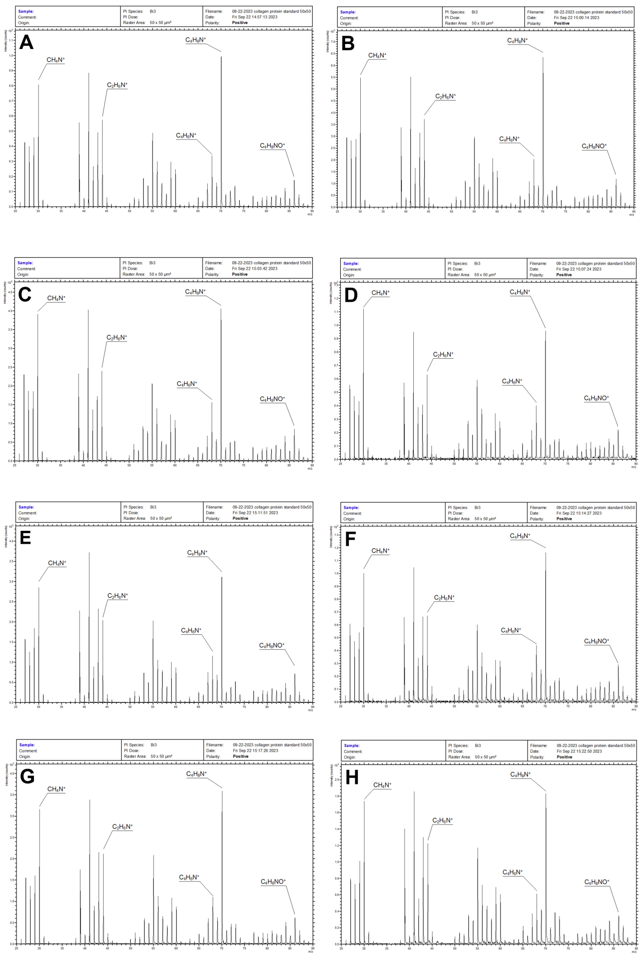

***Figure S5. ToF-SIMS spectra collected of the type-1 collagen protein standard (A-H)*** *The positive ion spectra shown above were taken of purified bovine type-1 collagen protein. Note the high intensities for the peaks $C_4H_6N^+$ and $C_4H_8N^+$ relative to the hemoglobin and BSA spectra shown in Figures S6 and S7, respectively. Also note the lower intensities for the peaks $C_4H_{10}N^+$ (valine) and $C_5H_{10}N^+$ (lysine/leucine) relative to the hemoglobin and BSA standards. These differences agree well with the known amino acid compositions for these different proteins [15-17]. Furthermore, the secondary ion $C_4H_8NO^+$ (hydroxyproline) is observed in the above type-1 collagen spectra (peaks for both $C_4H_8NO^+$ and $C_5H_{12}N^+$ are present, but*

*indistinguishable due to the large m/z range shown), but not within the hemoglobin or BSA spectra. Glycine constitutes about one-third of type-1 collagen amino acids, but its corresponding secondary ion $CH_4N^+$ is ubiquitous to most amino acids, hence the limited difference between the different protein standards [10, 15-17].*

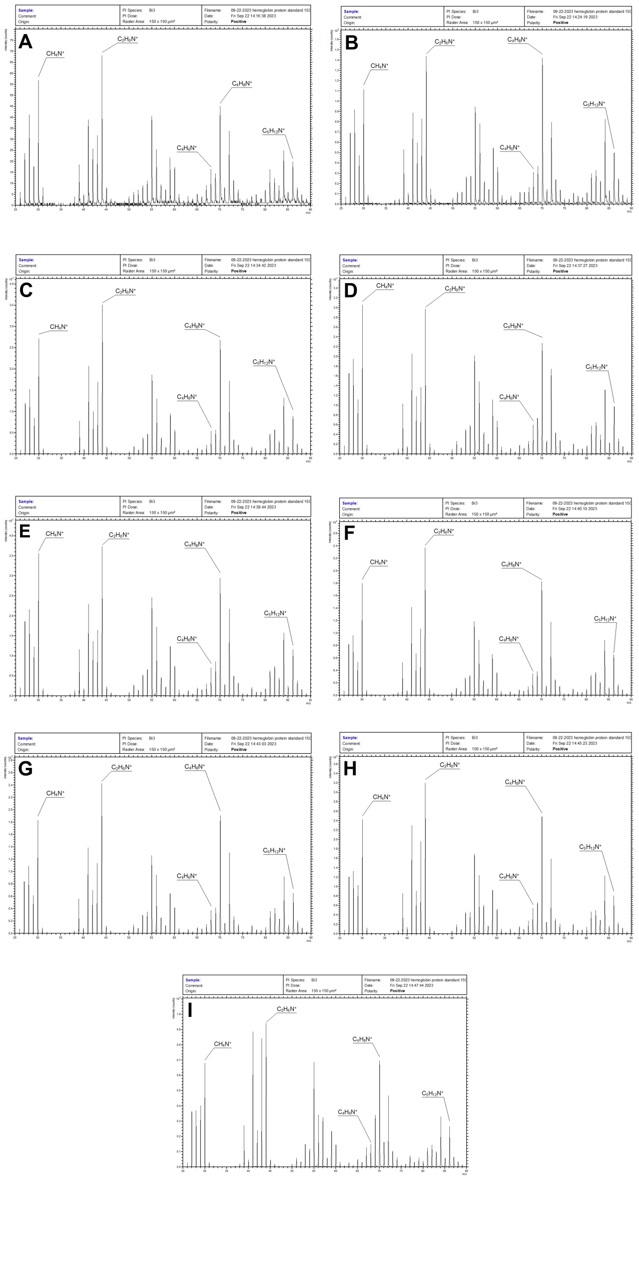

***Figure S6. ToF-SIMS spectra collected of the hemoglobin protein standard (A-I)*** *The positive ion spectra shown above were taken of purified porcine hemoglobin. Note the higher intensity of the secondary ion corresponding to alanine ($C_2H_6N^+$, 44.05) relative to that for proline ($C_4H_8N^+$, 70.07). The type-1 collagen spectra of Figure S5 tend to show an opposite trend, with higher intensities observed for $C_4H_8N^+$ relative to $C_2H_6N^+$. Additionally, the peaks corresponding to valine ($C_4H_{10}N^+$, 72.09) and lysine/leucine ($C_5H_{10}N^+$, 84.08) [10-12] record higher intensities relative to the type-1 collagen standard spectra. These amino acids are present in substantially higher abundances within hemoglobin relative to type-1 collagen [16, 17]. Also note that while the $C_5H_{12}N^+$ peak is present at a nominal m/z of 86, the $C_4H_8NO^+$ secondary ion peak is absent, likely due to the lack of hydroxyproline in hemoglobin.*

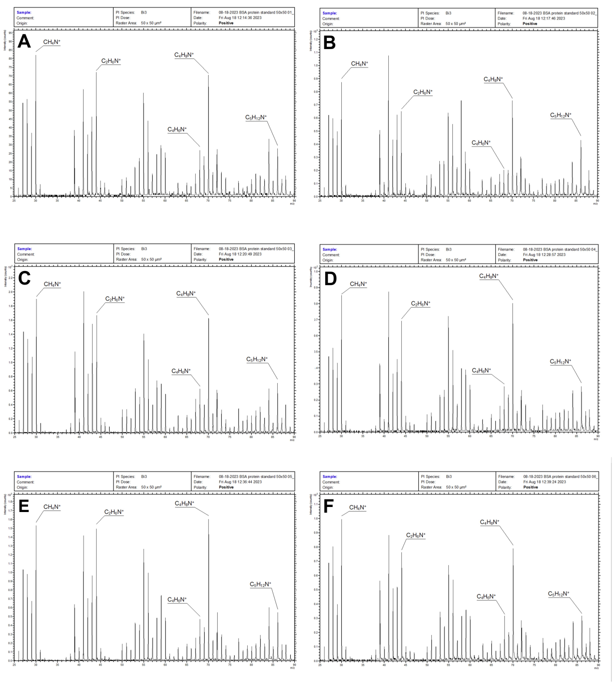


***Figure S7. ToF-SIMS spectra collected of the BSA protein standard (A-F)*** *The positive ion spectra shown above were taken of purified bovine serum albumin (BSA). The intensity for the secondary ion corresponding to alanine ($C_2H_6N^+$, 44.05) is similar to that for proline ($C_4H_8N^+$, 70.07). This agrees with BSA having a lower alanine content relative to hemoglobin, and a lower proline content than type-1 collagen. Furthermore, secondary ions corresponding to valine ($C_4H_{10}N^+$, 72.09) and lysine/leucine ($C_5H_{10}N^+$, 84.08) [10-12] are of higher intensities relative to the type-1 collagen standard spectra. Like the hemoglobin standard in Figure S6, these amino acids are present in substantially higher abundances within BSA relative to type-1 collagen [15-17]. Also while the $C_5H_{12}N^+$ peak is present at a nominal m/z of 86, the $C_4H_8NO^+$ secondary ion peak is absent, likely due to the lack of hydroxyproline in BSA.*

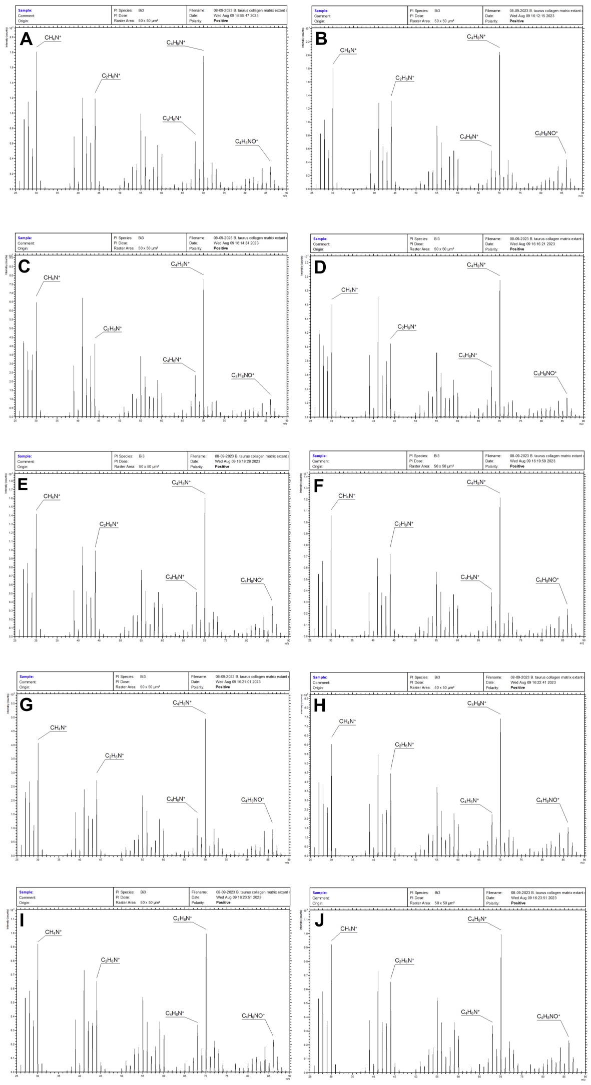

***Figure S8. ToF-SIMS spectra collected of extant B. taurus OBM (A-J)*** *The positive ion spectra shown above were taken of the demineralized extant B. taurus OBM. The high relative intensities for the peaks $C_4H_6N^+$ and $C_4H_8N^+$ agree with the type-1 collagen protein standard and which is not the case for the hemoglobin and BSA spectra shown in Figures S6 and S7, respectively. The peaks $C_4H_{10}N^+$ (valine) and $C_5H_{10}N^+$ (lysine/leucine) exhibit lower intensities relative to the hemoglobin and BSA standards. Furthermore, the secondary ion $C_4H_8NO^+$ (hydroxyproline) is observed in the*

*above type-1 collagen spectra (peaks for both $C_4H_8NO^+$ and $C_5H_{12}N^+$ are present, but indistinguishable due to the large m/z range shown), but not within the hemoglobin or BSA spectra. Glycine constitutes about one-third of type-1 collagen amino acids, but its corresponding secondary ion $CH_4N^+$ is ubiquitous to most amino acids, hence the limited difference between the above spectra and the non-collagen protein standards [10, 15-17]. Despite this sample being OBM and not purified type-1 collagen, the spectra closely match the purified type-1 collagen standard. This is likely due to the high type-1 collagen content (>90%) for OBM [18, 19]. This also indicates the collagenous scaffolding was analyzed rather than non-collagenous portions such as vascular tissue.*

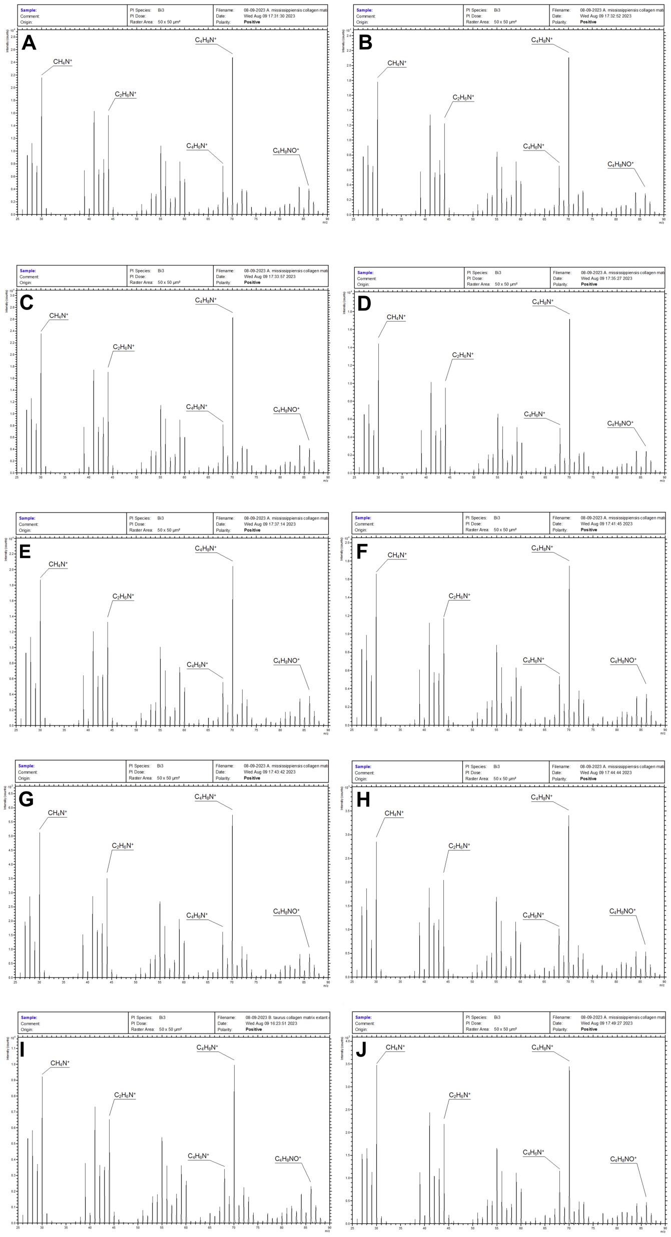


**Figure S9. ToF-SIMS spectra collected of extant *A. mississippiensis* OBM (A-J)** *The positive ion spectra shown above were taken of the demineralized extant A. mississippiensis OBM. Little difference relative to the B. taurus spectra (Figure S8) and the type-1 collagen protein standard spectra (Figure S5) is observable; hence, the description given by Figure S8 also fits for the spectra figured above.*

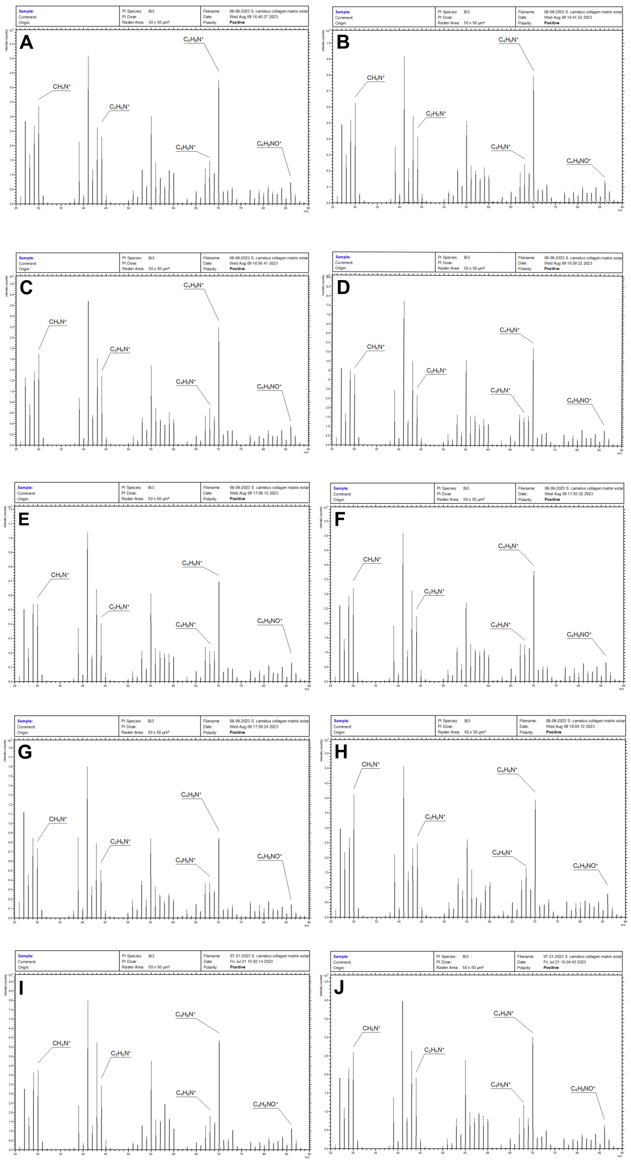


***Figure S10. ToF-SIMS spectra collected of extant S. camelus OBM (A-J)*** *The positive ion spectra shown above were taken of the demineralized extant S. camelus OBM. Little difference relative to the B. taurus spectra (Figure S8) and the type-1 collagen protein standard spectra (Figure S5) is observable; hence, the description given by Figure S8 also fits for the spectra figured above.*

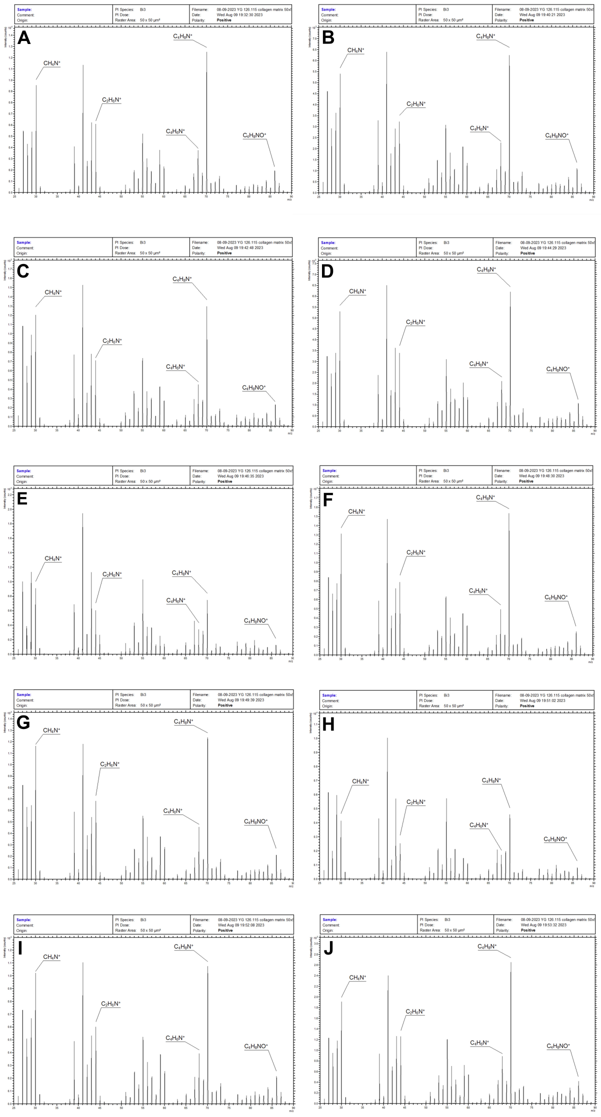

***Figure S11. ToF-SIMS spectra collected of OBM from specimen YG 126.115 (A-J)*** *The positive ion spectra shown above were taken of the demineralized YG 126.115 OBM. High secondary ion yields were observed for many of the protein-related fragment ions included in the PCA analysis, despite this being an ancient specimen. Little difference relative to the B. taurus spectra (Figure S8) and the type-1 collagen protein standard spectra (Figure S5) is observable; hence, the see Figure S8 legend.*

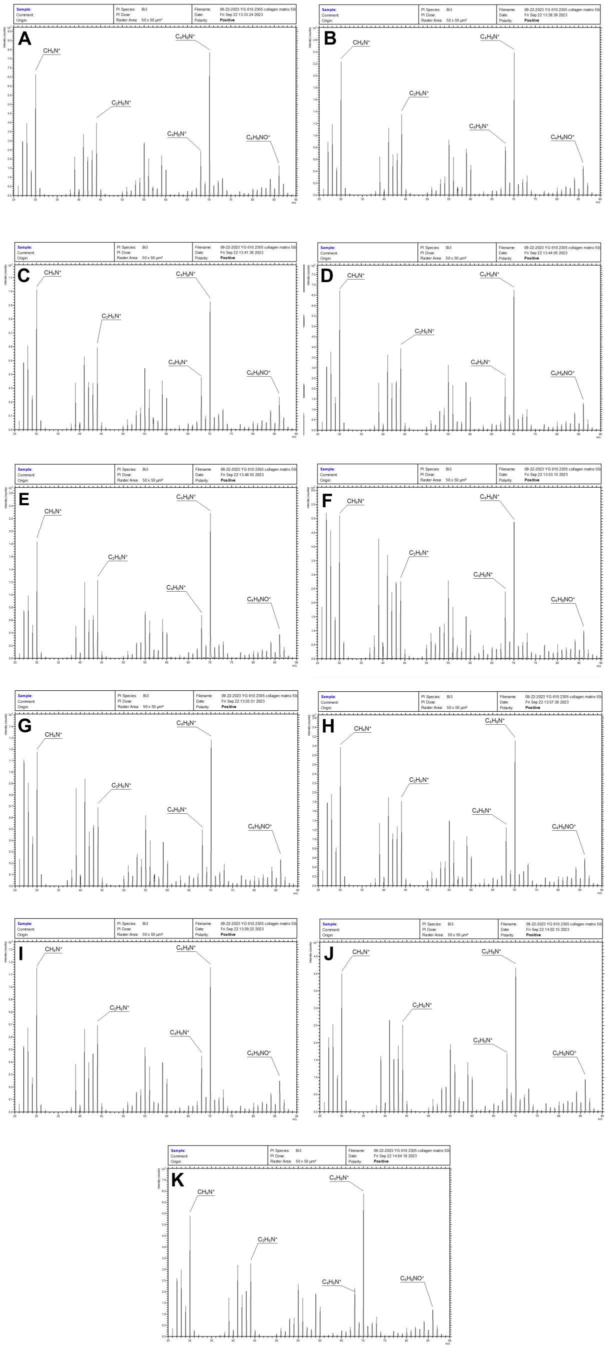


***Figure S12. ToF-SIMS spectra collected of OBM from specimen YG 610.2305 (A-K)*** *The positive ion spectra shown above were taken of the demineralized YG 610.2305*

*OBM. High secondary ion yields were observed for many of the protein-related fragment ions included in the PCA analysis, despite this being an ancient specimen. Little difference relative to the B. taurus spectra (Figure S8) and the type-1 collagen protein standard spectra (Figure S5) is observable; hence, the see Figure S8 legend.*

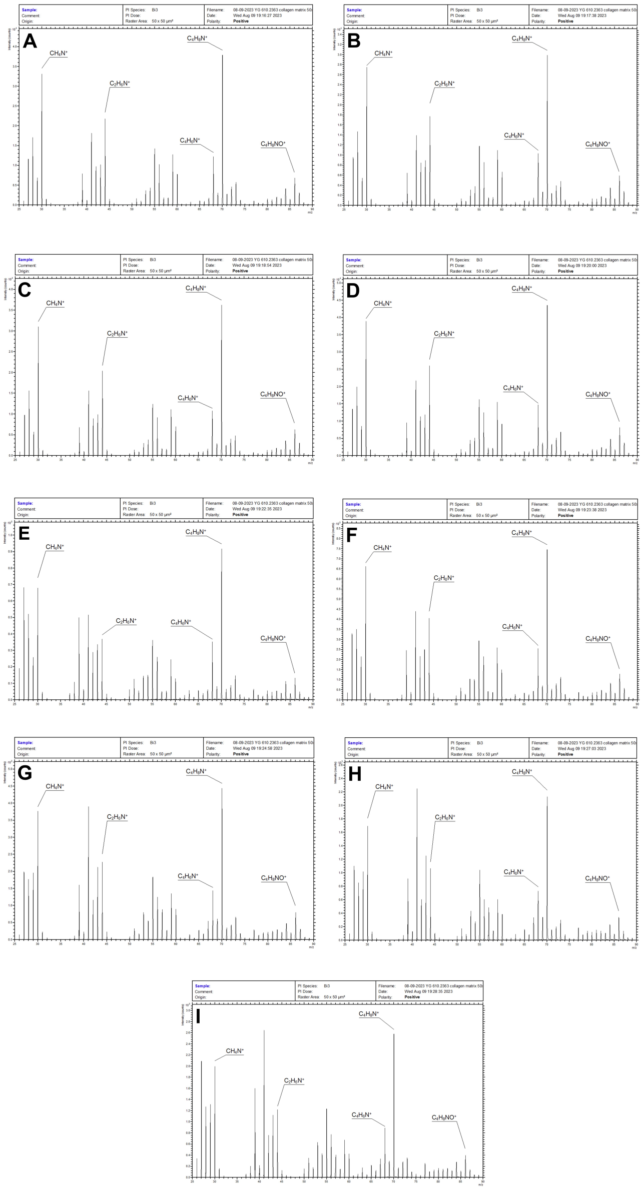

***Figure S13. ToF-SIMS spectra collected of OBM from specimen YG 610.2363 (A-I)*** *The positive ion spectra shown above were taken of the demineralized YG 610.2363 OBM. High secondary ion yields were observed for many of the protein-related fragment ions included in the PCA analysis, despite this being an ancient specimen. Little difference relative to the B. taurus spectra (Figure S8) and the type-1 collagen protein standard spectra (Figure S5) is observable; hence, the see Figure S8 legend.*

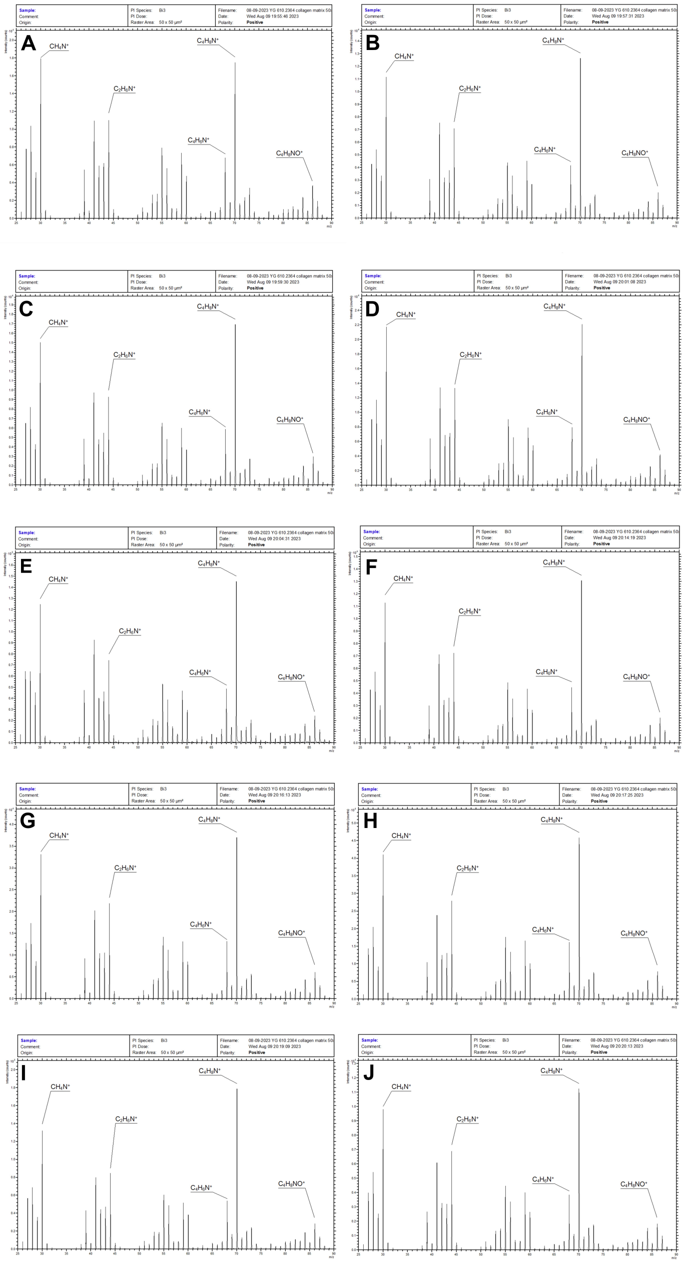


***Figure S14. ToF-SIMS spectra collected of OBM from specimen YG 610.2364 (A-J)*** *The positive ion spectra shown above were taken of the demineralized YG 610.2364 OBM. High secondary ion yields were observed for many of the protein-related fragment ions included in the PCA analysis, despite this being an ancient specimen. Little difference relative to the B. taurus spectra (Figure S8) and*

*the type-1 collagen protein standard spectra (Figure S5) is observable; hence, the see Figure S8 legend.*

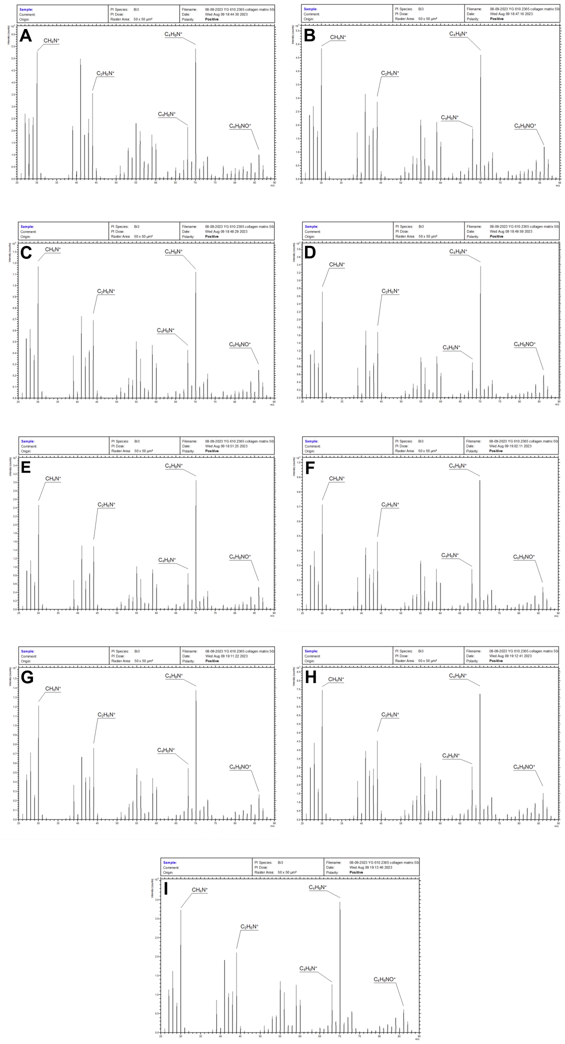


***Figure S15. ToF-SIMS spectra collected of OBM from specimen YG 610.2365 (A-I)*** *The positive ion spectra shown above were taken of the demineralized YG 610.2365 OBM. High secondary ion yields were observed for many of the protein-related fragment ions included in the PCA analysis, despite this being an ancient specimen. Little difference relative to the B. taurus spectra (Figure S8) and the type-1 collagen protein standard spectra (Figure S5) is observable; hence, the see Figure S8 legend.*

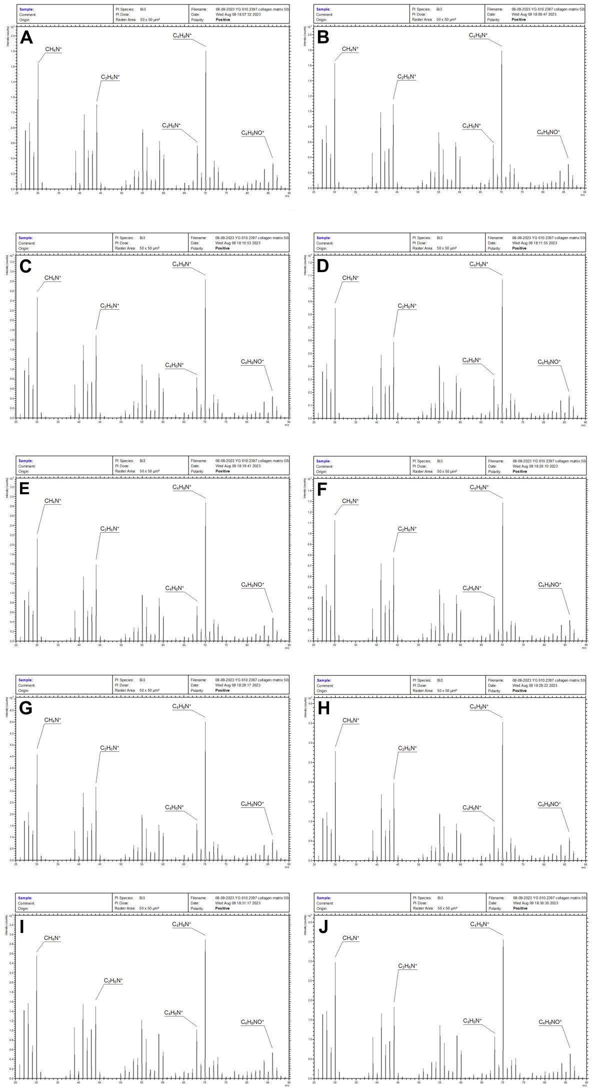


***Figure S16. ToF-SIMS spectra collected of OBM from specimen YG 610.2397 (A-J)*** *The positive ion spectra shown above were taken of the demineralized YG 610.2397 OBM. High secondary ion yields were observed for many of the protein-related fragment ions included in the PCA analysis, despite this being an ancient specimen. Little difference relative to the B. taurus spectra (Figure S8) and the type-1 collagen protein standard spectra (Figure S5) is observable; hence, the see Figure S8 legend.*

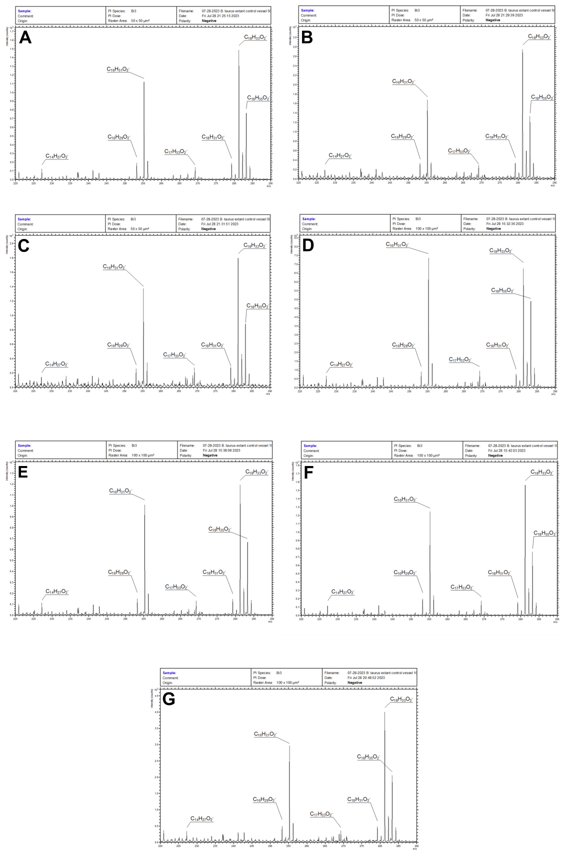


***Figure S17. ToF-SIMS spectra collected of extant B. taurus vascular tissue (A-G)*** *Negative ion spectra shown above were taken of the demineralized B. taurus vascular tissue post-collagenase digestion. Secondary ion peaks corresponding to m/z values for a variety of fatty acids are consistently observed. This is expected as the vascular tissue (basal endothelium) is cellular in nature, consisting substantially of phospholipids. Note the comparative intensities of the palmitic and oleic acid molecular ions vary between spectra. As discussed in the main text, this may be attributable to differences in membrane lipid composition compounded by the air-drying process exposing internal cytosolic membranes.*

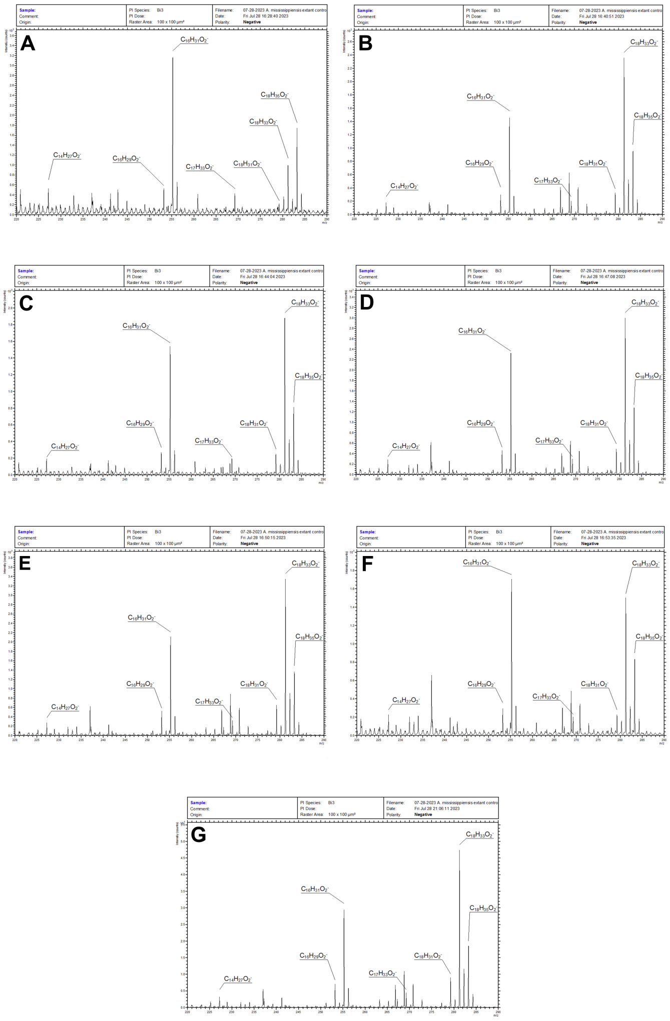


**Figure S18. ToF-SIMS spectra collected of extant A. mississippiensis vascular tissue (A-G)** *Negative ion spectra shown above were taken of the demineralized A. mississippiensis vascular tissue post-collagenase digestion. As with the other extant specimens, secondary ion peaks corresponding to m/z values for a variety of fatty acids are consistently observed. This is expected as the vascular tissue (basal endothelium) is cellular in nature, consisting substantially of phospholipids. Note the comparative intensities of the palmitic and oleic acid molecular ions vary substantially between spectra. As discussed in the main text, this may be attributable to differences in membrane lipid*

*composition compounded by the air-drying process exposing internal cytosolic membranes.*

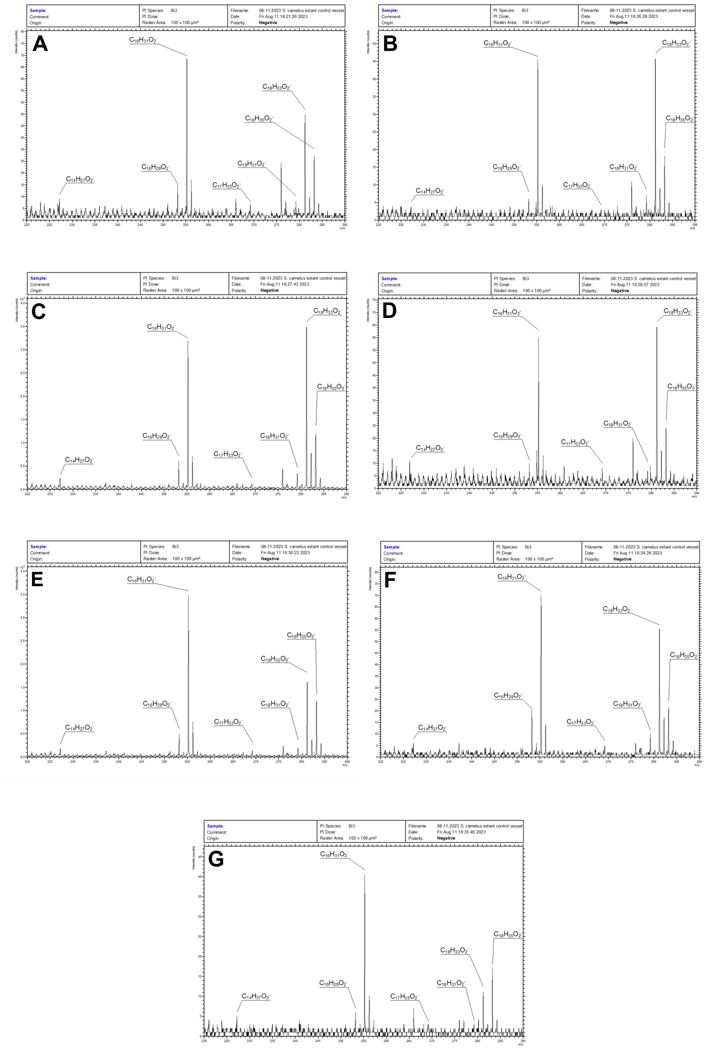


***Figure S19. ToF-SIMS spectra collected of extant S. camelus vascular tissue (A-G)*** *The negative ion spectra shown above were taken of the demineralized S. camelus vascular tissue post-collagenase digestion. As with the other extant specimens, secondary ion peaks corresponding to m/z*

*values for a variety of fatty acids are consistently observed. This is expected as the vascular tissue (basal endothelium) is cellular in nature, consisting substantially of phospholipids. Note that observed intensities for margaric and linoleic acids are attenuated relative to the extant B. taurus and A. mississippiensis vascular tissue. This possible difference may be taxon-related or could result from the age of the S. camelus bone as it has been stored in a laboratory setting for 16 years (the B. taurus and A. mississippiensis bones were obtained fresh for this study). Additionally, the comparative intensities of the palmitic and oleic acid molecular ions vary substantially between spectra. As discussed in the main text, this may be attributable to differences in membrane lipid composition compounded by the air-drying process exposing internal cytosolic membranes.*

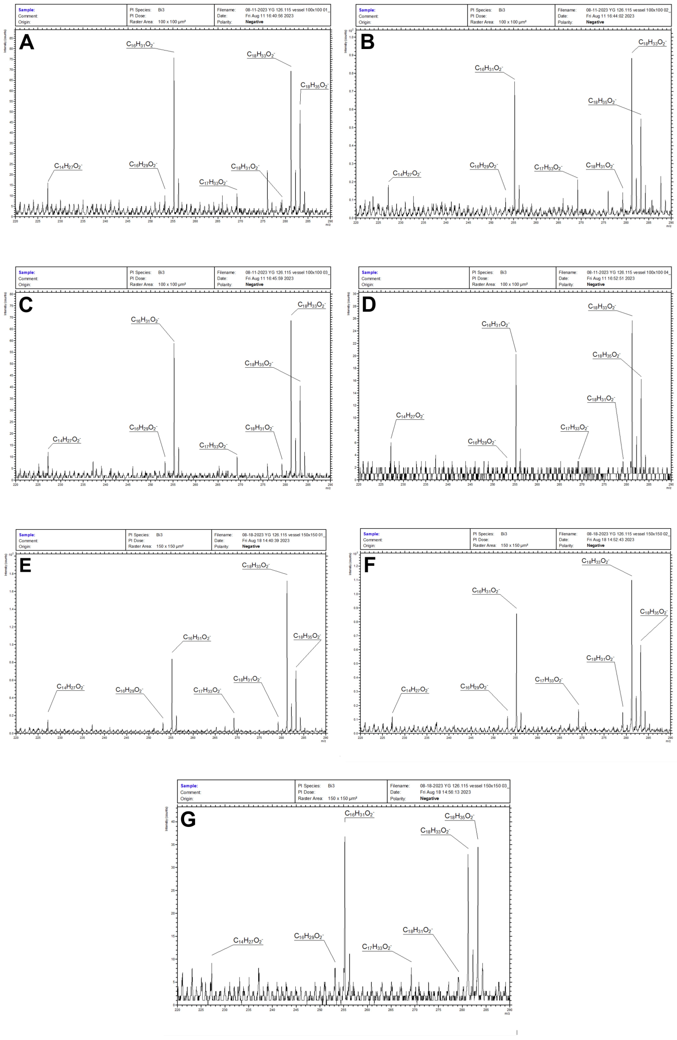


**Figure S20. ToF-SIMS spectra collected of vascular tissue from specimen YG 126.115 (A-G)** *The negative ion spectra shown above were taken of the demineralized YG 126.115 vascular tissue post-collagenase digestion. Unlike the other permafrost specimens, YG 126.115 consistently produced secondary ions with similar distribution and intensities to the extant control specimens shown in Figures S17-S19. Despite this specimen's ancient nature, molecular ions corresponding to both saturated and unsaturated fatty acids were observed. As the alkene bond(s) of unsaturated fatty acids make them particularly labile, this indicates this specimen is well-preserved and potentially a suitable candidate for palaeogenomic and palaeoproteomic*

*sequencing studies. YG 126.115 was also one of only two permafrost specimens (the other being YG 610.2364) in which structures matching osteocytes were observed.*

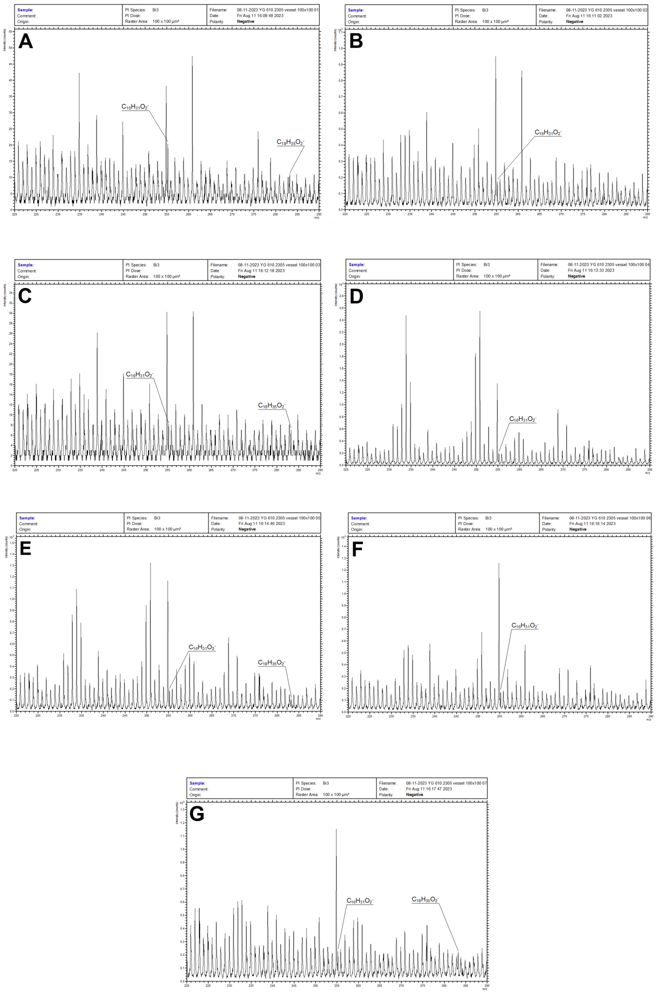


***Figure S21. ToF-SIMS spectra collected of vascular tissue from specimen YG 610.2305 (A-G)*** *The negative ion spectra shown above were taken of the demineralized YG 610.2305 vascular tissue post-collagenase digestion. Secondary ion peaks (of low intensity) corresponding to m/z values for palmitic and stearic acid are sporadically observed. Both are saturated fatty acids and expected to have a higher preservation potential relative to unsaturated*

*fatty acids (such as oleic acid, which was abundant in the extant vascular tissue spectra). However, the poor signal-to-noise ratio of these ion peaks and their sporadic distribution precludes any potential identification due to the ToF analyzer’s limited mass resolution. Thus the presence of fatty acids within the vascular tissue of specimen YG 610.2305 cannot be confirmed. Rather, the chemical composition of its “vascular tissue” is supported as having undergone substantial diagenesis. This is generally the case for vascular tissue from all 5 Little Blanche Creek permafrost specimens.*

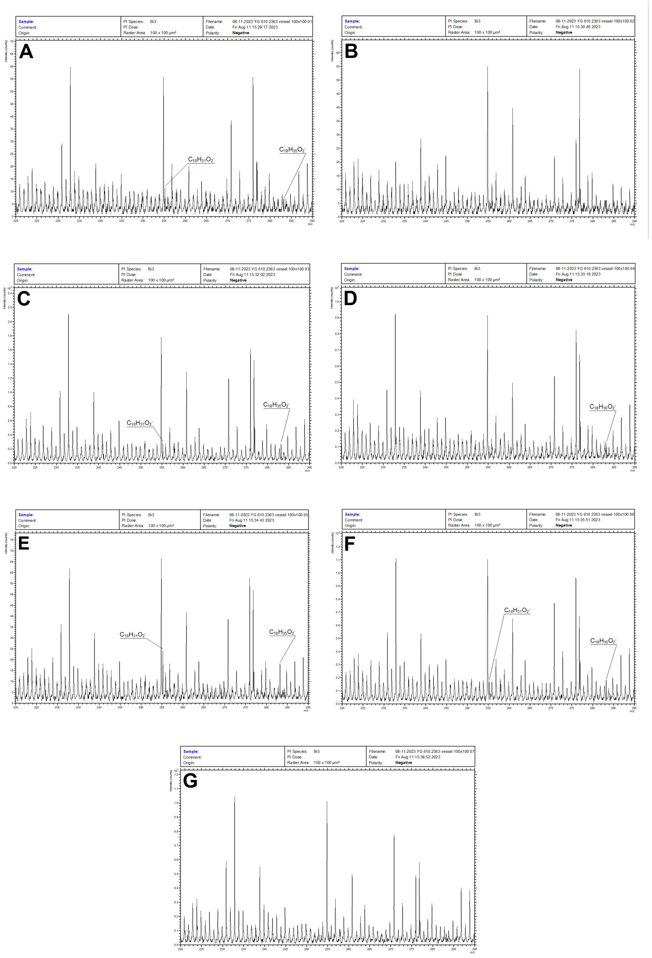

***Figure S22. ToF-SIMS spectra collected of vascular tissue from specimen YG 610.2363 (A-G)*** *The negative ion spectra shown above were taken of the demineralized YG 610.2363 vascular tissue post-collagenase digestion. As with the other Little Blanche Creek permafrost specimens, secondary ion peaks (of low intensity) corresponding to m/z values for palmitic and stearic acid are sporadically observed and cannot be confidently identified as to source molecule. This was also one of two specimens in which fungal hyphae were observed to have colonized the vascular canals.*

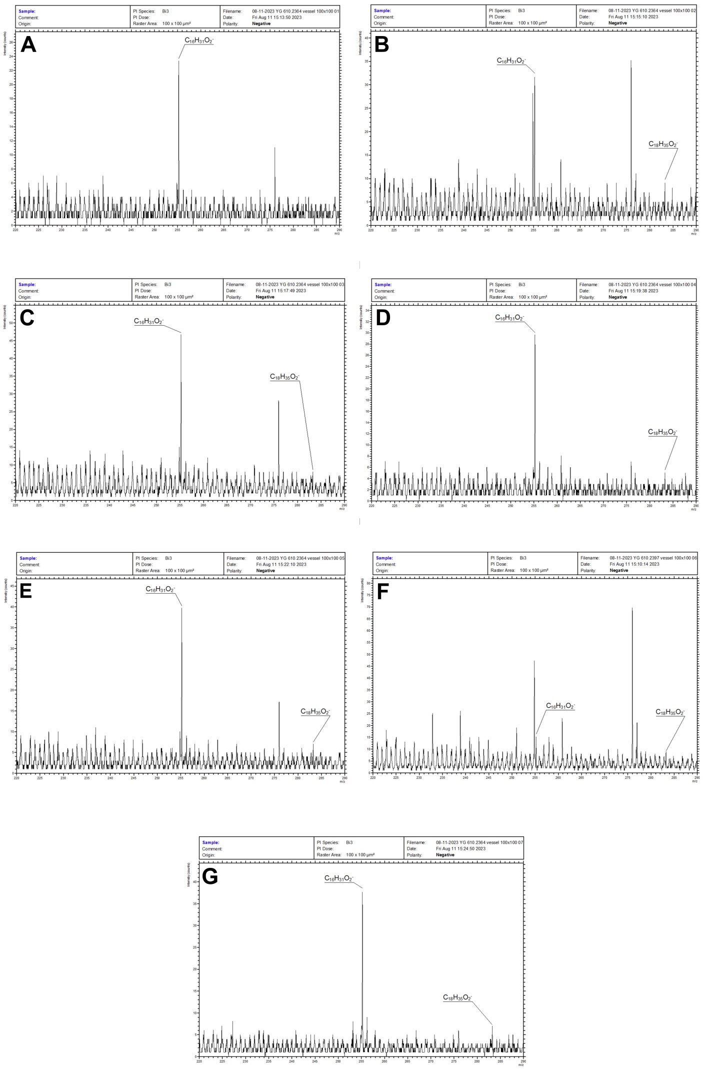

***Figure S23. ToF-SIMS spectra collected of vascular tissue from specimen YG 610.2364 (A-G)*** *The negative ion spectra shown above were taken of the demineralized YG 610.2364 vascular tissue post-collagenase digestion. As with the other Little Blanche Creek permafrost specimens, secondary ion peaks corresponding to m/z values for palmitic and stearic acid are sporadically observed and cannot be confidently identified as to source molecule. However, the intensity of the "fatty acid" ions is consistently higher for YG 610.2364 relative to the vascular tissue of the other Little Blanche Creek permafrost specimens, possibly suggesting better preservation. YG 610.2364 was also the only permafrost specimen from Little Blanche Creek in which structures matching osteocytes (not shown) were observed.*

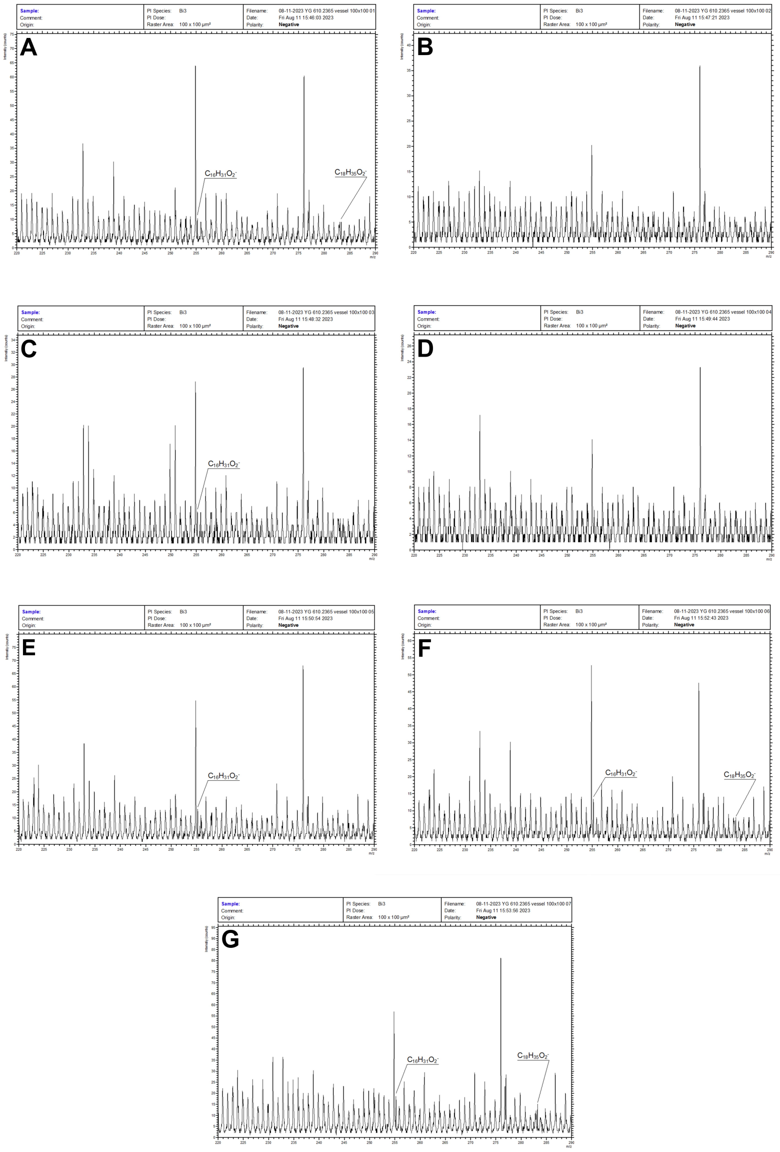


***Figure S24. ToF-SIMS spectra collected of vascular tissue from specimen YG 610.2365 (A-G)*** *The negative ion spectra shown above were taken of the demineralized YG 610.2365 vascular tissue post-collagenase digestion. Like the other Little Blanche Creek permafrost specimens, secondary ion peaks (of poor signal-to-noise) corresponding to m/z values for palmitic and stearic acid are sporadically observed but cannot be confidently identified as to source molecule. This was also one of two specimens in which*

*fungal hyphae were observed to have colonized the vascular canals.*

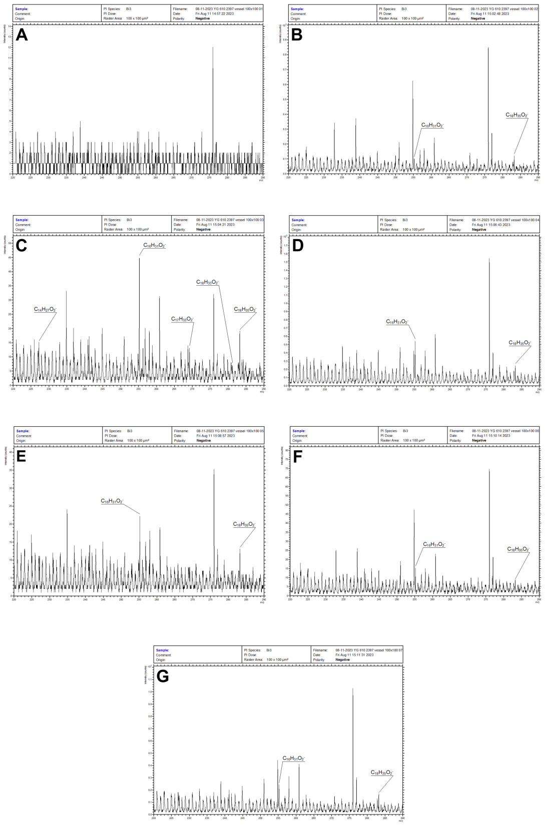


***Figure S25. ToF-SIMS spectra collected of vascular tissue from specimen YG 610.2397 (A-G)*** *The negative ion spectra shown above were taken of the demineralized YG 610.2397 vascular tissue post-collagenase digestion. Secondary ion peaks (generally of low intensity) corresponding to m/z values for palmitic and stearic acid are sporadically observed and cannot be confidently identified as to source molecule. Spectrum S25(C) does show a distribution of “fatty acid molecular ion” peaks closer to that of the extant controls, but little can be determined from this single lone spectrum.*
