## Supplementary figures and images for "Nanoscale Imaging and Microanalysis of Ice Age Bone Offers New Perspective on “Subfossils” and Fossilization"

### 01-16-2023 ostrich collagen 400x 01.TIF

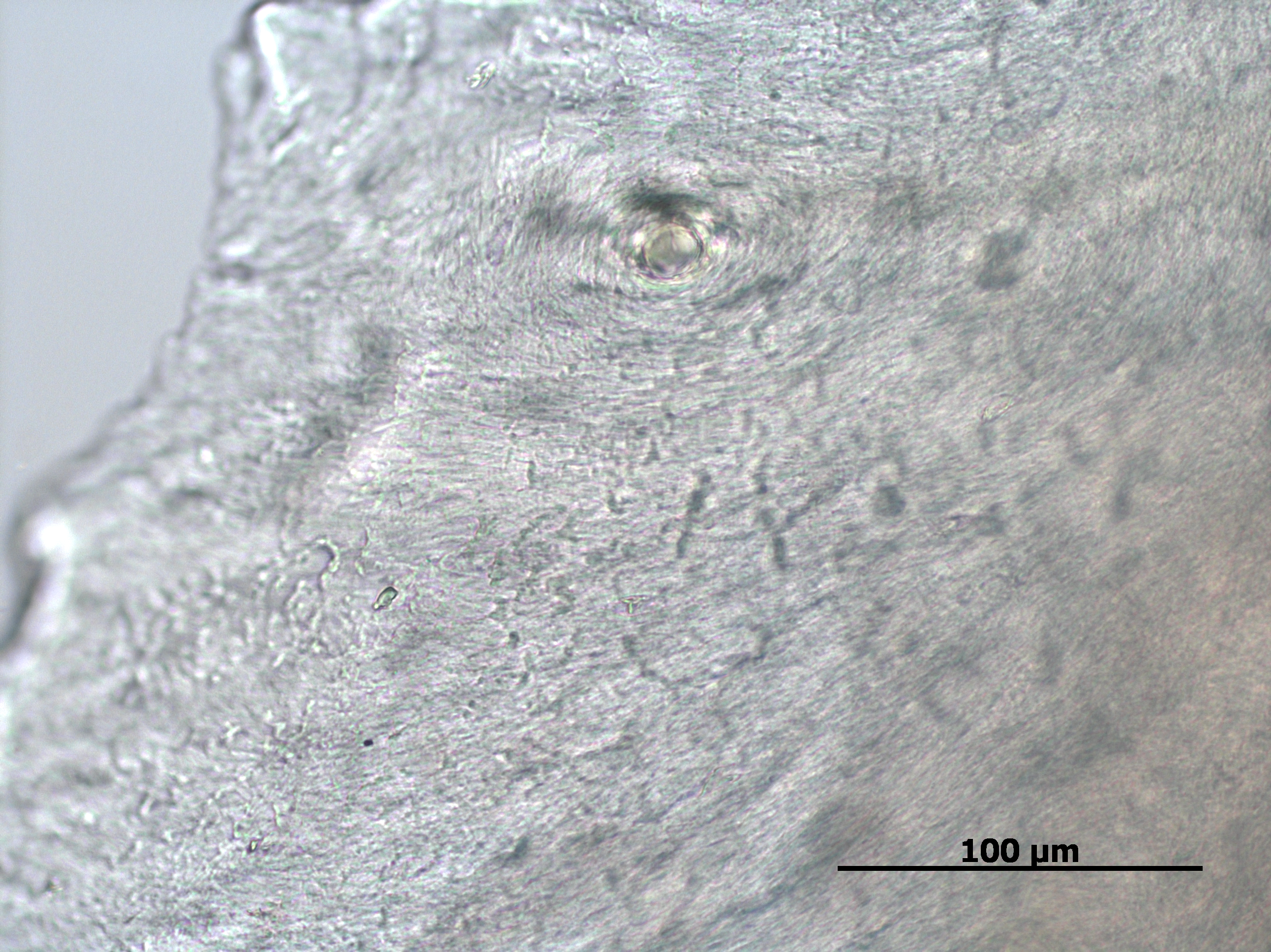

### 01-16-2023 steppe bison radius collagen 400x 01.TIF

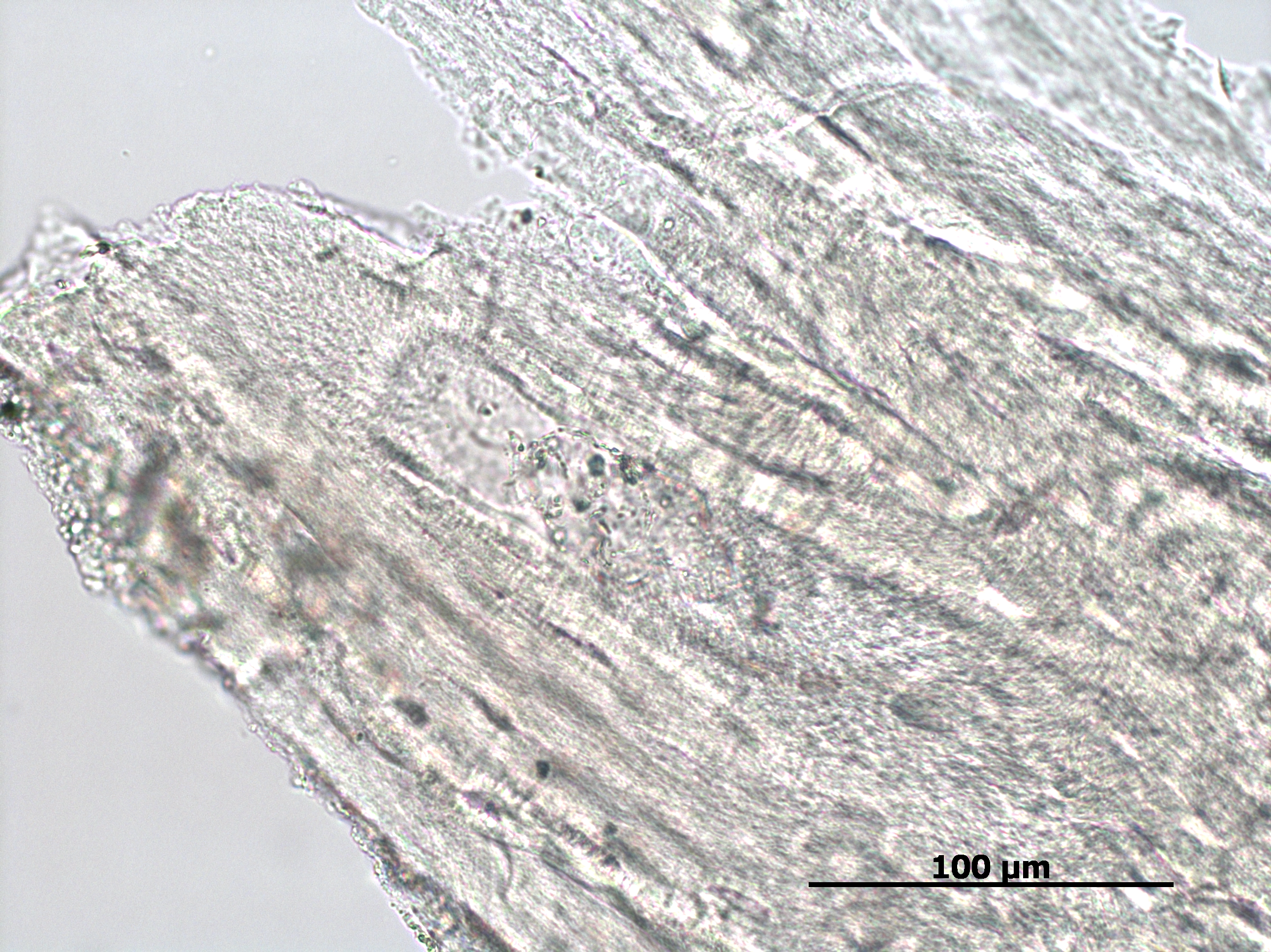

### 01-16-2023 steppe bison tibia with soft tissue collagen 400x 01.TIF

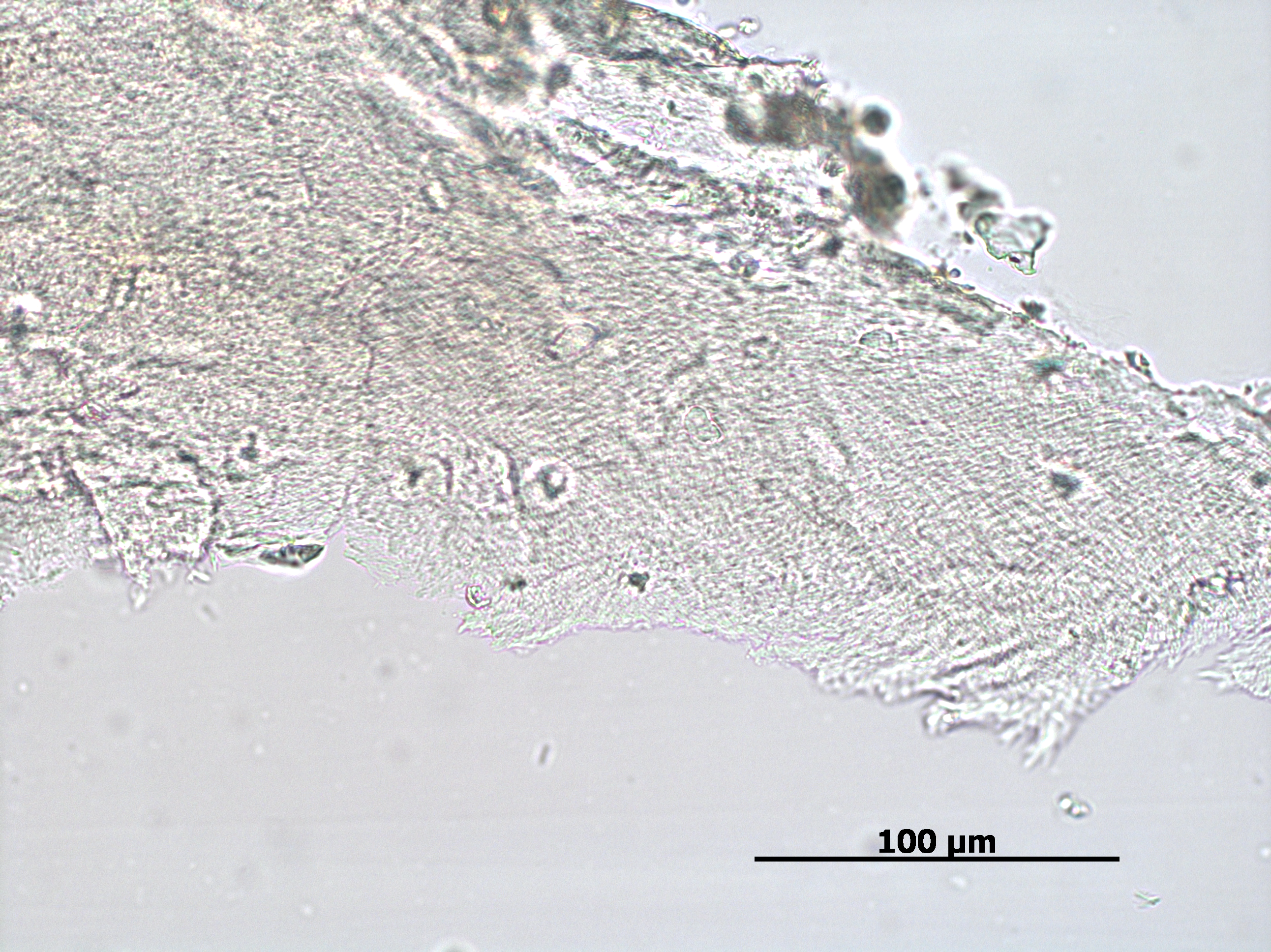

### 01-16-2023 woolly mammoth collagen 400x 01.TIF

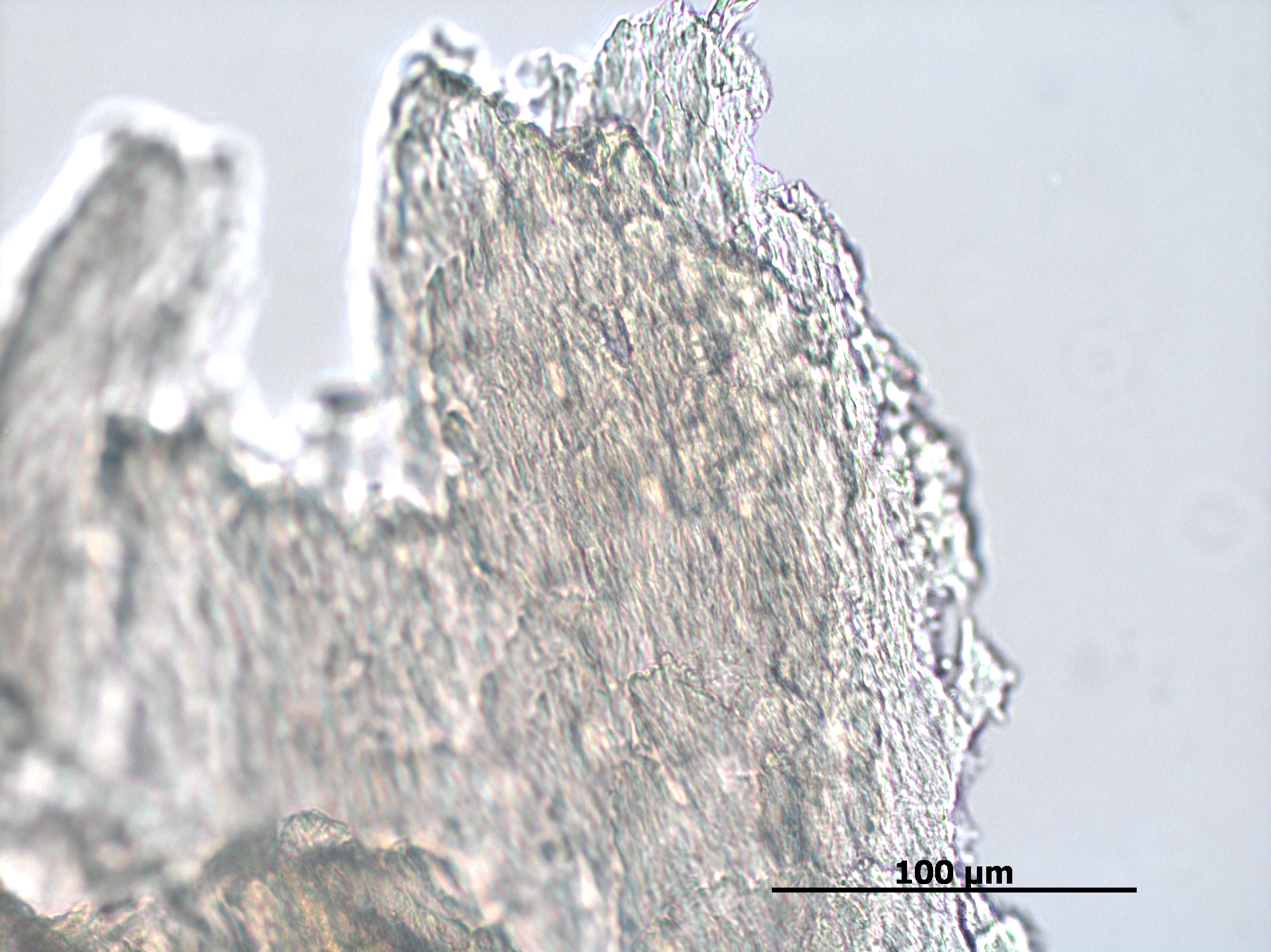

### 01-24-2023 ostrich vessel 400x 02.TIF

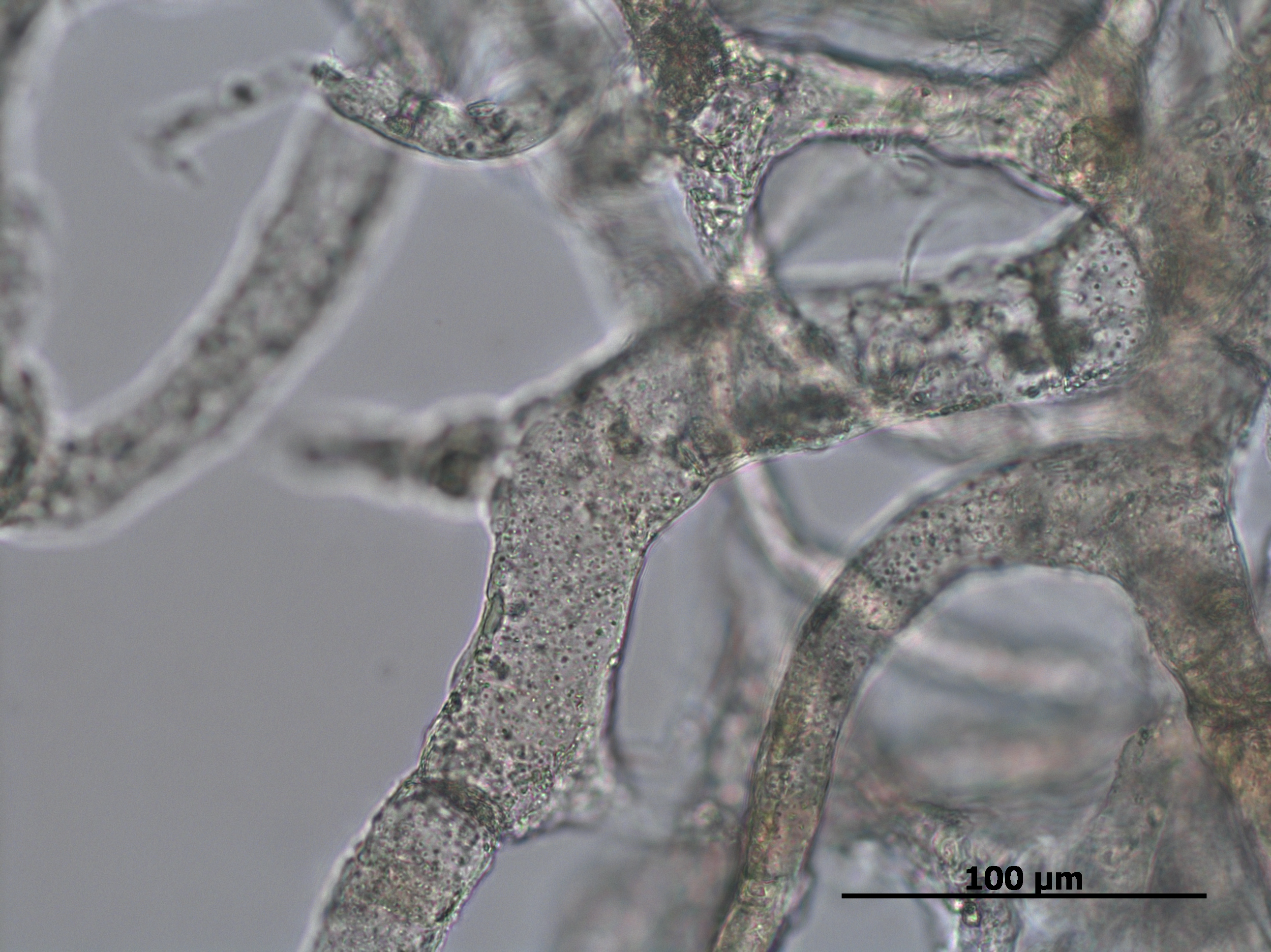

### 01-24-2023 yukon steppe bison tibia vessel 400x 01.TIF

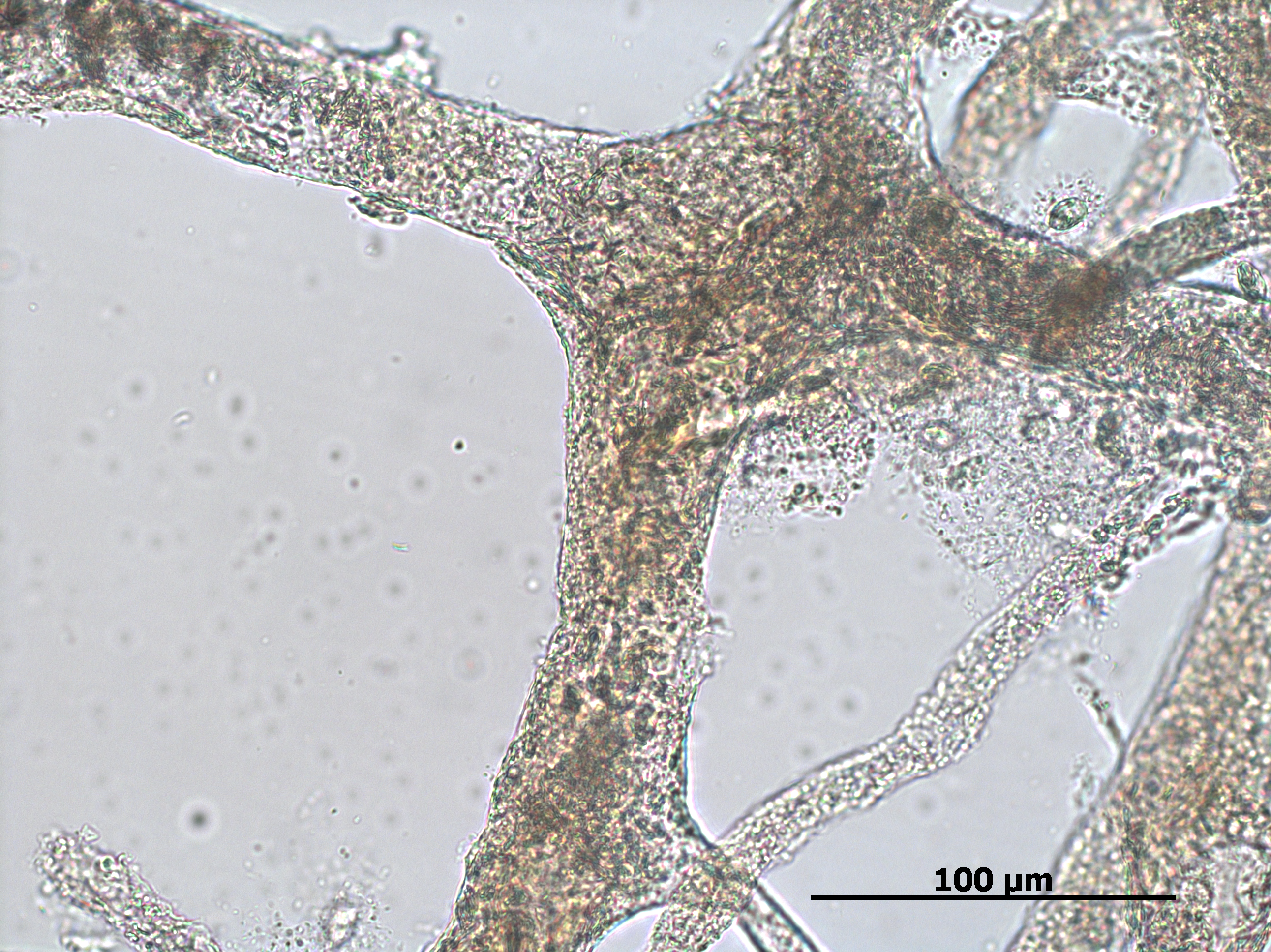

### 01-26-2023 steppe bison tibia osteocytes_m01.tif

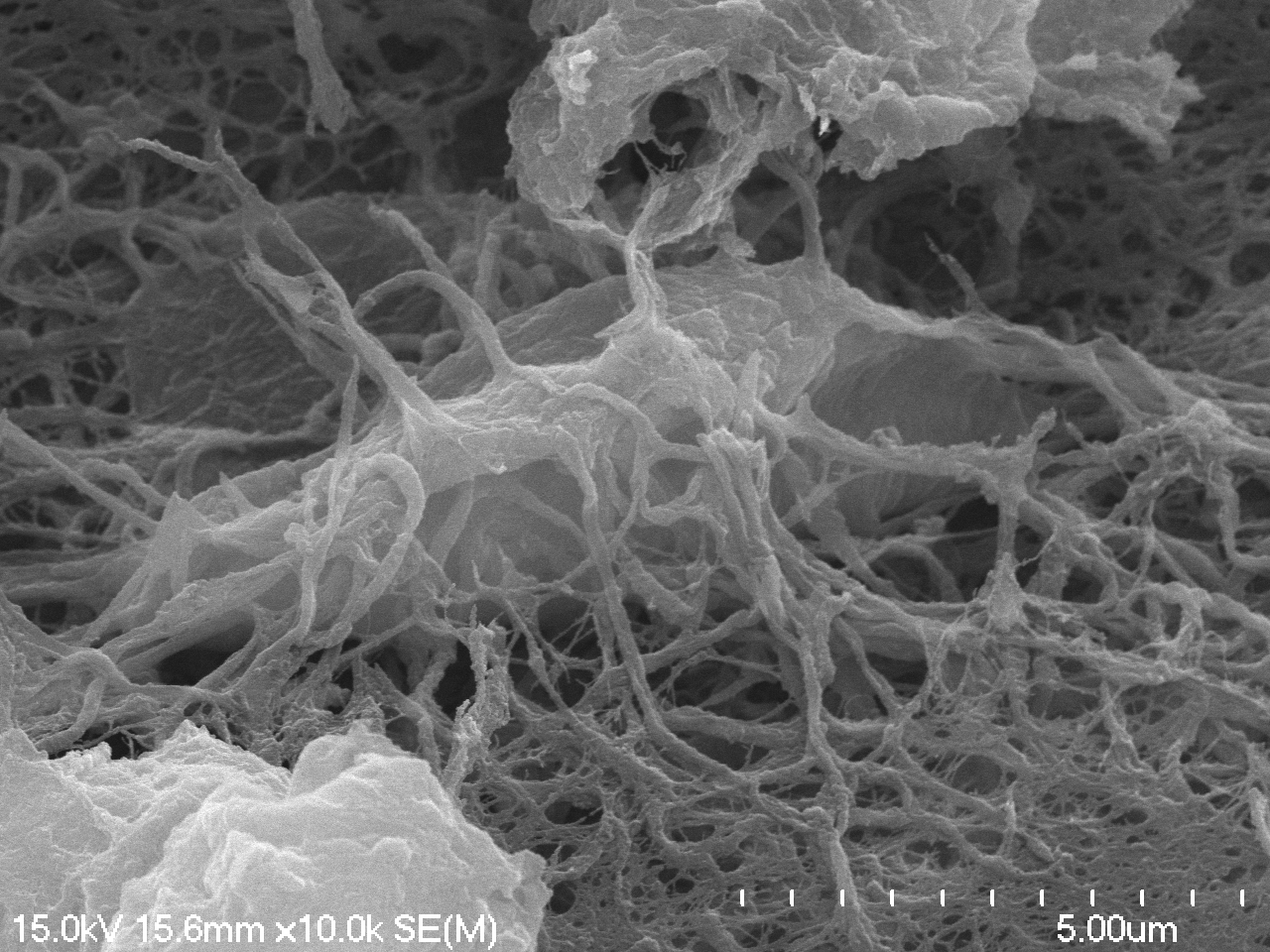

### 01-26-2023 steppe bison tibia vessels_m05.tif

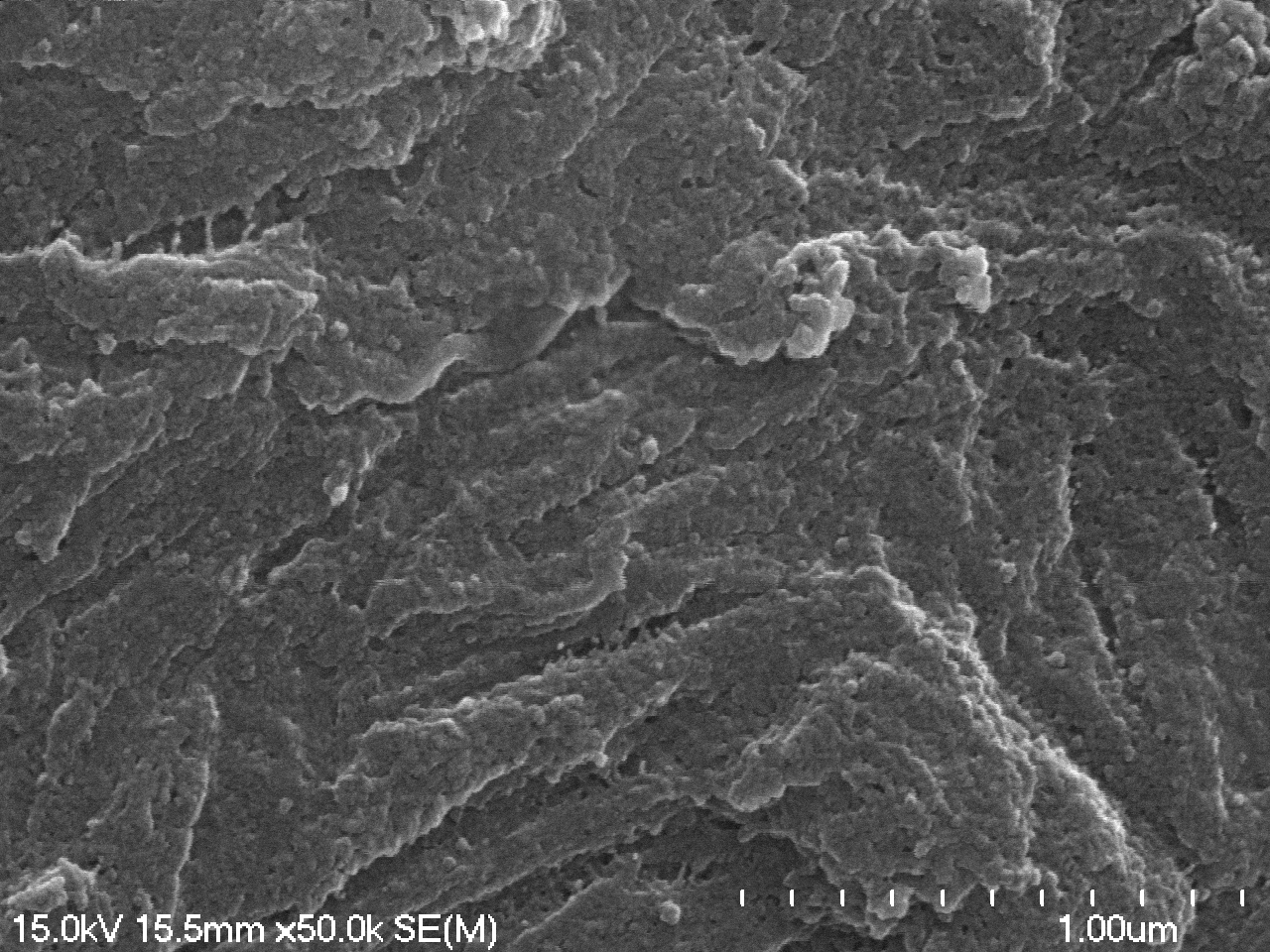

### 01-26-2023 woolly mammoth vessels_m05.tif

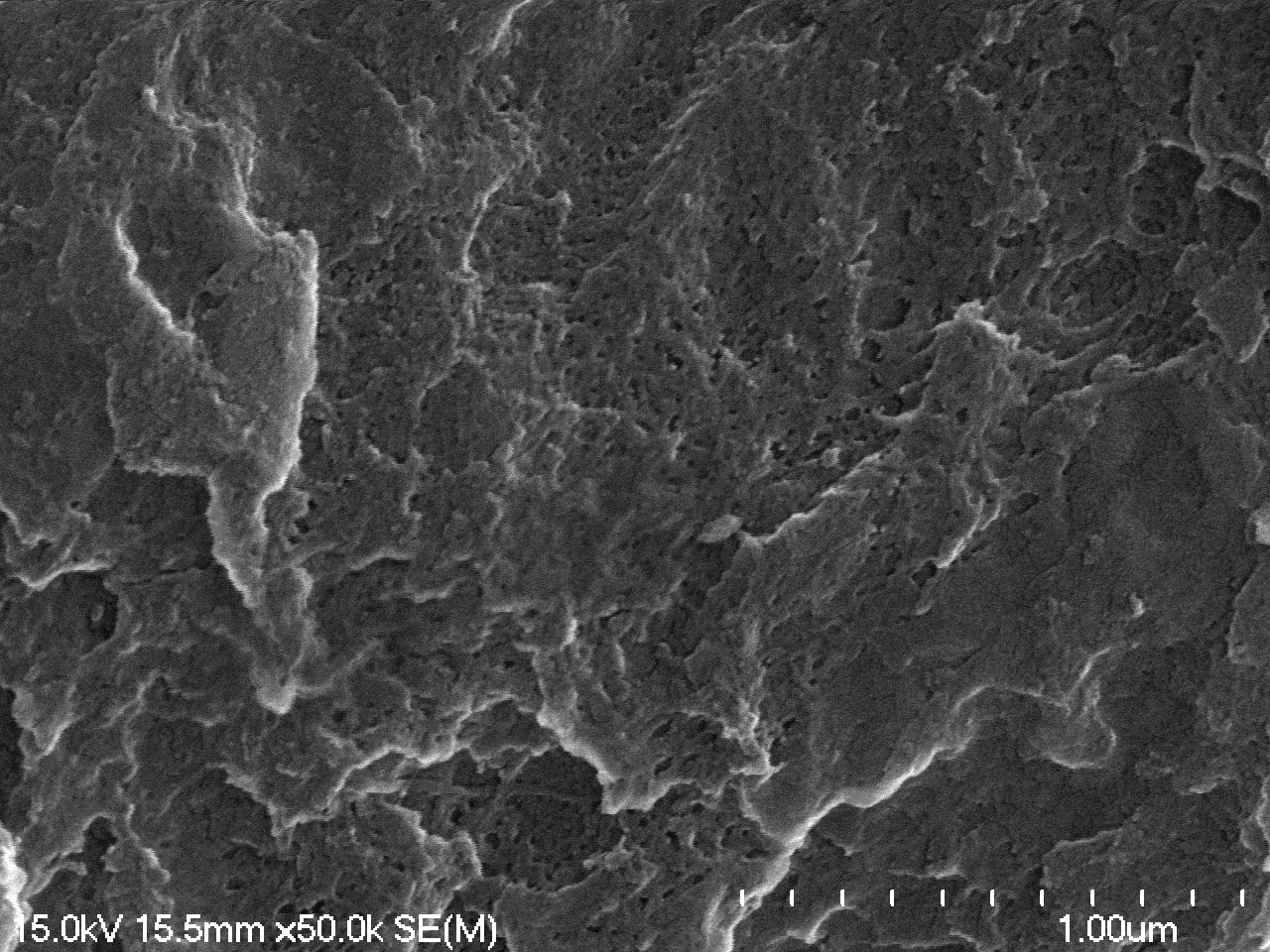

### 01-26-2023 woolly mammoth vessels_m06.tif

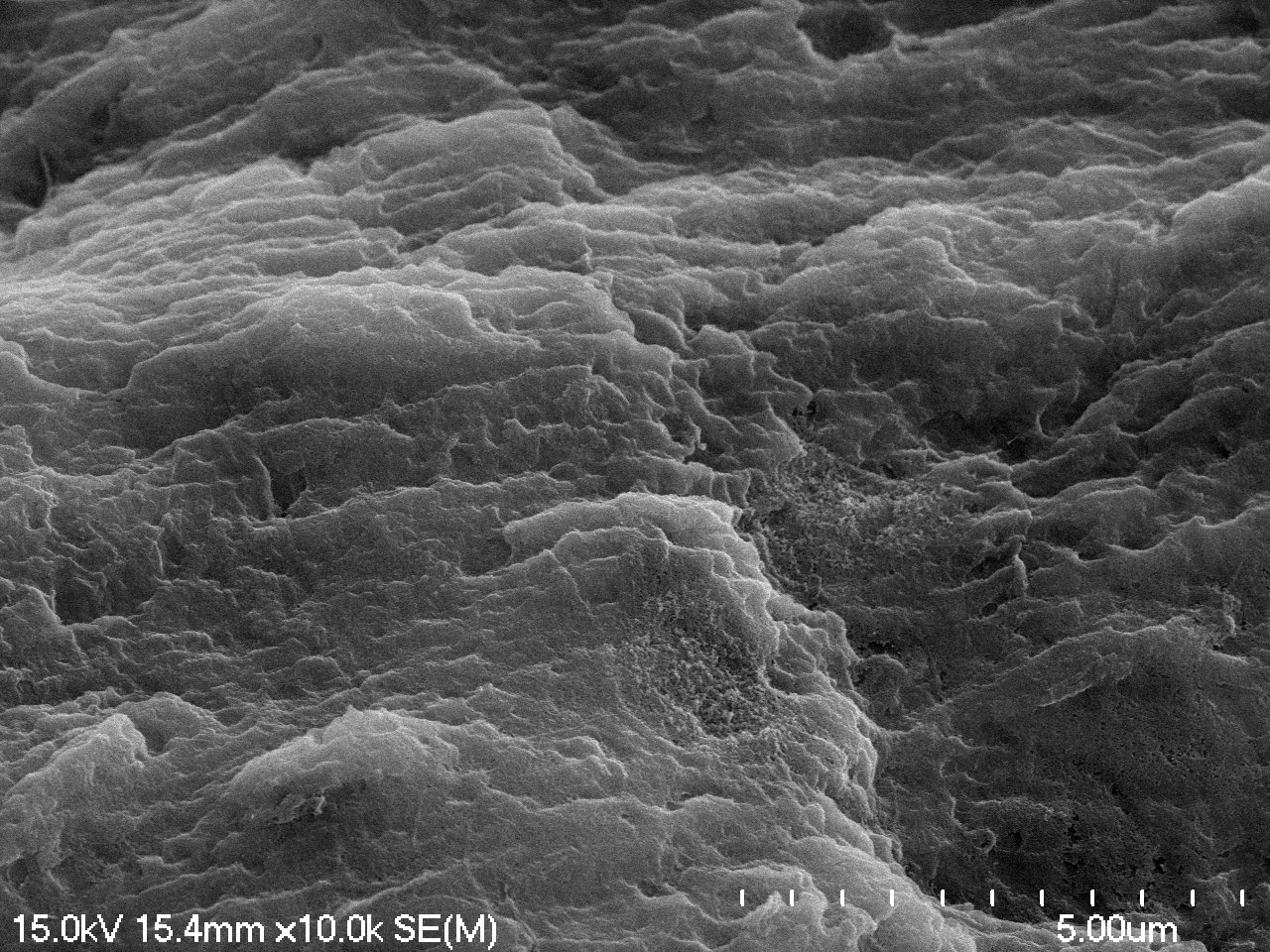

### 01-26-2023 woolly mammoth vessels_m07.tif

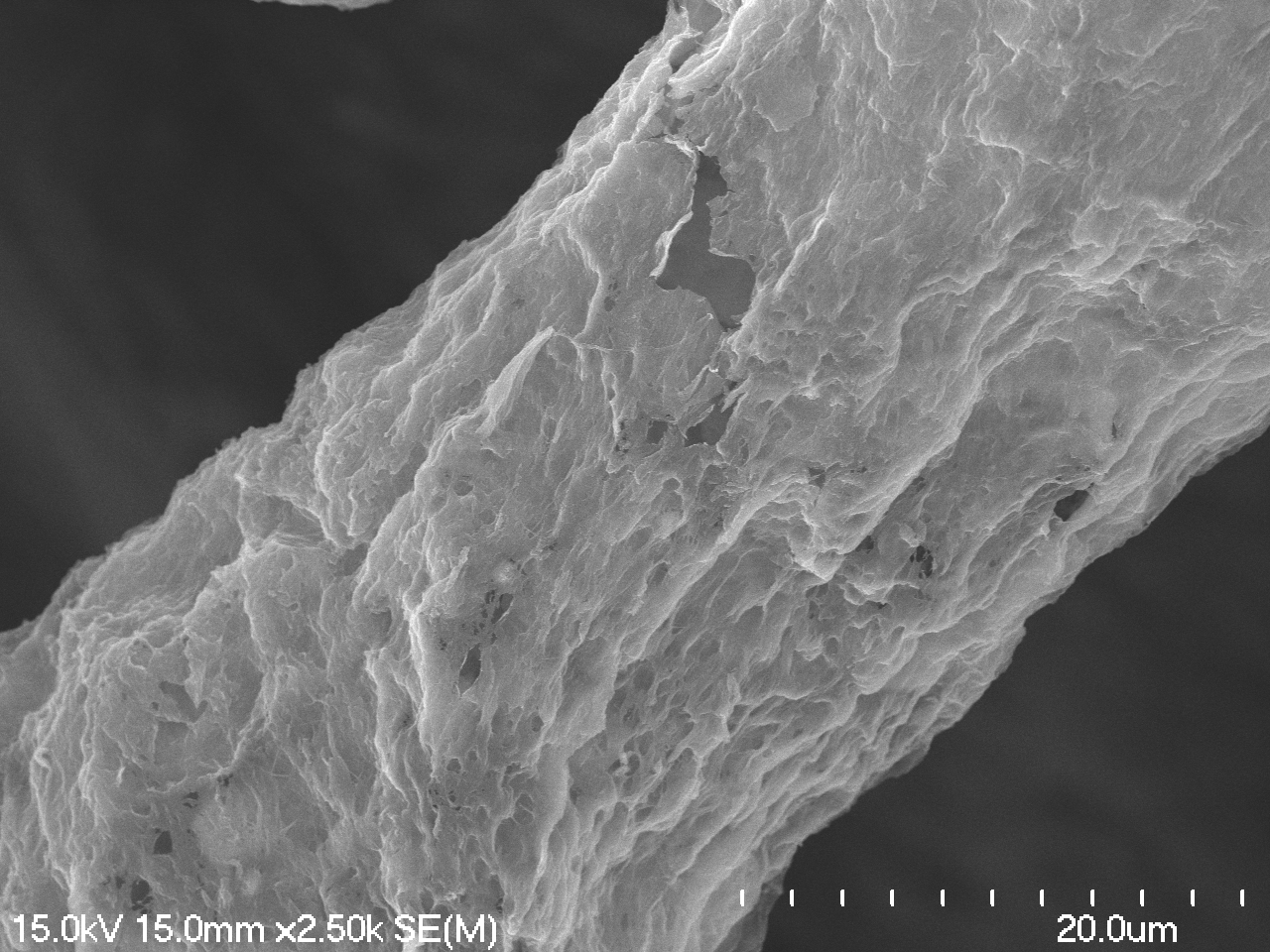

### 05-25-2023 cow collagen_m01_m03.tif

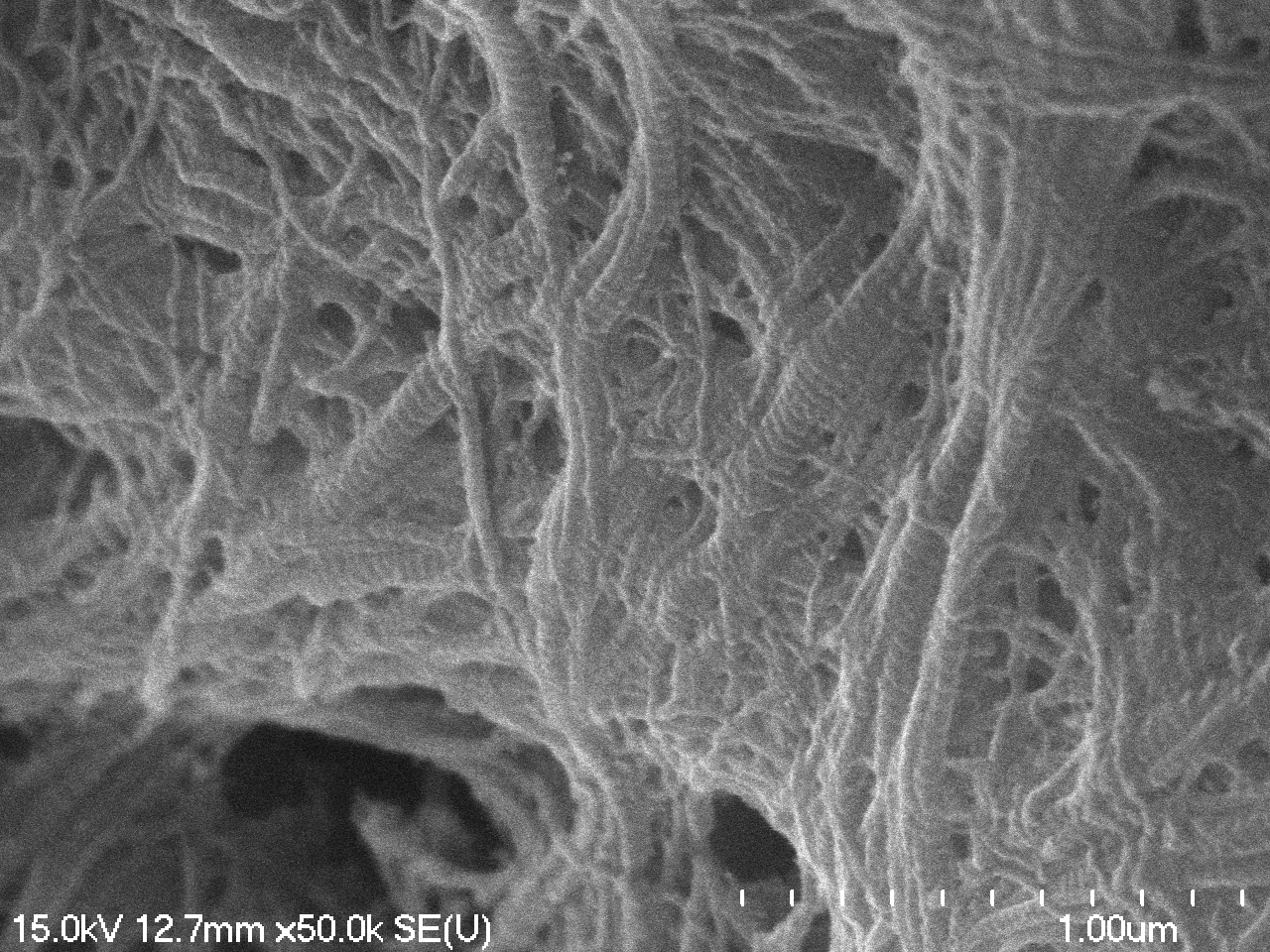

### 05-25-2023 cow collagen_m01_m05.tif

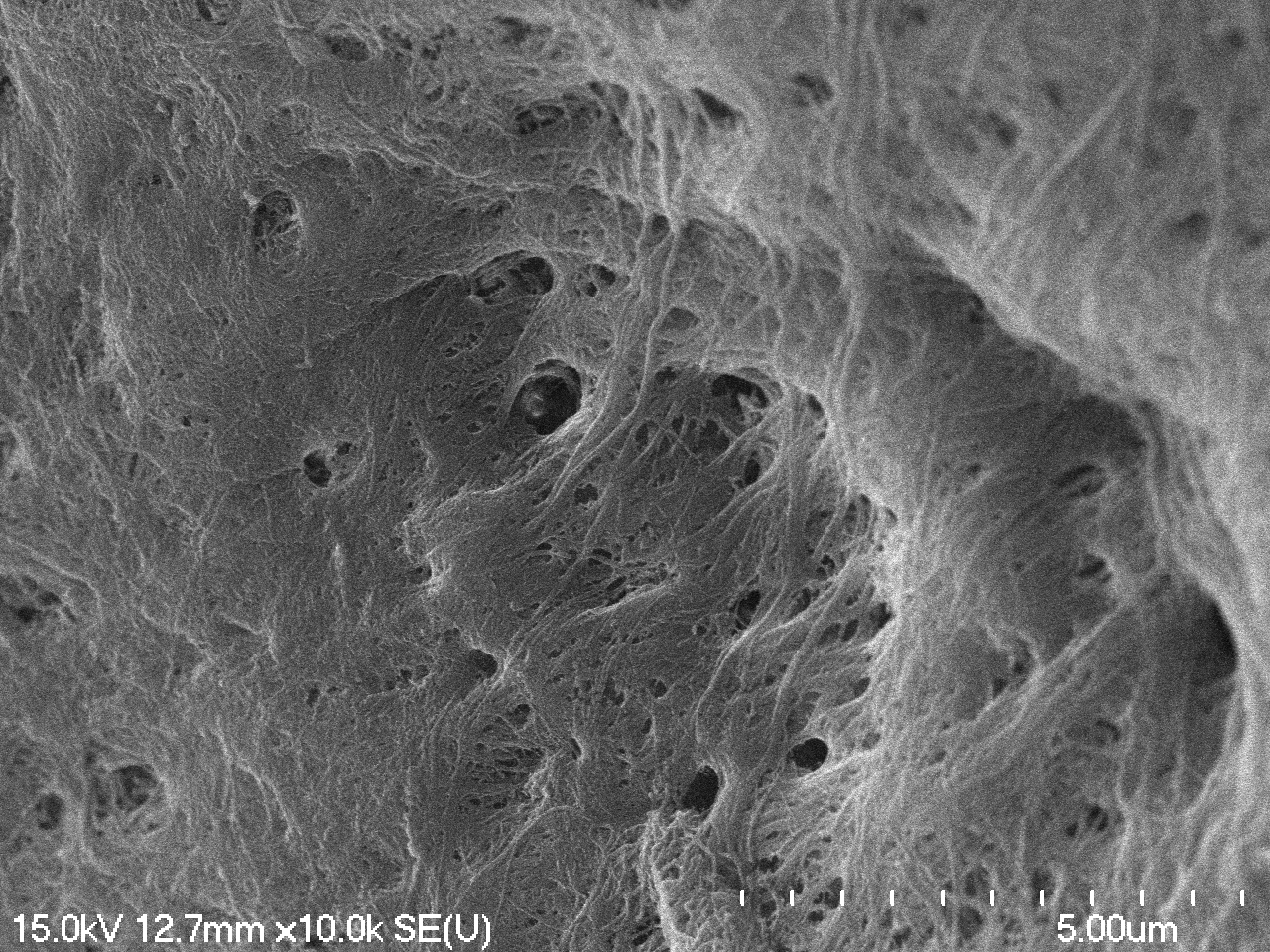

### 05-25-2023 cow collagen_m08.tif

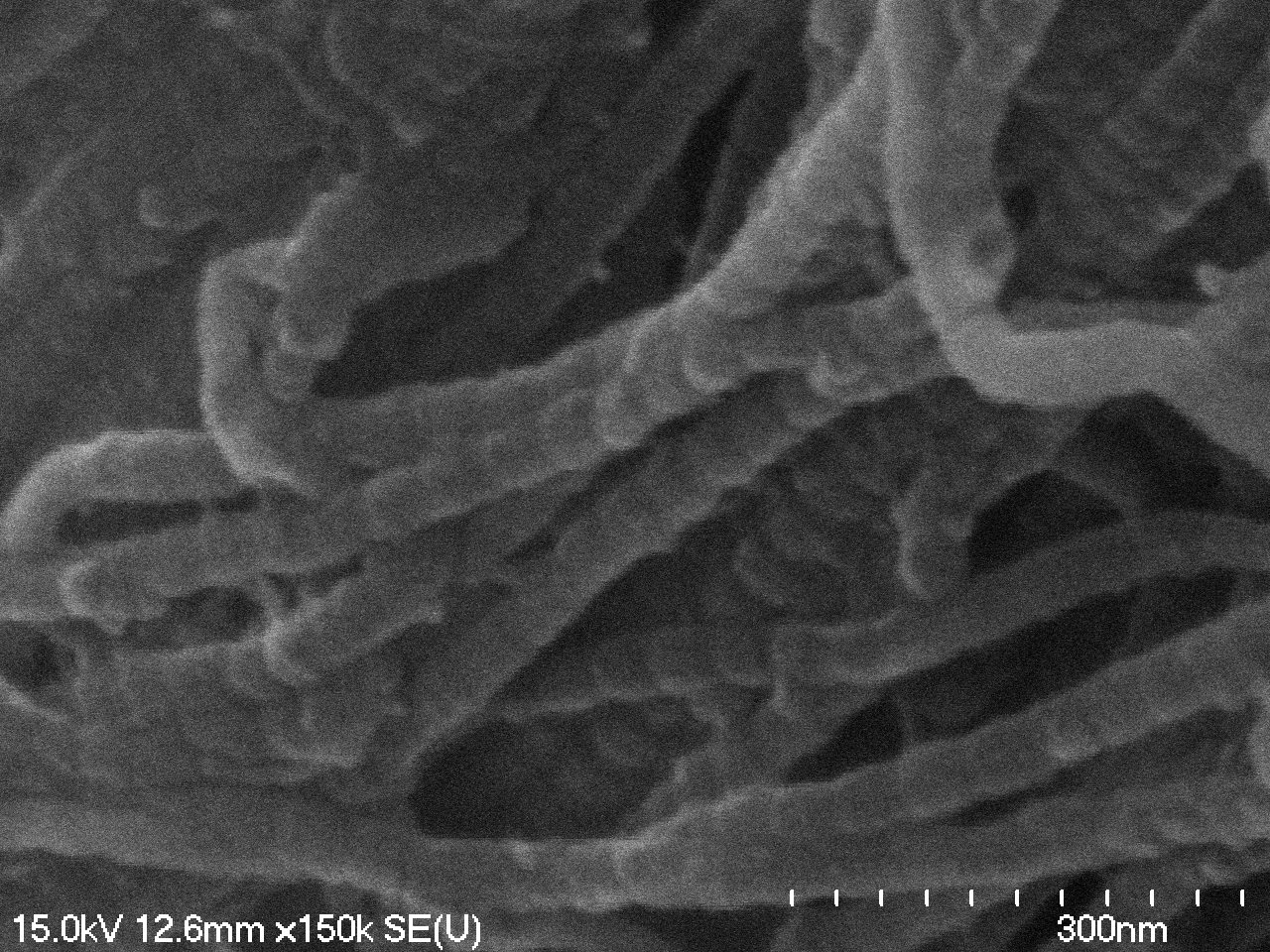

### 05-25-2023 horse collagen_m0_m01.tif

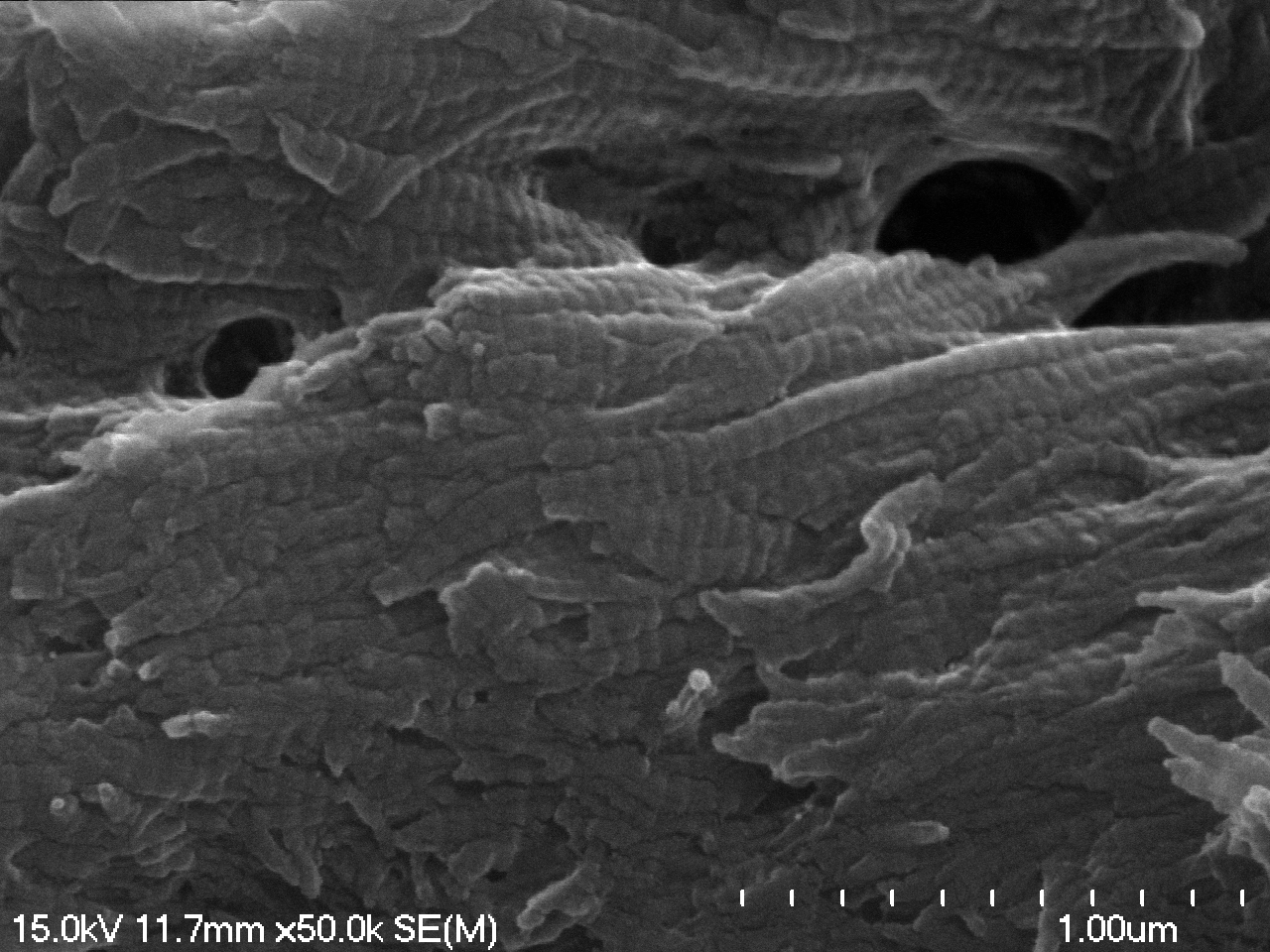

### 05-25-2023 horse collagen_m0_m02.tif

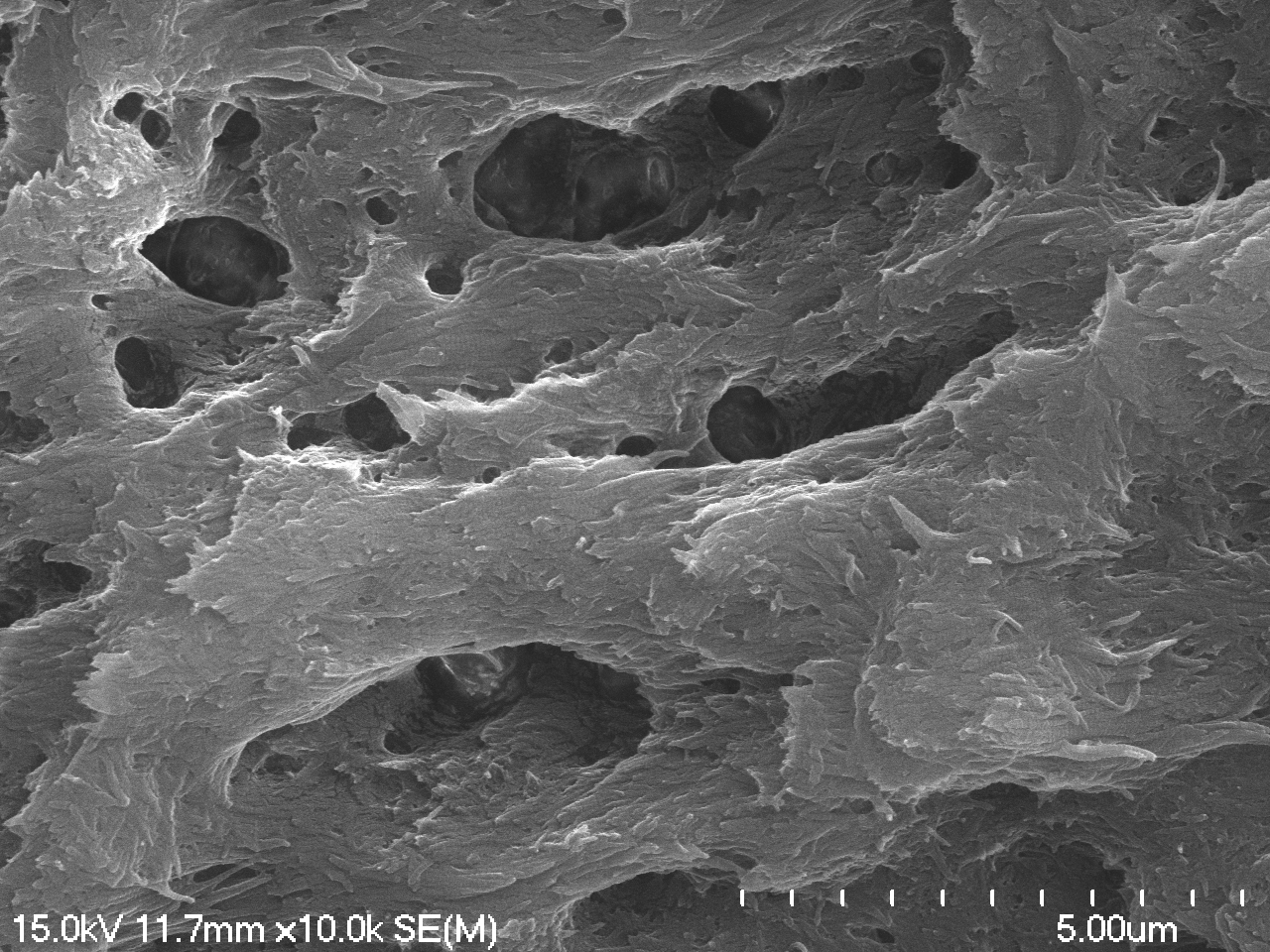

### 05-25-2023 horse collagen_m10.tif

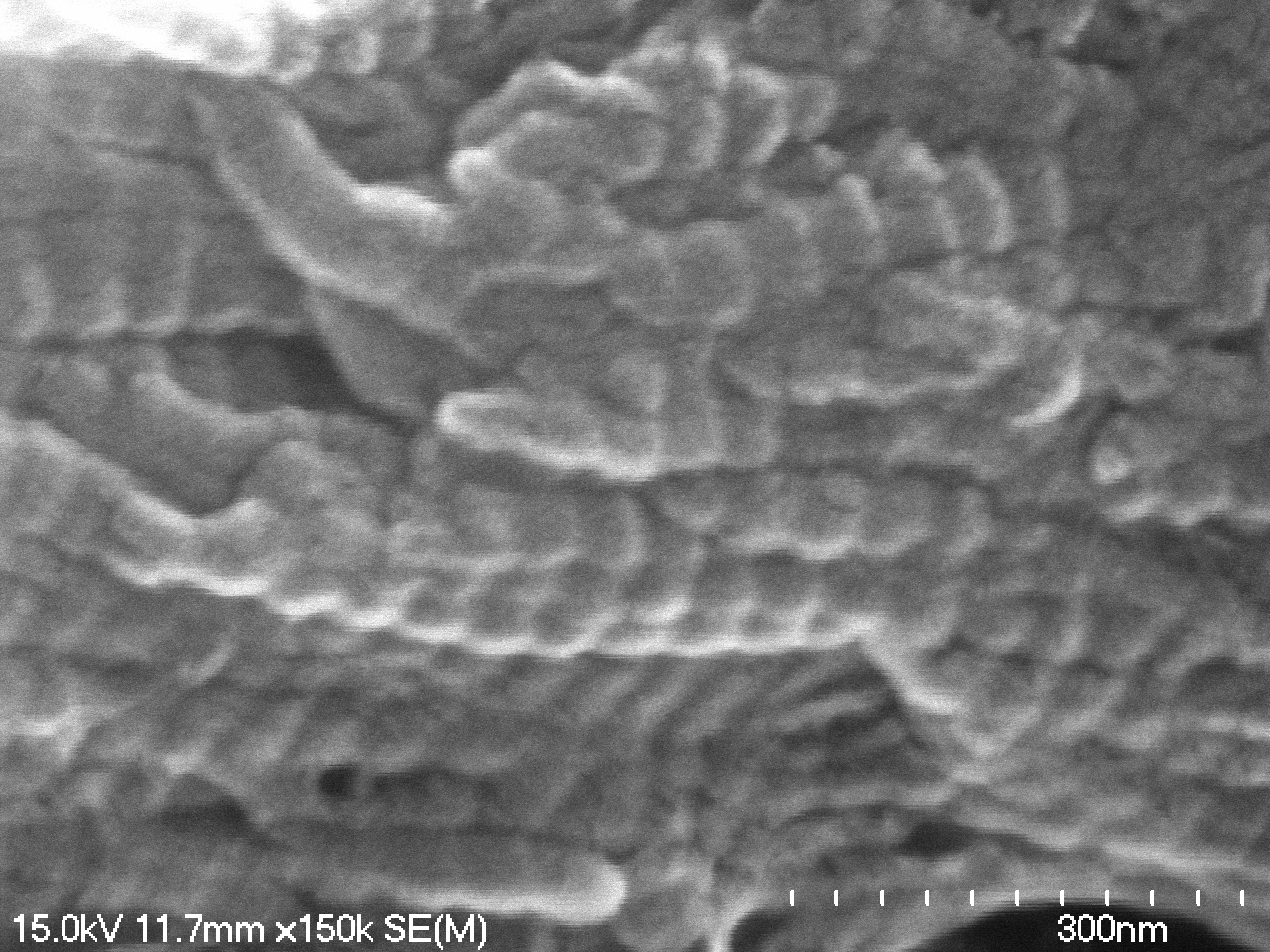

### 05-25-2023 mammoth collagen_m06_m01.tif

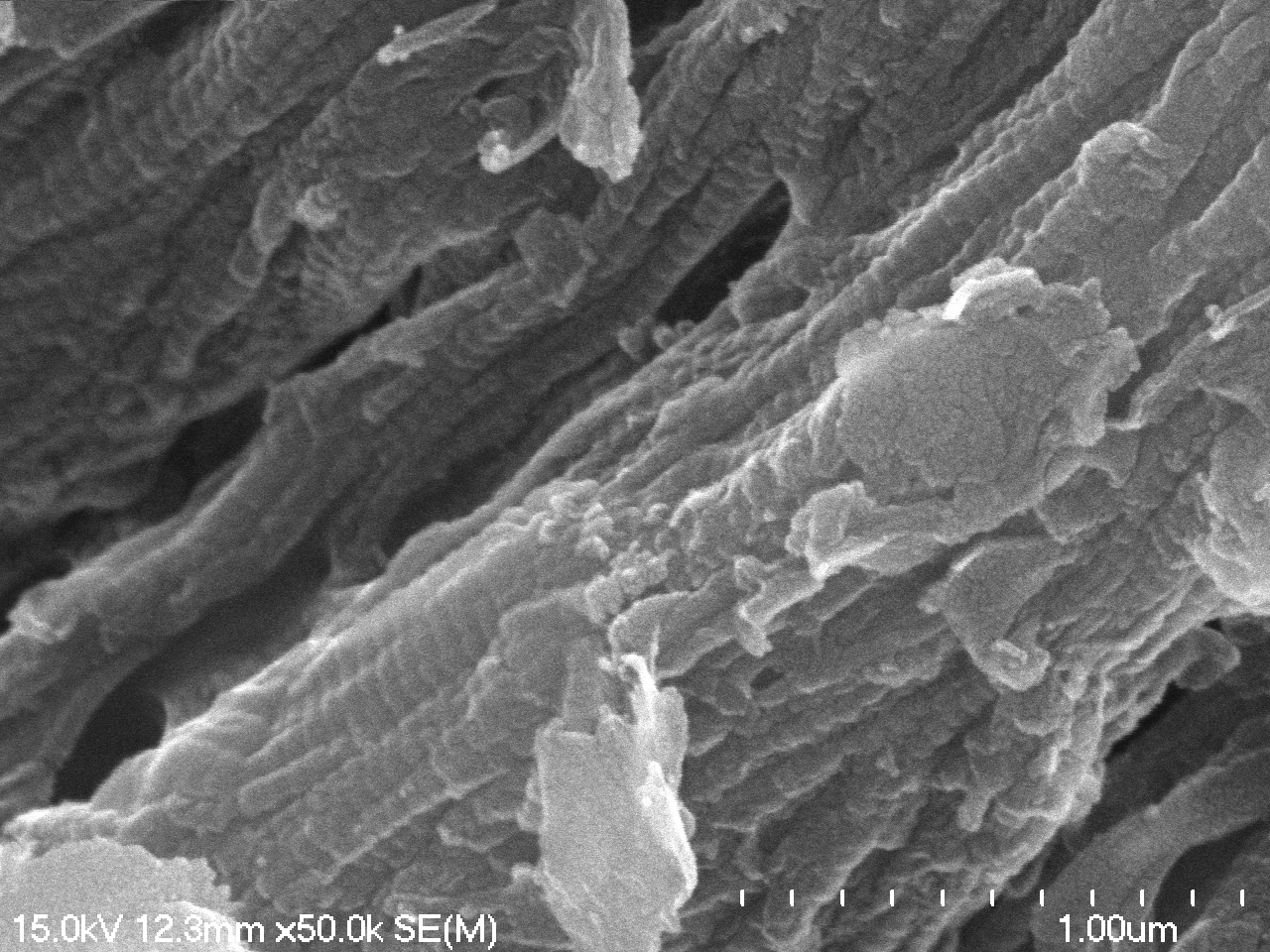

### 05-25-2023 mammoth collagen_m06_m05.tif

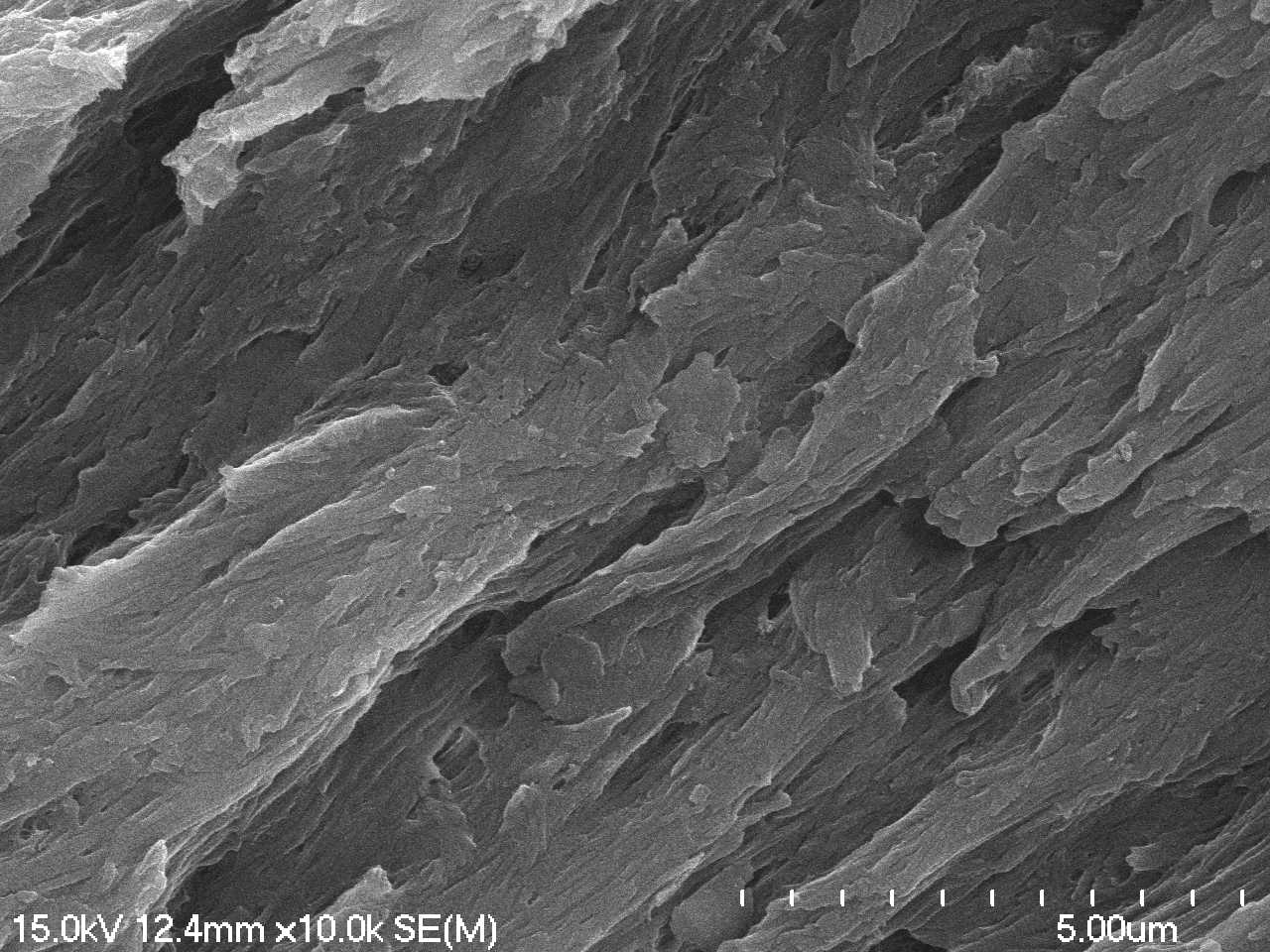

### 05-25-2023 mammoth collagen_m11.tif

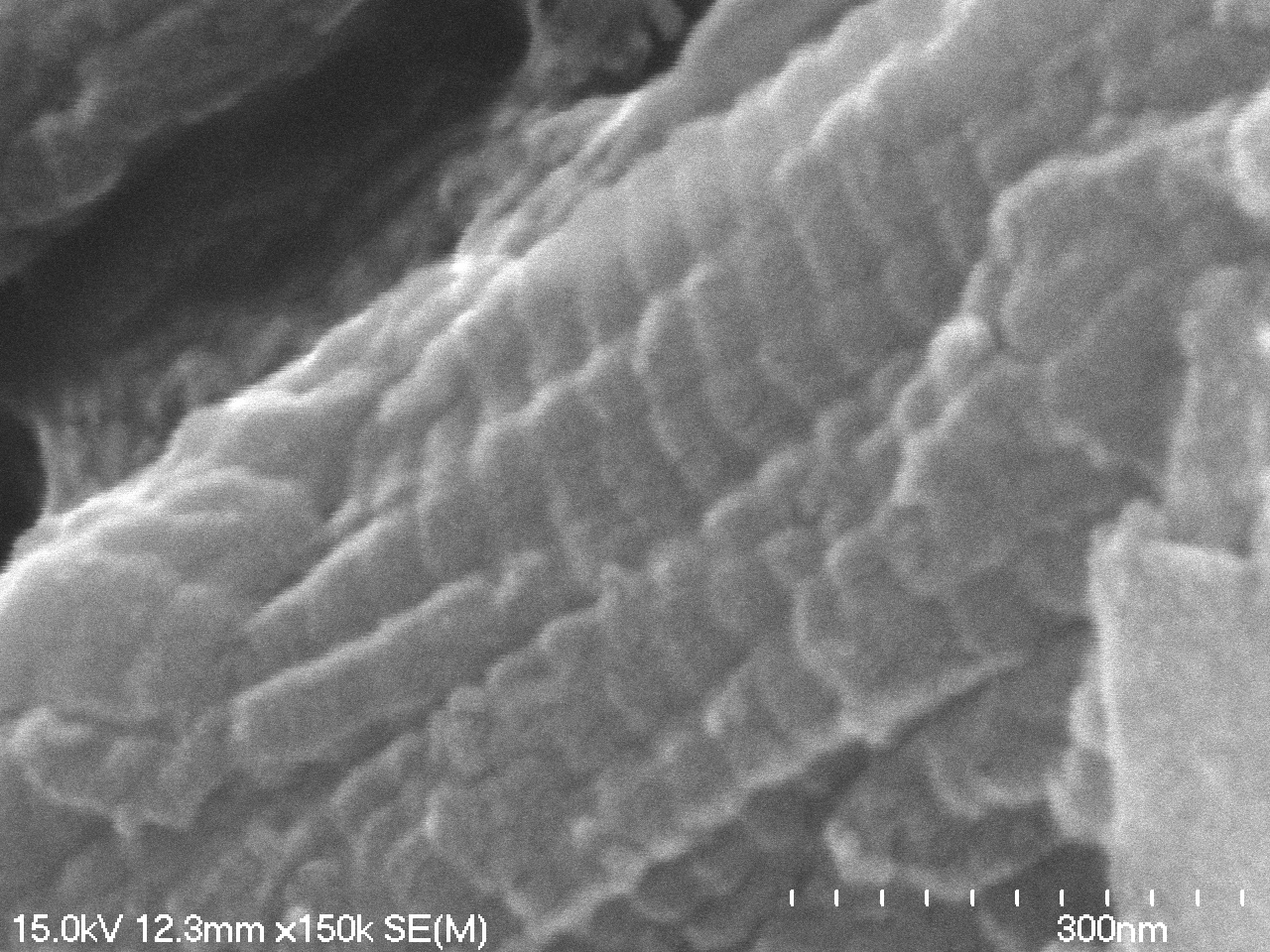

### 05-25-2023 reindeer collagen_m02_m05.tif

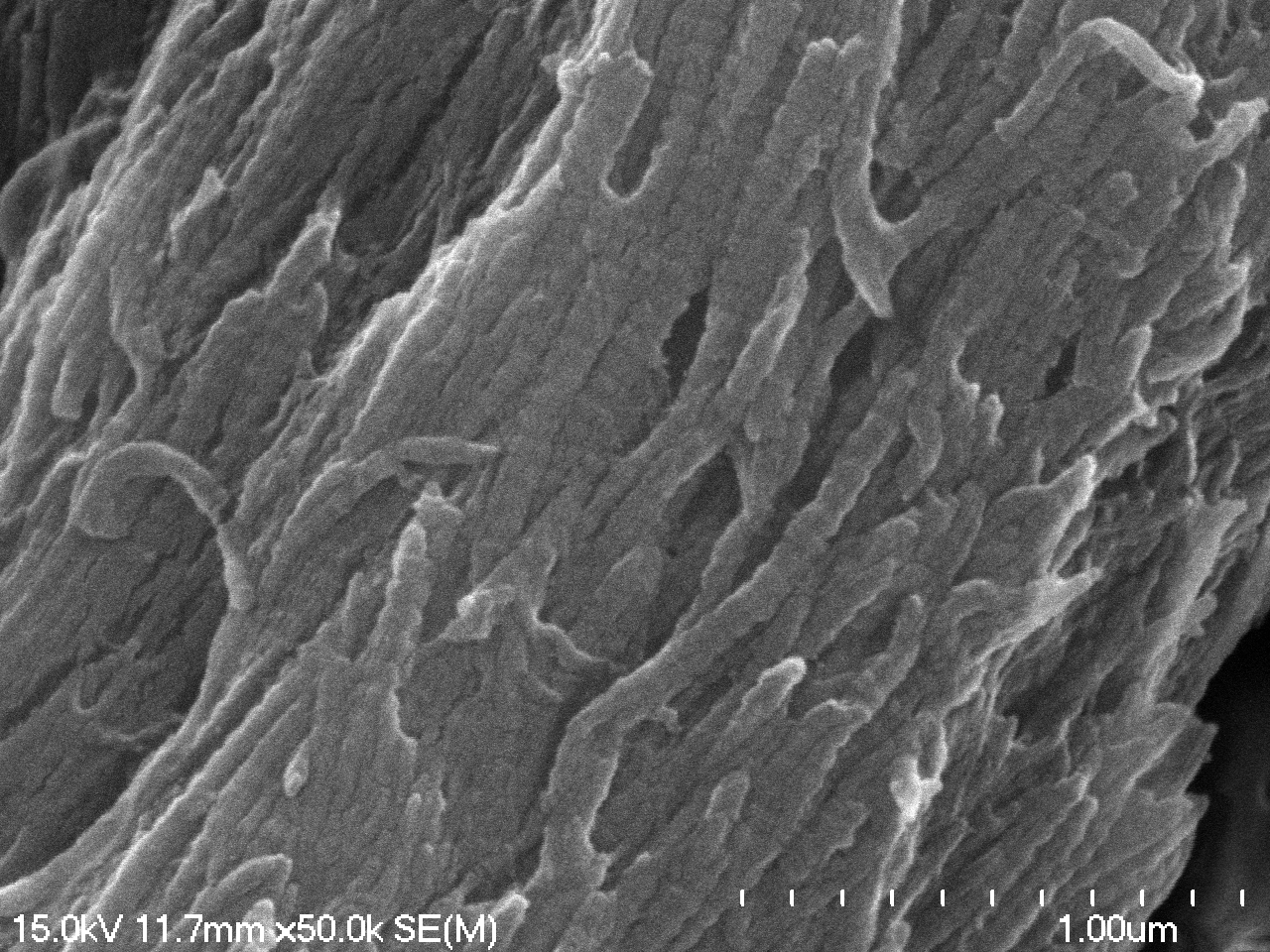

### 05-25-2023 reindeer collagen_m05.tif

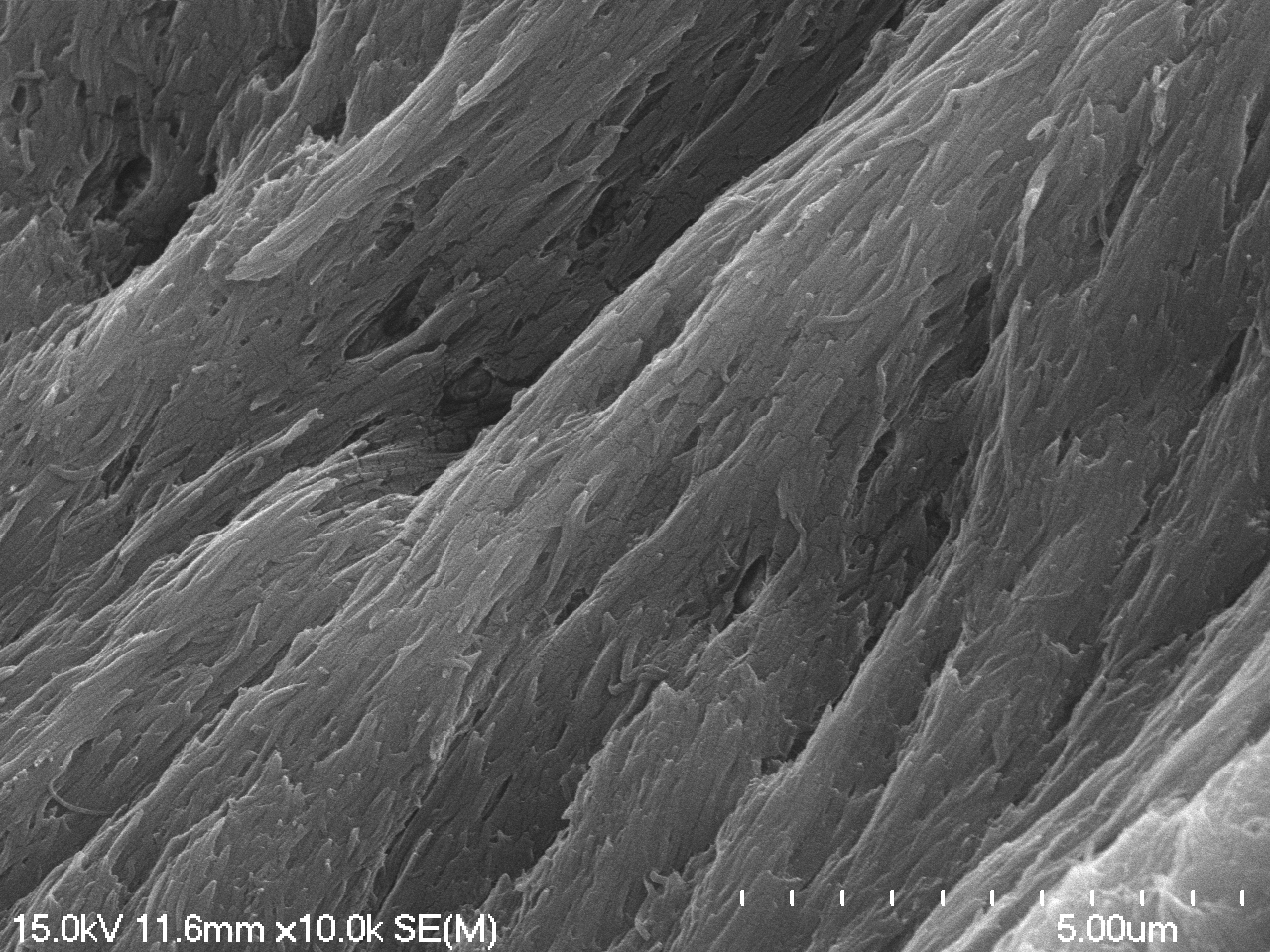

### 05-25-2023 reindeer collagen_m07.tif

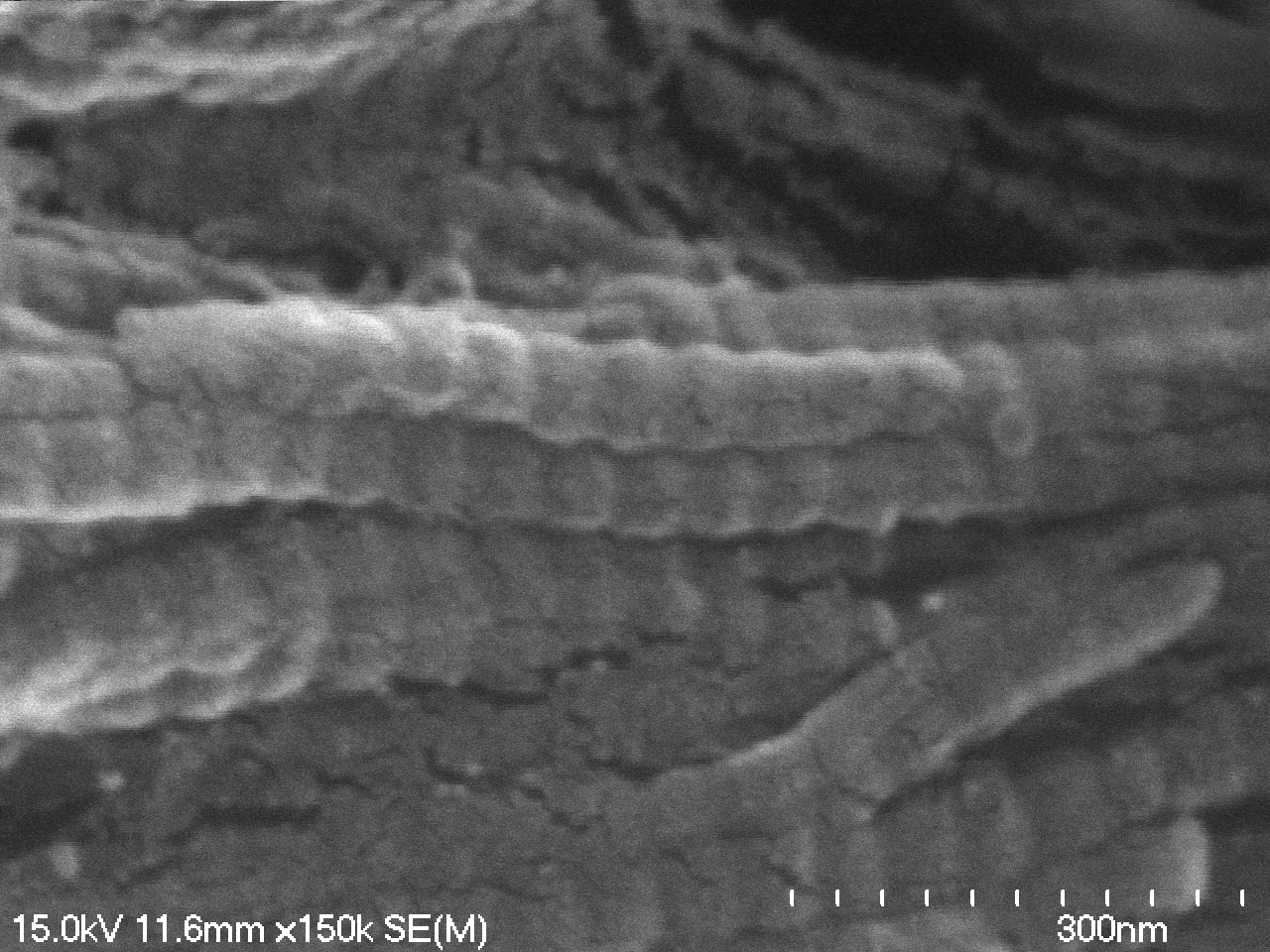

### 05-25-2023 steppe bison metatarsal collagen_m06.tif

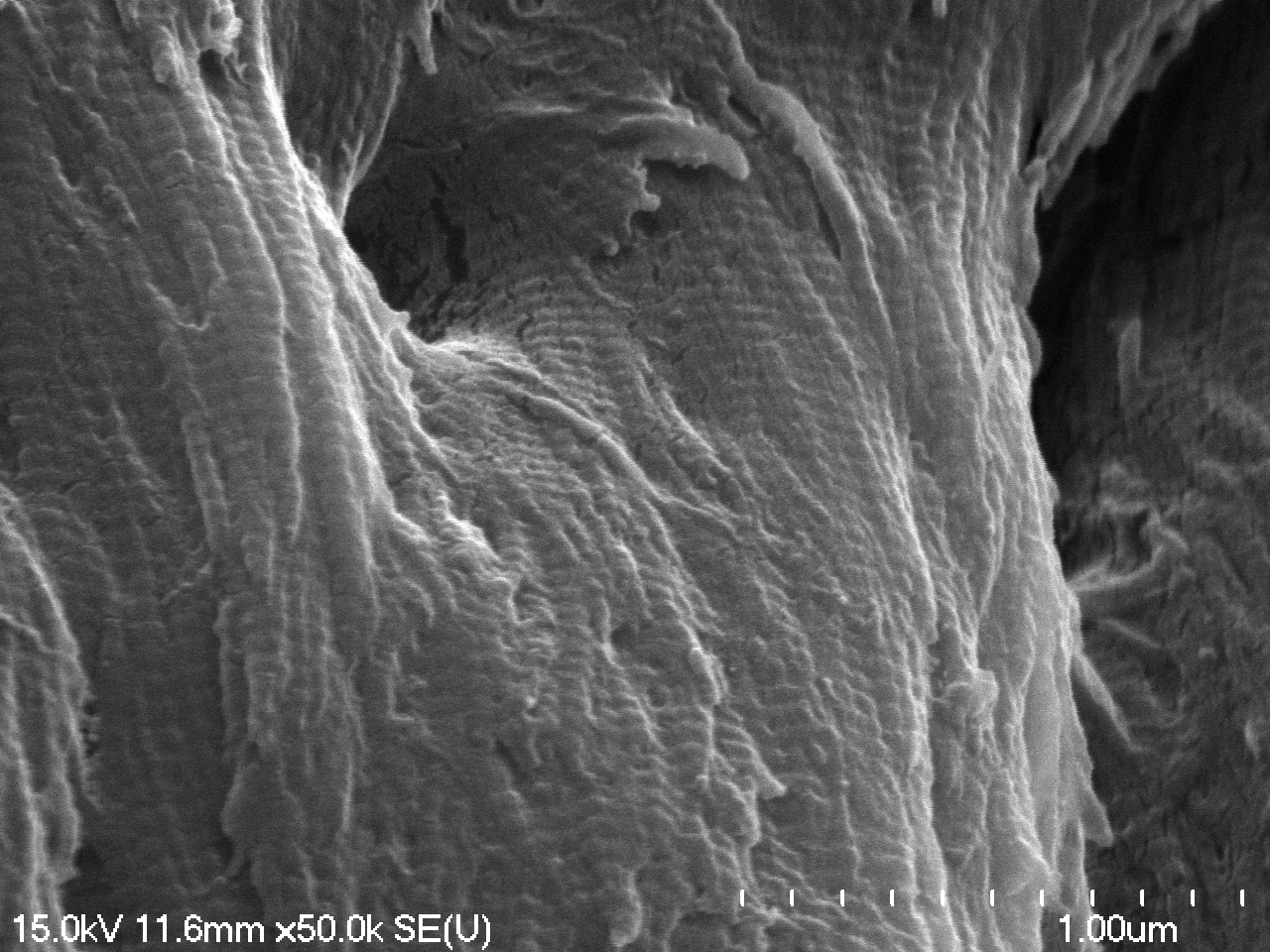

### 05-25-2023 steppe bison metatarsal collagen_m07.tif

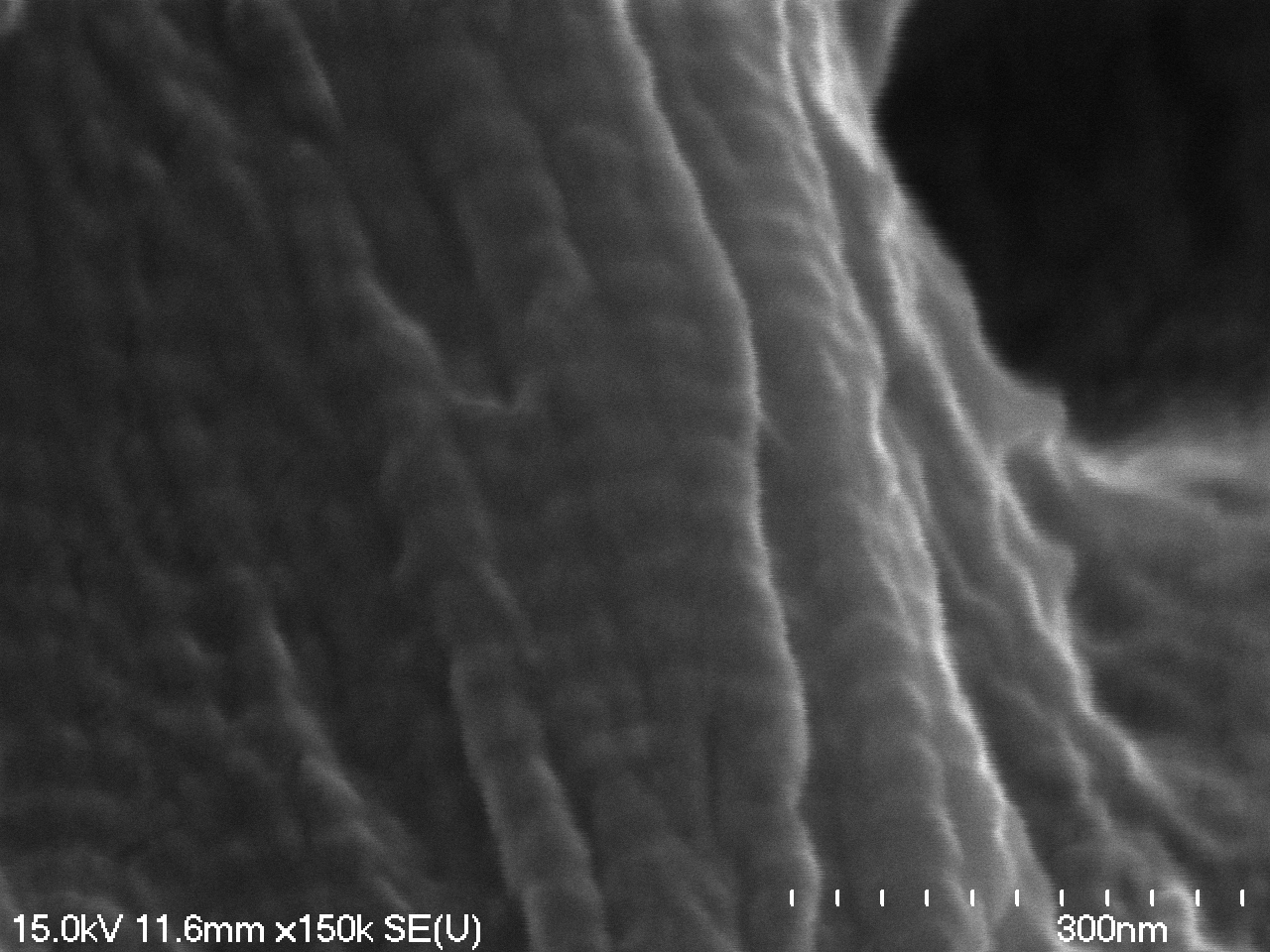

### 05-25-2023 steppe bison metatarsal collagen_m10.tif

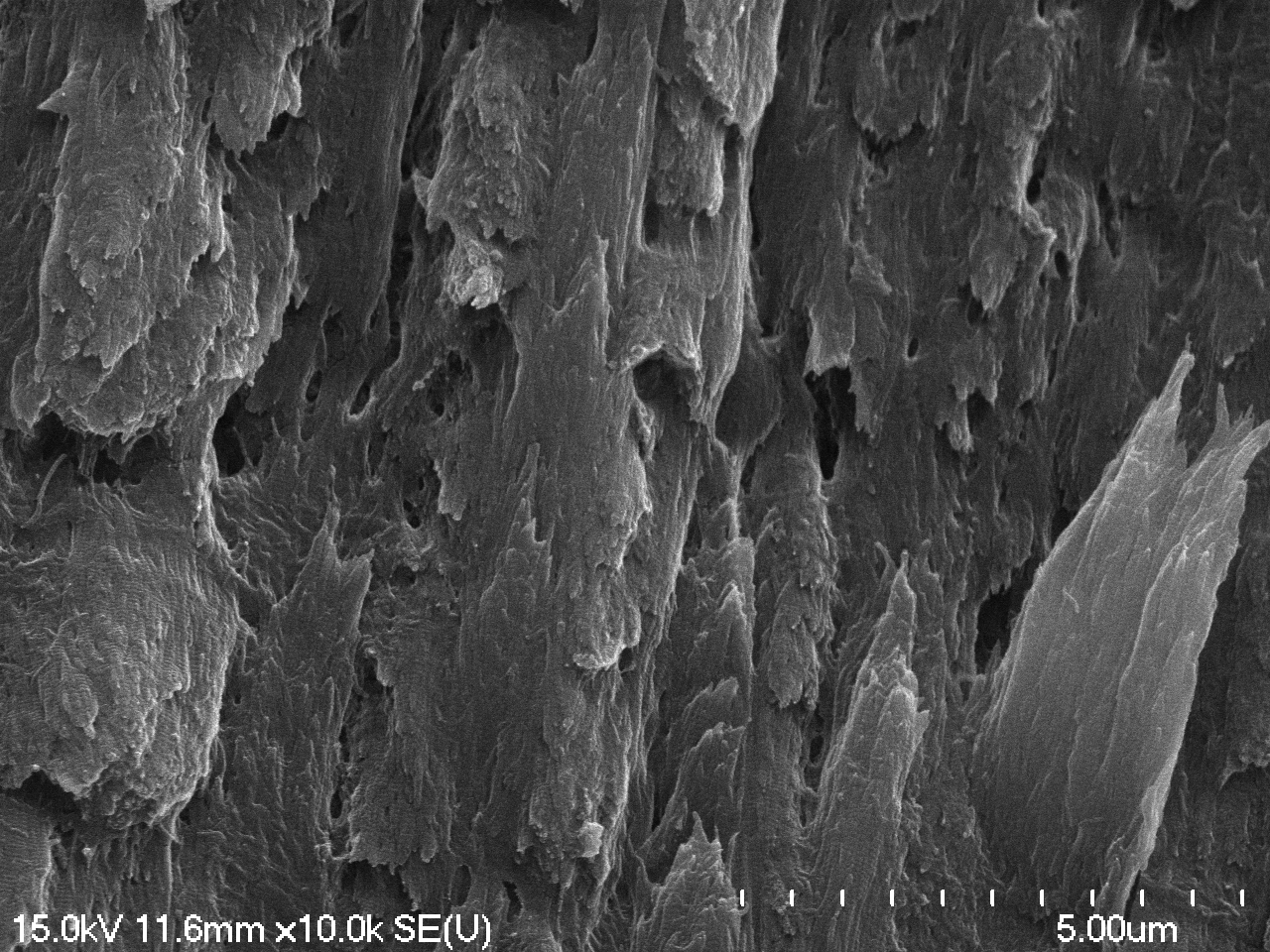

### 05-25-2023 steppe bison radius collagen_m08.tif

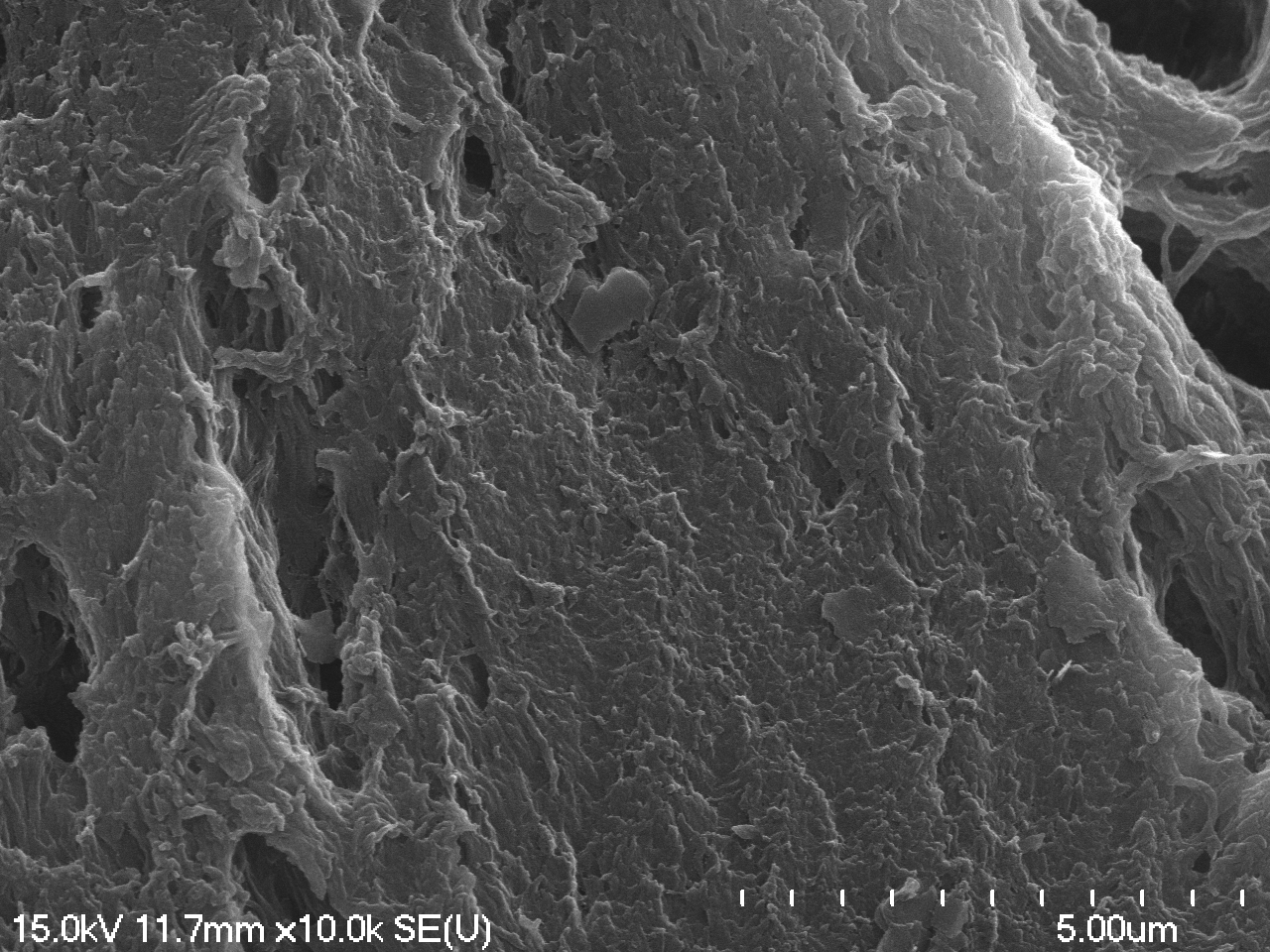

### 06-02-2023 horse metatarsal vessel_m01_m02.tif

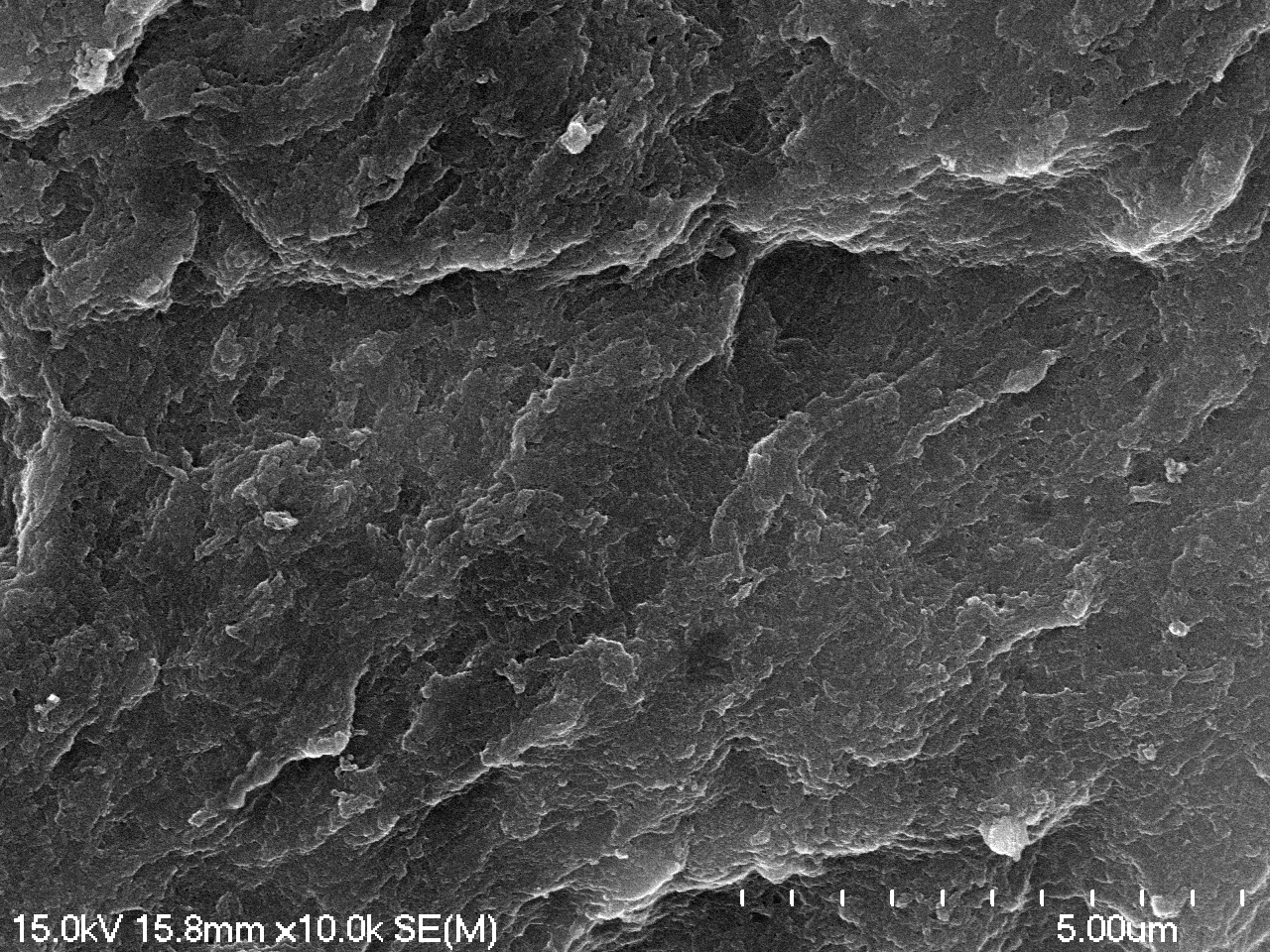

### 06-02-2023 reindeer antler vessel_m02.tif

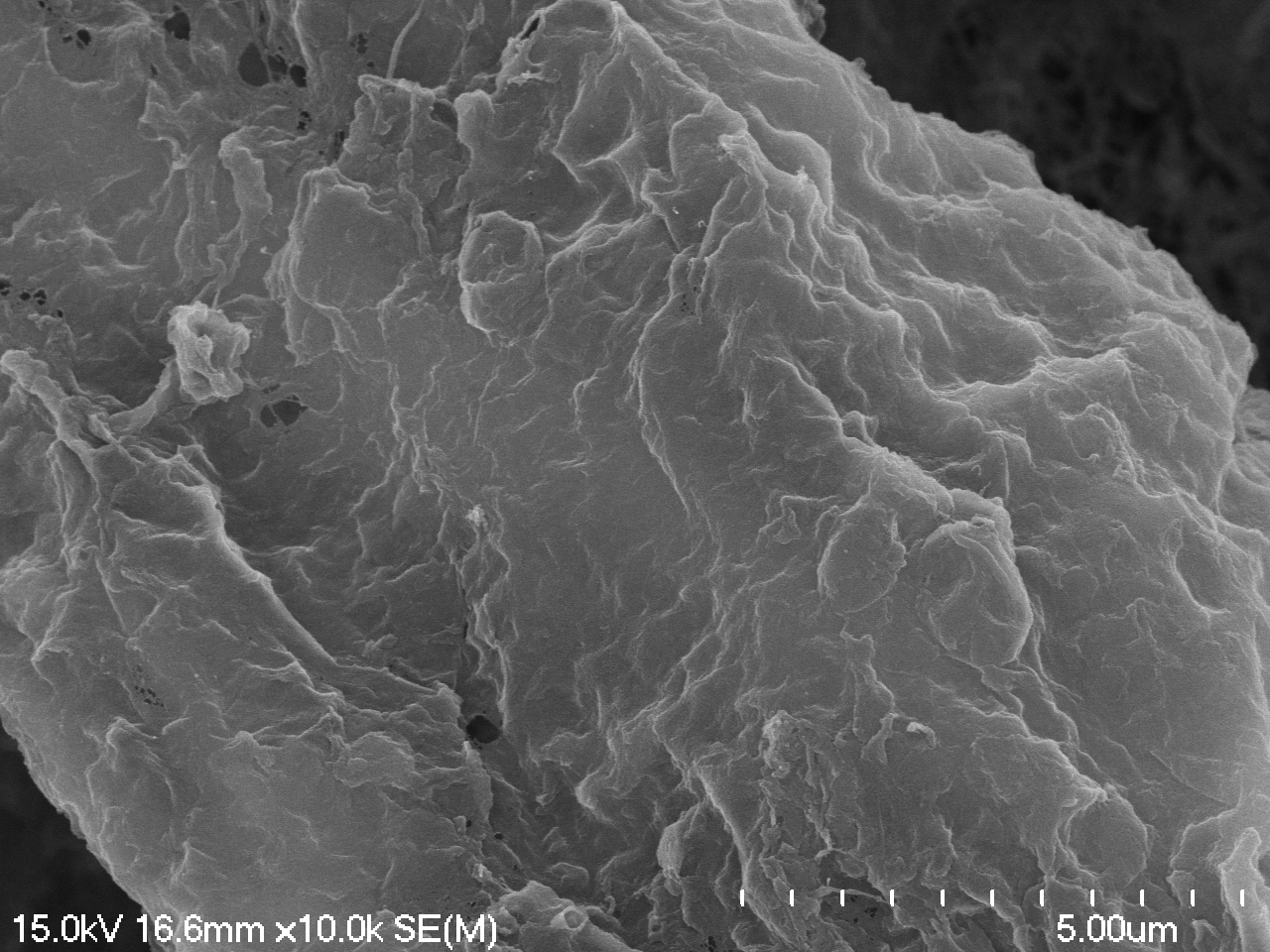

### 06-02-2023 reindeer antler vessel_m05.tif

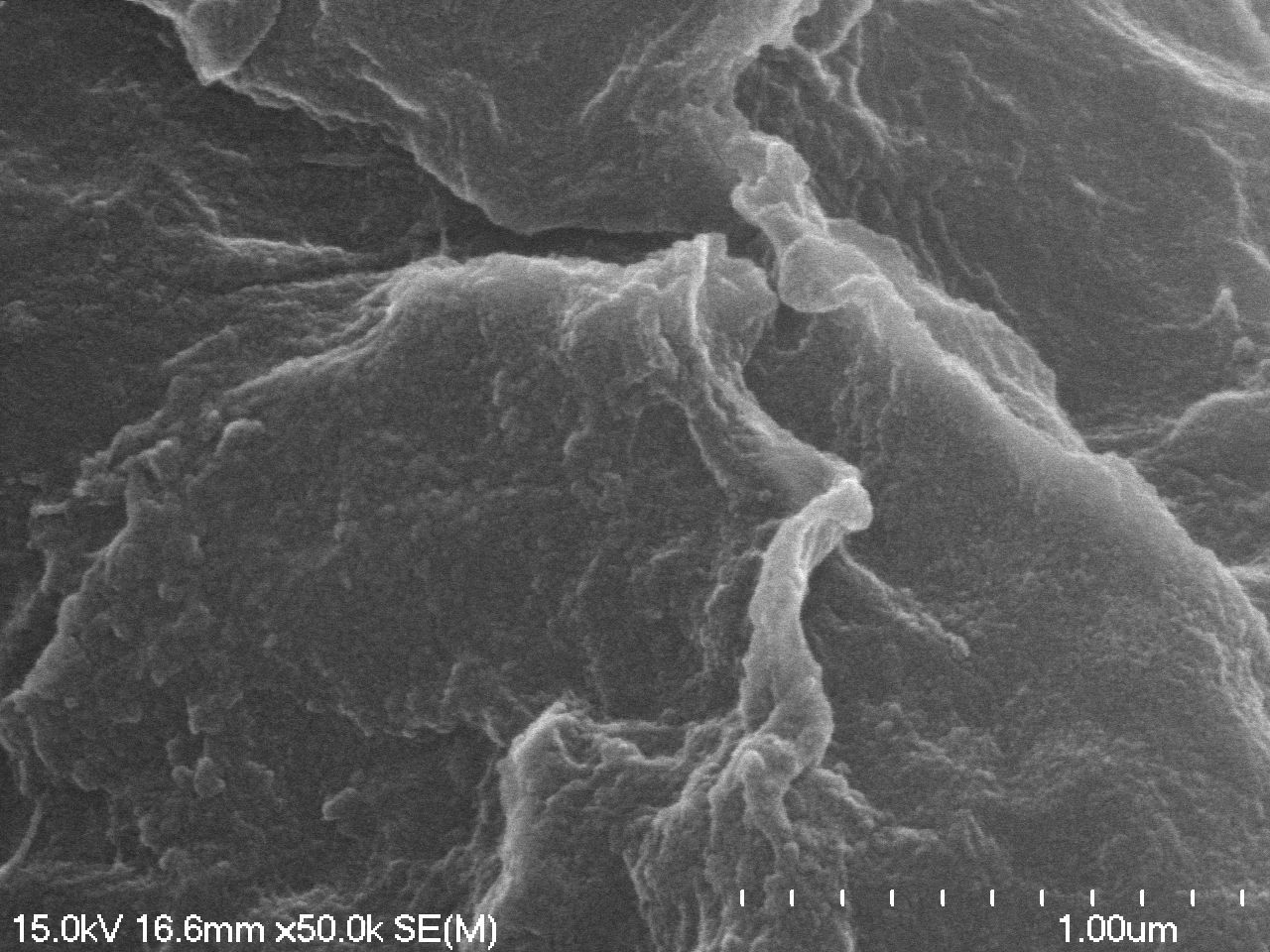

### 06-02-2023 stepp bison metatarsal vessel_m03.tif

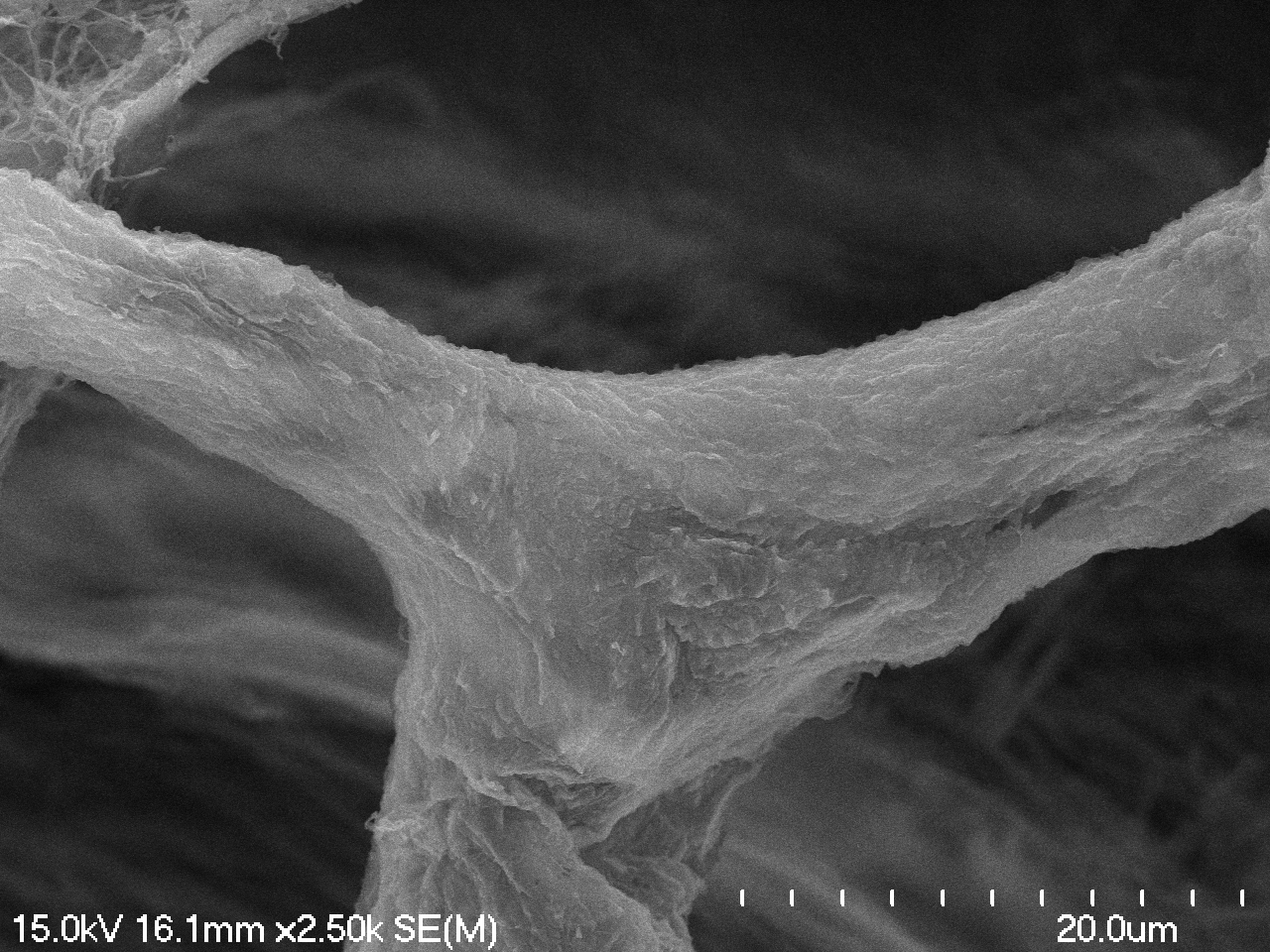

### 06-02-2023 stepp bison metatarsal vessel_m04.tif

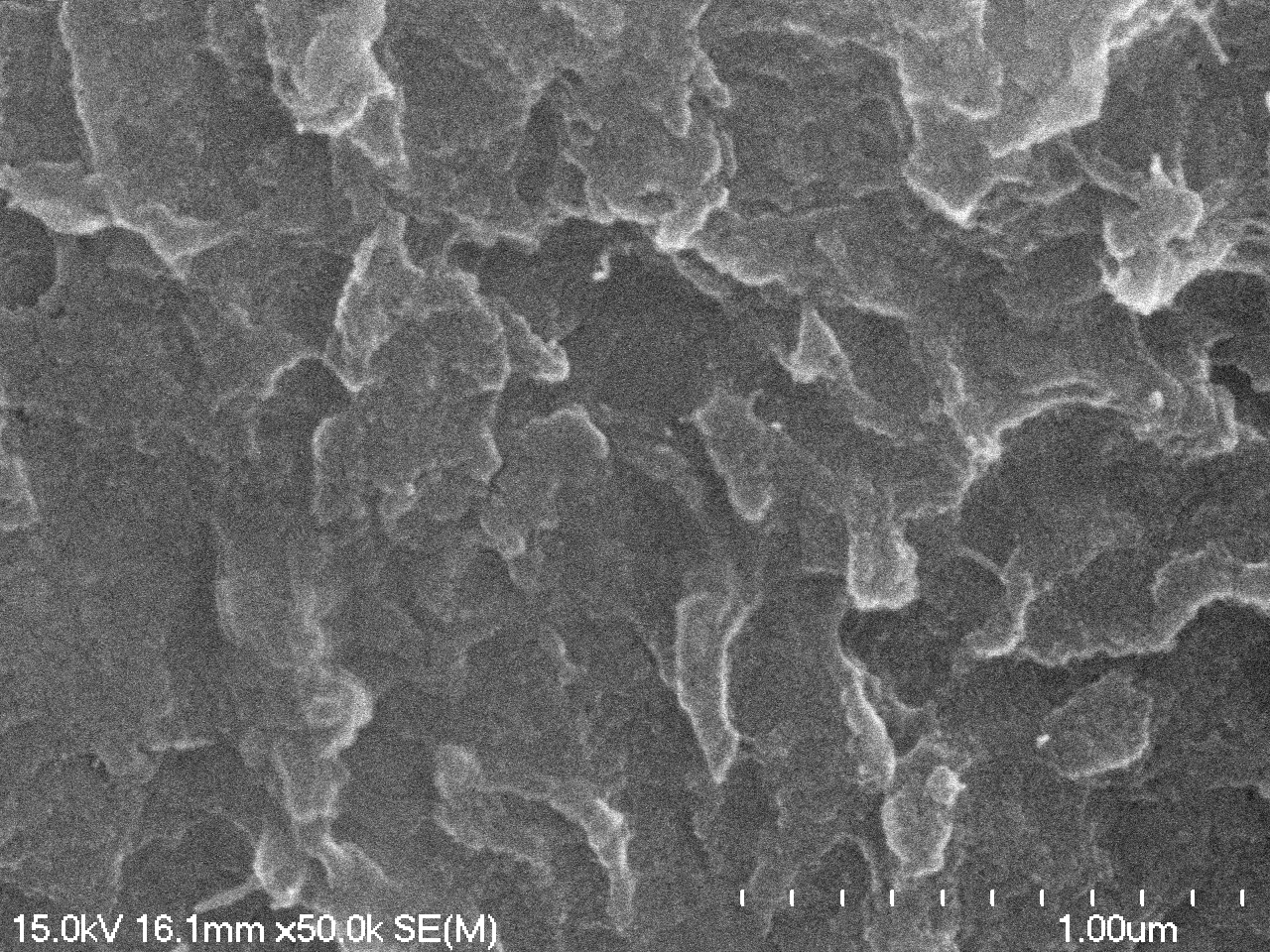

### 06-02-2023 stepp bison metatarsal vessel_m05.tif

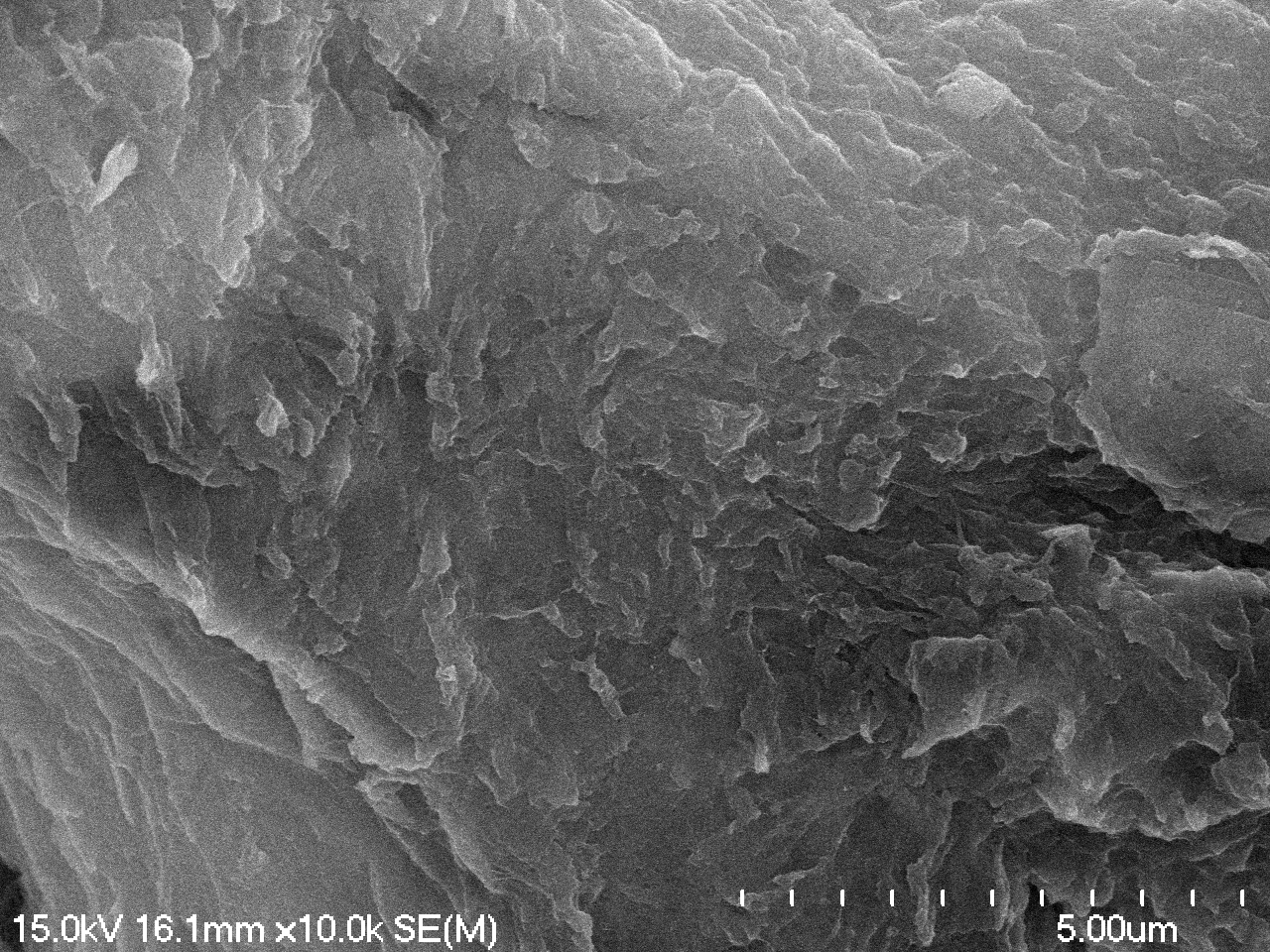

### 06-12-2023 reindeer collagen 400x 01.TIF

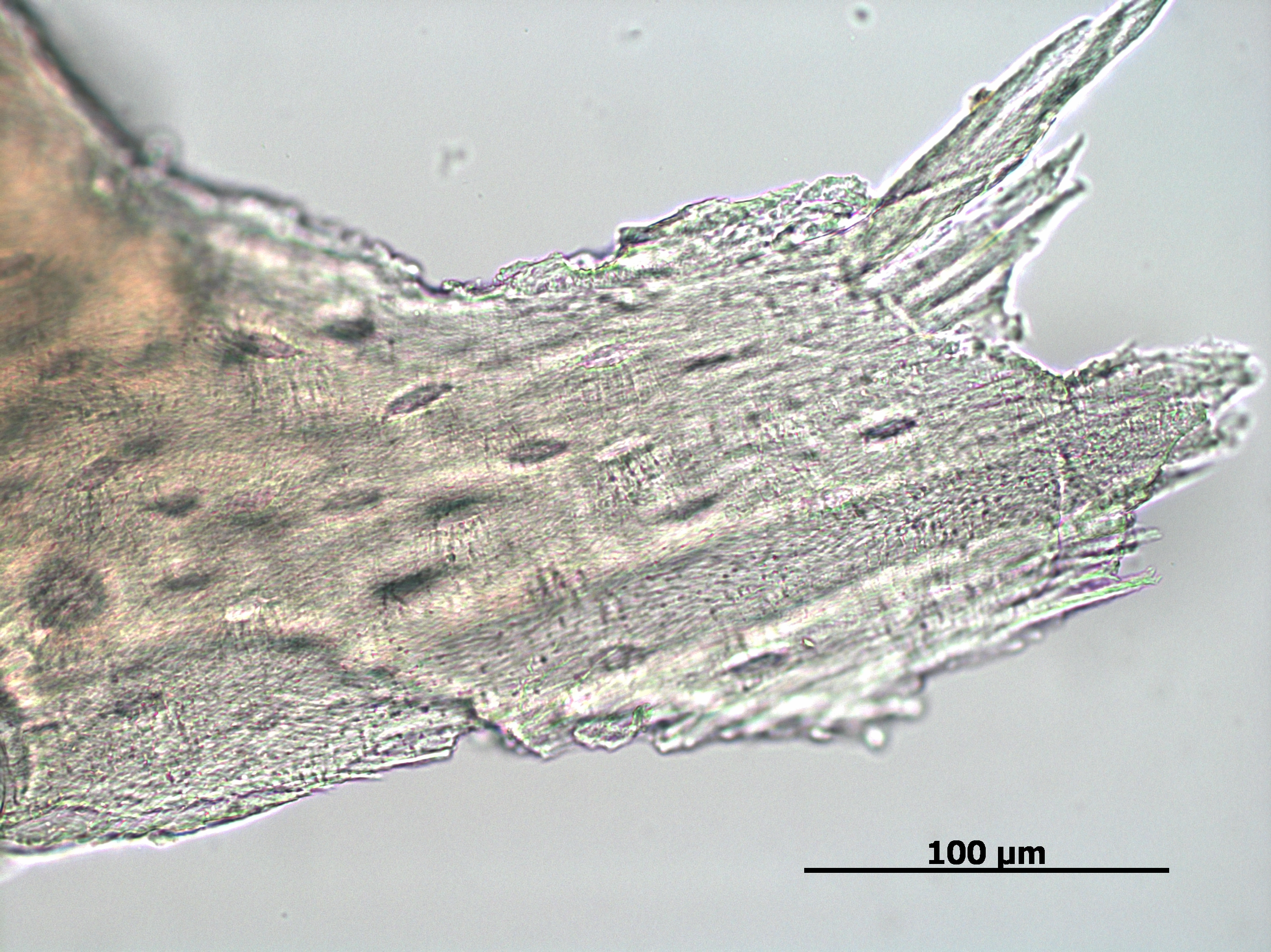

### 06-12-2023 steppe bison meta collagen 400x 02.TIF

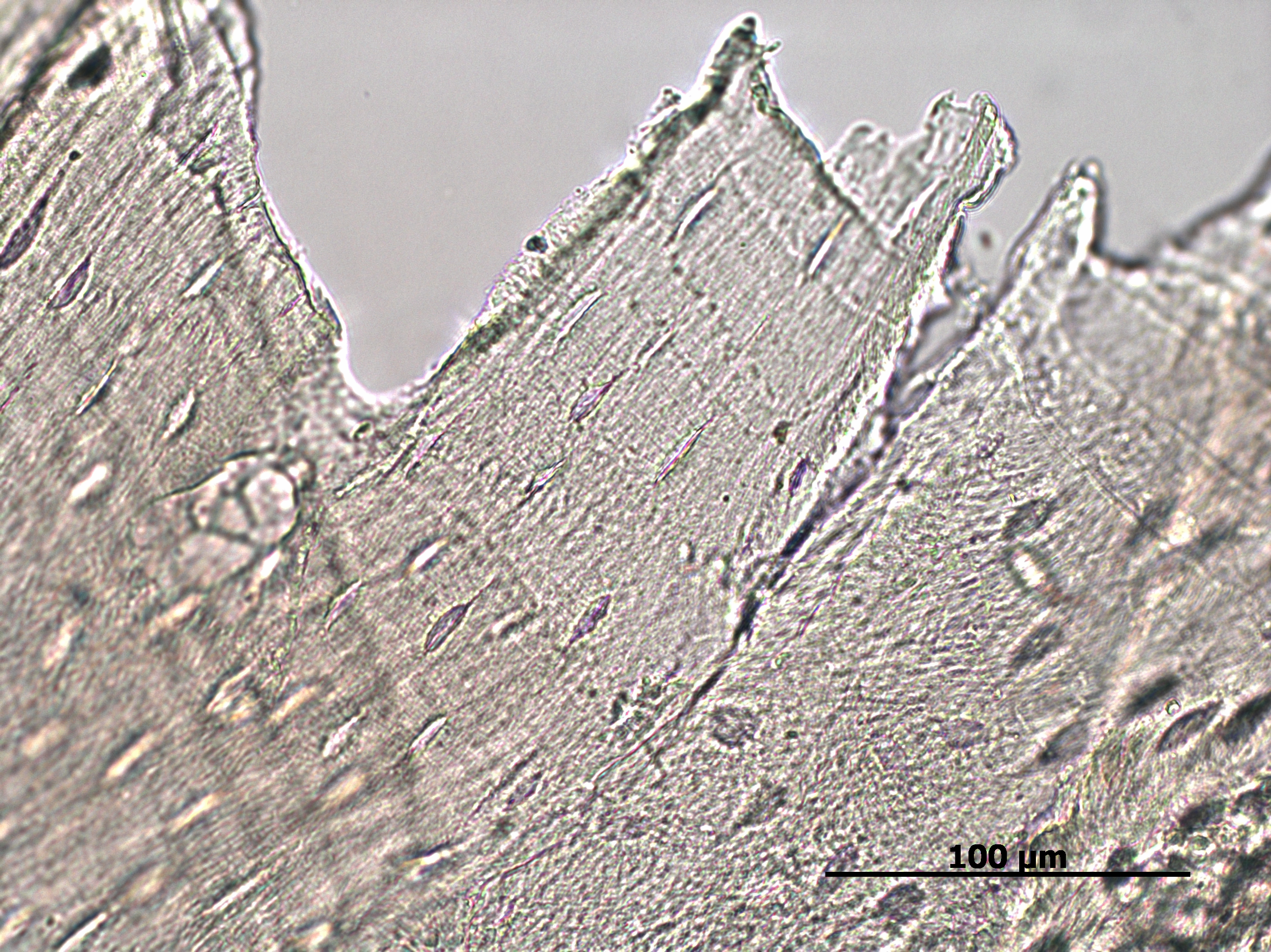

### 06-13-2023 horse collagen 400x 02.TIF

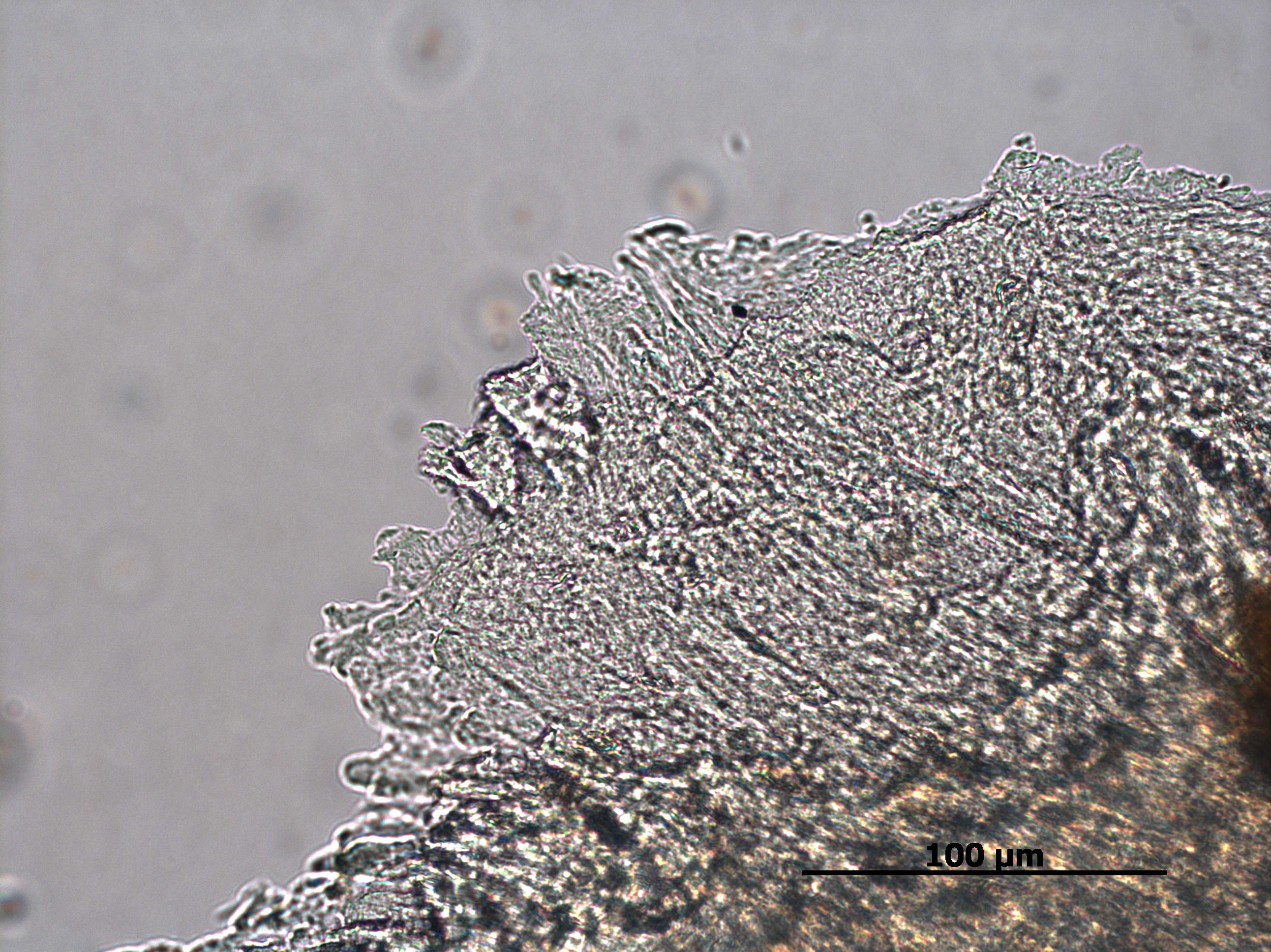

### 06-13-2023 reindeer antler vessel 400x 02.TIF

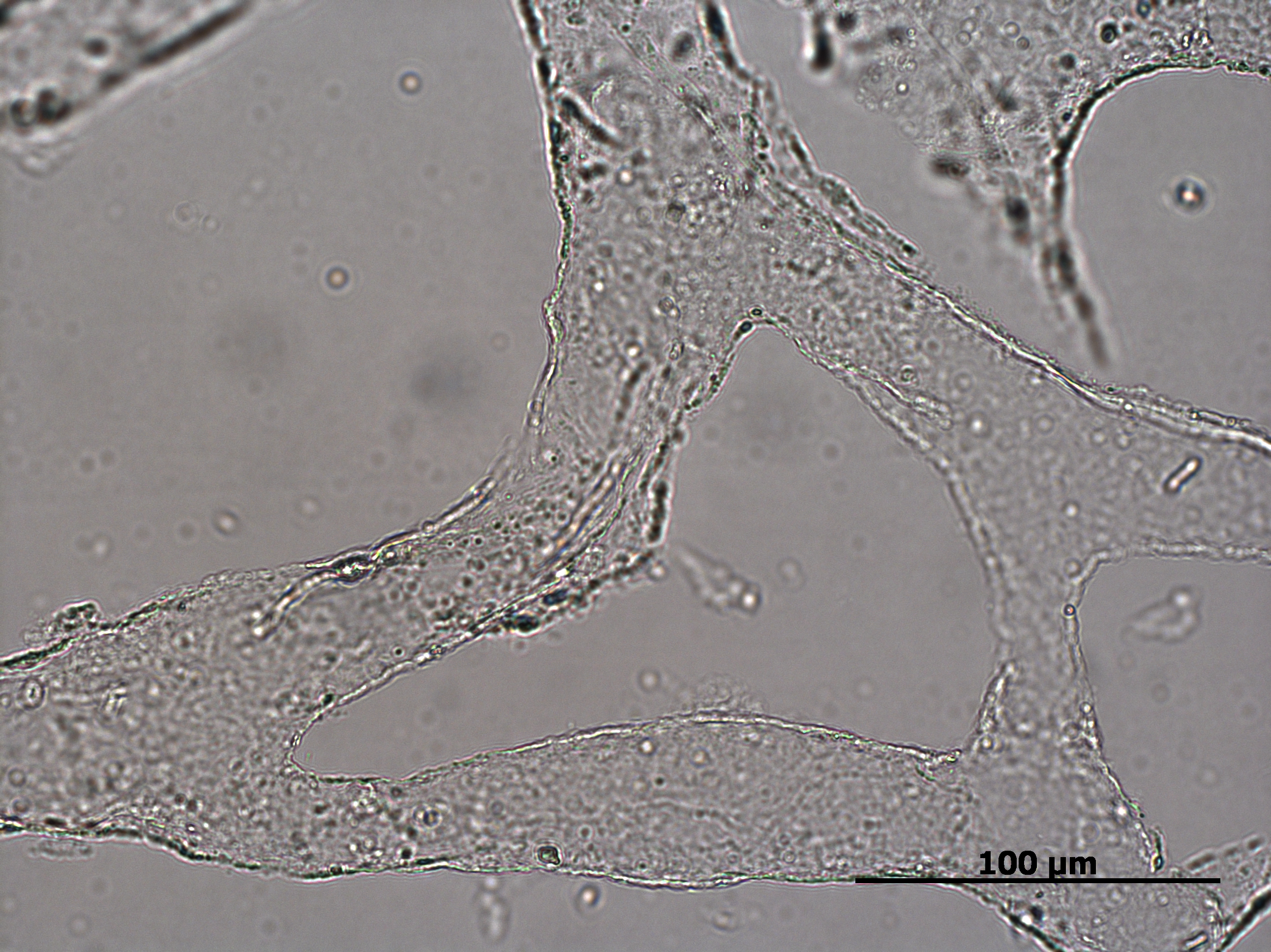

### 06-13-2023 steppe bison meta vessel 400x 02.TIF

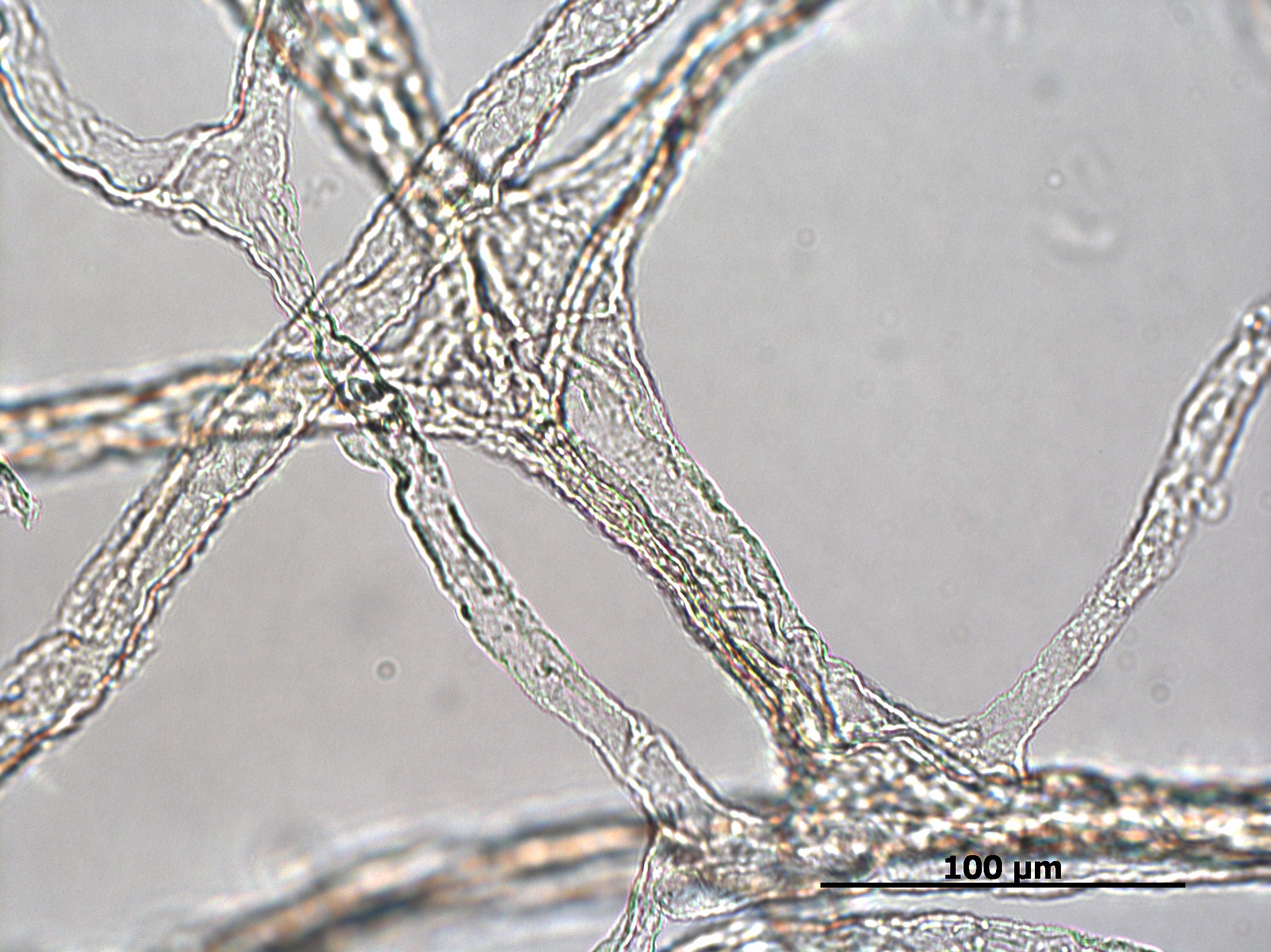

### 06-13-2023 steppe bison radius vessel 400x 01.TIF

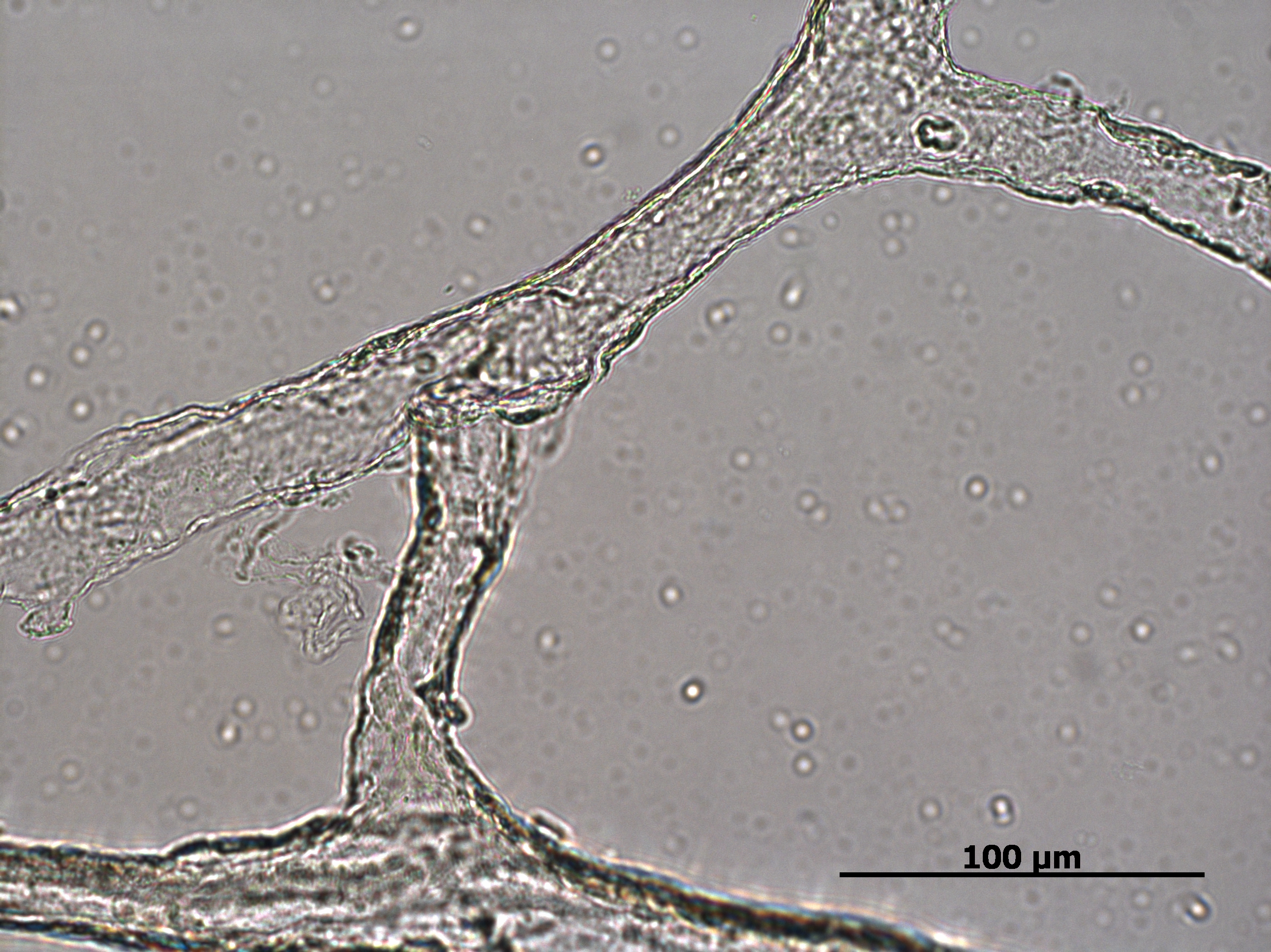

### 06-14-2023 cow humerus collagen 400x 01.TIF

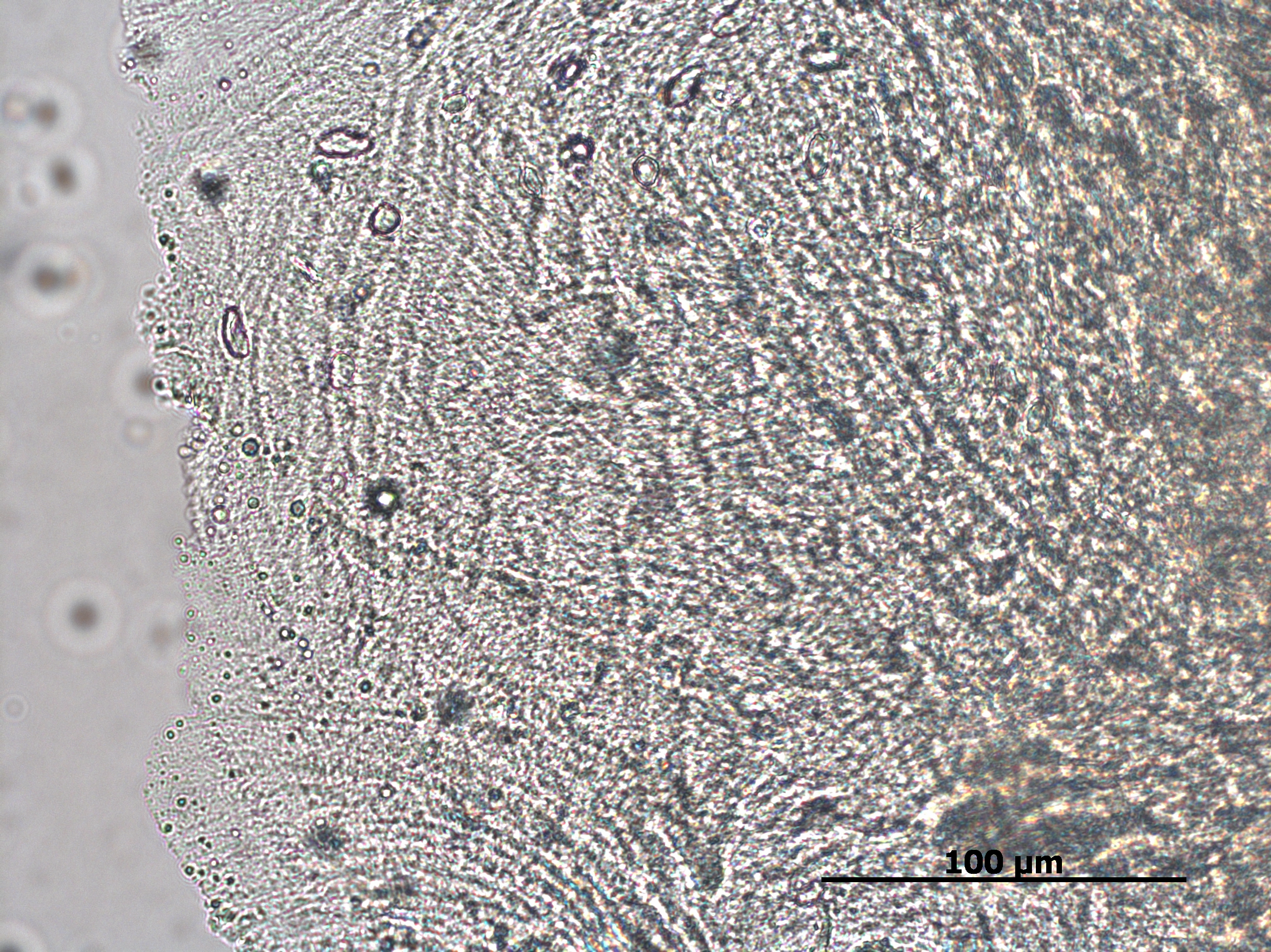

### 06-15-2023 horse meta vessel 400x 01.TIF

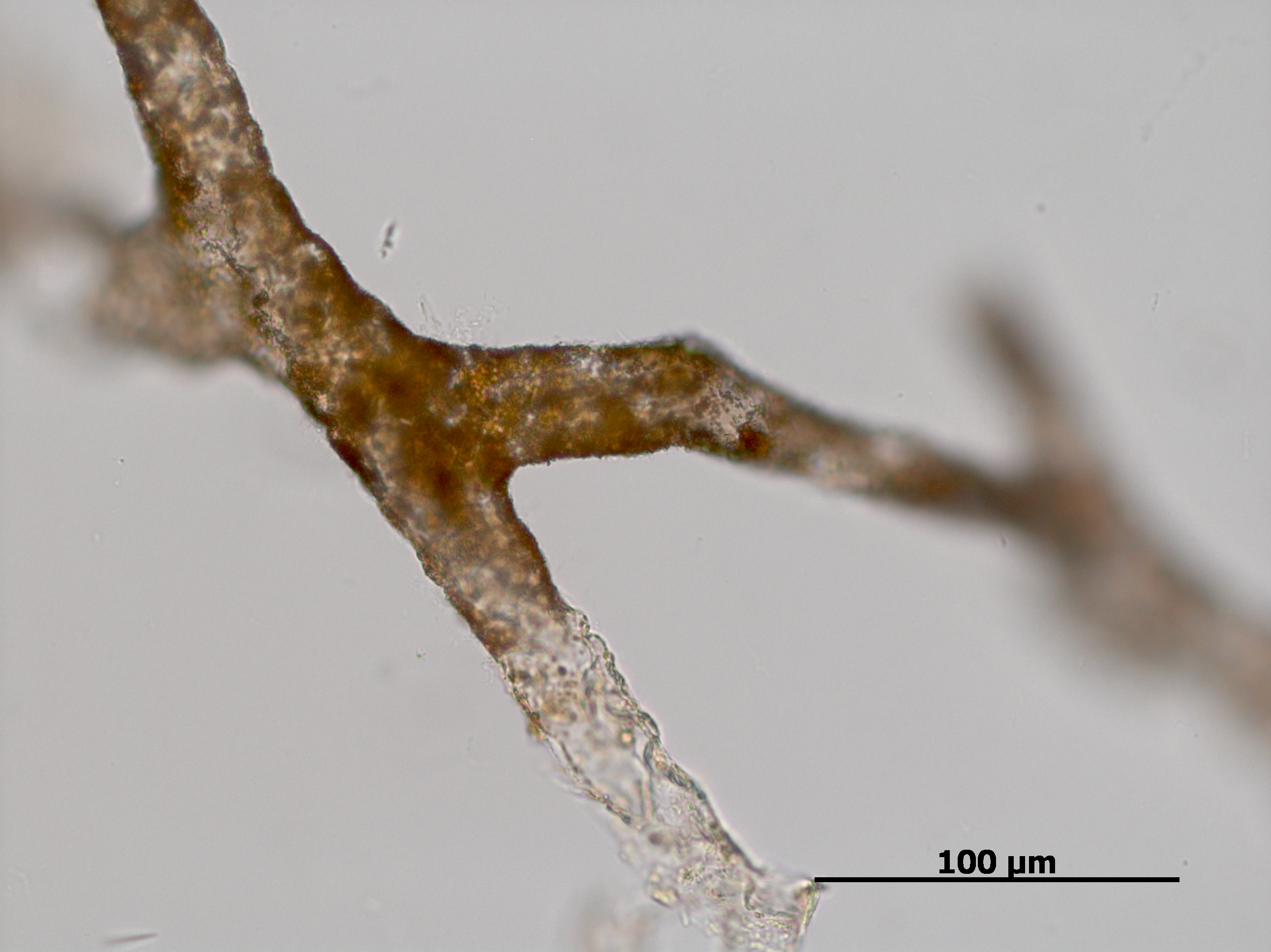

### 06-15-2023 mammoth innominate vessel 400x 01.TIF

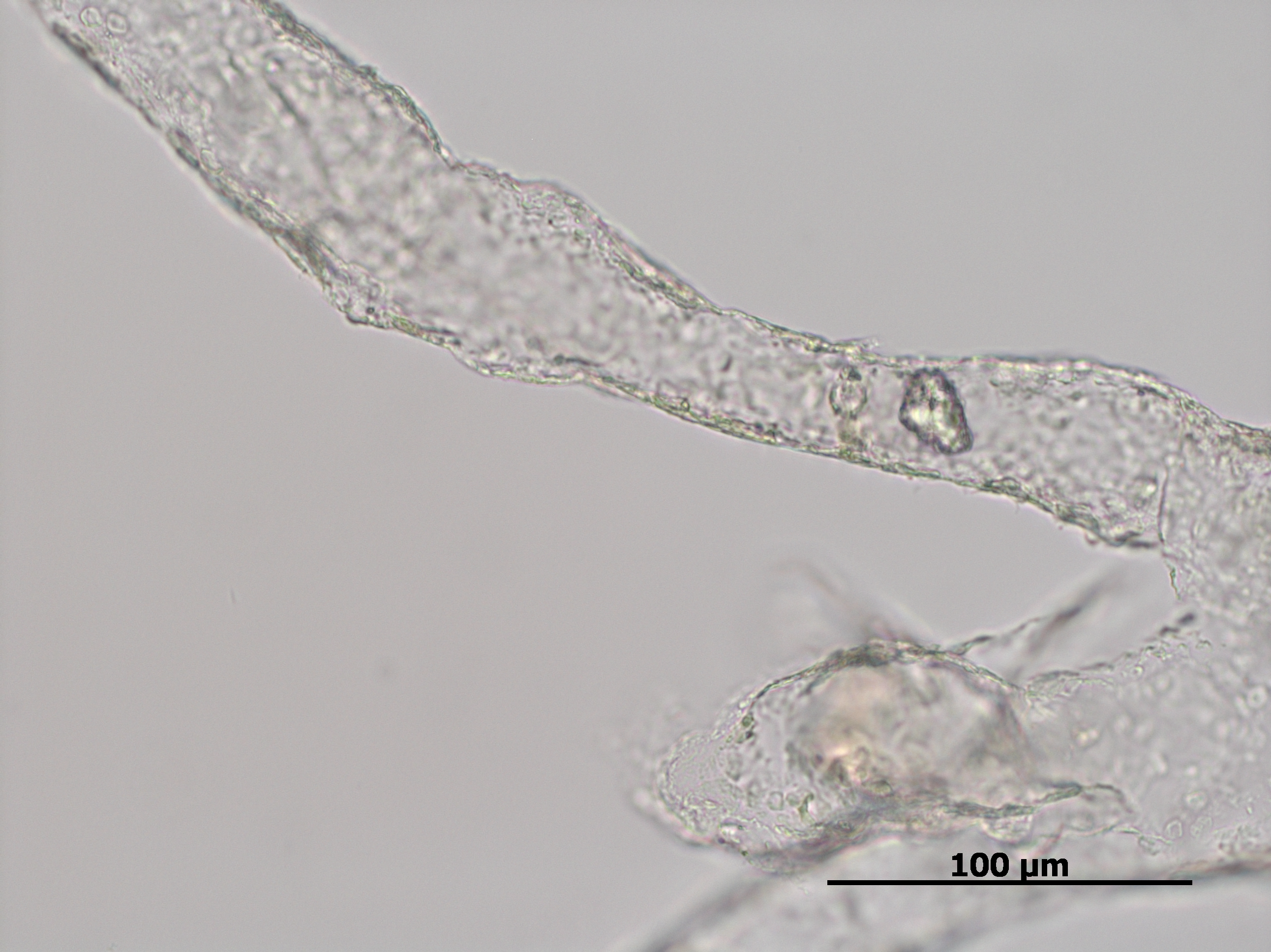

### 06-15-2023 reindeer antler vessels_m07.tif

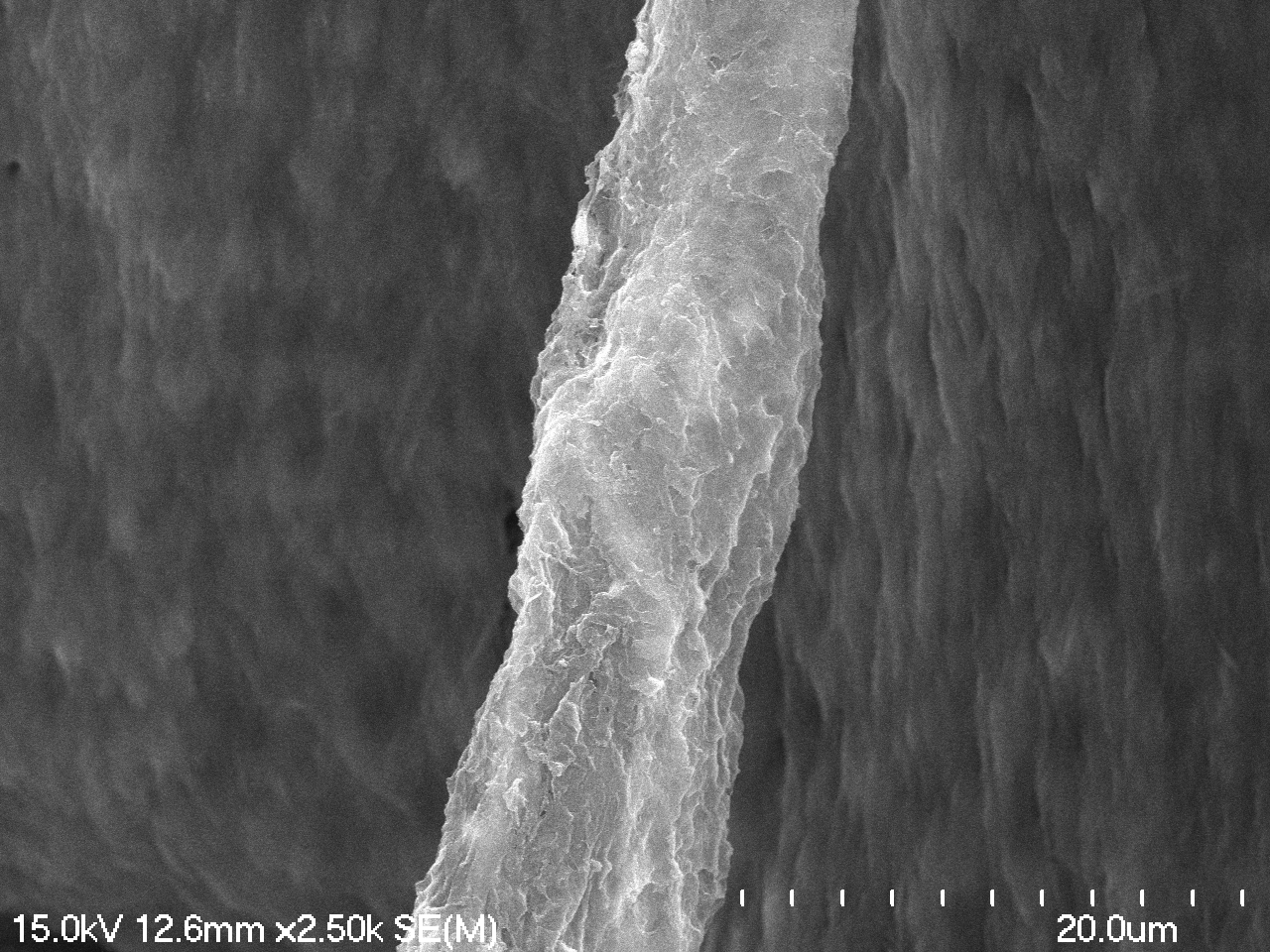

### 06-17-2023 horse meta vessel_m07.tif

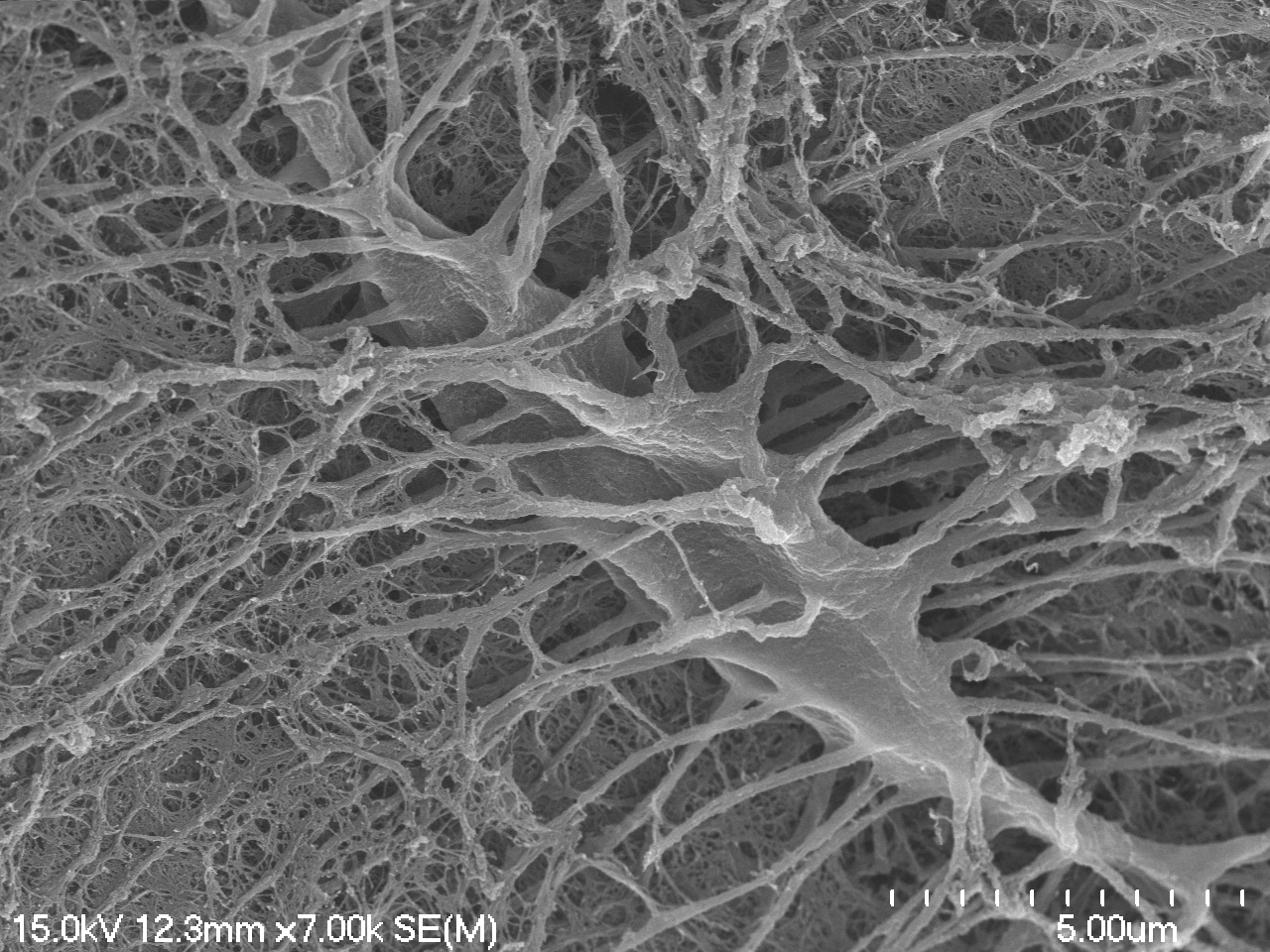

### 06-17-2023 horse meta vessel_m09.tif

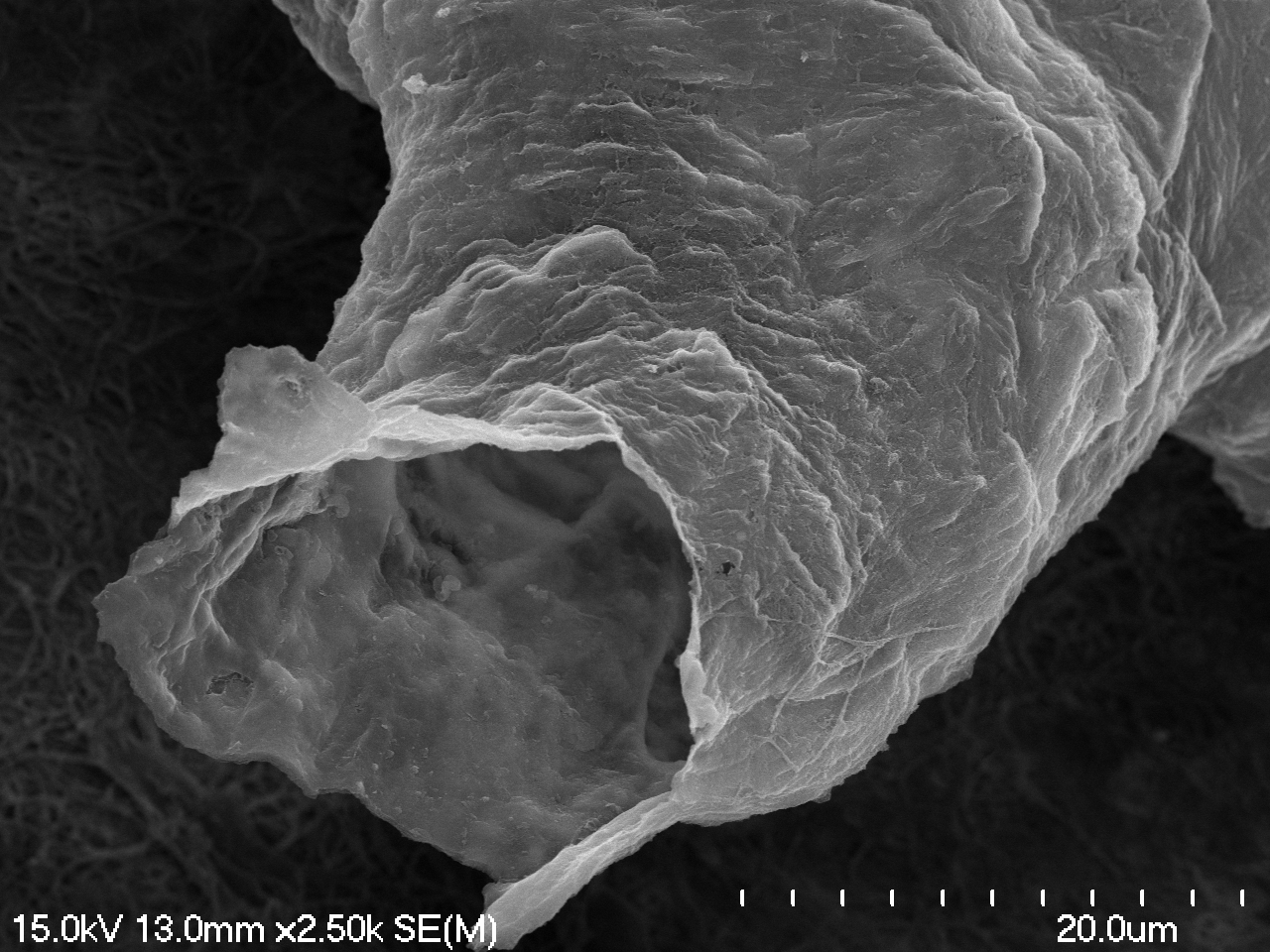

### 06-17-2023 horse meta vessel_m11.tif

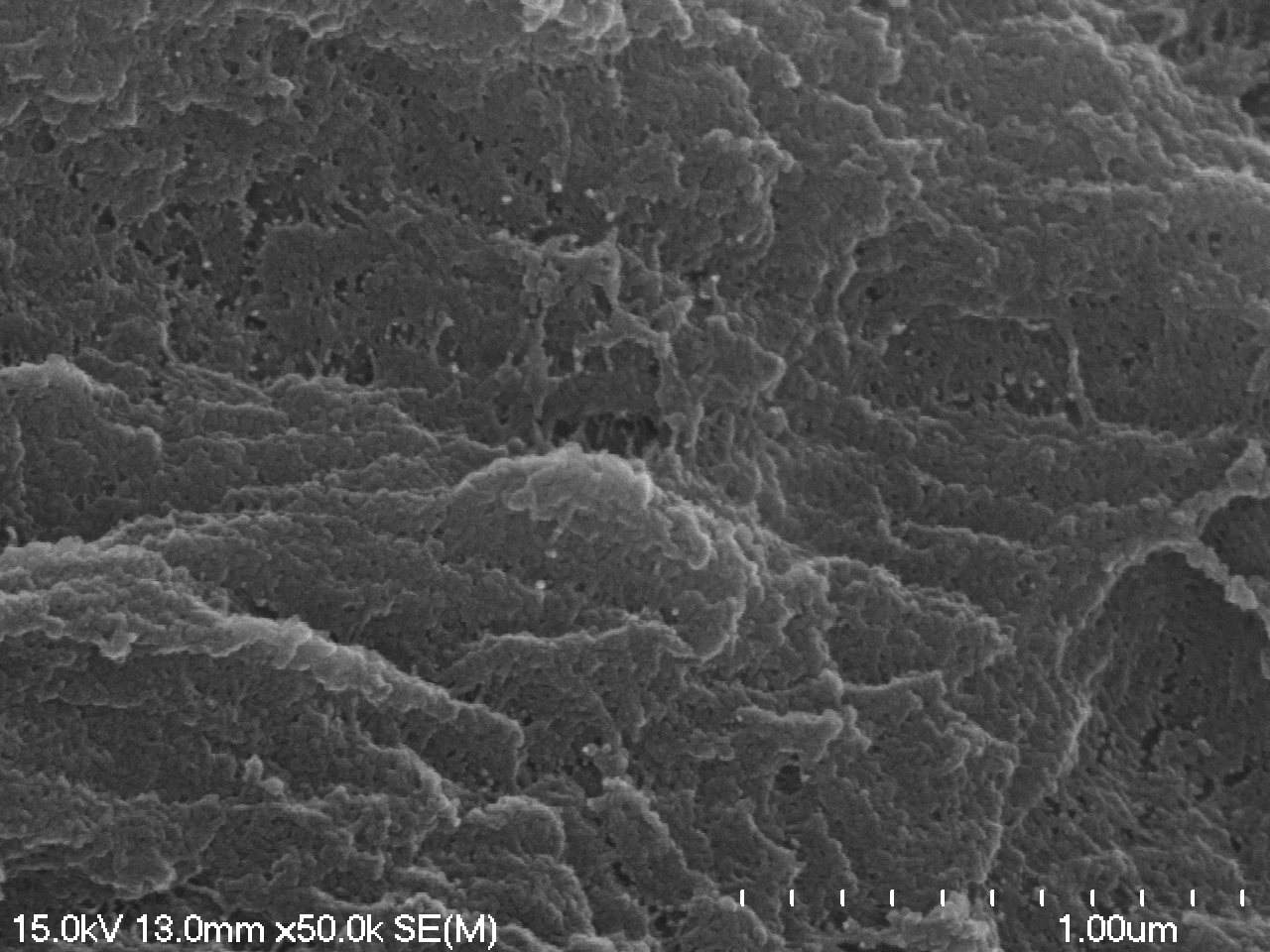

### 06-17-2023 steppe bison radius vessel_m01.tif

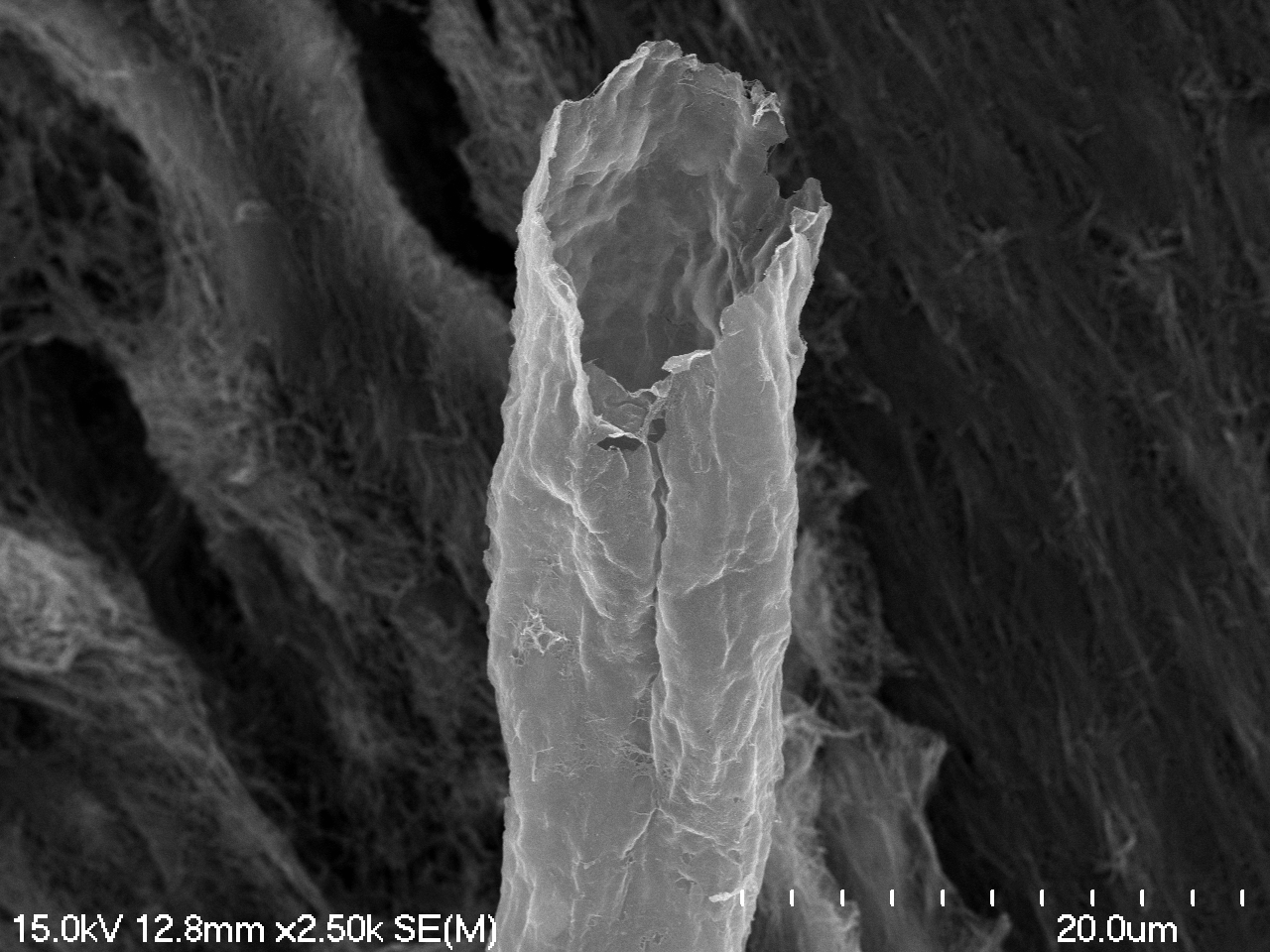

### 06-17-2023 steppe bison radius vessel_m02.tif

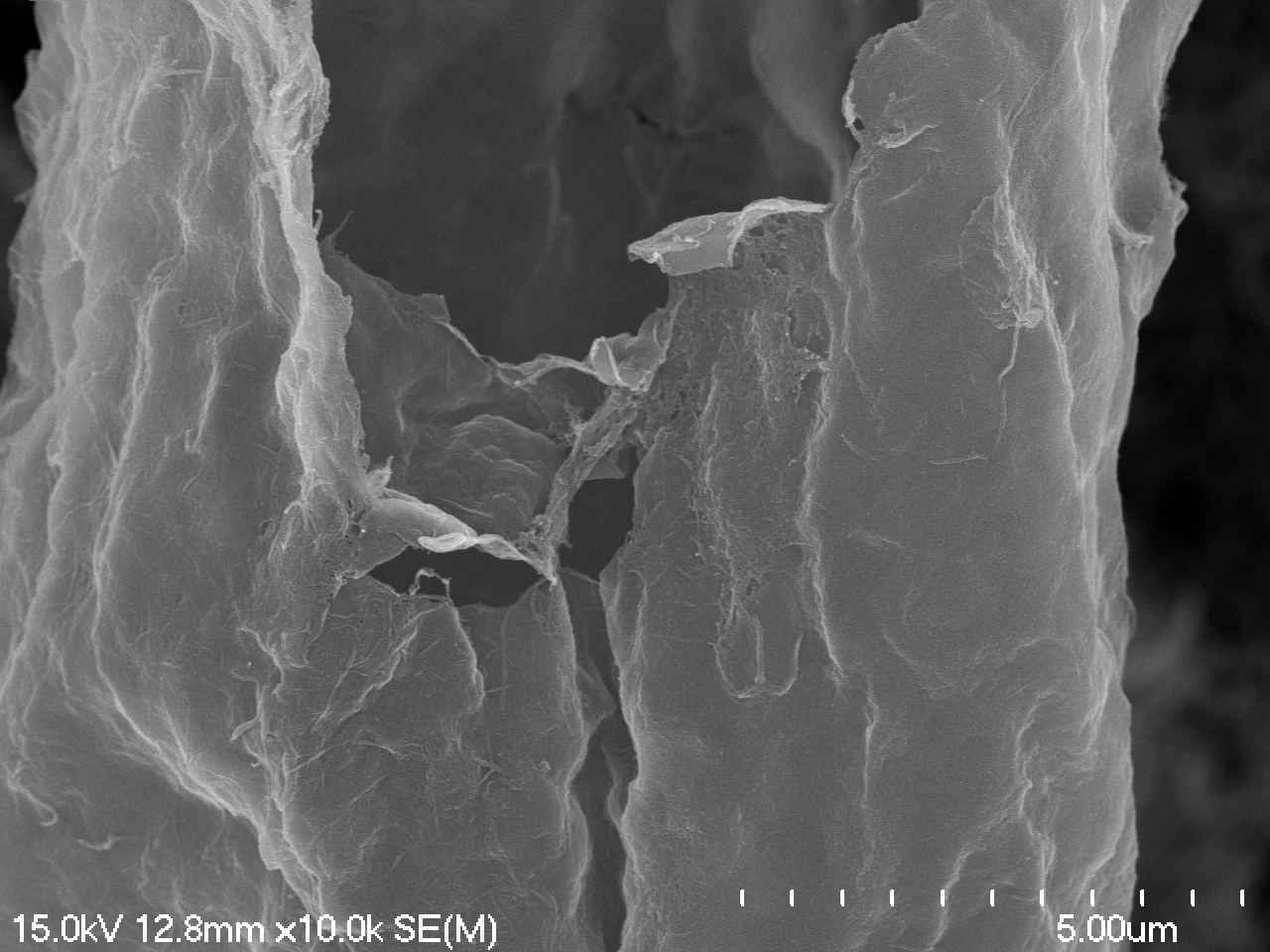

### 06-17-2023 steppe bison radius vessel_m03.tif

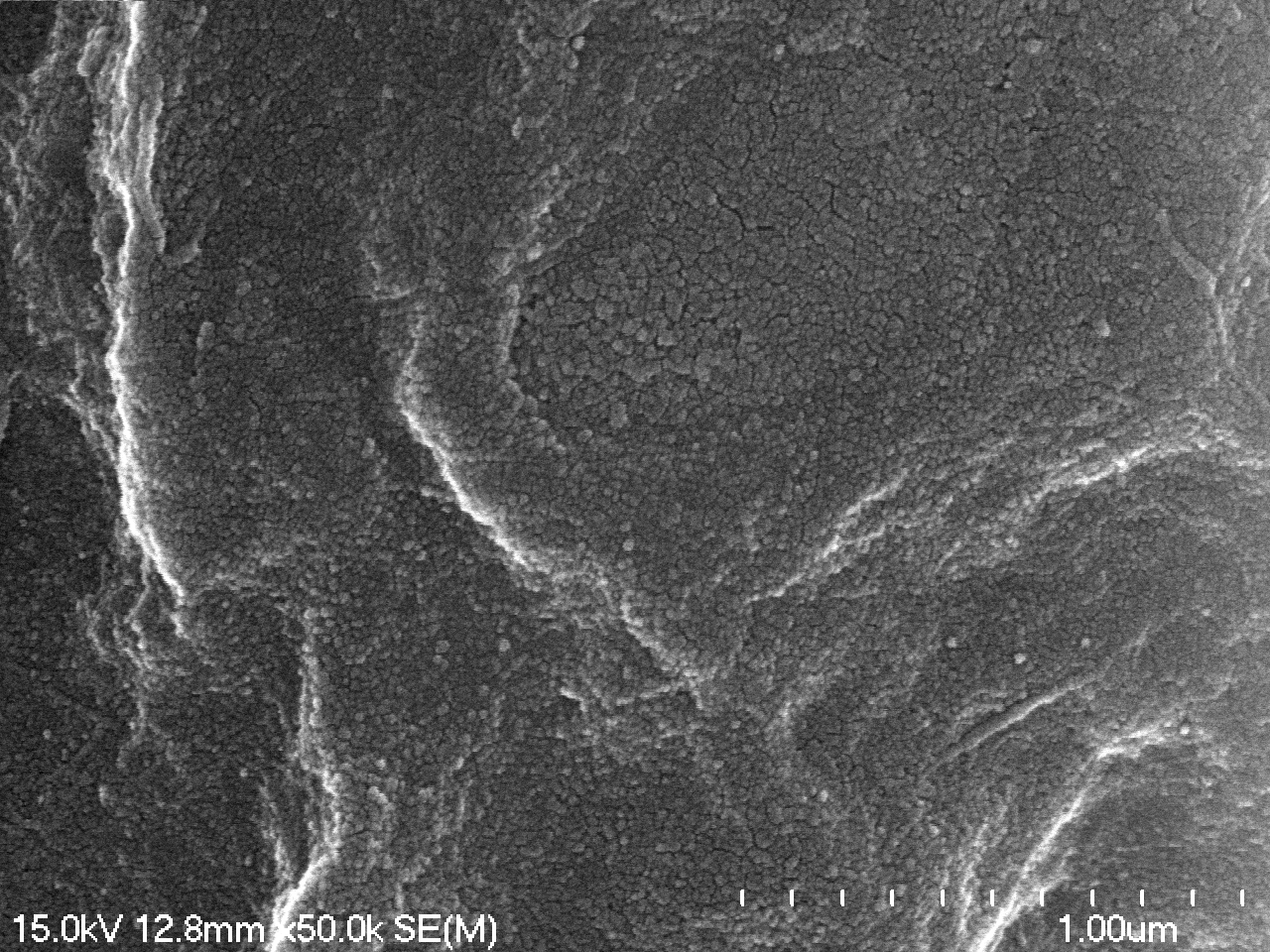

### 06-29-2023 cow humerus blood vessels_m01_m02.tif

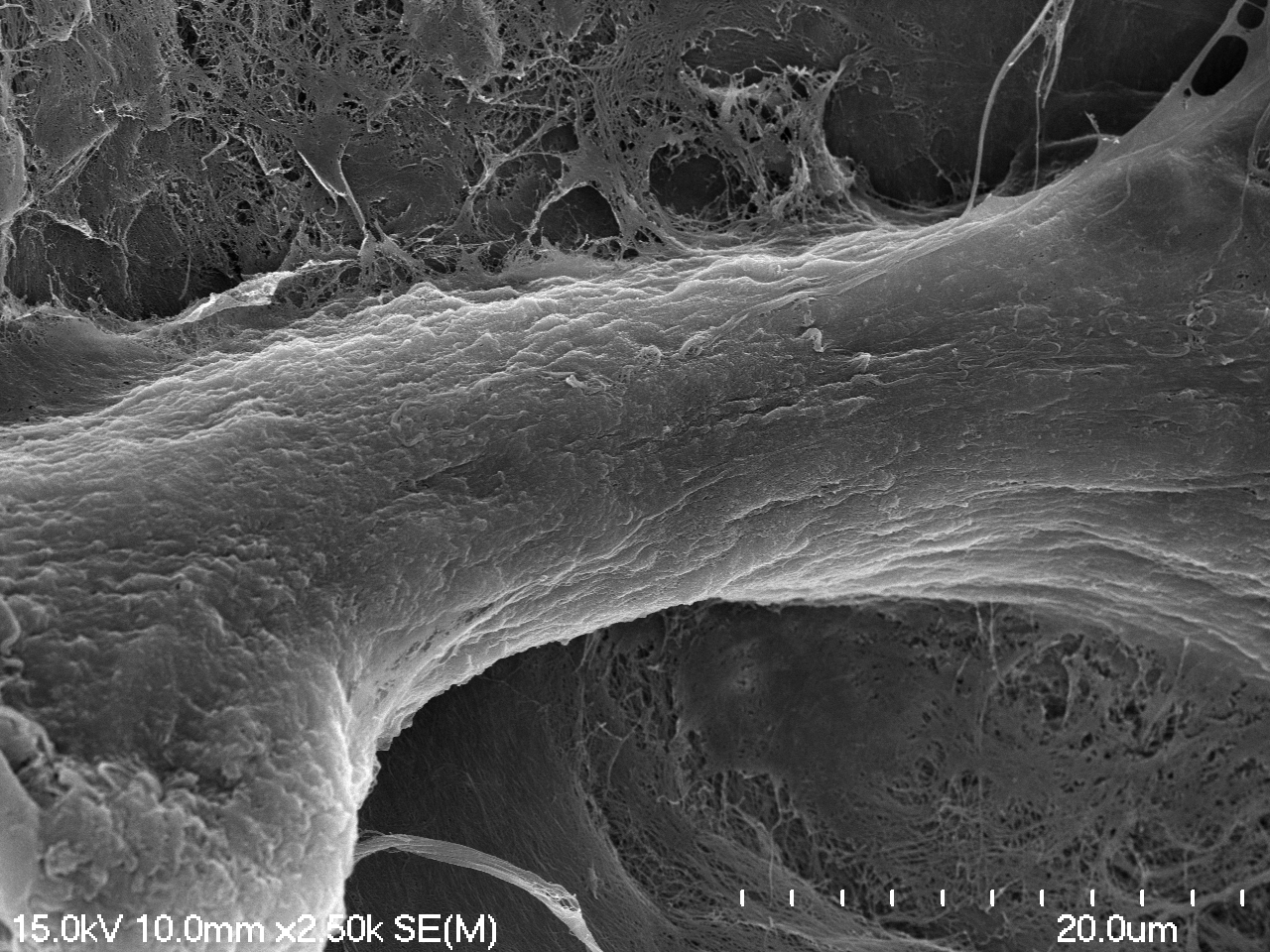

### 06-29-2023 cow humerus blood vessels_m11_m07.tif

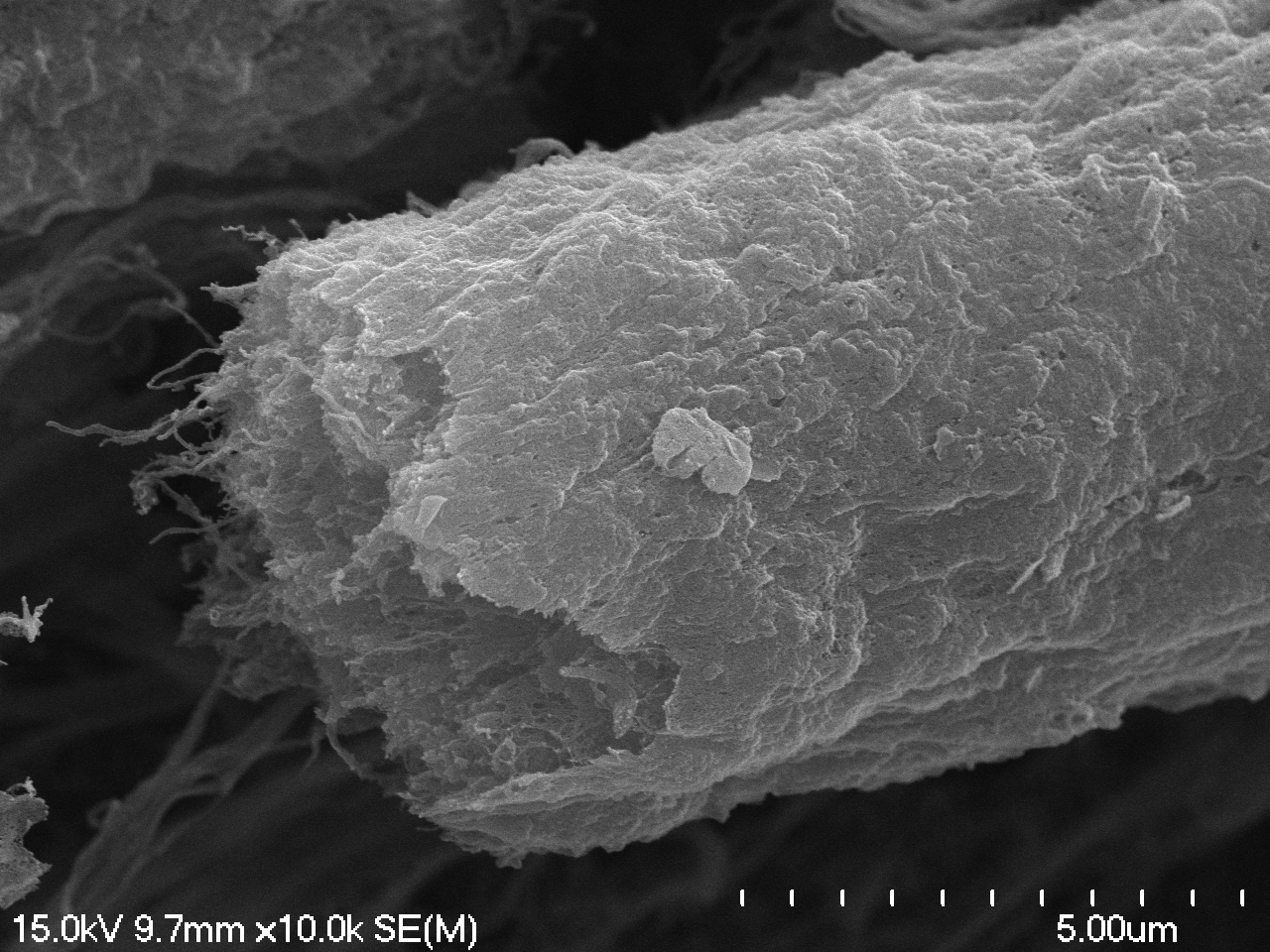

### 06-29-2023 cow humerus blood vessels_m11_m09.tif

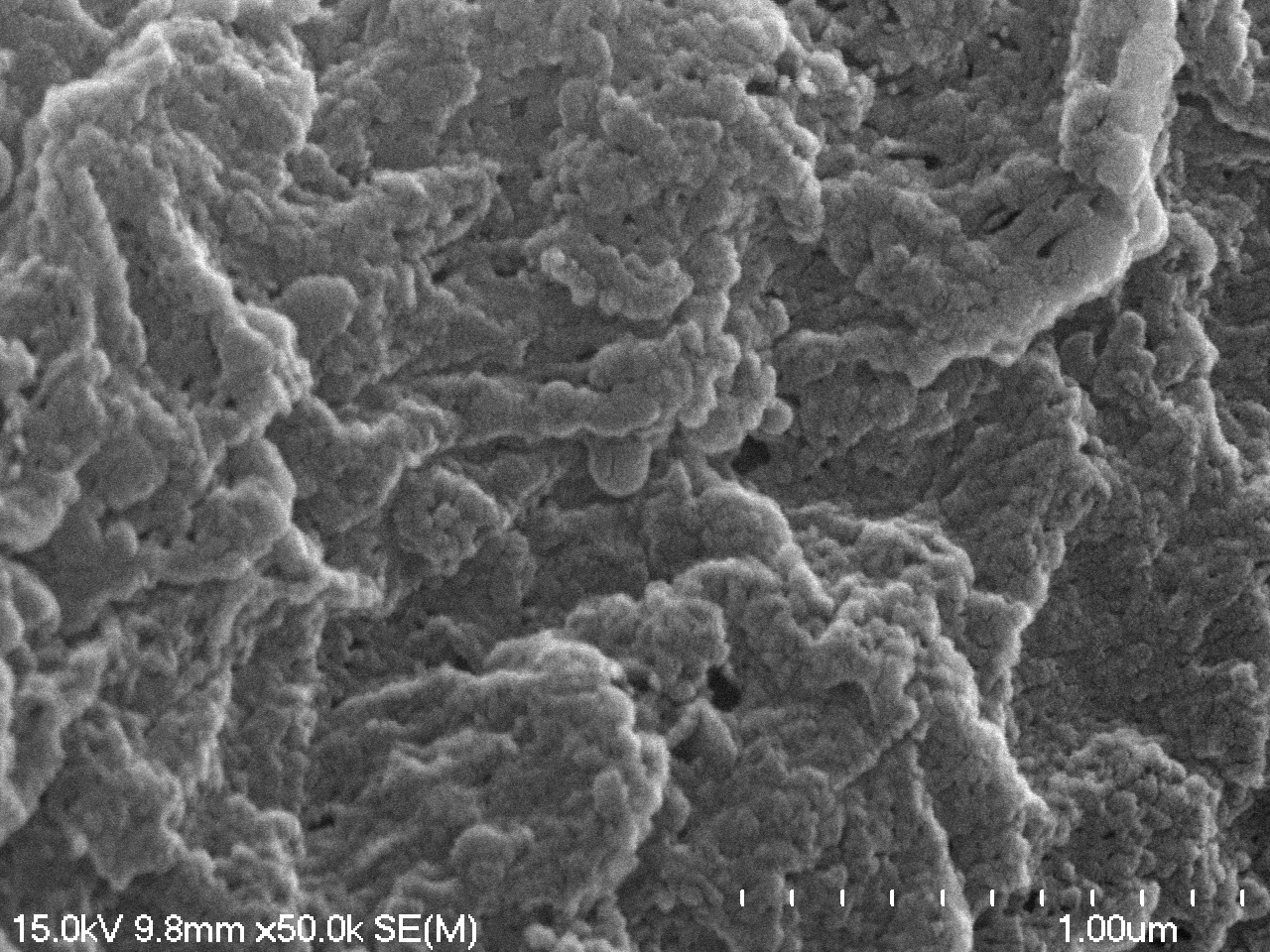

### 07-05-2023 alligator collagen 400x 02.TIF

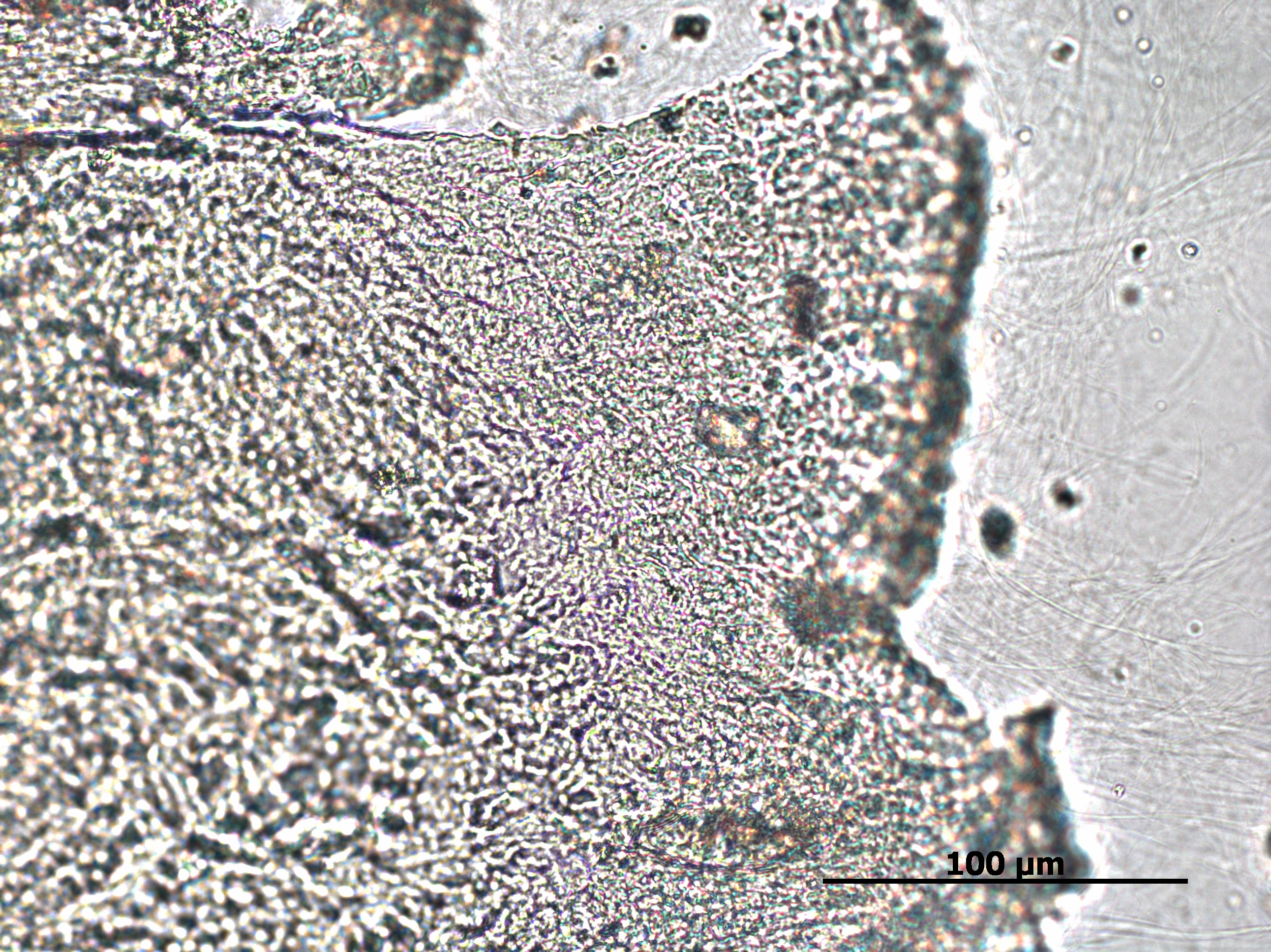

### 07-06-2023 alligator collagen_m04.tif

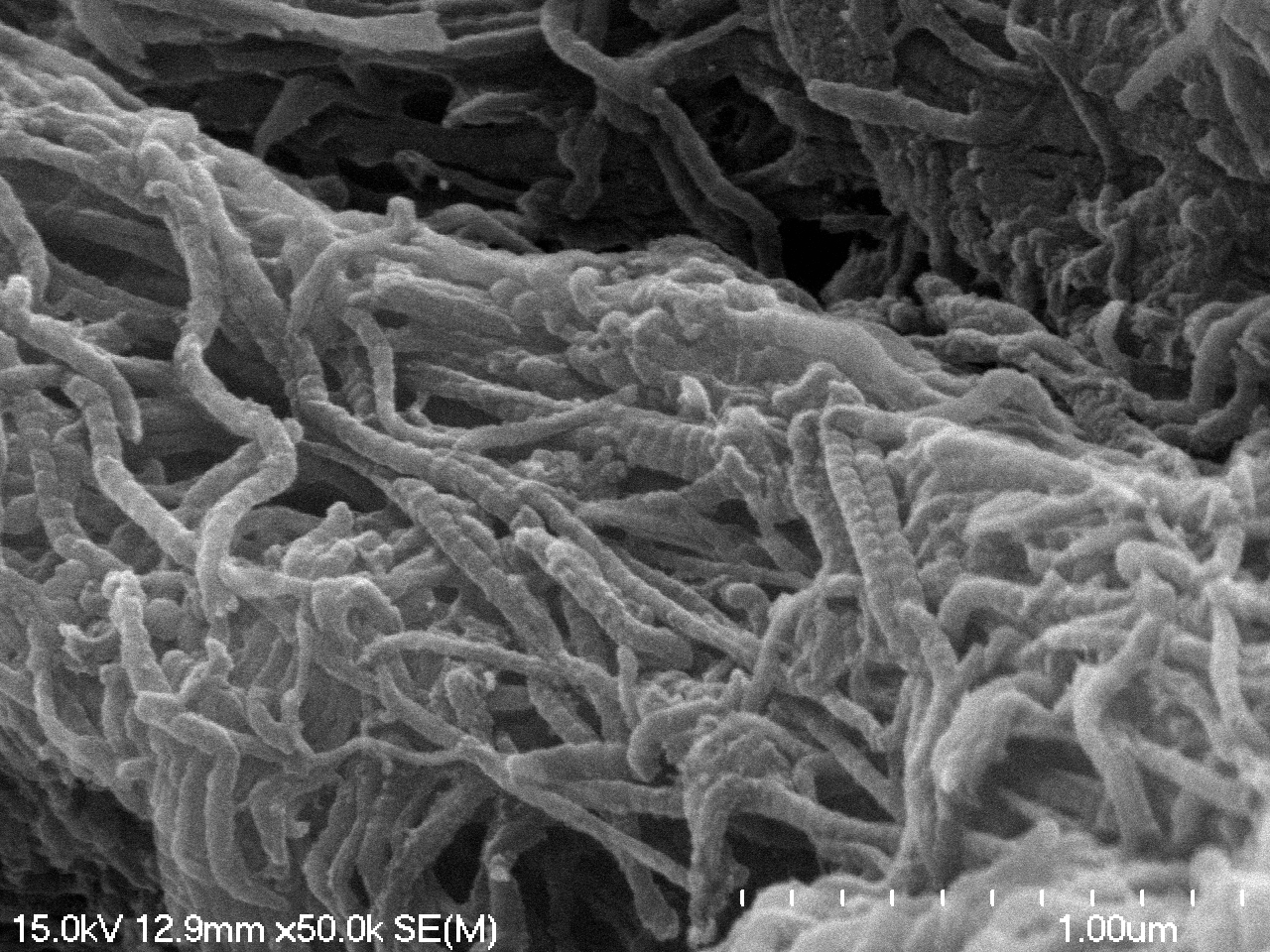

### 07-06-2023 alligator collagen_m07.tif

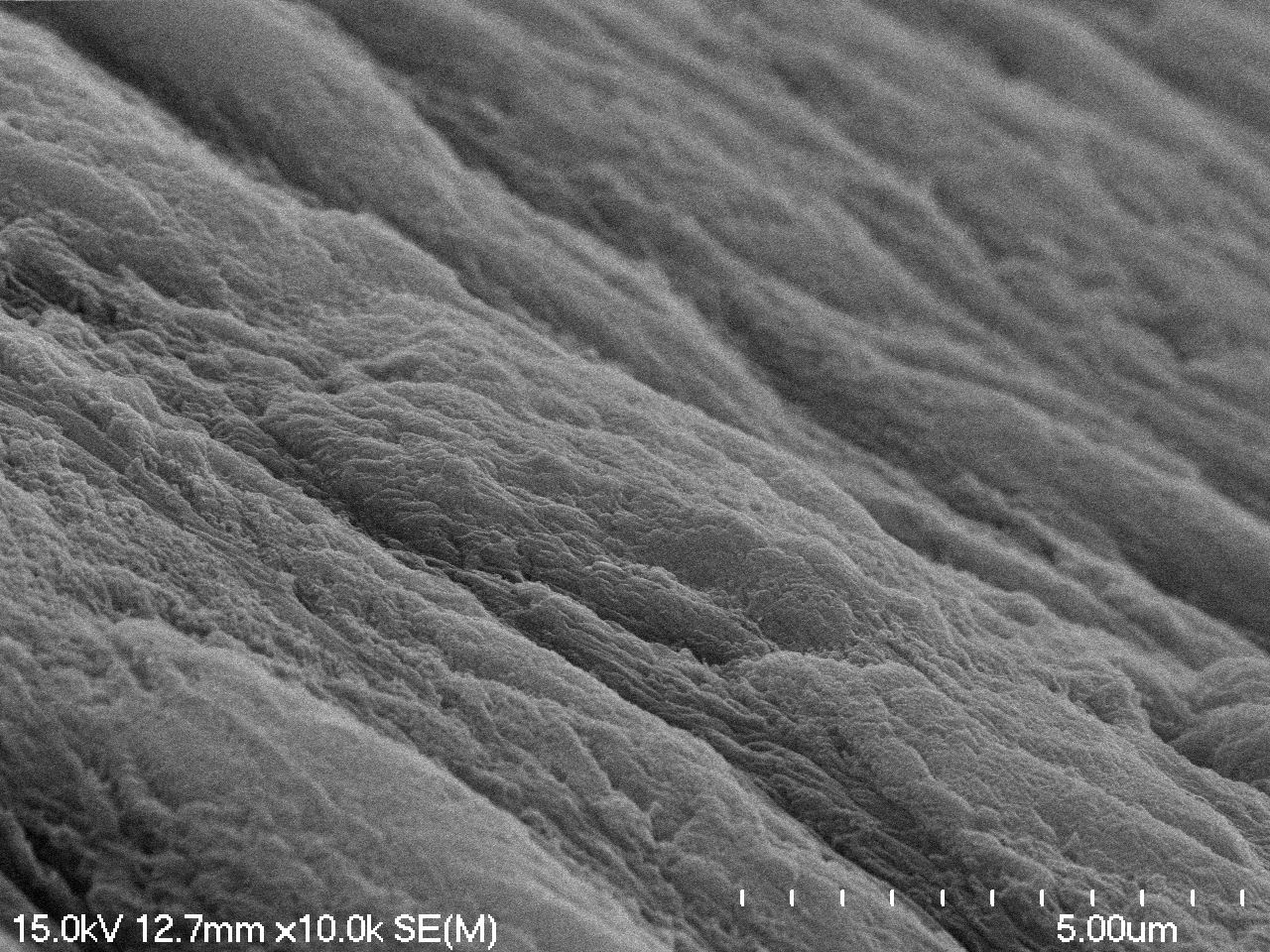

### 07-06-2023 ostrich collagen_m05.tif

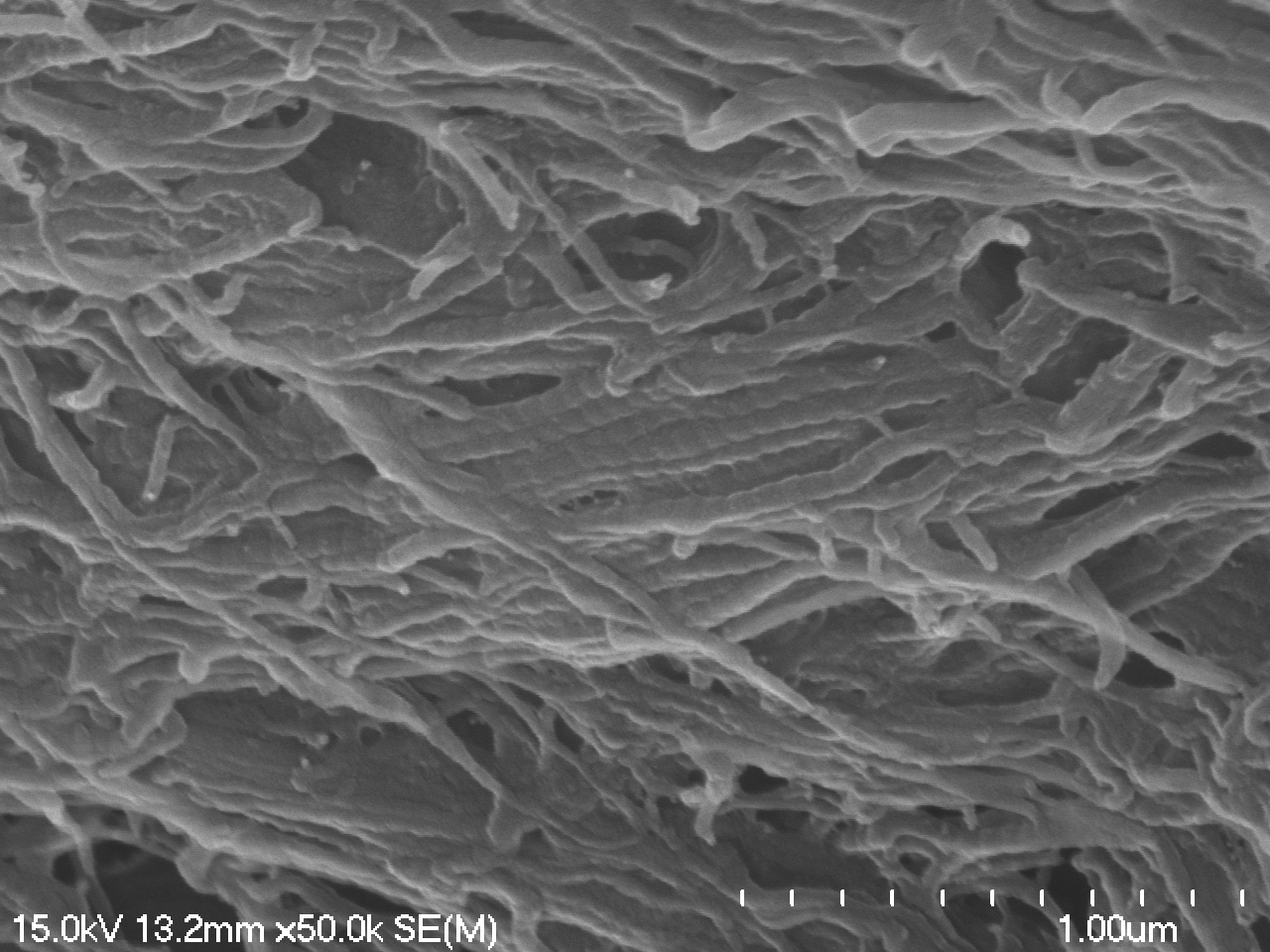

### 07-06-2023 ostrich collagen_m11.tif

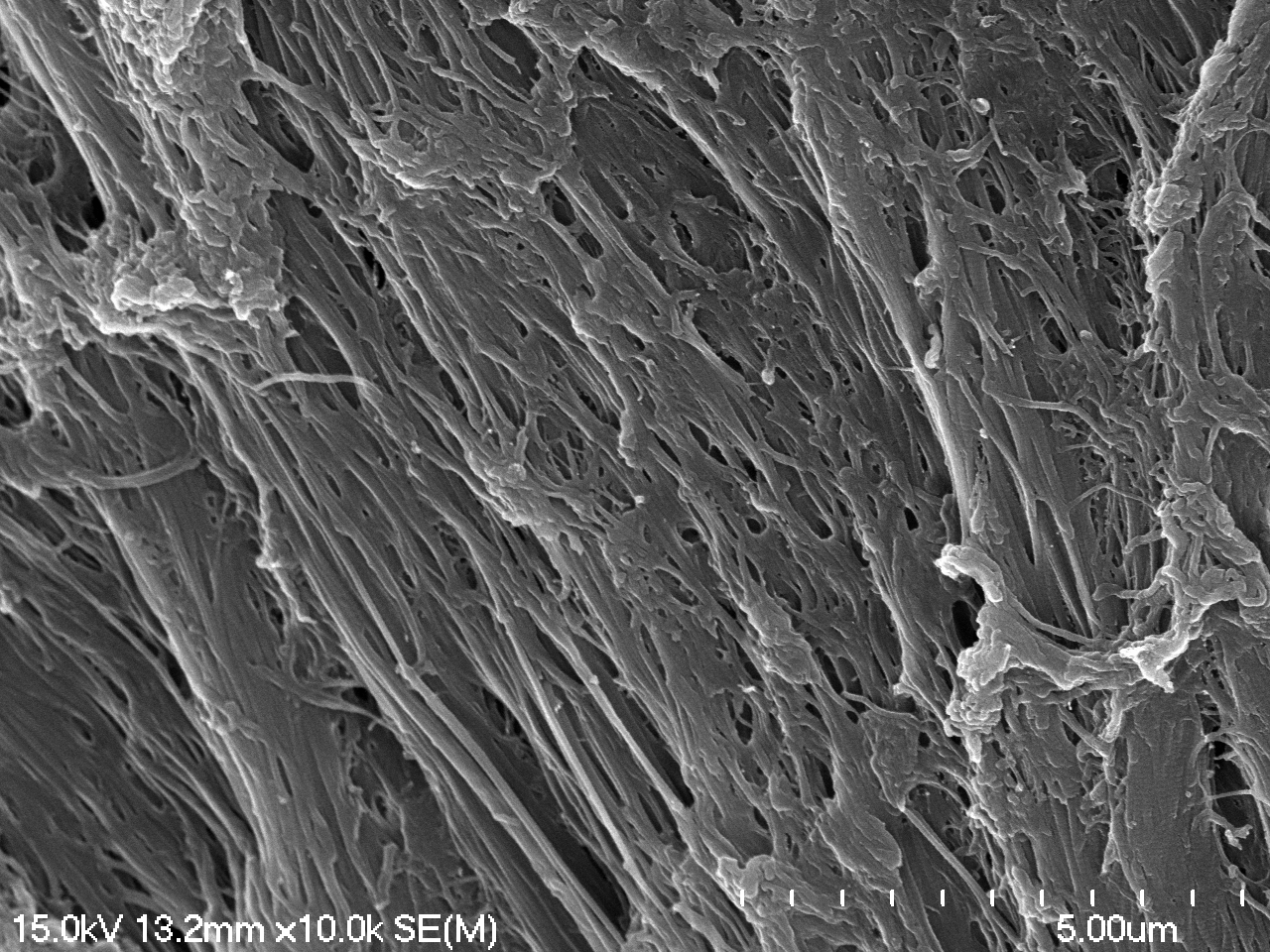

### 11-05-2023 A. mississippiensis vessels 400x 01.TIF

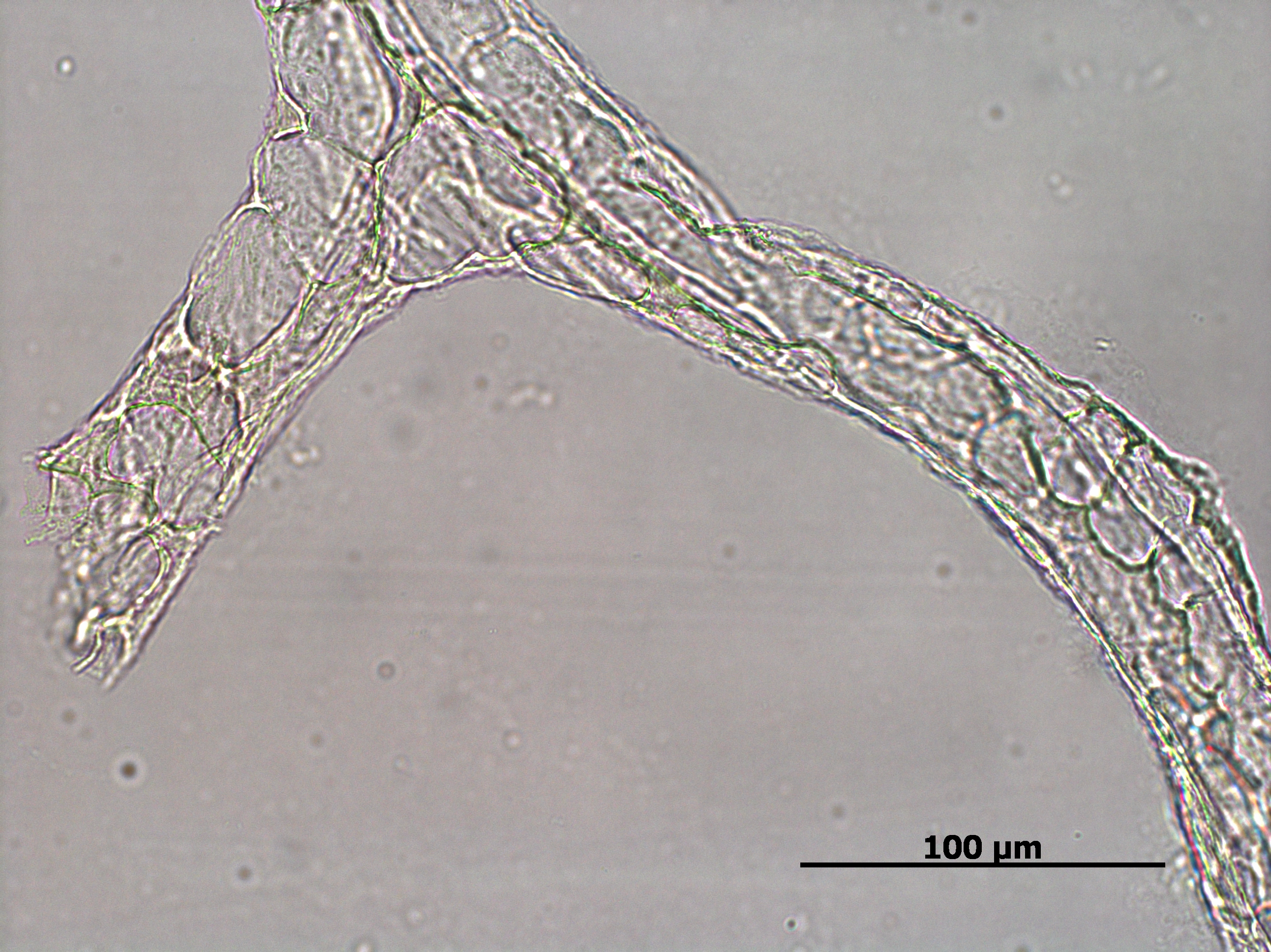

### 11-05-2023 B. taurus vessels 400x 02.TIF

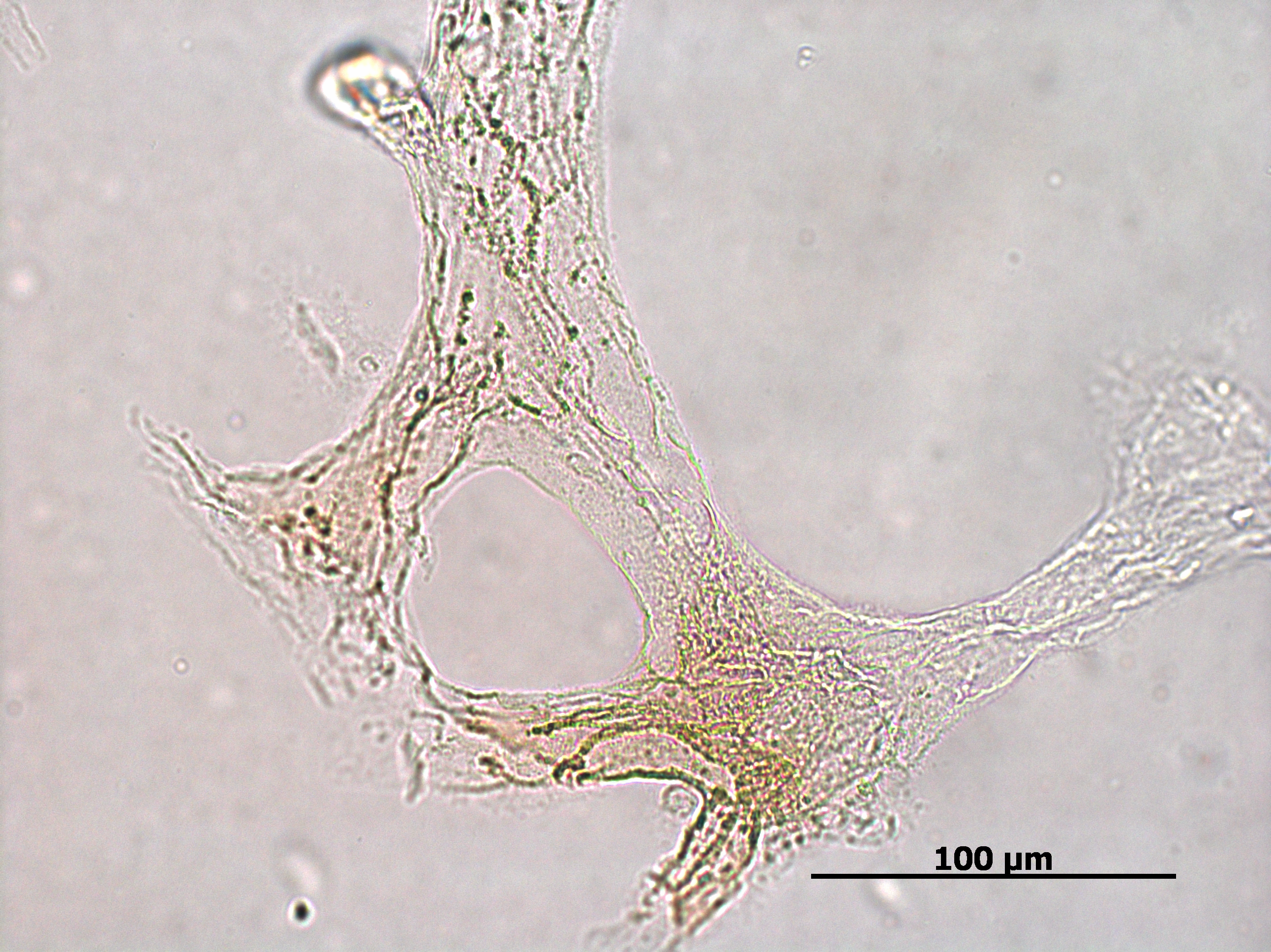

### 11-07-2023 alligator collagen_m07.tif

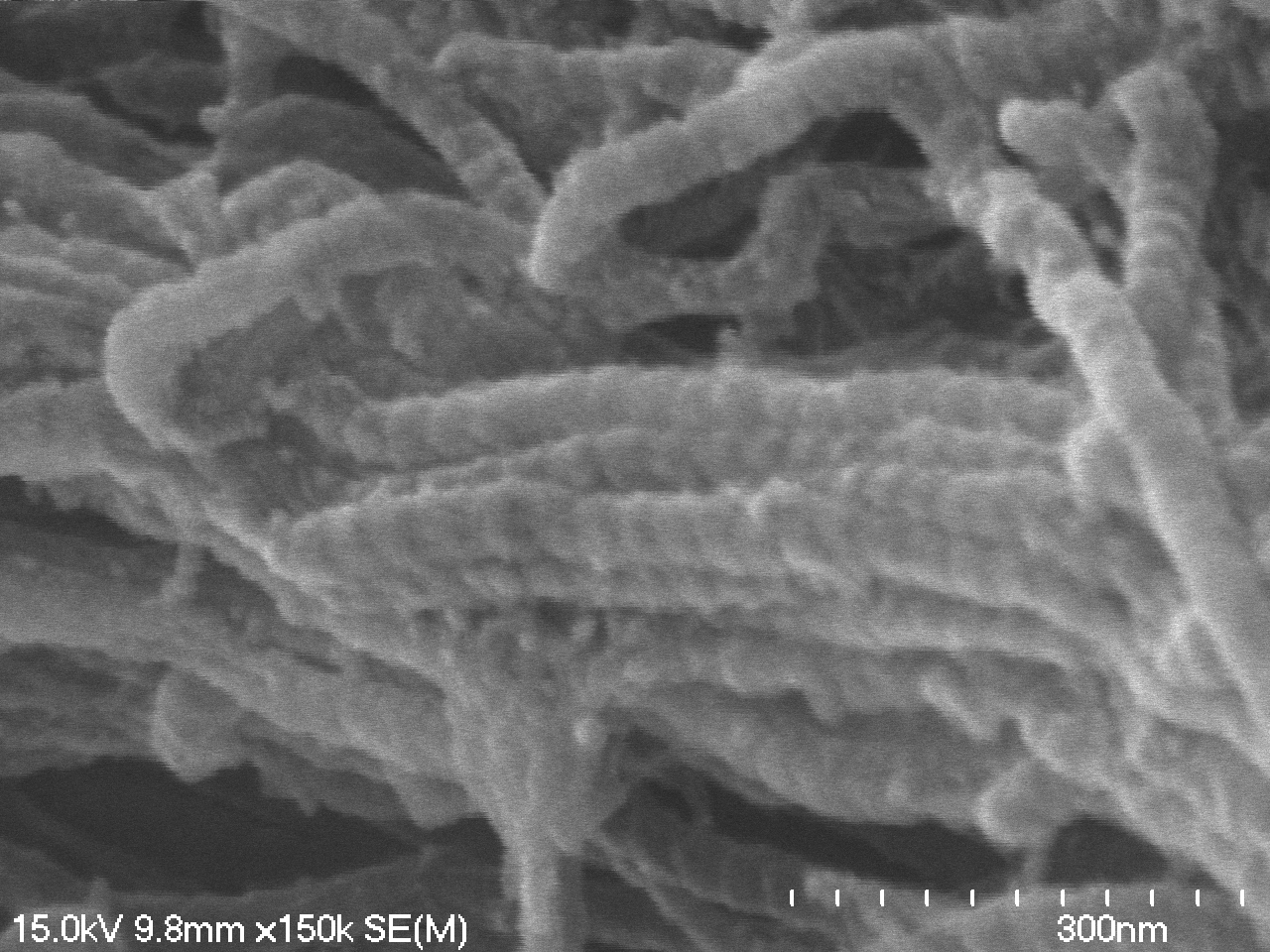

### 11-07-2023 alligator vessels_m05.tif

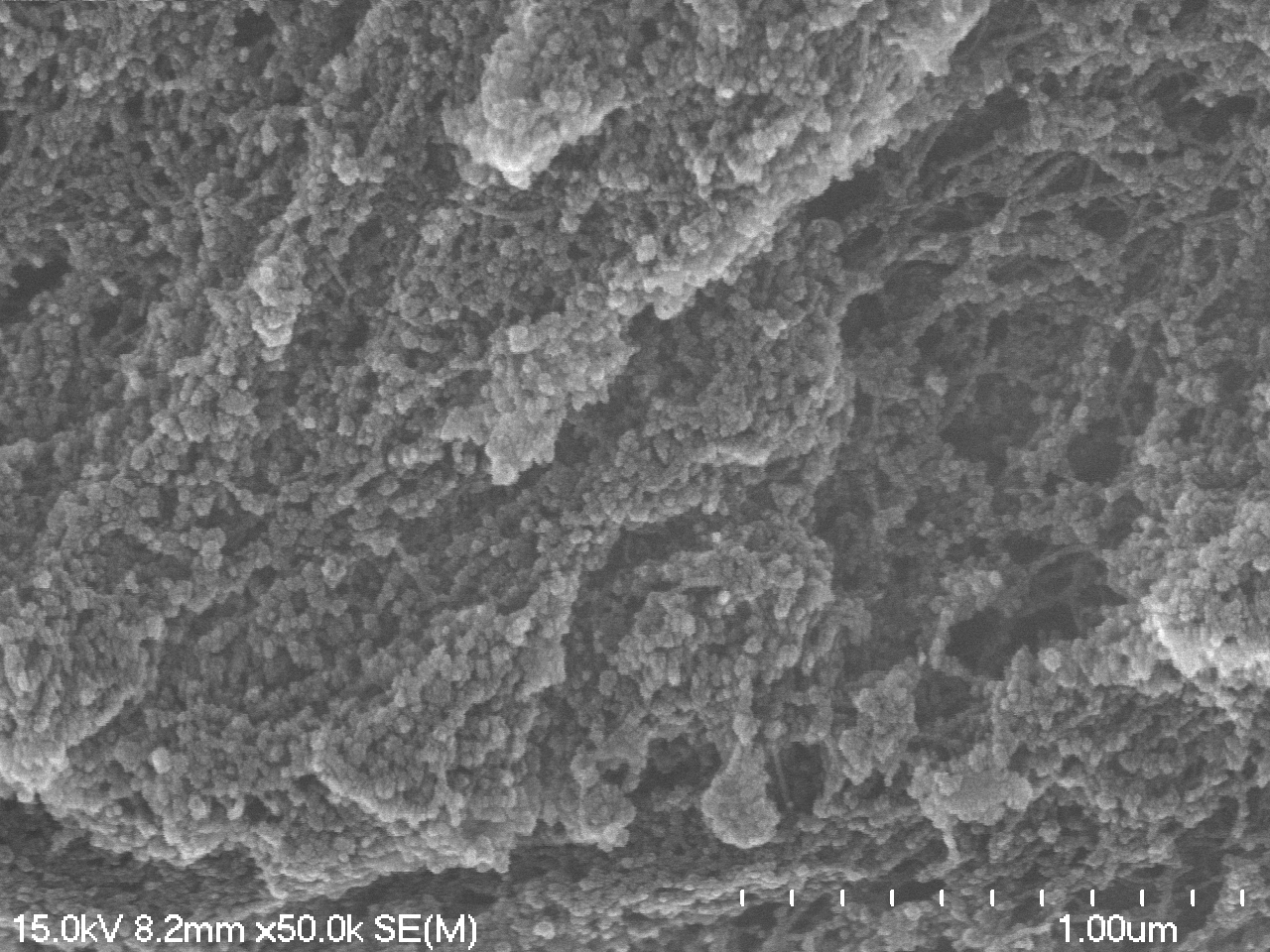

### 11-07-2023 alligator vessels_m06.tif

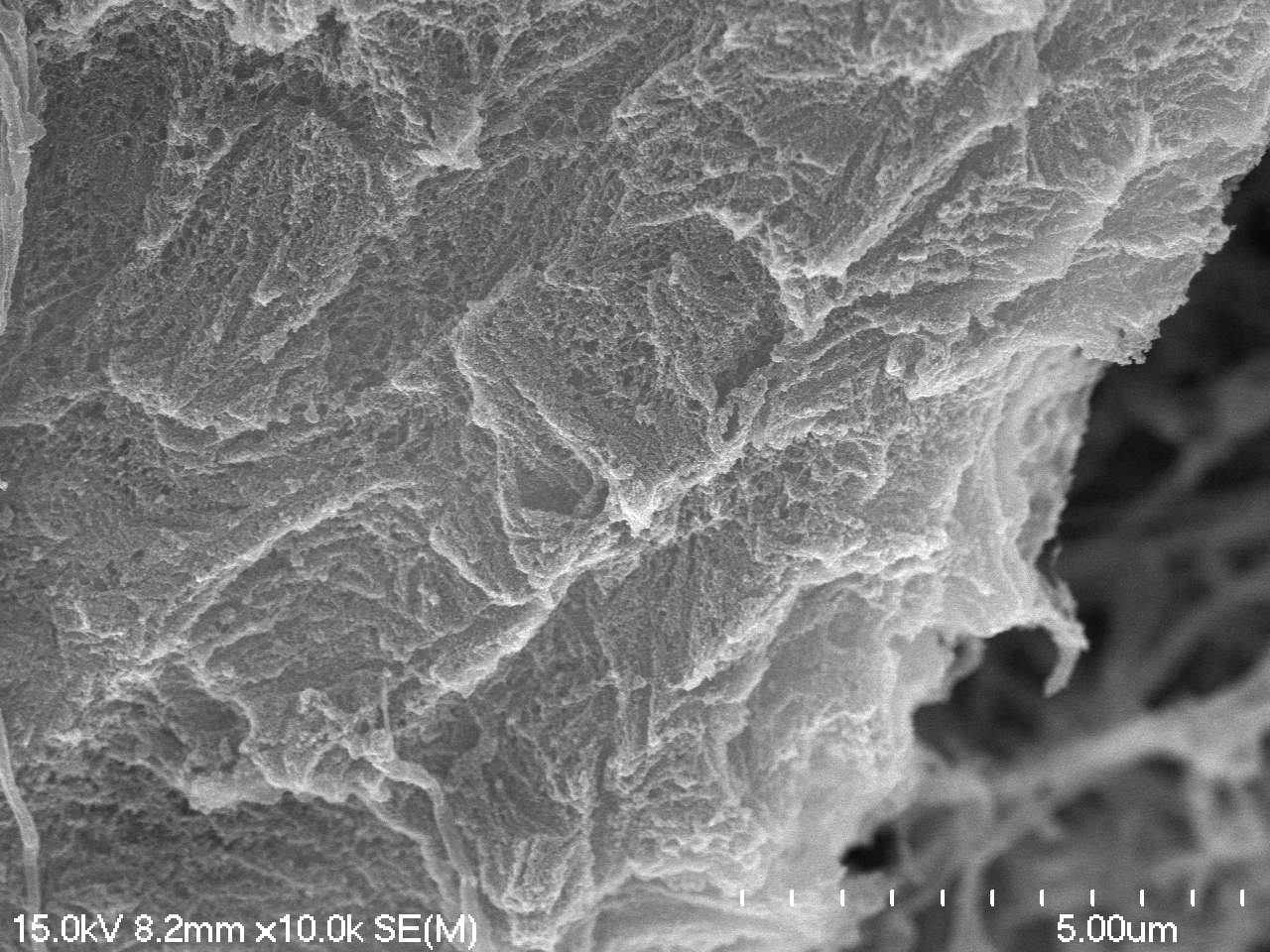

### 11-07-2023 alligator vessels_m08.tif

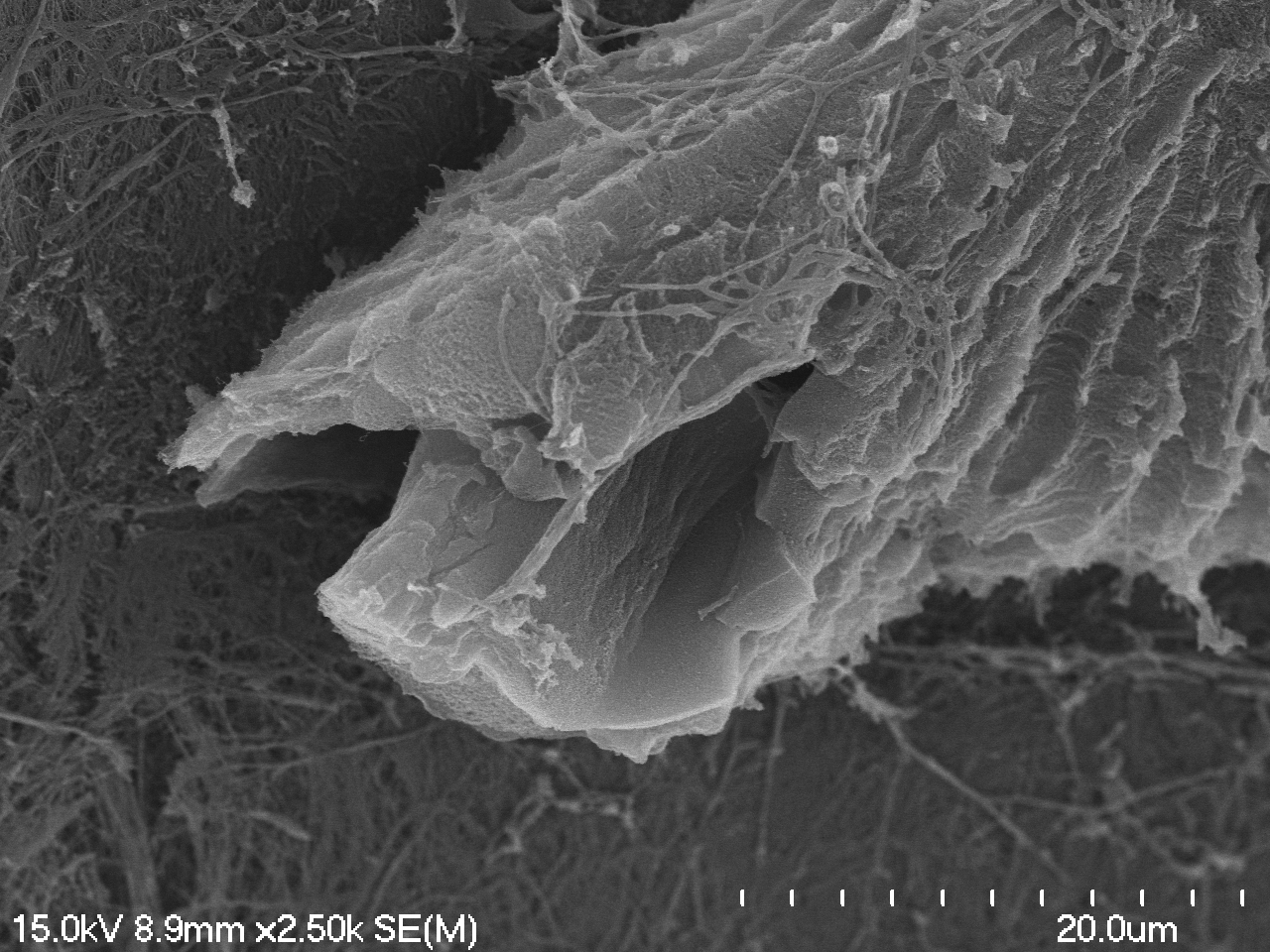

### 11-07-2023 cow collagen_m04_m02.tif

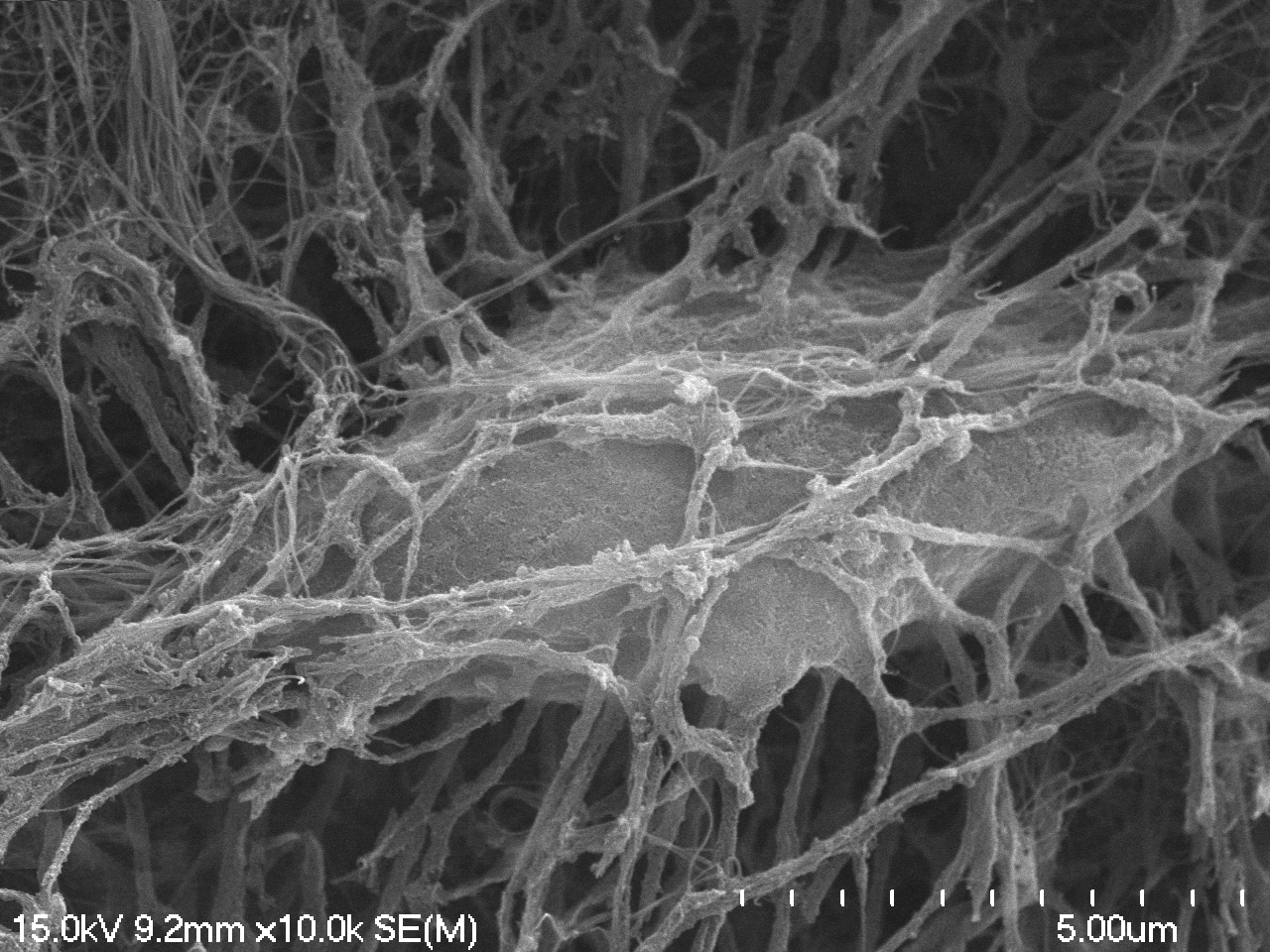

### 11-07-2023 ostrich collagen_m05.tif

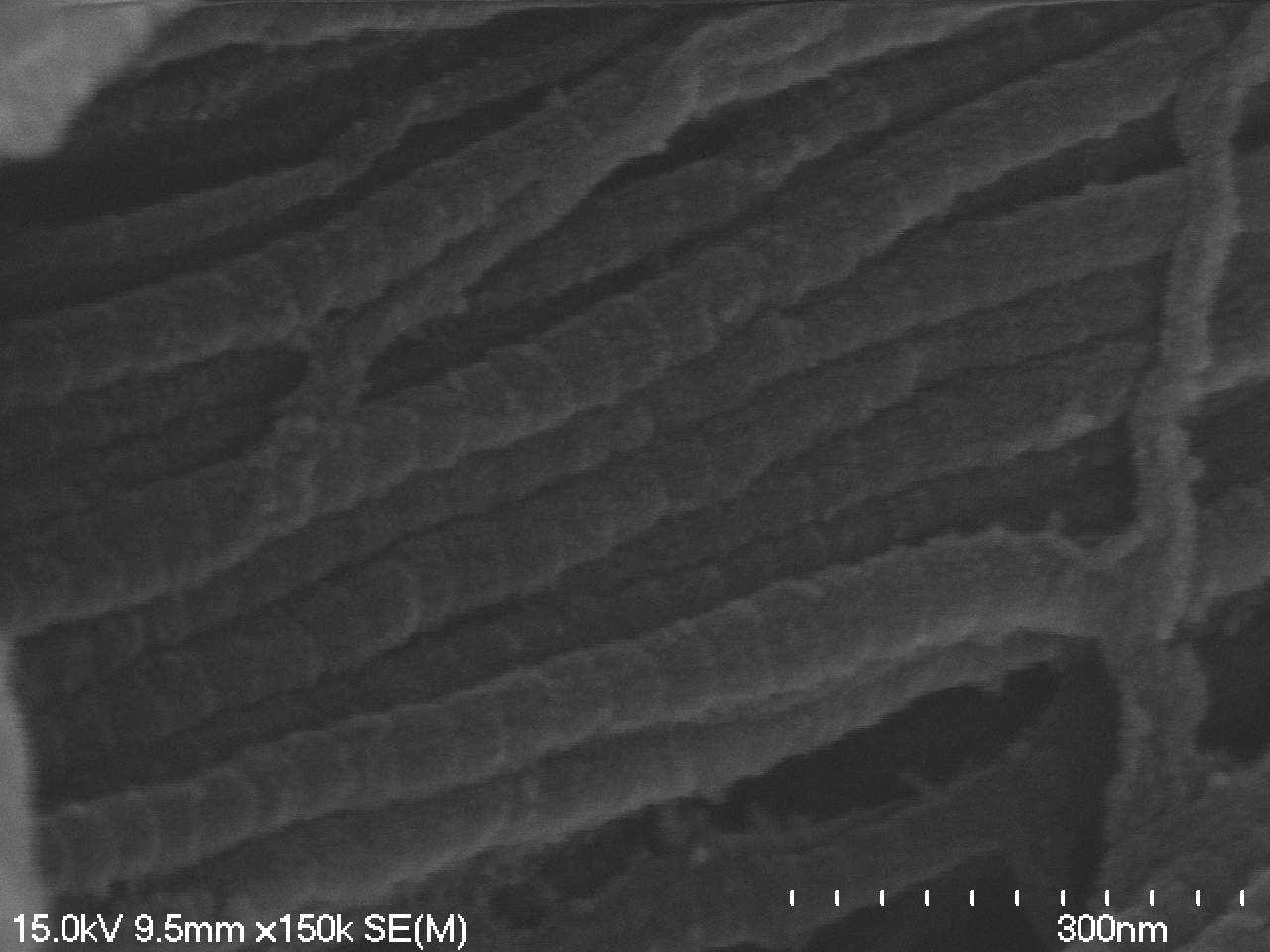

### 11-07-2023 ostrich vessels_m01.tif

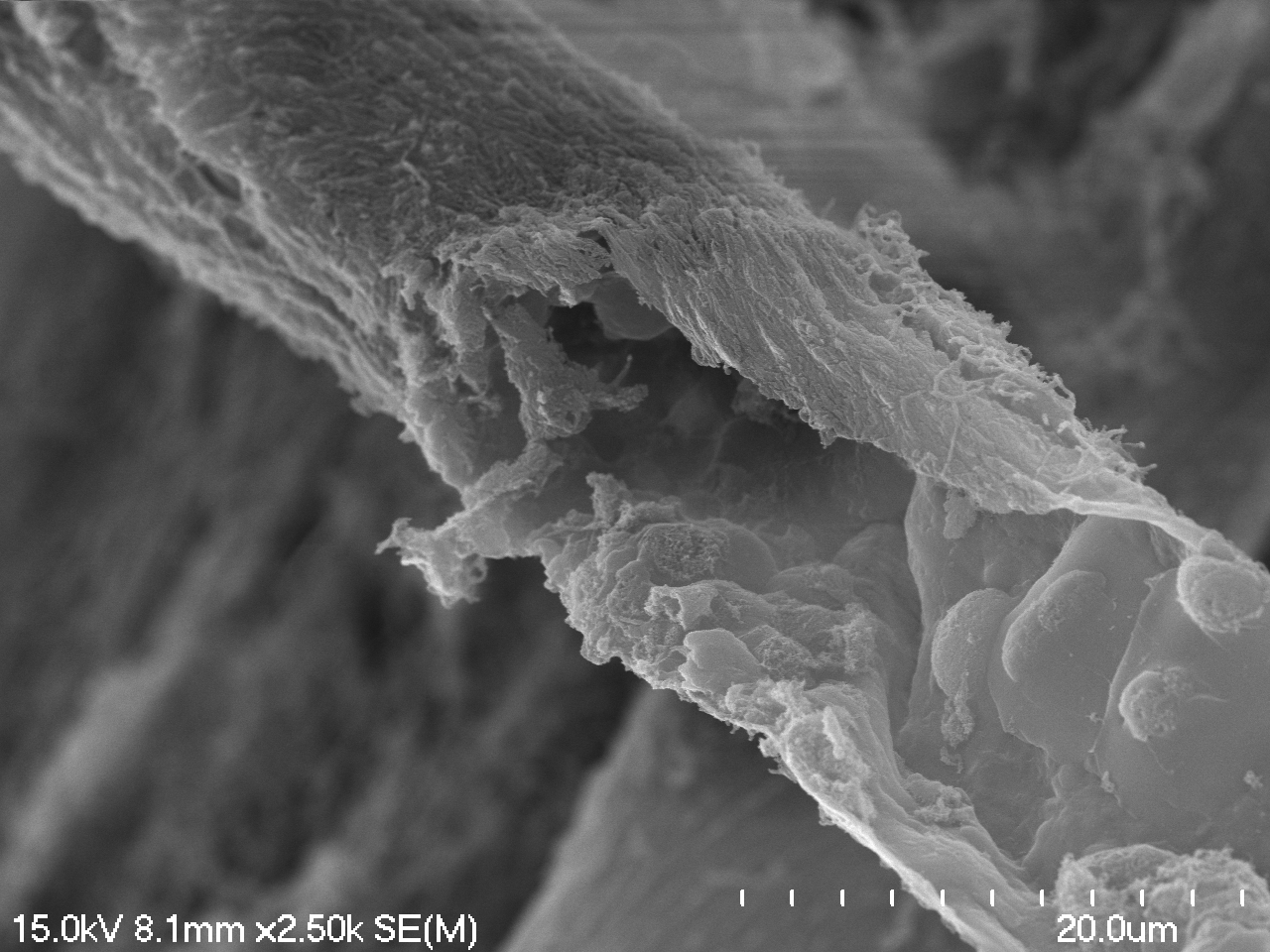

### 11-07-2023 ostrich vessels_m02.tif

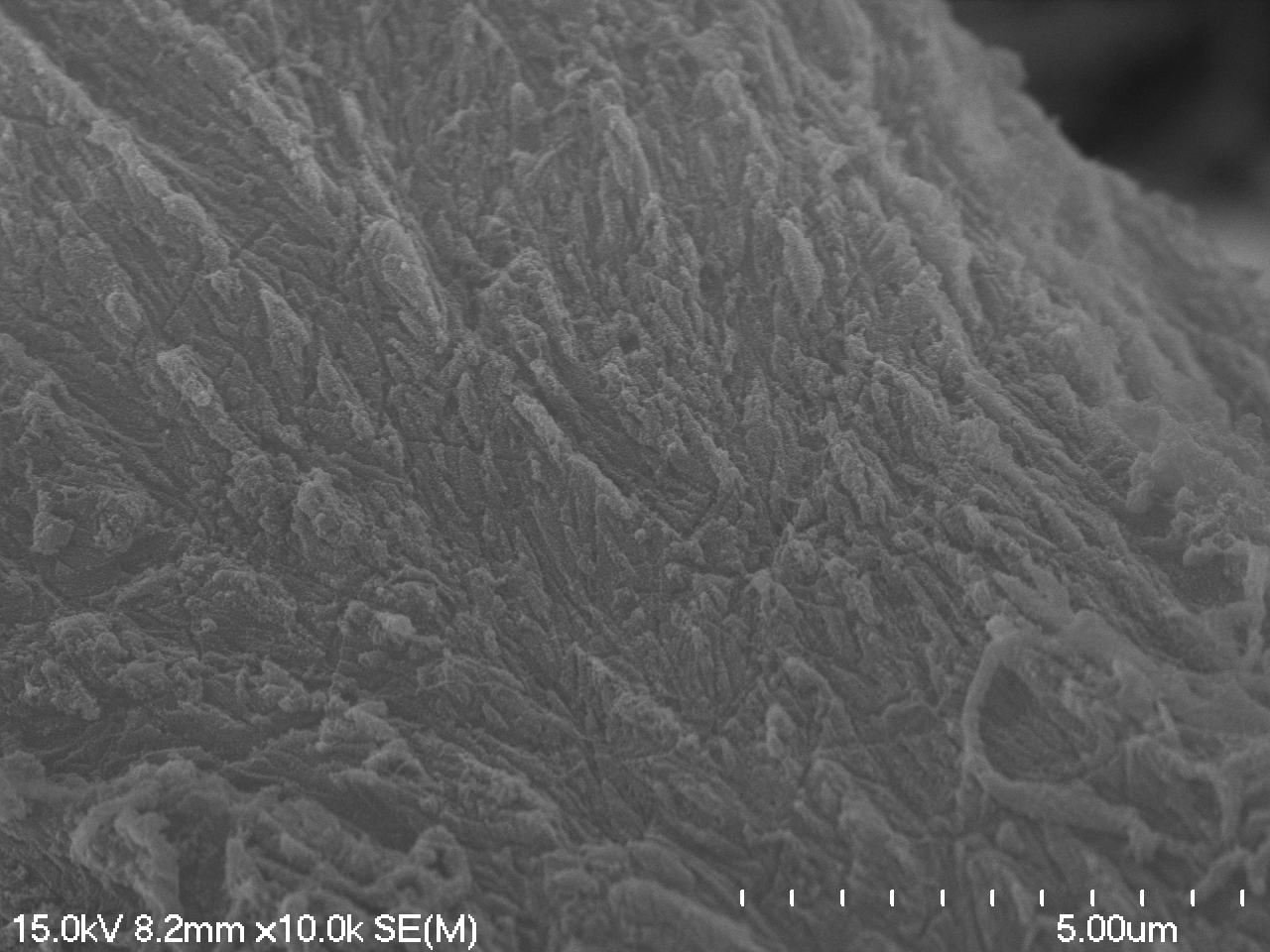

### 11-07-2023 ostrich vessels_m03.tif

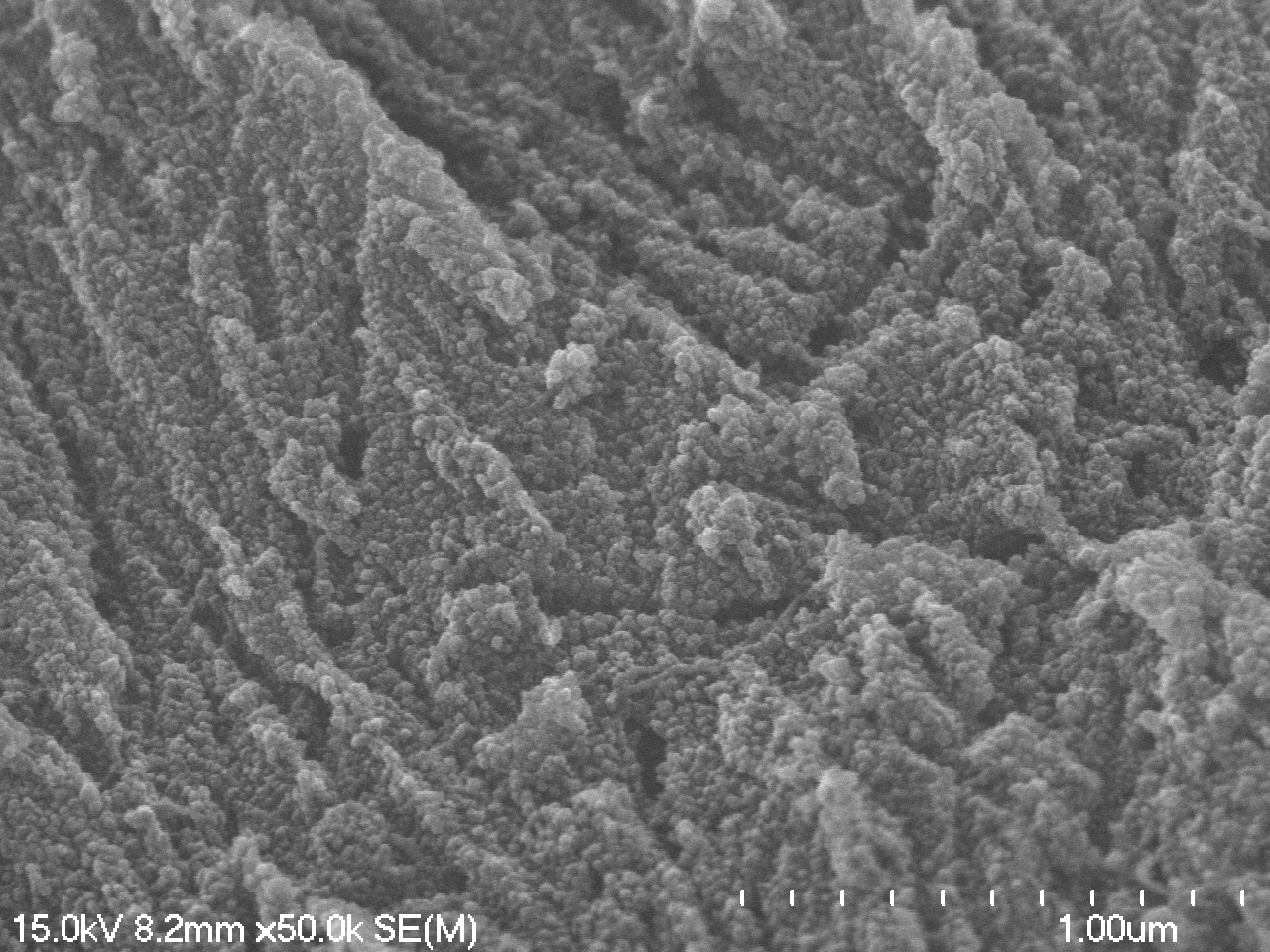

### 11-07-2023 steppe bison radius collagen_m03_m07.tif

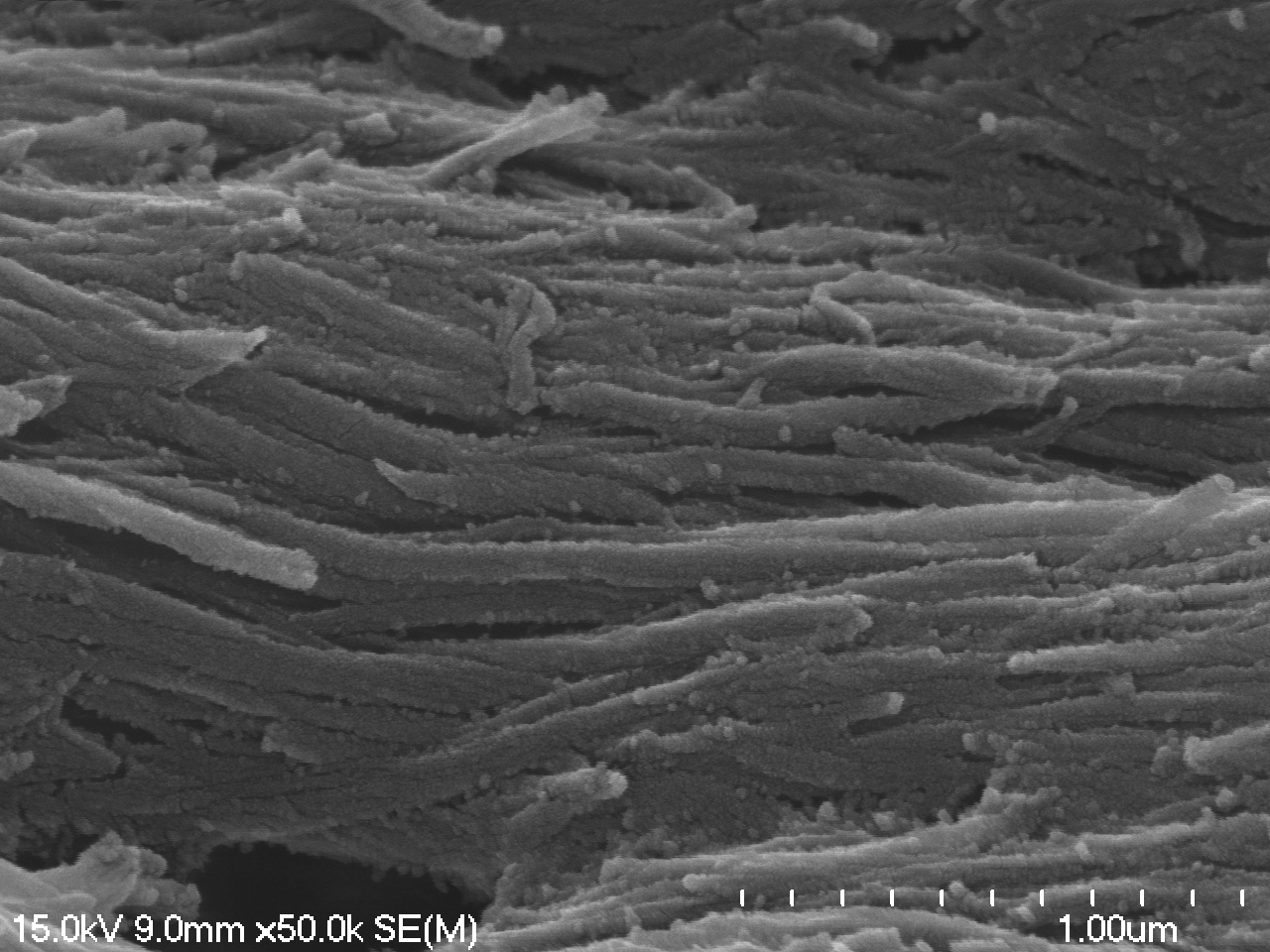

### 11-07-2023 steppe bison radius collagen_m06_m02.tif

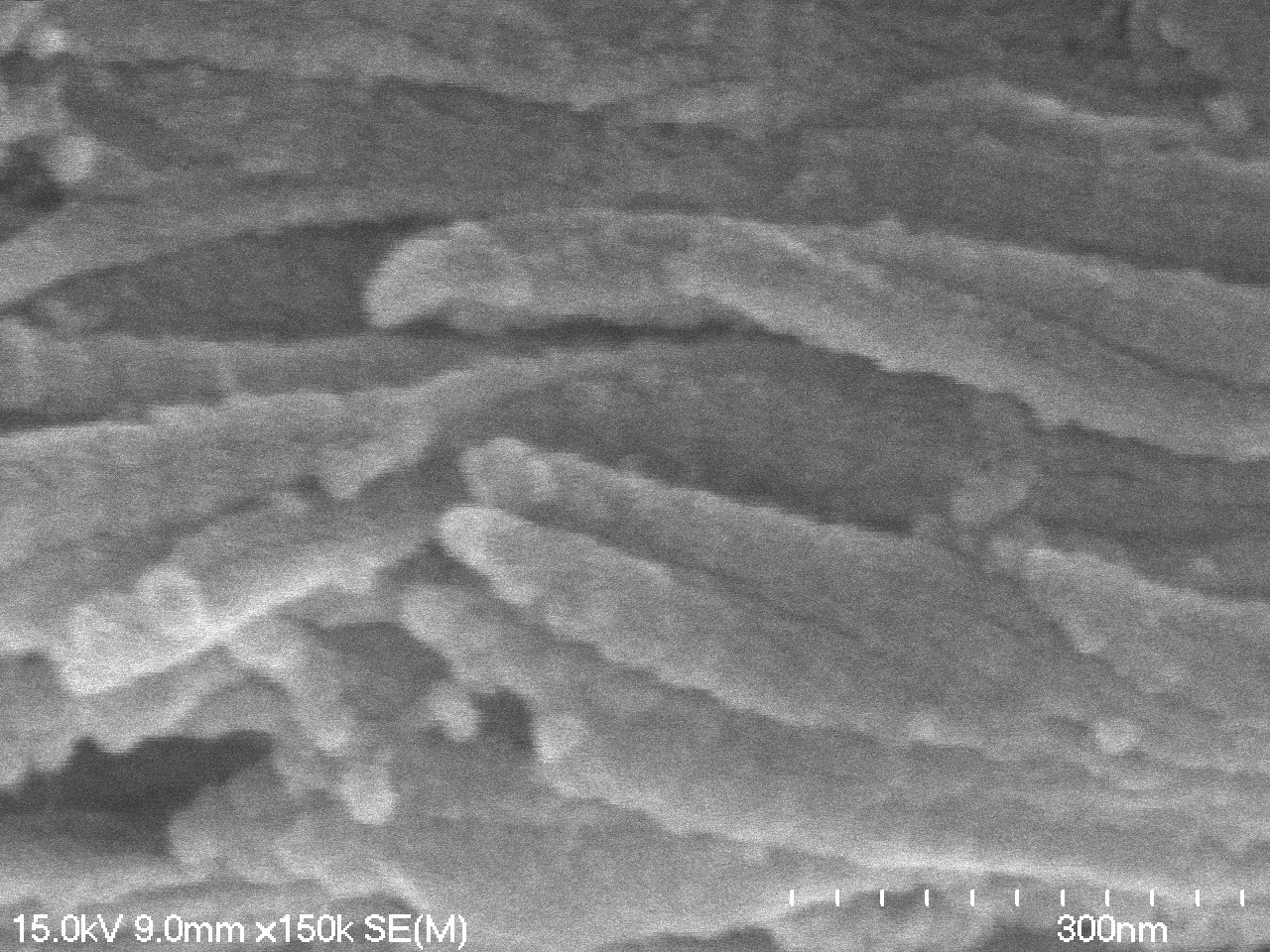

### 11-07-2023 steppe bison tibia vessels_m06.tif

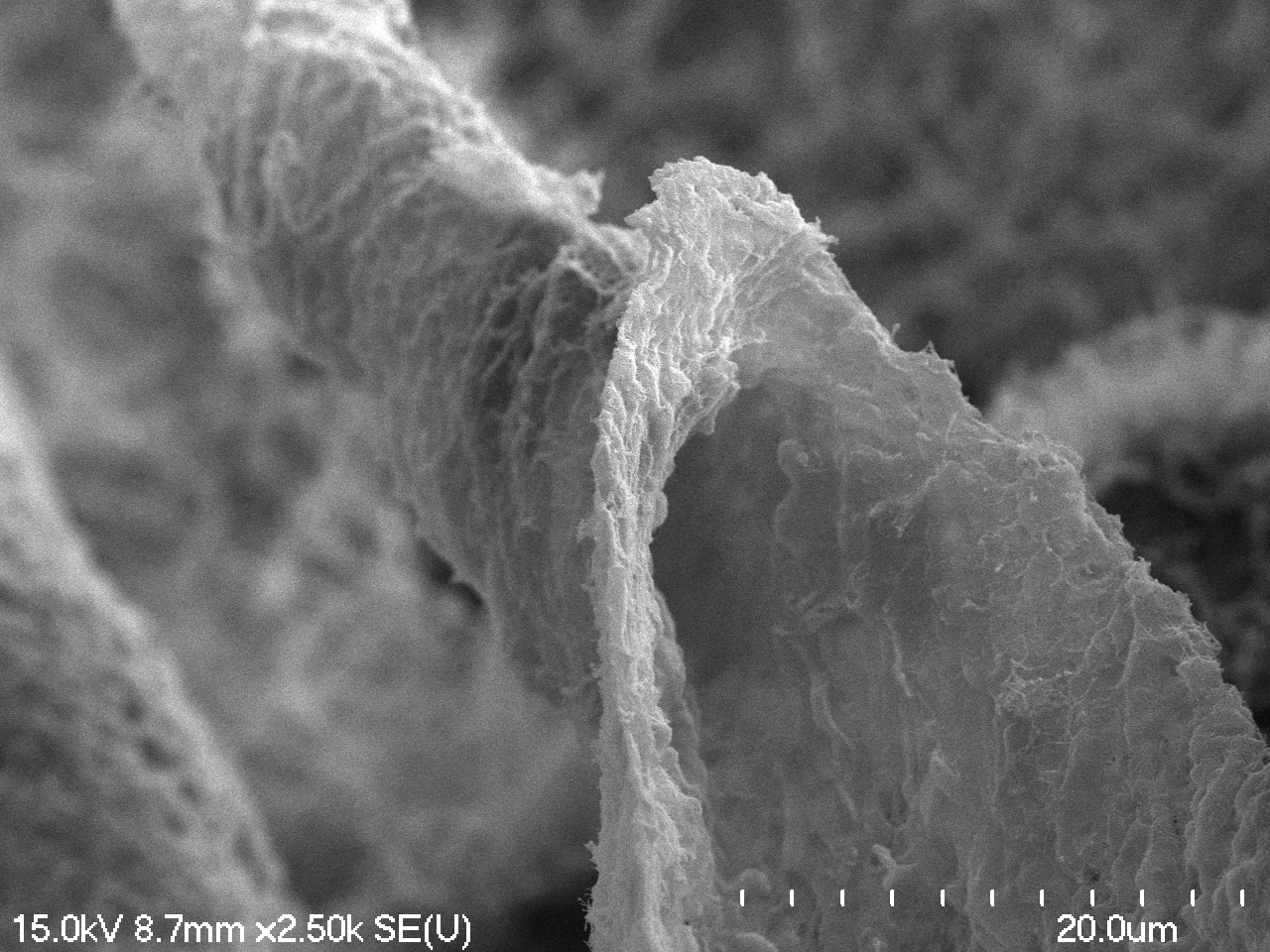

### 11-07-2023 steppe bison tibia vessels_m09.tif

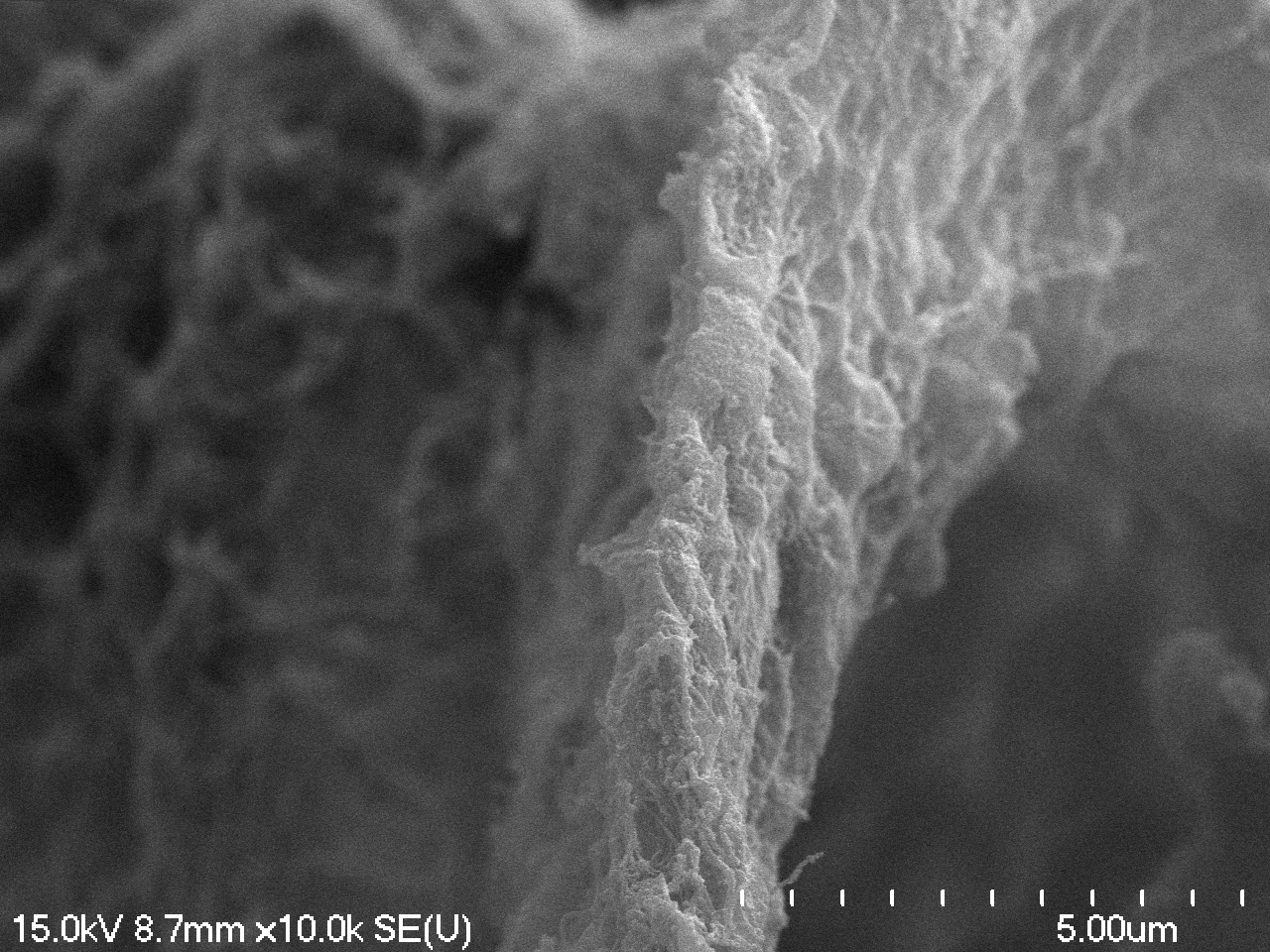

### 11-18-2023 soft tissue steppe bison tibia_m01.tif

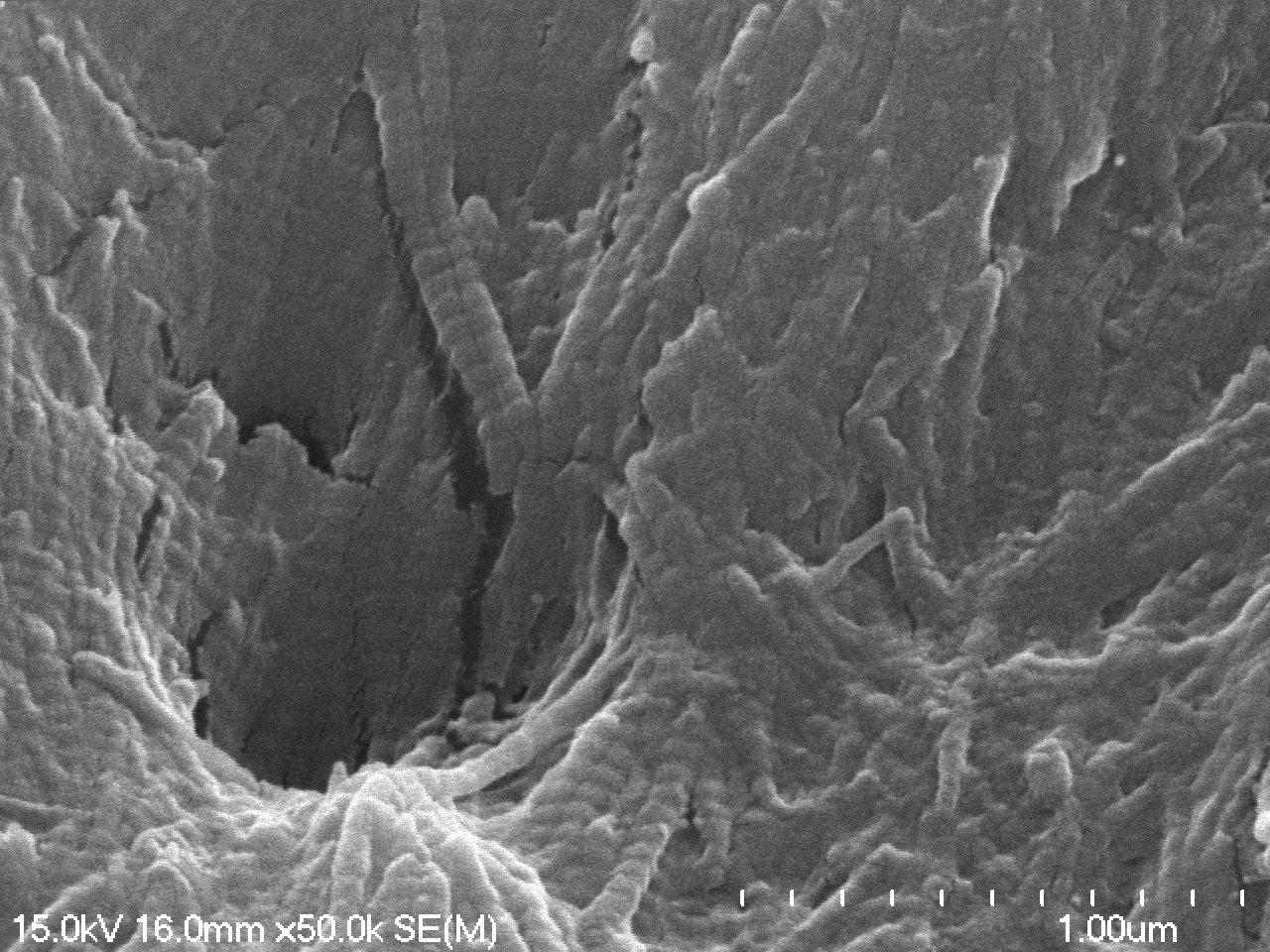

### 11-18-2023 soft tissue steppe bison tibia_m02.tif

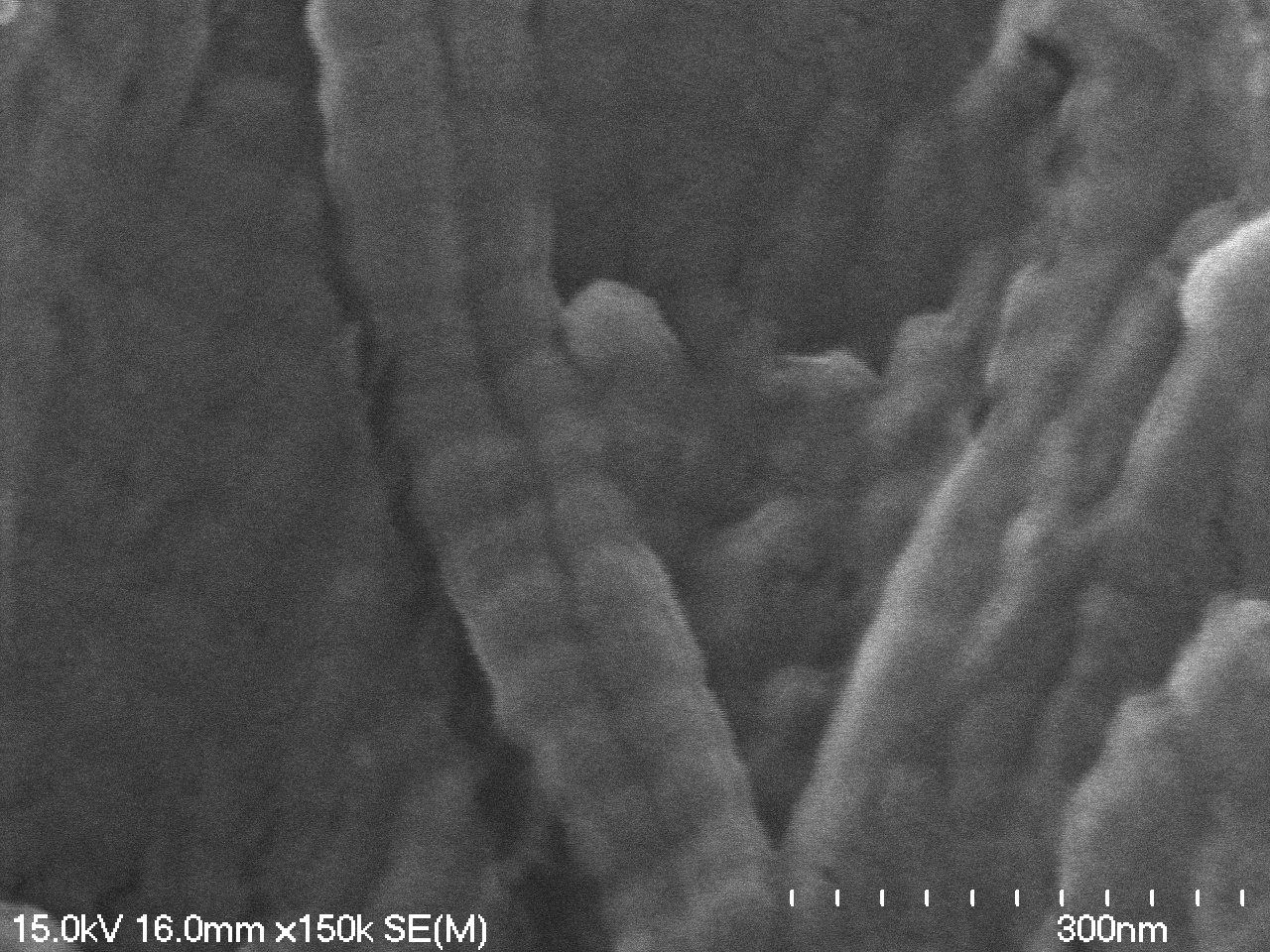

### 11-18-2023 soft tissue steppe bison tibia_m04.tif

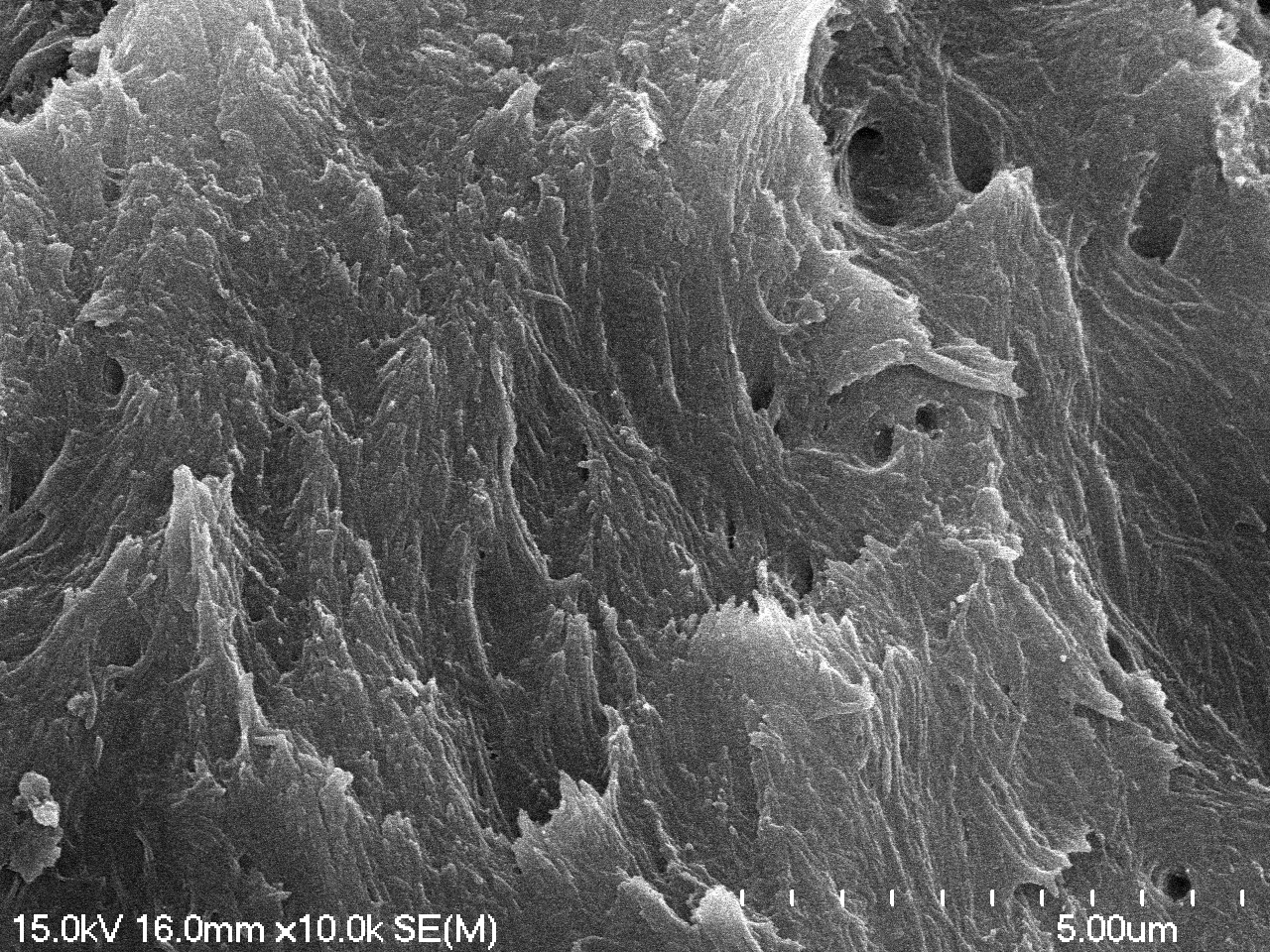

### S_m03.tif

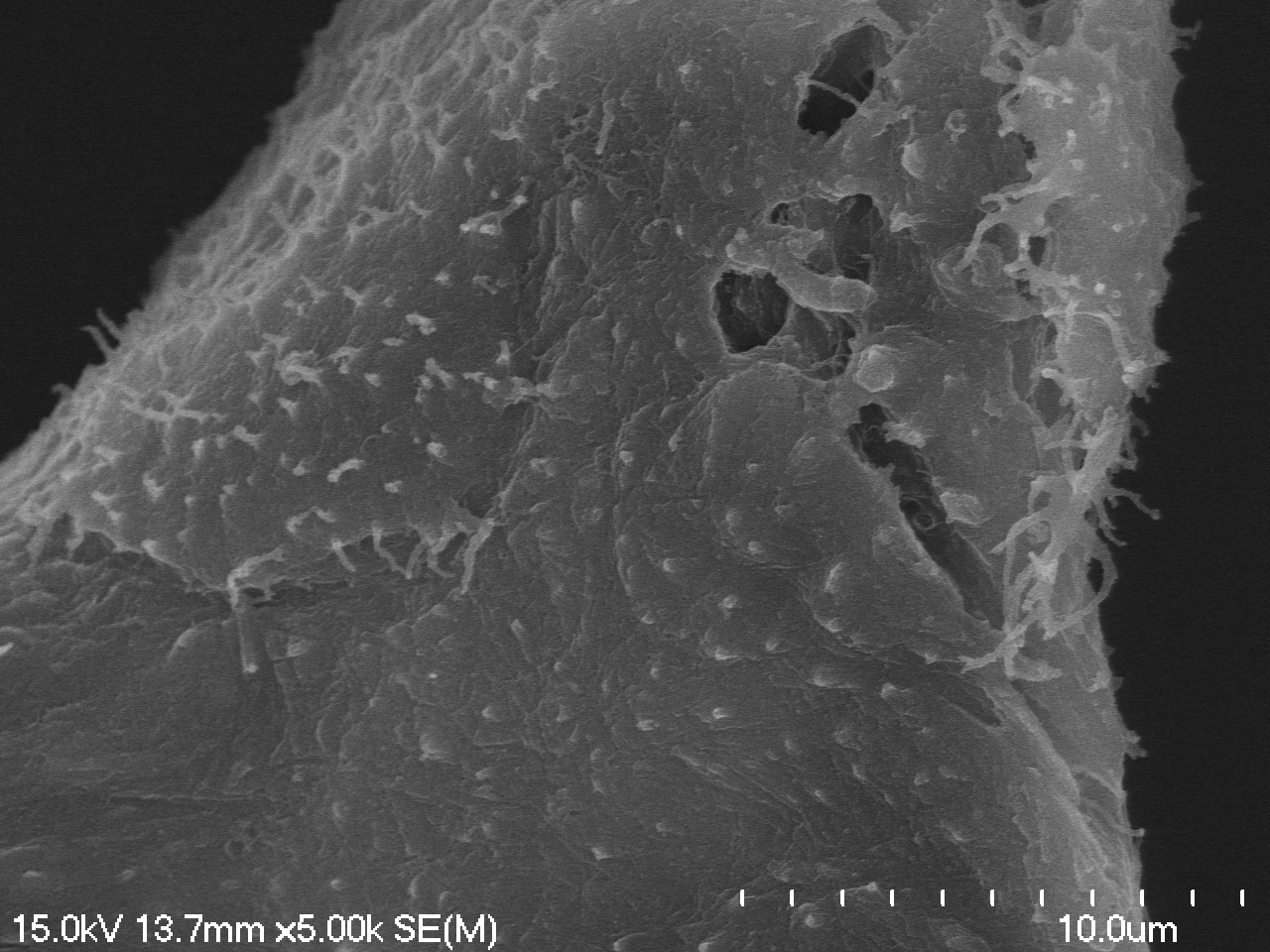
